# Estimating the power of sequence covariation for detecting conserved RNA structure

**DOI:** 10.1101/789404

**Authors:** Elena Rivas, Jody Clements, Sean R. Eddy

**Affiliations:** Department of Molecular and Cellular Biology, Harvard University, Cambridge, Massachusetts, USA; Janelia Research Campus, Howard Hughes Medical Institute, Ashburn, Virginia, USA; Howard Hughes Medical Institute, Harvard University, Cambridge, Massachusetts 02138, USA; John A. Paulson School of Engineering and Applied Sciences, Harvard University, Cambridge, Massachusetts 02138, USA

## Abstract

Pairwise sequence covariations are a signal of conserved RNA secondary structure. We describe a method for distinguishing when lack of covariation signal can be taken as evidence against a conserved RNA structure, as opposed to when a sequence alignment merely has insufficient variation to detect covariations. We find that alignments for several long noncoding RNAs previously shown to lack covariation support do have adequate covariation detection power, providing additional evidence against their proposed conserved structures.

---

Comparative analyses of pairwise covariations in RNA sequence alignments have a successful history in consensus RNA secondary structure prediction, where the existence of a conserved structure is assumed *a priori* ^1–7^. A statistically different question arises when covariation analysis is used to infer whether or not a genomic region is constrained by an evolutionarily conserved RNA secondary structure, as evidence for a structure-dependent function. For example, this question arises in analysis of long noncoding RNAs (lncRNAs) of uncertain mechanism. For this, one wants to determine if the covariation signal is distinguishable from a null hypothesis of primary sequence conservation patterns alone.

We previously introduced R-scape (RNA structural covariation above phylogenetic expectation), a method for evaluating the statistical significance of covariation support for conserved RNA basepairs ^8^. R-scape analyses found that the covariation evidence for proposed conserved structures of several long noncoding RNAs including HOTAIR^9^, SRA^10^, and the RepA region of Xist^11;12^ is not statistically significant^8^.

Lack of significant covariation signal does not necessarily mean there is no conserved RNA structure. An alignment could merely have too little sequence variation to detect significant covariation (Figure 1a). To know when an alignment has sufficient variation, we want to estimate the statistical power (the expected sensitivity) of detecting significant covariations. In a “low-power” alignment, covariation analysis is inconclusive because a conserved RNA secondary structure could be present without inducing sufficient covariation signal. In a high-power alignment, observing no supporting covariations does provide evidence against a conserved structure.

**Figure 1:**
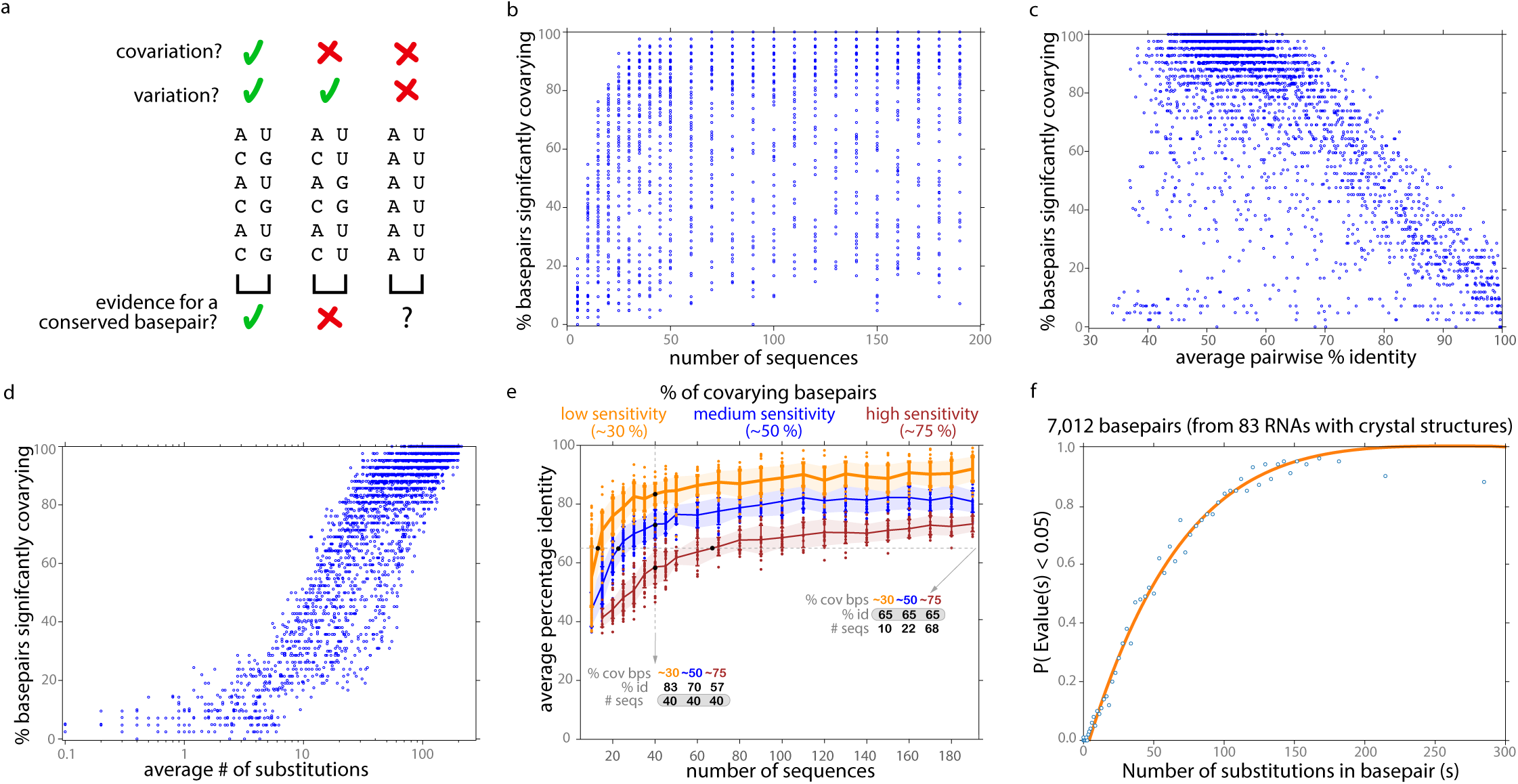
(a) Three different patterns for two alignment columns proposed to form a consensus basepair. **(a, left)** The two columns have variation and covariation (mutual information is 1 bit). This pattern is consistent with a basepair conserved throughout evolution. **(a, middle)** The two columns have variation but not covariation (mutual information is 0.0). These two positions are unlikely to form a basepair. **(a, right)** Two columns with no covariation and no variation. This pattern is consistent with a A-U basepair, but there is no evolutionary evidence for it. **(b,c,d) Scatter plots of power (% sensitivity) for detecting basepairs in simulated alignments.** Each point represents the fraction of 42 consensus basepairs in a simulated Cobalamin alignment detected with an R-scape E-value *<* 0.05, as a function of sequence number **(b)**, average pairwise % identity **(c)**, or inferred number of substitutions in two columns *s*_*ij*_ **(d). (e)** The same simulated alignments binned by low (yellow ∼ 27 − 32%), medium (blue ∼ 47 − 52%) or high (red ∼ 74 − 76%) sensitivity and scatter plotted, illustrating how detection power increases either by increasing sequence number or sequence diversity. **(f) Power of covariation as a function of the total number of substitutions.** The orange line is the fitted power(*s*) curve. Each blue dot represents the empirical data we fit to: the mean fraction of significantly covarying basepairs and mean *s*_*ij*_ in a set of 100 annotated basepairs from Rfam seed alignments, out of 7,012 total basepairs ordered by increasing number of substitutions.

Many details of an alignment affect covariation analysis, but we hypothesized that detection power should depend primarily on the total number of single residue substitutions *s*_*i,j*_ in two alignment columns *i* and *j* in a proposed consensus pair. We take the sequence phylogeny into account in inferring *s*_*i,j*_ by inferring a maximum likelihood tree, using the Fitch parsimony algorithm ^13^ to estimate a number of substitutions *s*_*i*_ at each column independently, and taking *s*_*i,j*_ = *s*_*i*_ + *s*_*j*_ (Methods).

We tested this idea using synthetic RNA alignments evolved under simulated pairwise constraints. Figure 1b-d show simulations based on a cobalamin riboswitch alignment (Rfam RF00174) of 430 sequences and 42 annotated consensus basepairs. We choose a random sequence as the root and evolve it down a sub-sampled and rescaled phylogenetic tree, using an evolutionary model that includes basepair substitutions, insertions, and deletions ^14^, to generate a synthetic alignment with a desired number of taxa and average percentage identity (Methods). We repeat this to create synthetic alignments over a wide range of sequence number and diversity. We use R-scape on each alignment to determine the number of basepairs with significant covariation support (E-value *<* 0.05). Neither the number nor the diversity of sequences in the alignment alone suffices to estimate detection power (Figure 1b,c), whereas *s*_*i,j*_ does have a good relationship to power (Figure 1d). Using either deeper or more diverse alignments increases *s*_*i,j*_ and detection power (Figure 1e).

We empirically fit the relationship between substitutions and detection power (at a significance threshold of E *<* 0.05) on a dataset of alignments of 87 RNA families in Rfam v14.0 with known three dimensional structures, consisting of 7,012 annotated basepairs (Figure 1f). The fitted curve enables estimation of power(*s*), the expected sensitivity for detecting any proposed basepair with *s* total substitutions. For an alignment with *B* total proposed basepairs, the expected fraction of basepairs with significant covariation support is 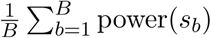. We use this number, which we call *alignment power*, to compare covariation support across different alignments with different numbers of proposed conserved basepairs. We define an arbitrary threshold of 10% alignment power to distinguish *low-power* from *high-power* alignments.

Only low-power alignments of conserved structural RNAs should lack significant covariation support. We analyzed all 3,016 seed alignments for known conserved structural RNAs in Rfam v14.1 and compared the fraction of basepairs with significant covariation support versus estimated alignment power (Figure 2a; Supplemental Table S1). Many Rfam alignments (66%, 1,985/3,016) have no statistically significant covariation support for any annotated consensus basepair, and almost all of these (98%, 1,945/1,985) are low-power alignments. Only 1% (40/3,016) are high-power alignments with no significant detected covariations (shaded in red in Figure 2a; Supplemental Table S2). Rfam, though curated, is a large compendium with a nonzero error rate. Upon examination, we believe these 40 families are enriched for inaccuracies. For example, the miR-1937 family (RF01942) (66% alignment power) is annotated in miRbase ^15^ as a tRNA sequence fragment unlikely to be a bona fide miRNA.

**Figure 2:**
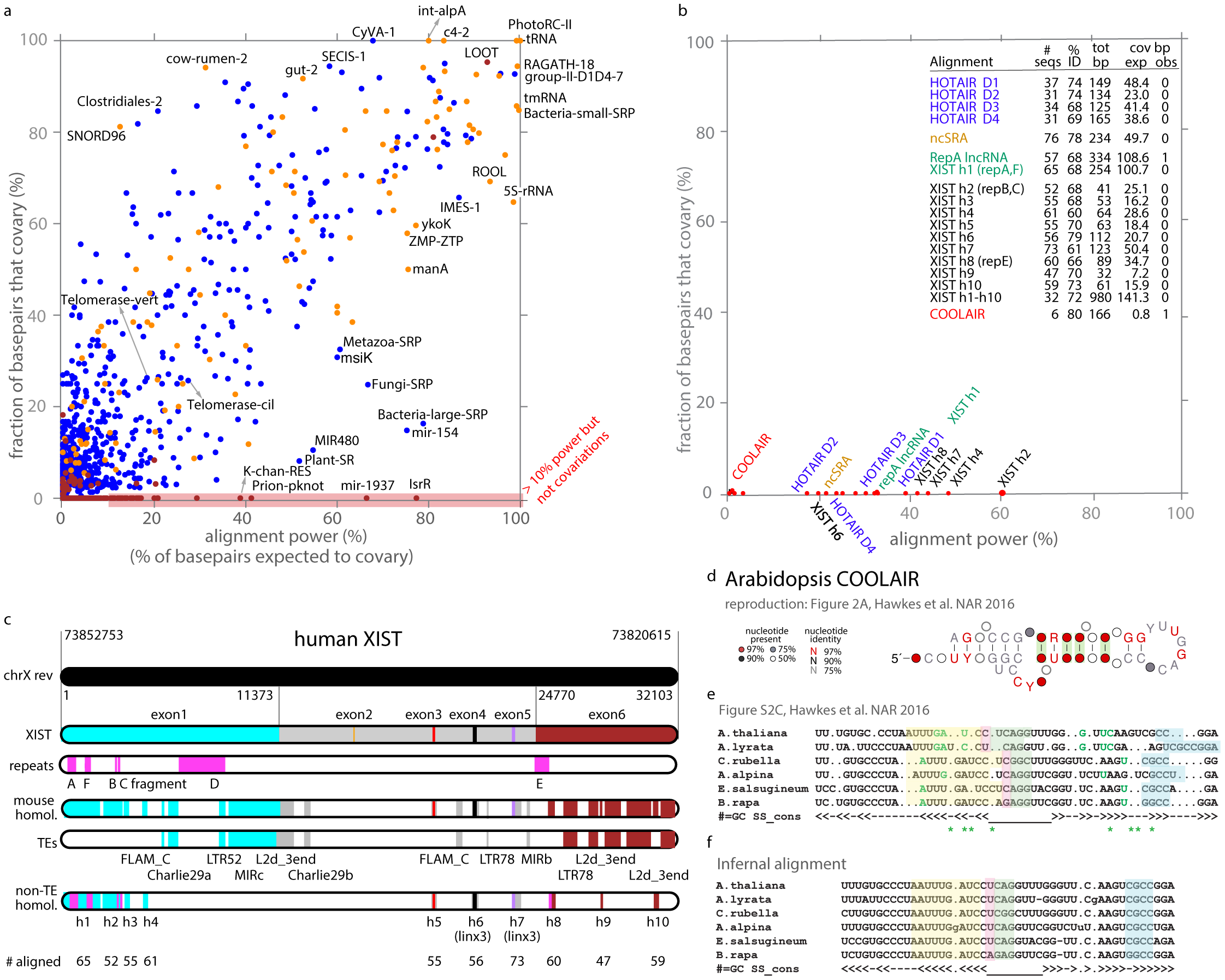
(a) Power of covariation for structural RNAs. Each point represents one of 2,209 Rfam families (seed alignments) with at least 10 annotated consensus basepairs, plotting the fraction of annotated basepairs that show a significant covariation signal (R-scape E*<* 0.05) versus “alignment power”, the fraction expected to show a significant signal. Points are color coded by positive predictive value (PPV): blue are PPV *>* 95%, yellow 50 − 95%, red *<* 50%. Red shaded region along the bottom indicates alignments with sufficient power, defined as *>* 10%, but no significant detected covariations. **(b) Results for HOTAIR, SRA, Xist, and COOLAIR lncRNA alignments**. Inset table shows details for each alignment, including the total number of annotated basepairs, the expected number that should show significant covariation (i.e. alignment power times total bp), and the number observed with significant covariation. Supplemental Table S3 describes all lncRNA alignments and proposed structures tested. **(c) Human XIST RNA conservation.** Representation of the locations of ten human/mouse unique sequence conserved regions h1-h10 in XIST/Xist, relative to XIST repeat regions A-F and to other human/mouse conserved sequence corresponding to ancient transposable elements (TEs). **(d) The proposed COOLAIR helix H10**, redrawn from ref. 21 displaying R-scape’s covariation annotation. **(e)** The proposed alignment of six homologous sequences for COOLAIR helix 10 resulting in the apparent covariations in (d). **(f) A revised COOLAIR H10 alignment more consistent with primary sequence conservation.** The apparent R-scape covariations from the original alignment in (e) disappear, while the proposed structure in (d) is maintained. Identical sets of residues in both alignments are shaded in the same color.

Previous analysis of several long noncoding RNAs including HOTAIR^9^, SRA^10^, and the Xist RepA region ^11;12^ found no significant covariation support for their proposed structures, but left open the possibility that the existing alignments lacked sufficient variation ^8^. We reanalyzed the four HOTAIR lncRNA domain alignments and consensus structures proposed by ref. 9, and the SRA alignment and consensus structure in ref. 10. All five alignments are high-power, estimated to be able to detect 23-50 significant basepair covariations each (Figure 2b; Supplemental Table S3). Although the covariation analysis of these lncRNAs has been a subject of disagreement^16;17^, these results provide new evidence for the view that HOTAIR and SRA do not have evolutionarily conserved RNA structures.

Xist RNA is perhaps the best studied lncRNA, but it remains unclear whether Xist’s role in X dosage compensation depends on any conserved RNA structure, as opposed to its sequence alone. Several different conserved structures have been proposed for the conserved 5’ RepA region of Xist^11;12;18^, two of which are based on covariation analysis of alignments of 10-13 sequences ^11;12^. Although R-scape finds no significant covariation support for the proposed RepA structures, our method finds that these are low-power alignments, so the R-scape covariation analyses are inconclusive (Figure 2b; Supplemental Table S3).

A conserved structure for the ∼1.3 kb Xist RepA lncRNA (including the conserved repeat A and F regions) has been proposed recently ^18^ from a deeper and more diverse alignment of 57 sequences. Although the R2R visualization program used by Liu et al. highlighted many potential covariations ^18^, statistical analysis by R-scape identifies only one significant covarying basepair with an E-value of 0.005, out of 334 proposed pairs. Our method judges this alignment to be high-power, estimated to be sufficient to detect about 110/334 basepairs.

The repeat A+F region is the most conserved region of Xist, but Xist is a large RNA and it is possible that other Xist regions could show covariation support for conserved RNA structure (Figure 2c). Starting with the human XIST genomic sequence, we used the *nhmmer* homology search program ^19^ to identify 21 regions of significant sequence similarity with mouse Xist (E-value *<* 10^−5^). Eleven regions correspond to insertions of well-studied ancient transposons according to Dfam analysis ^20^. For the remaining 10 unique sequence conserved regions, we iteratively built up alignments of homologs from 47-65 vertebrate species. All of these are high-power alignments; none show significant covariation support for any basepair (Figure 2b; Supplemental Table S3). In order to test for long range base pairing, we created a concatenated alignment of all ten XIST homology regions for 32 species. This concatenated alignment also has sufficient power but not covariations are observed.

Experimental evidence from chemical probing and crosslinking has been used in making structure predictions for the HOTAIR, SRA, and Xist lncRNAs. However, essentially any RNA, even a random RNA sequence, folds into some secondary structure. Lack of covariation signal in high-power RNA sequence alignments for these lncRNAs suggests that whatever structure they adopt is not detectably constraining their evolution, and thus may not be relevant for their function.

An important caveat in covariation analysis is that the input sequence alignment is assumed to be reasonably correct. Spurious apparent covariations can be created artifactually by sliding conserved primary sequence regions under proposed stems. We identified an example of this artifact in a proposed conserved structure for COOLAIR, an *Arabidopsis* lncRNA ^21^. The COOLAIR alignment is a low-power alignment of only 6 aligned sequences, yet R-scape identifies 6 significant covarying basepairs, 4 of them in one proposed helix (Figure 2d). Upon inspection, it appears that misalignment introduced artifactual covariations (Figure 2e). We realigned the COOLAIR sequences using Infernal ^22^ (Methods), which brought regions of strong primary sequence identity back into alignment (Figure 2f). The revised COOLAIR alignment is still low-power, and has only one significant supported basepair with a marginal E-value of 0.048.

The R-scape software now reports estimated statistical power calculations along with observed pairwise correlations. We expect that one important future use of covariation power analysis is to enable quantitative use of negative information by excluding pairs that are unlikely to be conserved basepairs because they have high-power and no significant covariation.

## Online Methods

### Estimating substitution number *s*_*ij*_

Given an input RNA sequence alignment, R-scape infers a maximum likelihood phylogenetic tree using FastTree (version 2.1.10) ^23^, then infers a maximum parsimony assignment of substitutions at each independent column *i* to each branch of the tree using our implementation of the Fitch algorithm ^13^. For phylogeny-aware statistical significance testing as described in ref. 8, R-scape then uses this information in tree-based simulations to construct synthetic negative control alignments that preserve average identity, composition, and phylogenetic relationships of the original alignment, while randomizing pairwise correlations. An empirical null distribution for a pairwise covariation statistic (default is the G-test statistic, related to mutual information) is then obtained from all pairs of columns *i, j* in these simulated alignments.

In the new method for statistical power estimation, the same tree and inferred maximum parsimony substitutions are used to obtain the total substitutions *s*_*i*_ (summed over branches) at each individual column *i*. For a proposed basepair involving two columns *i, j*, we use *s*_*ij*_ = *s*_*i*_ + *s*_*j*_, the sum of the independent variation at each column.

We also tested a more expensive variation where we parsimoniously infer pairwise substitutions jointly at all column pairs *i, j* to obtain *s*_*ij*_, rather than assuming column independence, which gave similar results (data not shown).

### Simulations

Simulated alignments were produced with the program R-scape-sim. Given an input sequence alignment with a consensus RNA secondary structure, R-scape-sim calculates a maximum likelihood phylogenetic tree with branch lengths. It sub-samples the original phylogenetic tree to a desired number of taxa, and linearly scales branch lengths to achieve a desired average percentage identity amongst the aligned sequences. One sequence is selected at random as a root and its evolution is simulated down the tree branches using a probabilistic evolutionary model. The evolutionary model consists of rate matrices for single (unpaired) residue substitution, pairwise (basepair) substitution, insertion, and deletion events^8;14^. We used this simulation procedure on the Rfam Cobalamin riboswitch alignment (RF00174) to generate 29,976 synthetic alignments with sequence number ranging from five to 190, and average percentage identity ranging from 30% to 100%. The simulated alignments were randomly downsampled to 3, 000 in order to produce the scatter plots of Figure 1.

### Empirical power(*s*) curve

For each of 7,012 annotated consensus basepairs in 87 RNA families Rfam v14.0 with known three dimensional structures, we use R-scape to calculate the statistical significance (expectation value, E-value) and estimate *s*_*ij*_ for each proposed pair in Rfam “seed” alignments. We binned proposed pairs with identical *s*_*ij*_ and calculated the frequency of pairs with significant support *E <* 0.05 in each bin. For 1, 653 such points for bins *s* = 0 to 1, 652, we fitted a polynomial of degree 10 by minimizing least-square-error. The choice of degree 10 was arbitrary, and simpler functions such as 1 − *e*^−*λs*^ did not fit as well.

In binning the data by integer *s*, the number of basepairs per bin is variable. For large values of *s* there are few basepairs per bin (often 0-2), leading to noisy data, which is why we fit all point for *s ≤* 150, and only those with at least 80% power for *s >* 150. For Figure 1f we plotted the Rfam data differently, in equal-size bins, by ranking all 7,012 basepairs by increasing E-values, dividing them into 70 equal bins, and calculating the means *s* and power(*s*) in each bin. This plot is less noisy at high *s*. We did not reestimate the fitted curve when we replotted. Fitted power(*s*) values starting from *s* = 0.012 are hardcoded in the R-scape source code; for *s >* 226 we set power(*s*) = 1.

Our approach treats power(*s*) as a function solely of *s*. This approximates away an important additional dependency on alignment length. R-scape E-values are multiple-test-corrected; the number of potential basepairs depends on the input alignment length. Detecting significant support for a basepair in a longer alignment requires more signal because the background of non-pairs is higher. We considered fitting power(*s, p*) to a range of different p-value thresholds *p* (i.e. before multiple test correction to E-values) but decided this was impractical. Instead the fitted power curve treats all alignments as approximately the same length. The actual lengths of the Rfam seed alignments used in Figure 1f range from 40 consensus columns (HIV retroviral Psi packing element) to 3,680 consensus columns (eukarya LSU rRNA).

### Alignment power

The expected sensitivity for detecting a basepair *b* with *s*_*b*_ substitutions is power(*s*_*b*_), from the empirical fitted curve shown in Figure 1f. In addition to reporting the pairs that significantly covary with their corresponding E-value, R-scape now also reports for each pair the inferred number of substitutions *s*_*b*_ and the estimated power(*s*_*b*_).

As an overall summary statistic for an alignment with a proposed structure, R-scape reports the total number of basepairs expected to have significant covariation support,

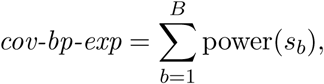

and the *alignment power*, defined as the fraction of base pairs expected to have significant support,

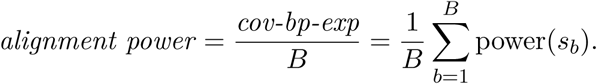

In this work, alignments with *>* 10% power are considered to have sufficient power.

### R-scape statistical test modes

R-scape has two statistical modes to test the presence of a conserved RNA structure. By default, R-scape considers all pairs as equivalent and performs an statistical test as to which of the all possible *L*(*L* − 1)*/*2 pairs (for an alignment of length *L*) are significantly covarying. This is R-scape’s default *one-set test*. Alternatively, if a consensus secondary structure is provided, R-scape allows an optional *two-set test* consisting of two independent tests on two different sets. One test is on the proposed structure (the set of basepairs); the other parallel test is on all other possible pairs in the alignment (the set of non basepairs). On the set of basepairs, R-scape extracts the alignment’s support for the annotated structure. On the set of non basepairs, R-scape identifies other possible covarying basepairs not present in the given structure.

Estimating the alignment power requires a proposed structure, thus the use of R-scape’s *two-set* mode. Under the *one-set* mode, R-scape still reports the power for each of the significantly covarying basepairs, assuming that those could be part of a structure. The covariation and covariation power analyses provided in this manuscript for all lncRNAs have been obtained with R-scape’s *two-set* mode on the proposed secondary structures.

### lncRNA alignment sources

HOTAIR domain 1-4 alignments (D1-D4) and proposed consensus structure used in ref. 9 were kindly provided to us by S. Somarowthu.

The SRA alignment and proposed consensus structure used in ref. 10 were unavailable to us. The proposed secondary structure of the human ncSRA was reproduced by hand from Supplementary Figure 1 of ref. 10. A SRA alignment was produced by imposing the human ncSRA proposed structure in the Multiz100way alignment of the ncSRA region obtained from the UCSC human genome browser (http://genome.ucsc.edu). This alignment includes 76 mammalian species.

The Xist repeat A region alignment used in ref. 11 and four alternative proposed consensus structures that we call RepA.S0 through RepA.S3 were reproduced from Supplemental Fig. 5 in ref. 11.

The Xist repeat A region alignment and proposed consensus structure in ref. 12 were kindly provided to us by W. Moss.

The RepA lncRNA alignment (spanning repeat A and repeat F regions) with a proposed consensus structure in ref. 18 was kindly provided to us by the authors.

We produced our own RepA lncRNA alignment of 65 sequences and got similar results: a high-power alignment sufficient to detect about 254 basepairs, but no significantly supported covarying pairs. The proposed structures for all ten XIST conserved regions were produced using R-scape.

As described in the main text, we used nhmmer ^19^ to identify 21 significant local alignments (at E *<* 10^−5^) between human XIST and mouse Xist, covering 79% of the XIST RNA sequence. We used nhmmer and Dfam ^20^ to determine that 11 of these conserved regions correspond to known transposable elements including retroposons (L2d 3end), DNA transposons (Charlie29a, Charlie29b), SINEs (FLAM C), and retroviral LTRs (LTR78). For the remaining 10 conserved regions we used an nhmmer profile of the mouse/human pairwise alignment to search a database of vertebrate genome sequences, resulting in 10 alignments consisting of 47-65 homologous sequences, which we name XIST h1 through XIST h10.

We used the *A. thaliana* COOLAIR lncRNA sequence and the consensus structure proposed in ref. 21 to construct a single-sequence Infernal profile ^22^, then used Infernal to align all six COOLAIR homologs to this profile.

All alignments (with consensus structure annotation, where applicable) are included in supplementary material in Stockholm format.

**Supplementary information** is linked to the online version of the paper.

## Acknowledgment

We thank the Centro de Ciencias de Benasque Pedro Pascual, Spain, where ideas for this manuscript were developed.

## Authors contribution

E.R. and S.R.E. designed the method and wrote the manuscript. E.R. wrote the code, designed and carried out the experiments.

## Author information

An R-scape web server is at eddylab.org/R-scape, with a link to download source code. The authors declare that they have no competing financial interests. Correspondence and requests for materials should be addressed to E.R.

