## Supplementary material for "Estimating the power of sequence covariation for detecting conserved RNA structure": Table_S1

### 1,411 RNA families with power, 1,031 with structural covariations

Table S1. Power of covariation for the complete Rfam v14.1 3,016 RNA families. The families are ordered by decreasing number of covariations and decreasing power. "bps" refers to basepairs annotated in the Rfam alignment; "pairs" refers to pairs annotated by R-scape as significant (E-value < 0.05). We used Rfam v14.1 seed alignments.

| | RNA family (seed alignment) | Sensitivity annotated bpairs that covary % (cov_bps/bps) | Power average power % | Positive Predictive Value covarying pairs in structure % (cov_bps/cov_pairs) | average substitutions per bpair | avg pairwise identity % | number of sequences |
|---|---|---|---|---|---|---|---|
| 1 | RF00005 tRNA | 100.0 (21/21) | 100.0 | 56.8 (21/37) | 1299.7 | 44.4 | 954 |
| 2 | RF01717 PhotoRC-II | 100.0 (13/13) | 99.2 | 92.8 (13/14) | 482.2 | 61.8 | 445 |
| 3 | RF00026 U6 | 100.0 (5/5) | 96.0 | 100.0 (5/5) | 171.0 | 70.9 | 188 |
| 4 | RF02944 c4-2 | 100.0 (24/24) | 83.3 | 82.8 (24/29) | 122.5 | 49.8 | 224 |
| 5 | RF02996 int-alpA | 100.0 (23/23) | 80.0 | 79.3 (23/29) | 105.6 | 51.2 | 167 |
| 6 | RF02965 CyVA-1 | 100.0 (18/18) | 67.8 | 100.0 (18/18) | 75.7 | 53.8 | 123 |
| 7 | RF02925 6A | 100.0 (10/10) | 56.0 | 100.0 (10/10) | 58.3 | 67.7 | 273 |
| 8 | RF00049 SNORD36 | 100.0 (4/4) | 42.5 | 100.0 (4/4) | 42.8 | 66.9 | 59 |
| 9 | RF00532 snoMe18S-Um1356 | 100.0 (3/3) | 33.3 | 100.0 (3/3) | 29.3 | 73.6 | 21 |
| 10 | RF00537 snoMe28S-Am2634 | 100.0 (3/3) | 23.3 | 100.0 (3/3) | 22.3 | 72.5 | 18 |
| 11 | RF00089 SNORD31 | 100.0 (2/2) | 20.0 | 100.0 (2/2) | 19.0 | 76.7 | 26 |
| 12 | RF01218 snR41 | 100.0 (3/3) | 13.3 | 100.0 (3/3) | 12.3 | 76.8 | 18 |
| 13 | RF00526 snoMe28S-U3344 | 100.0 (3/3) | 10.0 | 100.0 (3/3) | 12.7 | 77.8 | 8 |
| 14 | RF00529 snoMe28S-Am2589 | 100.0 (3/3) | 6.7 | 100.0 (3/3) | 9.3 | 70.3 | 16 |
| 15 | RF01196 snoZ30a | 100.0 (2/2) | 5.0 | 100.0 (2/2) | 7.0 | 78.4 | 10 |
| 16 | RF00282 SNORD48 | 100.0 (2/2) | 0.0 | 100.0 (2/2) | 4.5 | 90.4 | 8 |
| 17 | RF00270 SNORD61 | 100.0 (2/2) | 0.0 | 100.0 (2/2) | 5.5 | 71.3 | 36 |
| 18 | RF01593 plasmodium snoR16 | 100.0 (2/2) | 0.0 | 100.0 (2/2) | 3.0 | 77.2 | 3 |
| 19 | RF03000 LOOT | 95.3 (41/43) | 92.8 | 47.7 (41/86) | 317.2 | 48.3 | 368 |
| 20 | RF00442 ykkC-yxkD | 95.0 (19/20) | 83.5 | 100.0 (19/19) | 129.0 | 63.1 | 97 |
| 21 | RF03064 RAGATH-18 | 94.4 (17/18) | 99.4 | 73.9 (17/23) | 443.7 | 61.5 | 1420 |
| 22 | RF00031 SECIS 1 | 94.4 (17/18) | 58.3 | 100.0 (17/17) | 57.8 | 41.9 | 61 |
| 23 | RF03019 RT-16 | 94.1 (32/34) | 78.8 | 94.1 (32/34) | 108.7 | 48.5 | 104 |
| 24 | RF02947 cow-rumen-2 | 94.1 (16/17) | 31.2 | 84.2 (16/19) | 29.1 | 64.3 | 32 |
| 25 | RF01731 TwoAYGGAY | 93.3 (42/45) | 83.5 | 93.3 (42/45) | 130.8 | 48.8 | 210 |
| 26 | RF02967 DUF3800-VII | 93.1 (27/29) | 61.0 | 100.0 (27/27) | 70.3 | 52.0 | 139 |
| 27 | RF02924 skipping-rope | 92.8 (26/28) | 95.7 | 100.0 (26/26) | 451.8 | 53.5 | 1426 |
| 28 | RF02012 group-II-D1D4-7 | 92.7 (38/41) | 98.8 | 97.4 (38/39) | 328.4 | 51.6 | 244 |
| 29 | RF02958 drum | 92.6 (25/27) | 90.4 | 89.3 (25/28) | 152.1 | 58.1 | 464 |
| 30 | RF02913 pemK | 92.3 (12/13) | 95.4 | 92.3 (12/13) | 321.6 | 65.0 | 1542 |
| 31 | RF01699 Clostridiales-1 | 91.9 (57/62) | 80.2 | 100.0 (57/57) | 116.5 | 56.7 | 194 |
| 32 | RF02921 RT-14 | 91.9 (34/37) | 69.2 | 100.0 (34/34) | 77.6 | 57.6 | 143 |
| 33 | RF02990 gut-2 | 91.7 (11/12) | 52.5 | 91.7 (11/12) | 52.7 | 55.2 | 73 |
| 34 | RF03074 Rhodo-rpoB | 91.3 (21/23) | 80.9 | 95.5 (21/22) | 138.5 | 54.9 | 261 |
| 35 | RF03044 Proteo-phage-1 | 90.9 (20/22) | 35.5 | 100.0 (20/20) | 34.5 | 60.2 | 67 |
| 36 | RF02679 Pistol | 90.5 (19/21) | 40.5 | 100.0 (19/19) | 38.6 | 59.2 | 45 |
| 37 | RF00029 Intron gpII | 89.5 (17/19) | 65.3 | 100.0 (17/17) | 81.8 | 54.0 | 92 |
| 38 | RF02991 GP20-a | 89.5 (17/19) | 39.5 | 100.0 (17/17) | 37.5 | 53.1 | 54 |
| 39 | RF02003 group-II-D1D4-4 | 88.9 (24/27) | 82.2 | 92.3 (24/26) | 118.0 | 50.7 | 90 |
| 40 | RF03037 PAGEV | 88.1 (37/42) | 48.6 | 97.4 (37/38) | 47.8 | 53.6 | 86 |
| 41 | RF03057 nhaA-I | 87.5 (14/16) | 82.5 | 100.0 (14/14) | 182.9 | 58.7 | 281 |
| 42 | RF02401 ClpQY promoter | 87.5 (7/8) | 27.5 | 100.0 (7/7) | 26.5 | 70.6 | 39 |
| 43 | RF00020 U5 | 86.7 (26/30) | 88.3 | 78.8 (26/33) | 167.9 | 52.7 | 180 |
| 44 | RF00004 U2 | 86.7 (39/45) | 81.5 | 88.6 (39/44) | 180.1 | 59.4 | 208 |
| 45 | RF02840 Ref68 | 86.7 (26/30) | 38.3 | 100.0 (26/26) | 36.2 | 58.9 | 78 |
| 46 | RF00167 Purine | 86.4 (19/22) | 85.5 | 100.0 (19/19) | 151.8 | 54.7 | 133 |
| 47 | RF00023 tmRNA | 85.7 (102/119) | 99.2 | 85.0 (102/120) | 635.3 | 44.6 | 477 |
| 48 | RF03052 RAGATH-28 | 85.7 (12/14) | 41.4 | 100.0 (12/12) | 40.5 | 57.8 | 85 |
| 49 | RF03027 RT-6 | 85.7 (12/14) | 29.3 | 100.0 (12/12) | 27.6 | 57.4 | 59 |
| 50 | RF00230 T-box | 85.1 (40/47) | 55.1 | 100.0 (40/40) | 57.5 | 49.6 | 48 |
| 51 | RF02968 DUF3800-IX | 85.0 (17/20) | 84.5 | 85.0 (17/20) | 150.8 | 52.9 | 229 |
| 52 | RF00169 Bacteria small SRP | 84.8 (28/33) | 99.7 | 90.3 (28/31) | 367.9 | 54.1 | 261 |
| 53 | RF03003 GP20-b | 84.6 (11/13) | 70.8 | 68.8 (11/16) | 93.7 | 56.3 | 210 |
| 54 | RF00519 suhB | 84.6 (11/13) | 66.2 | 64.7 (11/17) | 81.8 | 56.2 | 87 |
| 55 | RF02937 Clostridiales-2 | 84.6 (11/13) | 20.8 | 100.0 (11/11) | 20.6 | 64.3 | 44 |
| 56 | RF02986 FuFi-1 | 84.5 (49/58) | 75.3 | 100.0 (49/49) | 94.8 | 51.6 | 170 |
| 57 | RF00174 Cobalamin | 83.3 (35/42) | 81.7 | 74.5 (35/47) | 380.0 | 52.2 | 430 |
| 58 | RF00133 SNORD33 | 83.3 (5/6) | 66.7 | 100.0 (5/5) | 71.8 | 62.9 | 72 |
| 59 | RF01852 tRNA-Sec | 83.3 (25/30) | 48.0 | 92.6 (25/27) | 49.2 | 57.0 | 109 |
| 60 | RF03015 Transposase-2 | 83.3 (15/18) | 45.5 | 100.0 (15/15) | 48.7 | 65.8 | 151 |
| 61 | RF02969 DUF3800-I | 82.3 (28/34) | 79.7 | 90.3 (28/31) | 133.9 | 54.3 | 321 |
| 62 | RF03084 DUF2815 | 81.8 (9/11) | 16.4 | 100.0 (9/9) | 16.5 | 65.7 | 22 |
| 63 | RF02004 group-II-D1D4-5 | 81.5 (44/54) | 88.9 | 93.6 (44/47) | 160.9 | 57.0 | 139 |
| 64 | RF02955 EGFOA | 81.2 (26/32) | 43.1 | 100.0 (26/26) | 42.8 | 52.7 | 73 |
| 65 | RF03047 RAGATH-16 | 81.2 (13/16) | 12.5 | 86.7 (13/15) | 13.2 | 70.2 | 18 |
| 66 | RF00015 U4 | 80.6 (25/31) | 89.7 | 92.6 (25/27) | 158.9 | 58.7 | 170 |
| 67 | RF02344 ar14 | 80.6 (25/31) | 83.2 | 100.0 (25/25) | 125.5 | 60.9 | 118 |
| 68 | RF02932 Betaproteobacteria-1 | 80.5 (29/36) | 61.7 | 93.5 (29/31) | 72.0 | 55.7 | 102 |
| 69 | RF01734 Fluoride | 80.0 (8/10) | 84.0 | 80.0 (8/10) | 214.1 | 57.8 | 287 |
| 70 | RF02221 sRNA-Xcc1 | 80.0 (16/20) | 46.0 | 80.0 (16/20) | 46.1 | 59.9 | 74 |
| 71 | RF00309 snosnR60 Z15 | 80.0 (4/5) | 30.0 | 100.0 (4/4) | 28.2 | 65.3 | 23 |
| 72 | RF02929 algC | 80.0 (4/5) | 20.0 | 100.0 (4/4) | 20.8 | 88.0 | 492 |
| 73 | RF00055 SNORD96 | 80.0 (4/5) | 6.0 | 100.0 (4/4) | 7.0 | 62.1 | 9 |

Continued on next page

| | RNA family (seed alignment) | Sensitivity annotated bpairs that covary % (cov_bps/bps) | Power average power % | Positive Predictive Value covarying pairs in structure % (cov_bps/cov_pairs) | average substitutions per bpair | avg pairwise identity % | number of sequences |
|---|---|---|---|---|---|---|---|
| 74 | RF00010 RNaseP bact a | 79.8 (91/114) | 91.0 | 93.8 (91/97) | 440.2 | 60.3 | 458 |
| 75 | RF00013 6S | 79.6 (39/49) | 84.1 | 97.5 (39/40) | 167.7 | 43.8 | 149 |
| 76 | RF02005 group-II-D1D4-6 | 79.3 (46/58) | 88.1 | 95.8 (46/48) | 194.5 | 51.4 | 174 |
| 77 | RF02953 DUF1874 | 79.3 (23/29) | 45.2 | 95.8 (23/24) | 44.9 | 61.8 | 76 |
| 78 | RF03082 dinG | 79.2 (19/24) | 39.6 | 100.0 (19/19) | 38.2 | 54.9 | 86 |
| 79 | RF00009 RNaseP nuc | 79.0 (49/62) | 77.2 | 100.0 (49/49) | 101.4 | 49.1 | 116 |
| 80 | RF01705 Flavo-1 | 78.9 (15/19) | 81.0 | 38.5 (15/39) | 201.7 | 59.1 | 201 |
| 81 | RF00557 L10 leader | 78.9 (15/19) | 73.2 | 100.0 (15/15) | 110.6 | 47.6 | 97 |
| 82 | RF01701 Cyano-1 | 78.6 (11/14) | 89.3 | 100.0 (11/11) | 266.4 | 61.7 | 265 |
| 83 | RF01695 C4 | 77.8 (21/27) | 88.5 | 70.0 (21/30) | 393.3 | 56.0 | 456 |
| 84 | RF01704 Downstream-peptide | 77.8 (14/18) | 83.3 | 100.0 (14/14) | 528.4 | 61.6 | 627 |
| 85 | RF03077 RT-2 | 77.8 (35/45) | 73.1 | 92.1 (35/38) | 102.7 | 49.5 | 195 |
| 86 | RF03081 DUF805 | 77.8 (7/9) | 20.0 | 100.0 (7/7) | 19.2 | 69.8 | 46 |
| 87 | RF01998 group-II-D1D4-1 | 77.3 (17/22) | 84.1 | 100.0 (17/17) | 275.0 | 57.6 | 237 |
| 88 | RF01051 c-di-GMP-I | 77.3 (17/22) | 75.5 | 100.0 (17/17) | 125.0 | 59.6 | 155 |
| 89 | RF01761 wcaG | 77.3 (17/22) | 70.0 | 89.5 (17/19) | 98.7 | 68.2 | 118 |
| 90 | RF03014 Transposase-1 | 76.9 (10/13) | 40.0 | 90.9 (10/11) | 39.5 | 66.0 | 176 |
| 91 | RF01690 Bacillaceae-1 | 76.5 (13/17) | 72.9 | 100.0 (13/13) | 94.3 | 63.7 | 95 |
| 92 | RF01999 group-II-D1D4-2 | 76.3 (29/38) | 79.7 | 96.7 (29/30) | 134.8 | 60.4 | 144 |
| 93 | RF02987 GA-cis | 76.2 (16/21) | 81.9 | 80.0 (16/20) | 119.6 | 59.9 | 481 |
| 94 | RF01749 pan | 76.2 (16/21) | 57.6 | 100.0 (16/16) | 62.1 | 52.6 | 74 |
| 95 | RF03075 DUF3800-VIII | 76.0 (19/25) | 72.0 | 95.0 (19/20) | 102.4 | 59.5 | 209 |
| 96 | RF00373 RNaseP arch | 75.6 (62/82) | 56.0 | 100.0 (62/62) | 60.1 | 55.2 | 70 |
| 97 | RF02001 group-II-D1D4-3 | 75.0 (36/48) | 96.7 | 94.7 (36/38) | 440.1 | 54.8 | 406 |
| 98 | RF02960 DUF2800 | 75.0 (30/40) | 50.2 | 96.8 (30/31) | 55.2 | 58.9 | 155 |
| 99 | RF00536 snoMe28S-Cm3227 | 75.0 (3/4) | 15.0 | 100.0 (3/3) | 15.5 | 90.4 | 13 |
| 100 | RF00531 snoMe28S-Gm3113 | 75.0 (3/4) | 12.5 | 100.0 (3/3) | 13.0 | 75.9 | 15 |
| 101 | RF00474 snosnR57 | 75.0 (3/4) | 5.0 | 100.0 (3/3) | 7.8 | 75.9 | 5 |
| 102 | RF00471 snosnR48 | 75.0 (3/4) | 2.5 | 100.0 (3/3) | 4.8 | 75.9 | 6 |
| 103 | RF02440 SpF59 sRNA | 72.7 (8/11) | 72.7 | 100.0 (8/8) | 166.2 | 72.1 | 410 |
| 104 | RF03070 ssNA-helicase | 72.7 (8/11) | 47.3 | 100.0 (8/8) | 48.0 | 70.9 | 104 |
| 105 | RF02916 atpB | 72.7 (16/22) | 42.7 | 100.0 (16/16) | 42.0 | 72.3 | 72 |
| 106 | RF01725 SAM-I-IV-variant | 72.4 (21/29) | 82.4 | 100.0 (21/21) | 430.9 | 56.3 | 437 |
| 107 | RF03069 malK-I | 72.2 (39/54) | 74.8 | 100.0 (39/39) | 137.0 | 58.5 | 322 |
| 108 | RF02914 DUF805b | 71.4 (5/7) | 88.6 | 100.0 (5/5) | 450.4 | 73.0 | 1230 |
| 109 | RF00059 TPP | 71.4 (20/28) | 70.7 | 100.0 (20/20) | 117.6 | 55.0 | 109 |
| 110 | RF02964 DUF3800-V | 71.4 (20/28) | 50.7 | 90.9 (20/22) | 51.0 | 51.6 | 79 |
| 111 | RF00619 U6atac | 71.4 (10/14) | 45.7 | 83.3 (10/12) | 43.7 | 71.5 | 62 |
| 112 | RF03032 narK | 71.4 (5/7) | 41.4 | 62.5 (5/8) | 42.0 | 70.1 | 308 |
| 113 | RF01748 nuoG | 71.4 (5/7) | 35.7 | 100.0 (5/5) | 33.0 | 67.5 | 31 |
| 114 | RF03080 RT-9 | 71.4 (10/14) | 24.3 | 100.0 (10/10) | 22.9 | 62.0 | 48 |
| 115 | RF02926 DUF2693-FD | 71.4 (10/14) | 24.3 | 100.0 (10/10) | 23.9 | 77.9 | 91 |
| 116 | RF02977 folE | 71.4 (5/7) | 15.7 | 100.0 (5/5) | 15.1 | 80.1 | 45 |
| 117 | RF02980 freshwater-2 | 70.6 (12/17) | 61.8 | 80.0 (12/15) | 94.3 | 61.4 | 200 |
| 118 | RF00011 RNaseP bact b | 70.5 (74/105) | 65.3 | 97.4 (74/76) | 99.9 | 66.8 | 114 |
| 119 | RF00162 SAM | 70.4 (19/27) | 72.6 | 67.8 (19/28) | 273.8 | 64.1 | 433 |
| 120 | RF03021 RT-18 | 70.0 (14/20) | 53.0 | 100.0 (14/14) | 53.5 | 62.1 | 141 |
| 121 | RF03026 RT-5 | 70.0 (21/30) | 41.3 | 95.5 (21/22) | 41.5 | 55.5 | 71 |
| 122 | RF01831 THF | 69.4 (25/36) | 70.0 | 100.0 (25/25) | 97.4 | 60.3 | 97 |
| 123 | RF03087 ROOL | 69.2 (119/172) | 93.4 | 60.7 (119/196) | 180.1 | 44.6 | 290 |
| 124 | RF01739 glnA | 69.2 (9/13) | 69.2 | 90.0 (9/10) | 674.7 | 70.8 | 956 |
| 125 | RF01687 Acido-Lenti-1 | 69.2 (9/13) | 57.7 | 100.0 (9/9) | 66.2 | 64.9 | 87 |
| 126 | RF00030 RNase MRP | 68.9 (51/74) | 57.4 | 98.1 (51/52) | 61.3 | 49.7 | 66 |
| 127 | RF02032 GOLLD | 68.4 (65/95) | 48.8 | 83.3 (65/78) | 47.9 | 49.2 | 35 |
| 128 | RF02984 DUF3800-X | 68.4 (13/19) | 42.6 | 100.0 (13/13) | 43.1 | 65.7 | 88 |
| 129 | RF00127 t44 | 68.2 (15/22) | 50.0 | 100.0 (15/15) | 58.1 | 63.7 | 78 |
| 130 | RF00080 yybP-ykoY | 68.0 (17/25) | 34.0 | 94.4 (17/18) | 31.8 | 39.2 | 29 |
| 131 | RF01055 MOCO RNA motif | 66.7 (28/42) | 71.4 | 93.3 (28/30) | 109.8 | 56.9 | 160 |
| 132 | RF02957 EFASI | 66.7 (24/36) | 54.2 | 100.0 (24/24) | 58.6 | 58.9 | 448 |
| 133 | RF00548 U11 | 66.7 (28/42) | 50.5 | 80.0 (28/35) | 53.1 | 58.9 | 72 |
| 134 | RF00095 snoPyro CD | 66.7 (2/3) | 46.7 | 100.0 (2/2) | 41.7 | 60.4 | 25 |
| 135 | RF00032 Histone3 | 66.7 (4/6) | 33.3 | 100.0 (4/4) | 32.2 | 48.3 | 46 |
| 136 | RF03020 RT-17 | 66.7 (8/12) | 28.3 | 100.0 (8/8) | 26.8 | 56.3 | 63 |
| 137 | RF00008 Hammerhead 3 | 66.7 (10/15) | 25.3 | 100.0 (10/10) | 24.0 | 69.3 | 82 |
| 138 | RF02684 Twister-P5 | 66.7 (12/18) | 13.9 | 100.0 (12/12) | 14.4 | 57.9 | 15 |
| 139 | RF01192 SNORD11B | 66.7 (4/6) | 13.3 | 100.0 (4/4) | 13.7 | 76.7 | 37 |
| 140 | RF00274 SNORD57 | 66.7 (2/3) | 6.7 | 100.0 (2/2) | 9.3 | 78.4 | 23 |
| 141 | RF00132 snoR24 | 66.7 (2/3) | 6.7 | 100.0 (2/2) | 7.0 | 80.6 | 12 |
| 142 | RF03092 Ocean-VII | 66.7 (4/6) | 5.0 | 100.0 (4/4) | 7.2 | 83.4 | 16 |
| 143 | RF01637 ceN47 | 66.7 (2/3) | 3.3 | 100.0 (2/2) | 8.0 | 76.0 | 6 |
| 144 | RF00539 snoMe28S-Cm788 | 66.7 (2/3) | 0.0 | 100.0 (2/2) | 2.0 | 89.7 | 5 |
| 145 | RF00070 SNORD29 | 66.7 (2/3) | 0.0 | 100.0 (2/2) | 2.0 | 75.2 | 10 |
| 146 | RF02034 IMES-1 | 65.7 (23/35) | 86.6 | 100.0 (23/23) | 207.8 | 71.6 | 219 |
| 147 | RF01764 yjdF | 65.2 (15/23) | 54.3 | 100.0 (15/15) | 59.4 | 67.3 | 62 |
| 148 | RF00001 5S rRNA | 64.7 (22/34) | 98.5 | 71.0 (22/31) | 306.1 | 56.1 | 712 |
| 149 | RF01685 6S-Flavo | 64.7 (11/17) | 54.7 | 100.0 (11/11) | 65.3 | 71.5 | 82 |
| 150 | RF03022 RT-10 | 64.5 (20/31) | 44.8 | 100.0 (20/20) | 44.6 | 58.1 | 113 |
| 151 | RF00168 Lysine | 64.2 (34/53) | 54.1 | 100.0 (34/34) | 55.4 | 49.2 | 47 |
| 152 | RF03055 RAGATH-30 | 63.6 (7/11) | 15.4 | 100.0 (7/7) | 16.0 | 69.2 | 33 |
| 153 | RF01754 radC | 62.9 (22/35) | 72.3 | 88.0 (22/25) | 105.7 | 58.9 | 130 |
| 154 | RF02033 HEARO | 62.9 (22/35) | 62.9 | 100.0 (22/22) | 84.9 | 59.5 | 110 |
| 155 | RF03013 nadA | 62.5 (15/24) | 69.2 | 100.0 (15/15) | 115.5 | 58.7 | 182 |
| 156 | RF02940 COG3860 | 62.5 (5/8) | 23.8 | 100.0 (5/5) | 23.0 | 76.1 | 42 |
| 157 | RF02976 Flavobacterium-1 | 62.2 (23/37) | 42.4 | 95.8 (23/24) | 44.8 | 62.3 | 210 |

Continued on next page

| | RNA family (seed alignment) | Sensitivity annotated bpairs that covary % (cov_bps/bps) | Power average power % | Positive Predictive Value covarying pairs in structure % (cov_bps/cov_pairs) | average substitutions per bpair | avg pairwise identity % | number of sequences |
|---|---|---|---|---|---|---|---|
| 158 | RF02933 ARRPOF | 62.2 (46/74) | 23.1 | 100.0 (46/46) | 22.3 | 56.2 | 35 |
| 159 | RF02956 DUF2693 | 62.1 (18/29) | 15.2 | 100.0 (18/18) | 15.5 | 65.0 | 63 |
| 160 | RF01706 Gut-1 | 61.9 (13/21) | 21.4 | 100.0 (13/13) | 20.4 | 62.5 | 38 |
| 161 | RF02541 LSU rRNA bacteria | 61.6 (521/846) | 57.5 | 95.9 (521/543) | 83.5 | 69.4 | 102 |
| 162 | RF03054 NMT1 | 61.5 (8/13) | 60.0 | 100.0 (8/8) | 128.2 | 62.3 | 382 |
| 163 | RF01068 mini-ykkC | 61.5 (8/13) | 55.4 | 100.0 (8/8) | 72.8 | 55.2 | 72 |
| 164 | RF01057 SAH riboswitch | 61.5 (8/13) | 50.8 | 100.0 (8/8) | 56.3 | 61.7 | 52 |
| 165 | RF02253 IRE II | 61.5 (8/13) | 32.3 | 100.0 (8/8) | 29.8 | 58.4 | 29 |
| 166 | RF03068 RT-3 | 60.6 (20/33) | 54.5 | 71.4 (20/28) | 59.2 | 66.9 | 231 |
| 167 | RF01707 JUMPstart | 60.0 (3/5) | 58.0 | 100.0 (3/3) | 71.8 | 78.2 | 128 |
| 168 | RF02915 DUF3800-VI | 60.0 (12/20) | 40.5 | 100.0 (12/12) | 40.2 | 60.8 | 87 |
| 169 | RF01727 SAM-SAH | 60.0 (6/10) | 36.0 | 100.0 (6/6) | 37.4 | 67.0 | 53 |
| 170 | RF02966 DUF3268 | 60.0 (21/35) | 35.7 | 100.0 (21/21) | 34.3 | 65.3 | 108 |
| 171 | RF00003 U1 | 60.0 (24/40) | 34.2 | 100.0 (24/24) | 32.0 | 63.7 | 100 |
| 172 | RF00522 PreQ1 | 60.0 (3/5) | 24.0 | 100.0 (3/3) | 21.2 | 65.7 | 35 |
| 173 | RF01718 Polynucleobacter-1 | 60.0 (18/30) | 16.0 | 100.0 (18/18) | 15.9 | 54.5 | 16 |
| 174 | RF00149 snoZ103 | 60.0 (3/5) | 6.0 | 75.0 (3/4) | 7.8 | 74.5 | 12 |
| 175 | RF01198 snoR69Y | 60.0 (3/5) | 6.0 | 100.0 (3/3) | 8.8 | 66.6 | 7 |
| 176 | RF00380 ykoK | 59.6 (28/47) | 77.2 | 93.3 (28/30) | 128.3 | 58.9 | 157 |
| 177 | RF00504 Glycine | 59.1 (13/22) | 45.0 | 100.0 (13/13) | 47.8 | 54.3 | 44 |
| 178 | RF02683 NiCo | 59.1 (13/22) | 40.0 | 100.0 (13/13) | 42.5 | 64.1 | 82 |
| 179 | RF01786 c-di-GMP-II | 58.6 (17/29) | 56.9 | 100.0 (17/17) | 63.1 | 55.2 | 54 |
| 180 | RF03005 lysM-TM7 | 58.6 (17/29) | 31.0 | 100.0 (17/17) | 30.2 | 64.7 | 129 |
| 181 | RF01746 mraW | 58.3 (21/36) | 50.0 | 100.0 (21/21) | 52.1 | 50.7 | 45 |
| 182 | RF01763 ykkC-III | 58.3 (7/12) | 36.7 | 100.0 (7/7) | 35.7 | 62.8 | 39 |
| 183 | RF02962 DUF3800-III | 58.1 (18/31) | 29.7 | 94.7 (18/19) | 29.0 | 61.5 | 55 |
| 184 | RF01750 ZMP-ZTP | 57.9 (11/19) | 75.3 | 73.3 (11/15) | 174.4 | 57.7 | 182 |
| 185 | RF01849 alpha tmRNA | 57.4 (31/54) | 58.7 | 96.9 (31/32) | 83.4 | 62.5 | 111 |
| 186 | RF02540 LSU rRNA archaea | 57.4 (451/786) | 56.8 | 97.4 (451/463) | 73.1 | 68.3 | 91 |
| 187 | RF00007 U12 | 57.1 (24/42) | 42.9 | 100.0 (24/24) | 42.4 | 62.2 | 62 |
| 188 | RF03034 nrdJ | 57.1 (12/21) | 34.3 | 100.0 (12/12) | 34.4 | 70.5 | 73 |
| 189 | RF01712 Methylobacterium-1 | 57.1 (16/28) | 32.1 | 100.0 (16/16) | 30.4 | 61.9 | 34 |
| 190 | RF00335 snoZ13 snr52 | 57.1 (4/7) | 27.1 | 100.0 (4/4) | 29.4 | 73.3 | 42 |
| 191 | RF02982 gltS | 57.1 (8/14) | 19.3 | 100.0 (8/8) | 18.9 | 75.2 | 108 |
| 192 | RF03072 raiA | 56.9 (33/58) | 62.8 | 82.5 (33/40) | 122.1 | 69.0 | 488 |
| 193 | RF00050 FMN | 56.5 (13/23) | 51.3 | 81.2 (13/16) | 82.2 | 69.2 | 144 |
| 194 | RF00233 Tymo tRNA-like | 56.5 (13/23) | 25.2 | 100.0 (13/13) | 23.6 | 69.0 | 28 |
| 195 | RF03071 DUF1646 | 56.2 (9/16) | 37.5 | 100.0 (9/9) | 37.2 | 62.6 | 146 |
| 196 | RF03012 Mu-gpT-DE | 55.5 (5/9) | 43.3 | 62.5 (5/8) | 78.7 | 74.2 | 316 |
| 197 | RF03085 abiF | 55.5 (5/9) | 32.2 | 100.0 (5/5) | 33.7 | 74.9 | 126 |
| 198 | RF03050 RAGATH-25 | 55.5 (5/9) | 7.8 | 100.0 (5/5) | 9.6 | 69.3 | 21 |
| 199 | RF02948 cow-rumen-4 | 55.5 (10/18) | 3.9 | 100.0 (10/10) | 6.4 | 61.8 | 8 |
| 200 | RF00379 ydaO-yuaA | 53.8 (14/26) | 52.7 | 82.3 (14/17) | 70.6 | 58.8 | 106 |
| 201 | RF03067 terC | 53.3 (8/15) | 50.0 | 100.0 (8/8) | 57.9 | 61.2 | 129 |
| 202 | RF03100 RAGATH-27 | 53.3 (8/15) | 7.3 | 100.0 (8/8) | 9.3 | 69.9 | 15 |
| 203 | RF03110 throat-1 | 53.1 (34/64) | 14.1 | 100.0 (34/34) | 15.0 | 71.5 | 27 |
| 204 | RF03046 Pseudomonadales-1 | 52.5 (21/40) | 23.5 | 87.5 (21/24) | 22.5 | 53.0 | 54 |
| 205 | RF00012 U3 | 52.4 (33/63) | 48.7 | 100.0 (33/33) | 50.8 | 60.1 | 87 |
| 206 | RF01857 Archaea SRP | 52.3 (69/132) | 58.6 | 98.6 (69/70) | 62.4 | 52.5 | 53 |
| 207 | RF00177 SSU rRNA bacteria | 52.2 (241/462) | 50.8 | 99.6 (241/242) | 70.7 | 73.3 | 99 |
| 208 | RF02954 DUF3577 | 52.2 (12/23) | 30.9 | 100.0 (12/12) | 30.4 | 65.1 | 86 |
| 209 | RF01794 sok | 51.9 (14/27) | 48.9 | 82.3 (14/17) | 59.8 | 58.4 | 91 |
| 210 | RF02951 DABA-DC-AT | 51.5 (17/33) | 32.4 | 100.0 (17/17) | 31.5 | 71.7 | 126 |
| 211 | RF01745 manA | 50.0 (29/58) | 75.5 | 61.7 (29/47) | 119.9 | 75.2 | 189 |
| 212 | RF00559 L21 leader | 50.0 (9/18) | 40.0 | 100.0 (9/9) | 40.0 | 55.3 | 38 |
| 213 | RF01708 L17DE | 50.0 (11/22) | 37.3 | 100.0 (11/11) | 39.0 | 63.7 | 52 |
| 214 | RF02278 Betaproteobacteria toxic sRNA | 50.0 (8/16) | 27.5 | 88.9 (8/9) | 27.7 | 70.8 | 51 |
| 215 | RF00538 snoCD11 | 50.0 (3/6) | 21.7 | 100.0 (3/3) | 20.0 | 77.3 | 26 |
| 216 | RF00325 SNORD53 SNORD92 | 50.0 (2/4) | 17.5 | 100.0 (2/2) | 18.0 | 74.5 | 37 |
| 217 | RF02961 DUF3800-II | 50.0 (10/20) | 16.0 | 90.9 (10/11) | 15.7 | 63.6 | 23 |
| 218 | RF00151 SNORD58 | 50.0 (2/4) | 12.5 | 100.0 (2/2) | 14.0 | 79.5 | 33 |
| 219 | RF01209 snR76 | 50.0 (2/4) | 10.0 | 100.0 (2/2) | 12.2 | 74.3 | 15 |
| 220 | RF00287 SNORD44 | 50.0 (1/2) | 10.0 | 100.0 (1/1) | 11.0 | 84.4 | 12 |
| 221 | RF00345 snoR1 | 50.0 (1/2) | 10.0 | 100.0 (1/1) | 12.0 | 73.7 | 13 |
| 222 | RF00071 SNORD73 | 50.0 (2/4) | 7.5 | 100.0 (2/2) | 9.0 | 79.1 | 25 |
| 223 | RF00086 SNORD27 | 50.0 (2/4) | 7.5 | 66.7 (2/3) | 8.8 | 75.6 | 8 |
| 224 | RF00300 snoZ221 snoR21b | 50.0 (2/4) | 5.0 | 100.0 (2/2) | 8.0 | 74.6 | 12 |
| 225 | RF00221 SNORD43 | 50.0 (2/4) | 2.5 | 100.0 (2/2) | 4.0 | 85.3 | 4 |
| 226 | RF00069 SNORD24 | 50.0 (2/4) | 2.5 | 100.0 (2/2) | 5.2 | 71.8 | 14 |
| 227 | RF01302 snoU36a | 50.0 (2/4) | 2.5 | 100.0 (2/2) | 3.5 | 76.2 | 15 |
| 228 | RF01221 snoU6-47 | 50.0 (2/4) | 2.5 | 100.0 (2/2) | 2.8 | 62.8 | 5 |
| 229 | RF00211 SNORD35 | 50.0 (2/4) | 0.0 | 100.0 (2/2) | 1.5 | 72.1 | 8 |
| 230 | RF01826 SAM V | 50.0 (2/4) | 0.0 | 100.0 (2/2) | 3.2 | 78.7 | 6 |
| 231 | RF01284 snoR8a | 50.0 (2/4) | 0.0 | 100.0 (2/2) | 2.8 | 73.3 | 7 |
| 232 | RF00525 Flavivirus DB | 47.4 (9/19) | 54.7 | 100.0 (9/9) | 81.8 | 73.0 | 111 |
| 233 | RF02917 Burkholderiales-2 | 47.4 (9/19) | 4.7 | 100.0 (9/9) | 5.9 | 76.6 | 19 |
| 234 | RF02943 Bacteroidales-2 | 47.3 (35/74) | 8.2 | 100.0 (35/35) | 10.0 | 74.4 | 43 |
| 235 | RF00618 U4atac | 46.7 (14/30) | 34.0 | 100.0 (14/14) | 33.7 | 62.5 | 61 |
| 236 | RF01844 SmY | 46.7 (7/15) | 34.0 | 100.0 (7/7) | 34.2 | 60.1 | 61 |
| 237 | RF02496 Atu At1 | 46.4 (13/28) | 25.0 | 100.0 (13/13) | 23.5 | 64.6 | 33 |
| 238 | RF00515 PyrR | 46.3 (19/41) | 51.7 | 95.0 (19/20) | 52.9 | 48.6 | 41 |
| 239 | RF01960 SSU rRNA eukarya | 45.9 (205/447) | 53.9 | 98.1 (205/209) | 70.1 | 62.6 | 91 |
| 240 | RF02988 GEBRO | 45.8 (33/72) | 13.5 | 100.0 (33/33) | 14.1 | 69.8 | 42 |
| 241 | RF01793 ffh | 45.5 (5/11) | 38.2 | 100.0 (5/5) | 38.9 | 64.4 | 89 |

Continued on next page

| | RNA family (seed alignment) | **Sensitivity** annotated bpairs that covary % (cov_bps/bps) | **Power** average power % | **Positive Predictive Value** covarying pairs in structure % (cov_bps/cov_pairs) | average substitutions per bpair | avg pairwise identity % | number of sequences |
|---|---|---|---|---|---|---|---|
| 242 | RF00066 U7 | 45.5 (5/11) | 36.4 | 100.0 (5/5) | 34.9 | 66.7 | 49 |
| 243 | RF03079 MISL | 45.5 (90/198) | 28.6 | 100.0 (90/90) | 27.1 | 57.0 | 44 |
| 244 | RF00634 SAM-IV | 45.2 (14/31) | 23.2 | 100.0 (14/14) | 24.0 | 70.4 | 40 |
| 245 | RF02379 csRNA | 45.0 (9/20) | 23.5 | 100.0 (9/9) | 22.7 | 79.3 | 48 |
| 246 | RF02345 ar15 | 44.8 (13/29) | 33.4 | 100.0 (13/13) | 35.0 | 71.4 | 61 |
| 247 | RF03025 RT-4 | 44.8 (13/29) | 19.0 | 92.8 (13/14) | 18.3 | 68.5 | 47 |
| 248 | RF00234 glmS | 44.7 (17/38) | 19.5 | 100.0 (17/17) | 18.7 | 59.1 | 18 |
| 249 | RF03018 RT-15 | 44.6 (25/56) | 27.9 | 100.0 (25/25) | 28.2 | 66.9 | 64 |
| 250 | RF01394 isrK | 44.4 (8/18) | 30.0 | 80.0 (8/10) | 30.9 | 60.6 | 52 |
| 251 | RF03039 Peptidase-S11 | 44.4 (8/18) | 23.3 | 100.0 (8/8) | 23.3 | 63.1 | 135 |
| 252 | RF01069 purD | 44.4 (12/27) | 22.6 | 100.0 (12/12) | 21.9 | 58.9 | 21 |
| 253 | RF01691 Bacillus-plasmid | 44.4 (4/9) | 16.7 | 100.0 (4/4) | 16.3 | 68.4 | 31 |
| 254 | RF00002 5 8S rRNA | 44.0 (11/25) | 40.4 | 100.0 (11/11) | 42.8 | 67.9 | 61 |
| 255 | RF01959 SSU rRNA archaea | 43.1 (197/457) | 44.3 | 100.0 (197/197) | 54.2 | 75.5 | 86 |
| 256 | RF01689 AdoCbl-variant | 42.9 (15/35) | 52.3 | 100.0 (15/15) | 64.4 | 78.6 | 144 |
| 257 | RF00213 snoR38 | 42.9 (3/7) | 24.3 | 100.0 (3/3) | 22.4 | 58.6 | 19 |
| 258 | RF03051 RAGATH-26 | 42.9 (3/7) | 4.3 | 100.0 (3/3) | 6.4 | 79.0 | 8 |
| 259 | RF03111 Zeta-pan | 42.9 (3/7) | 0.0 | 100.0 (3/3) | 3.0 | 80.3 | 11 |
| 260 | RF02942 Clostridiales-3 | 42.6 (23/54) | 28.9 | 100.0 (23/23) | 28.3 | 70.3 | 157 |
| 261 | RF02543 LSU rRNA eukarya | 42.4 (370/872) | 52.8 | 97.9 (370/378) | 68.9 | 65.9 | 88 |
| 262 | RF03028 RT-7 | 42.0 (21/50) | 23.8 | 100.0 (21/21) | 22.9 | 56.6 | 54 |
| 263 | RF03065 IS605-orfB-I | 41.9 (26/62) | 60.0 | 65.0 (26/40) | 77.9 | 61.2 | 259 |
| 264 | RF01709 Lacto-rpoB | 41.7 (5/12) | 40.8 | 100.0 (5/5) | 37.9 | 61.4 | 31 |
| 265 | RF00164 s2m | 41.7 (5/12) | 22.5 | 100.0 (5/5) | 22.8 | 78.4 | 38 |
| 266 | RF02946 cow-rumen-1 | 41.7 (5/12) | 15.8 | 100.0 (5/5) | 16.3 | 74.8 | 50 |
| 267 | RF00110 RybB | 41.7 (5/12) | 2.5 | 100.0 (5/5) | 4.6 | 82.5 | 8 |
| 268 | RF01688 Actino-pnp | 40.9 (9/22) | 45.0 | 100.0 (9/9) | 56.1 | 67.6 | 77 |
| 269 | RF01846 Fungi U3 | 40.5 (32/79) | 60.0 | 94.1 (32/34) | 69.0 | 56.4 | 66 |
| 270 | RF00140 Alpha RBS | 40.0 (8/20) | 35.5 | 88.9 (8/9) | 35.9 | 65.4 | 39 |
| 271 | RF00324 snoMBII-202 | 40.0 (2/5) | 28.0 | 66.7 (2/3) | 25.8 | 73.4 | 30 |
| 272 | RF02682 HDV-F-prausnitzii | 40.0 (6/15) | 26.7 | 100.0 (6/6) | 27.5 | 66.0 | 48 |
| 273 | RF02998 IS605-orfB-II | 40.0 (18/45) | 20.7 | 100.0 (18/18) | 20.9 | 69.9 | 56 |
| 274 | RF01072 TMV UPD-PK3 | 40.0 (4/10) | 15.0 | 100.0 (4/4) | 14.8 | 81.5 | 25 |
| 275 | RF00608 SNORD99 | 40.0 (2/5) | 12.0 | 100.0 (2/2) | 10.8 | 77.0 | 23 |
| 276 | RF00613 SNORD94 | 40.0 (2/5) | 10.0 | 100.0 (2/2) | 11.0 | 79.0 | 21 |
| 277 | RF00150 SNORD42 | 40.0 (2/5) | 4.0 | 100.0 (2/2) | 5.8 | 73.9 | 7 |
| 278 | RF00458 IRES Cripavirus | 40.0 (24/60) | 3.7 | 100.0 (24/24) | 6.4 | 54.8 | 7 |
| 279 | RF00204 snoR12 | 40.0 (2/5) | 2.0 | 66.7 (2/3) | 5.8 | 79.7 | 9 |
| 280 | RF02784 SorX | 40.0 (6/15) | 2.0 | 100.0 (6/6) | 3.5 | 78.5 | 8 |
| 281 | RF01317 CRISPR-DR4 | 40.0 (2/5) | 2.0 | 100.0 (2/2) | 3.6 | 83.6 | 33 |
| 282 | RF00203 snoR160 | 40.0 (2/5) | 0.0 | 100.0 (2/2) | 2.0 | 74.3 | 9 |
| 283 | RF01084 TLS-PK3 | 39.5 (17/43) | 3.7 | 100.0 (17/17) | 5.9 | 58.6 | 8 |
| 284 | RF02035 IMES-2 | 38.5 (15/39) | 63.3 | 93.8 (15/16) | 189.4 | 79.3 | 366 |
| 285 | RF00114 S15 | 38.5 (5/13) | 36.9 | 100.0 (5/5) | 38.2 | 61.0 | 78 |
| 286 | RF01290 SNORD10 | 38.5 (5/13) | 18.5 | 62.5 (5/8) | 18.7 | 64.9 | 22 |
| 287 | RF02923 HTH-XRE | 38.5 (5/13) | 15.4 | 83.3 (5/6) | 15.8 | 77.4 | 60 |
| 288 | RF00057 RyhB | 38.5 (5/13) | 11.5 | 100.0 (5/5) | 10.7 | 75.3 | 26 |
| 289 | RF03058 sul1 | 38.5 (5/13) | 10.0 | 100.0 (5/5) | 11.2 | 77.0 | 54 |
| 290 | RF03016 RT-12 | 37.8 (14/37) | 19.4 | 93.3 (14/15) | 19.2 | 56.3 | 42 |
| 291 | RF03086 chrB-b | 37.5 (6/16) | 51.2 | 100.0 (6/6) | 79.0 | 68.9 | 313 |
| 292 | RF01692 Bacteroid-trp | 37.5 (9/24) | 37.1 | 100.0 (9/9) | 35.8 | 54.2 | 35 |
| 293 | RF00558 L20 leader | 37.5 (9/24) | 30.8 | 100.0 (9/9) | 30.2 | 53.4 | 43 |
| 294 | RF02952 dfrA-dnaX | 37.5 (3/8) | 17.5 | 100.0 (3/3) | 19.1 | 74.7 | 138 |
| 295 | RF02930 aspS | 37.5 (9/24) | 16.7 | 100.0 (9/9) | 16.7 | 68.0 | 58 |
| 296 | RF00617 cHP | 37.5 (3/8) | 8.8 | 100.0 (3/3) | 10.2 | 72.7 | 59 |
| 297 | RF03041 poplar-1 | 37.5 (12/32) | 8.4 | 92.3 (12/13) | 10.0 | 69.1 | 22 |
| 298 | RF00173 Hairpin | 37.5 (3/8) | 0.0 | 100.0 (3/3) | 1.2 | 82.2 | 5 |
| 299 | RF03007 Mahella-1 | 36.4 (8/22) | 7.7 | 88.9 (8/9) | 9.6 | 64.2 | 19 |
| 300 | RF03062 xerDC | 35.9 (14/39) | 13.1 | 100.0 (14/14) | 13.9 | 75.9 | 72 |
| 301 | RF00053 mir-7 | 35.7 (10/28) | 33.9 | 100.0 (10/10) | 36.5 | 65.2 | 57 |
| 302 | RF02276 Hammerhead II | 35.3 (6/17) | 17.6 | 100.0 (6/6) | 17.4 | 65.3 | 24 |
| 303 | RF01732 asd | 34.9 (15/43) | 29.5 | 100.0 (15/15) | 28.2 | 70.4 | 31 |
| 304 | RF03045 proV | 34.6 (9/26) | 6.5 | 100.0 (9/9) | 8.4 | 71.2 | 16 |
| 305 | RF01820 RsaE | 34.5 (10/29) | 7.6 | 100.0 (10/10) | 8.1 | 75.9 | 15 |
| 306 | RF01760 traJ-II | 33.3 (11/33) | 40.0 | 91.7 (11/12) | 39.5 | 70.3 | 47 |
| 307 | RF00530 snoMe28S-Cm2645 | 33.3 (1/3) | 30.0 | 100.0 (1/1) | 28.0 | 79.0 | 20 |
| 308 | RF01848 ACEA U3 | 33.3 (17/51) | 29.4 | 100.0 (17/17) | 27.5 | 47.6 | 28 |
| 309 | RF03078 chrB-a | 33.3 (4/12) | 25.0 | 66.7 (4/6) | 27.7 | 68.7 | 163 |
| 310 | RF00609 SNORD100 | 33.3 (2/6) | 20.0 | 100.0 (2/2) | 20.2 | 76.1 | 26 |
| 311 | RF03053 OTKONC | 33.3 (10/30) | 9.0 | 55.5 (10/18) | 10.5 | 74.9 | 43 |
| 312 | RF03049 RAGATH-24 | 33.3 (9/27) | 7.8 | 100.0 (9/9) | 9.4 | 73.3 | 22 |
| 313 | RF03006 M23 | 33.3 (4/12) | 7.5 | 100.0 (4/4) | 7.5 | 73.0 | 31 |
| 314 | RF01183 SNORD19B | 33.3 (1/3) | 6.7 | 100.0 (1/1) | 9.0 | 77.5 | 17 |
| 315 | RF00533 snoMe18S-Gm1358 | 33.3 (1/3) | 6.7 | 100.0 (1/1) | 9.7 | 81.2 | 12 |
| 316 | RF02918 MDR-NUDIX | 33.3 (5/15) | 6.0 | 100.0 (5/5) | 7.1 | 70.0 | 15 |
| 317 | RF03010 mcrA | 33.3 (2/6) | 5.0 | 100.0 (2/2) | 8.5 | 80.9 | 16 |
| 318 | RF02451 ncr1241 | 33.3 (5/15) | 3.3 | 100.0 (5/5) | 5.5 | 72.0 | 8 |
| 319 | RF00295 snoTBR7 | 33.3 (1/3) | 3.3 | 100.0 (1/1) | 6.3 | 78.5 | 10 |
| 320 | RF00521 SAM alpha | 33.3 (2/6) | 3.3 | 100.0 (2/2) | 5.8 | 70.5 | 40 |
| 321 | RF00477 snosnR66 | 33.3 (2/6) | 3.3 | 100.0 (2/2) | 6.5 | 83.7 | 4 |
| 322 | RF00276 SNORD52 | 33.3 (1/3) | 3.3 | 100.0 (1/1) | 8.0 | 81.1 | 20 |
| 323 | RF00360 snoZ107 R87 | 33.3 (2/6) | 1.7 | 100.0 (2/2) | 4.3 | 74.3 | 9 |
| 324 | RF01179 snoU83 | 33.3 (1/3) | 0.0 | 100.0 (1/1) | 2.7 | 64.1 | 7 |
| 325 | RF01594 snoR18 | 33.3 (1/3) | 0.0 | 100.0 (1/1) | 1.3 | 83.7 | 3 |

Continued on next page

| | RNA family (seed alignment) | **Sensitivity** annotated bpairs that covary % (cov_bps/bps) | **Power** average power % | **Positive Predictive Value** covarying pairs in structure % (cov_bps/cov_pairs) | average substitutions per bpair | avg pairwise identity % | number of sequences |
|---|---|---|---|---|---|---|---|
| 326 | RF01711 Lnt | 33.3 (2/6) | 0.0 | 100.0 (2/2) | 2.8 | 76.5 | 10 |
| 327 | RF01348 CRISPR-DR38 | 33.3 (1/3) | 0.0 | 100.0 (1/1) | 0.7 | 81.7 | 5 |
| 328 | RF01599 plasmodium snoR24 | 33.3 (1/3) | 0.0 | 100.0 (1/1) | 2.7 | 79.6 | 4 |
| 329 | RF01285 snoU31b | 33.3 (1/3) | 0.0 | 100.0 (1/1) | 3.3 | 70.2 | 7 |
| 330 | RF01288 snoR31 | 33.3 (1/3) | 0.0 | 100.0 (1/1) | 2.7 | 70.8 | 6 |
| 331 | RF00017 Metazoa SRP | 32.5 (39/120) | 60.6 | 100.0 (39/39) | 71.5 | 68.1 | 91 |
| 332 | RF02928 Actinomyces-1 | 32.4 (11/34) | 40.9 | 100.0 (11/11) | 44.0 | 70.4 | 236 |
| 333 | RF00685 mir-36 | 32.1 (9/28) | 31.1 | 100.0 (9/9) | 30.2 | 53.9 | 46 |
| 334 | RF01066 6C | 32.0 (8/25) | 10.4 | 100.0 (8/8) | 11.9 | 71.8 | 18 |
| 335 | RF02194 HPnc0260 | 31.8 (7/22) | 21.8 | 100.0 (7/7) | 21.4 | 56.5 | 31 |
| 336 | RF02542 SSU rRNA microsporidia | 31.4 (115/366) | 34.9 | 99.1 (115/116) | 36.1 | 67.6 | 46 |
| 337 | RF03009 malK-III | 31.2 (5/16) | 6.2 | 100.0 (5/5) | 6.5 | 69.7 | 39 |
| 338 | RF02941 COG3943 | 31.0 (9/29) | 11.0 | 100.0 (9/9) | 11.4 | 69.7 | 27 |
| 339 | RF01747 msiK | 30.8 (4/13) | 60.0 | 100.0 (4/4) | 81.5 | 68.2 | 120 |
| 340 | RF01071 OLE | 30.8 (49/159) | 21.9 | 96.1 (49/51) | 21.2 | 62.8 | 20 |
| 341 | RF02931 Bacilli-1 | 30.8 (4/13) | 10.0 | 100.0 (4/4) | 11.9 | 86.2 | 145 |
| 342 | RF02963 DUF3800-IV | 30.6 (11/36) | 4.2 | 91.7 (11/12) | 6.7 | 59.6 | 10 |
| 343 | RF01693 Bacteroidales-1 | 30.4 (7/23) | 23.9 | 100.0 (7/7) | 23.2 | 75.2 | 41 |
| 344 | RF03097 RAGATH-21 | 30.4 (7/23) | 10.4 | 100.0 (7/7) | 11.0 | 76.3 | 38 |
| 345 | RF01728 STAXI | 30.3 (10/33) | 32.1 | 76.9 (10/13) | 30.4 | 57.6 | 30 |
| 346 | RF01859 Phe leader | 30.0 (12/40) | 37.8 | 100.0 (12/12) | 37.0 | 64.2 | 71 |
| 347 | RF03099 RAGATH-23 | 30.0 (6/20) | 2.0 | 100.0 (6/6) | 3.9 | 78.5 | 11 |
| 348 | RF02993 ilvH | 29.4 (5/17) | 4.7 | 100.0 (5/5) | 5.6 | 81.3 | 17 |
| 349 | RF03030 salivarius-1 | 29.4 (5/17) | 3.5 | 100.0 (5/5) | 6.7 | 72.5 | 19 |
| 350 | RF00711 mir-449 | 28.6 (8/28) | 40.7 | 100.0 (8/8) | 41.5 | 57.7 | 56 |
| 351 | RF00604 SNORD88 | 28.6 (2/7) | 20.0 | 100.0 (2/2) | 19.6 | 75.4 | 35 |
| 352 | RF03036 osmY | 28.6 (4/14) | 9.3 | 100.0 (4/4) | 10.6 | 70.6 | 38 |
| 353 | RF03001 leuA-Halobacteria | 28.6 (2/7) | 8.6 | 100.0 (2/2) | 8.3 | 76.1 | 152 |
| 354 | RF00580 SNORD91 | 28.6 (2/7) | 5.7 | 100.0 (2/2) | 7.7 | 79.1 | 24 |
| 355 | RF00288 snoZ30 | 28.6 (2/7) | 5.7 | 100.0 (2/2) | 6.6 | 82.7 | 20 |
| 356 | RF02959 DUF3085 | 27.8 (5/18) | 5.0 | 100.0 (5/5) | 6.7 | 69.9 | 16 |
| 357 | RF00016 SNORD14 | 27.3 (3/11) | 9.1 | 100.0 (3/3) | 11.2 | 67.7 | 18 |
| 358 | RF02927 Actino-ugpB | 27.3 (3/11) | 0.9 | 100.0 (3/3) | 3.5 | 72.4 | 12 |
| 359 | RF01847 Plant U3 | 27.1 (19/70) | 23.1 | 100.0 (19/19) | 22.5 | 61.6 | 24 |
| 360 | RF03103 RAGATH-34 | 26.9 (7/26) | 8.1 | 100.0 (7/7) | 10.4 | 73.9 | 24 |
| 361 | RF02949 Cupriavidus-1 | 26.9 (7/26) | 0.0 | 100.0 (7/7) | 2.7 | 62.7 | 5 |
| 362 | RF00037 IRE I | 26.7 (4/15) | 33.3 | 100.0 (4/4) | 35.4 | 76.7 | 61 |
| 363 | RF03063 Streptomyces-metK | 26.7 (4/15) | 11.3 | 100.0 (4/4) | 11.7 | 76.4 | 38 |
| 364 | RF03048 RAGATH-20 | 26.7 (4/15) | 2.0 | 100.0 (4/4) | 4.9 | 82.5 | 9 |
| 365 | RF02342 ar7 | 26.5 (9/34) | 20.9 | 100.0 (9/9) | 19.8 | 65.7 | 29 |
| 366 | RF00006 Vault | 26.3 (5/19) | 25.8 | 100.0 (5/5) | 25.3 | 59.3 | 73 |
| 367 | RF02938 COG2827 | 26.3 (5/19) | 18.4 | 100.0 (5/5) | 18.4 | 70.1 | 48 |
| 368 | RF00616 LhrC | 26.3 (5/19) | 2.6 | 100.0 (5/5) | 3.7 | 90.6 | 12 |
| 369 | RF01702 Cyano-2 | 26.1 (12/46) | 36.1 | 100.0 (12/12) | 37.3 | 63.5 | 57 |
| 370 | RF01724 SAM-Chlorobi | 26.1 (6/23) | 2.2 | 100.0 (6/6) | 3.9 | 76.6 | 9 |
| 371 | RF02972 engA | 25.9 (7/27) | 20.7 | 87.5 (7/8) | 21.3 | 70.3 | 154 |
| 372 | RF00025 Telomerase-cil | 25.7 (9/35) | 27.4 | 100.0 (9/9) | 25.6 | 54.6 | 20 |
| 373 | RF02995 IMES-5 | 25.6 (11/43) | 13.5 | 100.0 (11/11) | 14.0 | 69.4 | 52 |
| 374 | RF00024 Telomerase-vert | 25.5 (27/106) | 18.0 | 100.0 (27/27) | 17.6 | 61.5 | 37 |
| 375 | RF00096 U8 | 25.0 (6/24) | 43.3 | 100.0 (6/6) | 43.2 | 67.5 | 55 |
| 376 | RF01067 ATPC | 25.0 (6/24) | 40.4 | 100.0 (6/6) | 42.2 | 73.8 | 90 |
| 377 | RF00391 RtT | 25.0 (6/24) | 26.2 | 75.0 (6/8) | 25.7 | 68.3 | 57 |
| 378 | RF00555 L13 leader | 25.0 (4/16) | 25.0 | 100.0 (4/4) | 24.2 | 68.6 | 26 |
| 379 | RF00517 serC | 25.0 (3/12) | 21.7 | 100.0 (3/3) | 22.5 | 75.3 | 70 |
| 380 | RF00099 SNORD22 | 25.0 (1/4) | 20.0 | 100.0 (1/1) | 19.2 | 79.7 | 49 |
| 381 | RF03059 raiA-hairpin | 25.0 (2/8) | 17.5 | 100.0 (2/2) | 18.5 | 81.0 | 80 |
| 382 | RF01762 whalefall-1 | 25.0 (4/16) | 13.1 | 100.0 (4/4) | 12.7 | 71.6 | 14 |
| 383 | RF03060 uup | 25.0 (2/8) | 11.2 | 100.0 (2/2) | 11.0 | 69.7 | 203 |
| 384 | RF00280 SNORD51 | 25.0 (1/4) | 10.0 | 100.0 (1/1) | 12.2 | 86.6 | 14 |
| 385 | RF00368 sroB | 25.0 (4/16) | 10.0 | 100.0 (4/4) | 10.7 | 67.9 | 16 |
| 386 | RF02685 RAGATH-5 | 25.0 (3/12) | 9.2 | 60.0 (3/5) | 9.3 | 74.7 | 39 |
| 387 | RF02992 hya | 25.0 (3/12) | 7.5 | 100.0 (3/3) | 9.8 | 77.4 | 27 |
| 388 | RF00528 snoMe28S-Gm1083 | 25.0 (1/4) | 7.5 | 100.0 (1/1) | 9.5 | 85.7 | 11 |
| 389 | RF00572 SNORD66 | 25.0 (1/4) | 7.5 | 100.0 (1/1) | 9.0 | 81.9 | 20 |
| 390 | RF00085 SNORD28 | 25.0 (1/4) | 7.5 | 100.0 (1/1) | 7.5 | 75.9 | 20 |
| 391 | RF00273 SNORD59 | 25.0 (1/4) | 5.0 | 100.0 (1/1) | 8.0 | 77.9 | 24 |
| 392 | RF00588 SNORD41 | 25.0 (1/4) | 5.0 | 100.0 (1/1) | 5.8 | 77.2 | 20 |
| 393 | RF03008 malK-II | 25.0 (2/8) | 5.0 | 100.0 (2/2) | 7.5 | 80.5 | 94 |
| 394 | RF00479 snosnR71 | 25.0 (1/4) | 5.0 | 100.0 (1/1) | 8.5 | 73.7 | 7 |
| 395 | RF01316 CRISPR-DR3 | 25.0 (1/4) | 5.0 | 100.0 (1/1) | 4.0 | 84.0 | 20 |
| 396 | RF03073 RT-19 | 25.0 (4/16) | 3.1 | 100.0 (4/4) | 4.7 | 80.8 | 23 |
| 397 | RF01170 snoU61 | 25.0 (1/4) | 2.5 | 100.0 (1/1) | 4.8 | 74.7 | 8 |
| 398 | RF00160 snoZ159 | 25.0 (1/4) | 2.5 | 100.0 (1/1) | 4.5 | 82.0 | 10 |
| 399 | RF02459 Virus CITE 4 | 25.0 (6/24) | 1.7 | 100.0 (6/6) | 3.5 | 72.0 | 5 |
| 400 | RF02844 RefA9 | 25.0 (4/16) | 1.2 | 100.0 (4/4) | 2.9 | 62.2 | 4 |
| 401 | RF02939 COG2908 | 25.0 (3/12) | 0.8 | 100.0 (3/3) | 2.0 | 82.2 | 9 |
| 402 | RF00356 snoR32 R81 | 25.0 (1/4) | 0.0 | 100.0 (1/1) | 1.0 | 83.2 | 6 |
| 403 | RF01426 snoR126 | 25.0 (1/4) | 0.0 | 100.0 (1/1) | 1.2 | 67.3 | 4 |
| 404 | RF02519 ToxI | 25.0 (1/4) | 0.0 | 100.0 (1/1) | 0.5 | 87.8 | 4 |
| 405 | RF00153 SNORD62 | 25.0 (1/4) | 0.0 | 100.0 (1/1) | 1.2 | 86.2 | 14 |
| 406 | RF01866 ceN33 | 25.0 (1/4) | 0.0 | 100.0 (1/1) | 2.5 | 90.0 | 5 |
| 407 | RF01280 snoR14 | 25.0 (1/4) | 0.0 | 100.0 (1/1) | 0.5 | 79.3 | 7 |
| 408 | RF00475 snosnR69 | 25.0 (1/4) | 0.0 | 100.0 (1/1) | 2.5 | 64.1 | 6 |
| 409 | RF02896 S774 | 25.0 (3/12) | 0.0 | 100.0 (3/3) | 2.8 | 74.8 | 4 |

Continued on next page

| | RNA family (seed alignment) | Sensitivity annotated bpairs that covary % (cov_bps/bps) | Power average power % | Positive Predictive Value covarying pairs in structure % (cov_bps/cov_pairs) | average substitutions per bpair | avg pairwise identity % | number of sequences |
|---|---|---|---|---|---|---|---|
| 410 | RF00612 SNORD75 | 25.0 (1/4) | 0.0 | 100.0 (1/1) | 3.0 | 74.5 | 5 |
| 411 | RF00281 SNORD47 | 25.0 (1/4) | 0.0 | 100.0 (1/1) | 3.5 | 82.5 | 20 |
| 412 | RF01502 Fungi SRP | 24.8 (29/117) | 66.7 | 100.0 (29/29) | 72.5 | 43.6 | 50 |
| 413 | RF01256 snR43 | 24.1 (13/54) | 16.7 | 100.0 (13/13) | 16.3 | 64.3 | 20 |
| 414 | RF01730 Termite-leu | 23.8 (5/21) | 3.3 | 100.0 (5/5) | 6.2 | 65.6 | 20 |
| 415 | RF01054 preQ1-II | 23.5 (4/17) | 8.8 | 100.0 (4/4) | 9.8 | 70.5 | 14 |
| 416 | RF00435 ROSE | 23.5 (8/34) | 5.0 | 100.0 (8/8) | 6.8 | 61.8 | 13 |
| 417 | RF02680 PreQ1-III | 23.1 (6/26) | 13.5 | 100.0 (6/6) | 13.3 | 81.9 | 28 |
| 418 | RF03076 Streptomyces-metH | 23.1 (3/13) | 12.3 | 75.0 (3/4) | 11.9 | 66.4 | 43 |
| 419 | RF03083 cow-rumen-3 | 23.1 (9/39) | 5.1 | 100.0 (9/9) | 7.0 | 69.2 | 8 |
| 420 | RF03093 queA | 23.1 (3/13) | 3.1 | 100.0 (3/3) | 4.8 | 71.9 | 8 |
| 421 | RF02950 D12-methyl | 23.1 (3/13) | 2.3 | 100.0 (3/3) | 3.8 | 84.7 | 19 |
| 422 | RF03066 COG3610-DE | 22.9 (8/35) | 10.6 | 100.0 (8/8) | 11.5 | 73.6 | 59 |
| 423 | RF00645 MIR169 2 | 22.7 (5/22) | 37.7 | 83.3 (5/6) | 44.2 | 52.9 | 100 |
| 424 | RF01735 epsC | 22.5 (9/40) | 14.2 | 100.0 (9/9) | 14.3 | 76.7 | 30 |
| 425 | RF00353 snoR31 Z110 Z27 | 22.2 (2/9) | 5.5 | 100.0 (2/2) | 8.1 | 67.2 | 8 |
| 426 | RF02435 SpF41 sRNA | 22.2 (6/27) | 5.2 | 100.0 (6/6) | 6.9 | 69.3 | 9 |
| 427 | RF03040 PGK | 22.2 (4/18) | 2.8 | 100.0 (4/4) | 4.9 | 70.9 | 18 |
| 428 | RF02399 NsiR1 | 21.4 (3/14) | 10.7 | 100.0 (3/3) | 10.9 | 73.3 | 17 |
| 429 | RF00163 Hammerhead 1 | 21.4 (3/14) | 10.7 | 100.0 (3/3) | 12.1 | 67.8 | 29 |
| 430 | RF01721 Pseudomon-groES | 21.4 (6/28) | 7.1 | 100.0 (6/6) | 7.4 | 71.1 | 18 |
| 431 | RF00507 Corona FSE | 21.1 (4/19) | 4.7 | 80.0 (4/5) | 6.7 | 66.5 | 23 |
| 432 | RF00236 ctRNA pGA1 | 20.8 (5/24) | 7.1 | 100.0 (5/5) | 7.8 | 74.2 | 15 |
| 433 | RF02678 Hatchet | 20.8 (5/24) | 2.5 | 100.0 (5/5) | 5.2 | 54.9 | 8 |
| 434 | RF02999 ivy-DE | 20.7 (6/29) | 21.4 | 100.0 (6/6) | 21.6 | 76.0 | 248 |
| 435 | RF01867 CC2171 | 20.7 (6/29) | 7.6 | 100.0 (6/6) | 8.1 | 66.8 | 13 |
| 436 | RF02457 Virus CITE 2 | 20.3 (12/59) | 3.2 | 100.0 (12/12) | 5.4 | 71.9 | 9 |
| 437 | RF02975 DUF3800-XI | 20.0 (2/10) | 40.0 | 66.7 (2/3) | 43.7 | 72.1 | 256 |
| 438 | RF00181 SNORD113 | 20.0 (1/5) | 28.0 | 100.0 (1/1) | 25.8 | 68.0 | 59 |
| 439 | RF00284 SNORD74 | 20.0 (1/5) | 26.0 | 100.0 (1/1) | 24.4 | 78.5 | 22 |
| 440 | RF01116 Yfr1 | 20.0 (3/15) | 25.3 | 75.0 (3/4) | 24.4 | 69.0 | 37 |
| 441 | RF00054 SNORD25 | 20.0 (1/5) | 10.0 | 100.0 (1/1) | 11.6 | 68.1 | 12 |
| 442 | RF01178 snoR77Y | 20.0 (1/5) | 6.0 | 100.0 (1/1) | 7.0 | 70.0 | 21 |
| 443 | RF03017 RT-13 | 20.0 (4/20) | 4.0 | 100.0 (4/4) | 6.2 | 71.6 | 13 |
| 444 | RF02974 Fibro-purF | 20.0 (3/15) | 2.7 | 100.0 (3/3) | 4.0 | 81.8 | 12 |
| 445 | RF00186 SNORD101 | 20.0 (1/5) | 2.0 | 100.0 (1/1) | 3.6 | 85.0 | 26 |
| 446 | RF00350 snoZ152 | 20.0 (1/5) | 2.0 | 100.0 (1/1) | 4.0 | 72.1 | 9 |
| 447 | RF02979 freshwater-1 | 20.0 (3/15) | 2.0 | 75.0 (3/4) | 4.1 | 75.4 | 8 |
| 448 | RF00349 snoR11 | 20.0 (1/5) | 0.0 | 100.0 (1/1) | 1.8 | 86.0 | 5 |
| 449 | RF03091 Clostridium-PBP | 20.0 (1/5) | 0.0 | 100.0 (1/1) | 4.6 | 81.1 | 11 |
| 450 | RF00187 SNORD102 | 20.0 (1/5) | 0.0 | 100.0 (1/1) | 1.8 | 88.2 | 11 |
| 451 | RF01816 RsaA | 19.5 (8/41) | 1.7 | 100.0 (8/8) | 4.1 | 66.3 | 7 |
| 452 | RF00566 SNORA35 | 19.4 (6/31) | 11.9 | 100.0 (6/6) | 12.6 | 65.9 | 29 |
| 453 | RF00638 MIR159 | 19.2 (5/26) | 22.3 | 83.3 (5/6) | 22.9 | 53.1 | 66 |
| 454 | RF00022 GcvB | 19.1 (9/47) | 21.9 | 100.0 (9/9) | 21.3 | 64.6 | 27 |
| 455 | RF00239 mir-124 | 19.0 (4/21) | 25.2 | 100.0 (4/4) | 25.5 | 73.7 | 56 |
| 456 | RF01989 SECIS 3 | 18.8 (3/16) | 8.8 | 100.0 (3/3) | 9.6 | 81.7 | 24 |
| 457 | RF01767 SMK box riboswitch | 18.8 (3/16) | 6.2 | 100.0 (3/3) | 6.9 | 72.3 | 25 |
| 458 | RF02973 Enterococcus-1 | 18.2 (8/44) | 10.0 | 100.0 (8/8) | 11.4 | 78.5 | 78 |
| 459 | RF01759 sucC | 18.2 (2/11) | 9.1 | 100.0 (2/2) | 10.0 | 83.1 | 18 |
| 460 | RF02902 AaHKsRNA22 | 18.2 (6/33) | 3.0 | 100.0 (6/6) | 4.6 | 67.3 | 7 |
| 461 | RF00321 snoZ247 | 18.2 (2/11) | 0.0 | 50.0 (2/4) | 2.5 | 82.2 | 8 |
| 462 | RF01714 Ocean-V | 18.2 (2/11) | 0.0 | 100.0 (2/2) | 0.8 | 81.5 | 3 |
| 463 | RF02030 tp2 | 17.9 (5/28) | 12.5 | 100.0 (5/5) | 12.9 | 64.5 | 24 |
| 464 | RF00257 mir-194 | 17.9 (5/28) | 11.4 | 100.0 (5/5) | 11.4 | 69.8 | 29 |
| 465 | RF00694 mir-137 | 17.6 (6/34) | 17.1 | 100.0 (6/6) | 16.4 | 63.2 | 28 |
| 466 | RF00630 P26 | 17.6 (3/17) | 8.8 | 100.0 (3/3) | 9.6 | 70.4 | 27 |
| 467 | RF00171 Tombus 5 | 17.6 (6/34) | 2.6 | 100.0 (6/6) | 4.0 | 76.2 | 9 |
| 468 | RF00386 Entero 5 CRE | 17.2 (5/29) | 38.3 | 100.0 (5/5) | 49.5 | 83.7 | 160 |
| 469 | RF01065 23S-methyl | 17.1 (6/35) | 16.0 | 100.0 (6/6) | 15.9 | 61.6 | 19 |
| 470 | RF01736 flg-Rhizobiales | 17.1 (6/35) | 5.7 | 100.0 (6/6) | 7.3 | 69.2 | 13 |
| 471 | RF00185 Flavi CRE | 16.7 (4/24) | 42.1 | 100.0 (4/4) | 42.3 | 65.9 | 84 |
| 472 | RF01716 PhotoRC-I | 16.7 (2/12) | 23.3 | 100.0 (2/2) | 24.3 | 81.8 | 65 |
| 473 | RF00514 His leader | 16.7 (6/36) | 18.9 | 100.0 (6/6) | 18.6 | 64.4 | 33 |
| 474 | RF00147 SNORD34 | 16.7 (1/6) | 16.7 | 100.0 (1/1) | 16.2 | 69.9 | 14 |
| 475 | RF00575 SNORD70 | 16.7 (2/12) | 9.2 | 100.0 (2/2) | 10.6 | 78.9 | 21 |
| 476 | RF01410 BsrC | 16.7 (2/12) | 8.3 | 100.0 (2/2) | 10.7 | 76.1 | 29 |
| 477 | RF01723 Rhizobiales-2 | 16.7 (5/30) | 7.7 | 100.0 (5/5) | 8.7 | 78.0 | 15 |
| 478 | RF00188 SNORD103 | 16.7 (1/6) | 6.7 | 100.0 (1/1) | 9.0 | 80.7 | 41 |
| 479 | RF01315 CRISPR-DR2 | 16.7 (1/6) | 3.3 | 100.0 (1/1) | 4.5 | 78.0 | 19 |
| 480 | RF03108 Methylosinus-1 | 16.7 (6/36) | 2.5 | 100.0 (6/6) | 5.2 | 77.0 | 12 |
| 481 | RF00202 snoR66 | 16.7 (1/6) | 1.7 | 100.0 (1/1) | 2.8 | 87.3 | 6 |
| 482 | RF01299 SNORD2 | 16.7 (1/6) | 1.7 | 100.0 (1/1) | 4.5 | 76.0 | 17 |
| 483 | RF02877 NmsRb | 16.7 (4/24) | 1.2 | 100.0 (4/4) | 3.6 | 76.0 | 6 |
| 484 | RF01660 ceN88 | 16.7 (6/36) | 0.6 | 100.0 (6/6) | 1.3 | 88.6 | 6 |
| 485 | RF02067 STnc310 | 16.7 (2/12) | 0.0 | 100.0 (2/2) | 1.4 | 90.1 | 8 |
| 486 | RF00344 snoZ267 | 16.7 (1/6) | 0.0 | 100.0 (1/1) | 2.7 | 81.0 | 5 |
| 487 | RF01854 Bacteria large SRP | 16.3 (16/98) | 78.8 | 100.0 (16/16) | 103.2 | 75.3 | 92 |
| 488 | RF00091 SNORA62 | 16.0 (4/25) | 4.4 | 100.0 (4/4) | 5.5 | 76.0 | 18 |
| 489 | RF01733 atoC | 15.8 (3/19) | 1.6 | 100.0 (3/3) | 4.4 | 65.4 | 8 |
| 490 | RF01544 TB11Cs5H3 | 15.8 (3/19) | 0.5 | 100.0 (3/3) | 2.5 | 67.3 | 5 |
| 491 | RF01263 snR191 | 15.6 (12/77) | 4.0 | 100.0 (12/12) | 6.1 | 68.7 | 11 |
| 492 | RF00556 L19 leader | 15.4 (2/13) | 6.2 | 100.0 (2/2) | 7.1 | 65.2 | 10 |
| 493 | RF02887 mgtC leader | 15.4 (2/13) | 0.0 | 100.0 (2/2) | 2.2 | 74.2 | 4 |

Continued on next page

| | RNA family (seed alignment) | Sensitivity annotated bpairs that covary % (cov_bps/bps) | Power average power % | Positive Predictive Value covarying pairs in structure % (cov_bps/cov_pairs) | average substitutions per bpair | avg pairwise identity % | number of sequences |
|---|---|---|---|---|---|---|---|
| 494 | RF00396 SNORA13 | 15.0 (6/40) | 27.5 | 100.0 (6/6) | 27.4 | 67.7 | 50 |
| 495 | RF00111 SdsR RyeB | 15.0 (3/20) | 7.5 | 100.0 (3/3) | 8.4 | 75.6 | 17 |
| 496 | RF00573 SNORD67 | 15.0 (3/20) | 5.5 | 75.0 (3/4) | 7.3 | 77.4 | 15 |
| 497 | RF01720 Pseudomon-Rho | 15.0 (3/20) | 4.5 | 100.0 (3/3) | 6.0 | 78.1 | 16 |
| 498 | RF00695 MIR398 | 15.0 (3/20) | 2.5 | 100.0 (3/3) | 4.0 | 58.9 | 14 |
| 499 | RF01710 Lacto-usp | 15.0 (3/20) | 2.5 | 100.0 (3/3) | 4.8 | 65.9 | 6 |
| 500 | RF02898 AaHKsRNA20 | 15.0 (3/20) | 0.5 | 100.0 (3/3) | 2.3 | 74.8 | 6 |
| 501 | RF00641 mir-154 | 14.8 (4/27) | 75.2 | 100.0 (4/4) | 95.5 | 59.6 | 121 |
| 502 | RF00018 CsrB | 14.8 (8/54) | 21.1 | 100.0 (8/8) | 20.2 | 71.5 | 38 |
| 503 | RF01070 sucA | 14.8 (4/27) | 15.9 | 100.0 (4/4) | 15.9 | 71.2 | 35 |
| 504 | RF01089 PK-repBA | 14.8 (4/27) | 3.3 | 100.0 (4/4) | 5.0 | 62.9 | 7 |
| 505 | RF00166 PrrB RsmZ | 14.7 (5/34) | 18.8 | 100.0 (5/5) | 18.4 | 74.4 | 37 |
| 506 | RF01752 psaA | 14.5 (9/62) | 20.2 | 90.0 (9/10) | 19.2 | 75.8 | 31 |
| 507 | RF00156 SNORA70 | 14.3 (3/21) | 15.2 | 100.0 (3/3) | 15.1 | 77.1 | 43 |
| 508 | RF02348 tracrRNA | 14.3 (1/7) | 12.8 | 100.0 (1/1) | 12.6 | 80.3 | 35 |
| 509 | RF01755 rmf | 14.3 (4/28) | 11.4 | 100.0 (4/4) | 11.7 | 72.0 | 16 |
| 510 | RF00343 snoZ122 | 14.3 (1/7) | 11.4 | 100.0 (1/1) | 11.6 | 65.1 | 23 |
| 511 | RF01225 ACA64 | 14.3 (4/28) | 10.3 | 80.0 (4/5) | 11.5 | 72.3 | 30 |
| 512 | RF00191 SNORA57 | 14.3 (4/28) | 7.8 | 100.0 (4/4) | 9.1 | 78.8 | 26 |
| 513 | RF00136 SNORD81 | 14.3 (1/7) | 5.7 | 100.0 (1/1) | 7.4 | 82.0 | 22 |
| 514 | RF03033 NLPC-P60 | 14.3 (3/21) | 5.2 | 100.0 (3/3) | 7.5 | 70.2 | 17 |
| 515 | RF02402 Afu 263 | 14.3 (2/14) | 5.0 | 100.0 (2/2) | 5.7 | 77.2 | 19 |
| 516 | RF00444 PrrF | 14.3 (4/28) | 4.6 | 100.0 (4/4) | 5.7 | 81.4 | 18 |
| 517 | RF02978 folP | 14.3 (4/28) | 3.9 | 100.0 (4/4) | 5.0 | 81.4 | 28 |
| 518 | RF01737 flpD | 14.3 (2/14) | 3.6 | 100.0 (2/2) | 6.5 | 68.8 | 8 |
| 519 | RF03061 uxuA | 14.3 (2/14) | 3.6 | 100.0 (2/2) | 6.4 | 77.5 | 33 |
| 520 | RF01168 SNORD126 | 14.3 (1/7) | 2.9 | 100.0 (1/1) | 4.1 | 84.6 | 6 |
| 521 | RF02922 RAGATH-32 | 14.3 (2/14) | 2.1 | 100.0 (2/2) | 4.5 | 81.5 | 19 |
| 522 | RF00135 snoZ223 | 14.3 (1/7) | 1.4 | 100.0 (1/1) | 4.9 | 83.4 | 5 |
| 523 | RF01287 snoR27 | 14.3 (1/7) | 0.0 | 100.0 (1/1) | 1.9 | 69.0 | 6 |
| 524 | RF01186 SNORD127 | 14.3 (1/7) | 0.0 | 100.0 (1/1) | 0.3 | 86.6 | 13 |
| 525 | RF00577 SNORD72 | 14.3 (1/7) | 0.0 | 100.0 (1/1) | 2.1 | 85.7 | 16 |
| 526 | RF02732 JA04 | 14.3 (2/14) | 0.0 | 100.0 (2/2) | 1.5 | 85.5 | 6 |
| 527 | RF01856 Protozoa SRP | 13.8 (8/58) | 31.4 | 100.0 (8/8) | 28.7 | 45.4 | 18 |
| 528 | RF01411 BsrF | 13.8 (4/29) | 4.8 | 100.0 (4/4) | 7.0 | 74.1 | 11 |
| 529 | RF00838 mir-252 | 13.8 (4/29) | 3.4 | 100.0 (4/4) | 5.9 | 54.4 | 10 |
| 530 | RF00190 SNORA16 | 13.6 (3/22) | 10.4 | 100.0 (3/3) | 12.0 | 72.9 | 29 |
| 531 | RF00075 mir-166 | 13.6 (3/22) | 10.0 | 100.0 (3/3) | 10.3 | 57.6 | 31 |
| 532 | RF02347 ar45 | 13.6 (6/44) | 8.4 | 100.0 (6/6) | 9.5 | 85.9 | 36 |
| 533 | RF00546 snopsi28S-3316 | 13.5 (5/37) | 21.4 | 100.0 (5/5) | 20.5 | 70.3 | 33 |
| 534 | RF03098 RAGATH-22 | 13.5 (5/37) | 2.4 | 100.0 (5/5) | 4.9 | 80.5 | 18 |
| 535 | RF01729 Termite-flg | 13.3 (4/30) | 4.3 | 100.0 (4/4) | 6.2 | 68.4 | 13 |
| 536 | RF02920 Fusobacteriales-1 | 13.3 (4/30) | 1.3 | 100.0 (4/4) | 3.8 | 71.7 | 13 |
| 537 | RF02396 sau-5949 | 13.3 (2/15) | 0.7 | 100.0 (2/2) | 1.5 | 85.5 | 7 |
| 538 | RF02838 Ref55 | 13.3 (2/15) | 0.0 | 100.0 (2/2) | 1.4 | 68.2 | 4 |
| 539 | RF01742 lactis-plasmid | 13.2 (5/38) | 5.3 | 100.0 (5/5) | 5.6 | 83.8 | 14 |
| 540 | RF00229 IRES Picorna | 13.1 (8/61) | 25.6 | 100.0 (8/8) | 29.6 | 83.5 | 92 |
| 541 | RF00210 IRES Picorna 2 | 13.1 (16/122) | 11.4 | 100.0 (16/16) | 12.7 | 82.6 | 92 |
| 542 | RF00028 Intron gpI | 13.1 (8/61) | 10.5 | 100.0 (8/8) | 11.9 | 44.8 | 12 |
| 543 | RF00553 SCARNA1 | 13.0 (6/46) | 15.4 | 100.0 (6/6) | 16.0 | 71.0 | 29 |
| 544 | RF00238 ctRNA pND324 | 13.0 (3/23) | 7.0 | 100.0 (3/3) | 8.3 | 73.0 | 48 |
| 545 | RF02985 ftsZ-DE | 13.0 (3/23) | 1.3 | 100.0 (3/3) | 3.4 | 82.2 | 10 |
| 546 | RF02983 Fibrobacter-1 | 13.0 (3/23) | 0.4 | 75.0 (3/4) | 1.4 | 74.8 | 4 |
| 547 | RF00027 let-7 | 12.9 (4/31) | 34.5 | 100.0 (4/4) | 36.3 | 67.8 | 63 |
| 548 | RF02553 YrlA | 12.9 (4/31) | 1.6 | 100.0 (4/4) | 3.6 | 66.0 | 5 |
| 549 | RF00430 SNORA54 | 12.9 (4/31) | 0.6 | 100.0 (4/4) | 2.3 | 60.2 | 5 |
| 550 | RF00584 SNORD105 | 12.5 (1/8) | 6.2 | 100.0 (1/1) | 8.6 | 82.1 | 15 |
| 551 | RF00283 SCARNA18 | 12.5 (1/8) | 5.0 | 100.0 (1/1) | 6.5 | 86.5 | 20 |
| 552 | RF03023 rpfG | 12.5 (2/16) | 4.4 | 100.0 (2/2) | 6.9 | 85.2 | 28 |
| 553 | RF01482 AdoCbl riboswitch | 12.5 (5/40) | 3.8 | 83.3 (5/6) | 6.0 | 64.5 | 7 |
| 554 | RF02051 STnc450 | 12.5 (2/16) | 1.9 | 100.0 (2/2) | 3.5 | 77.2 | 12 |
| 555 | RF03095 RAGATH-17 | 12.5 (2/16) | 1.2 | 100.0 (2/2) | 3.1 | 79.0 | 7 |
| 556 | RF03109 Thermales-rpoB | 12.5 (2/16) | 0.6 | 50.0 (2/4) | 3.1 | 87.4 | 14 |
| 557 | RF00137 SNORD83 | 12.5 (1/8) | 0.0 | 100.0 (1/1) | 1.4 | 84.2 | 7 |
| 558 | RF01074 RF site1 | 12.5 (1/8) | 0.0 | 100.0 (1/1) | 0.8 | 81.8 | 4 |
| 559 | RF00524 R2 retro el | 12.2 (5/41) | 9.5 | 100.0 (5/5) | 10.8 | 67.5 | 15 |
| 560 | RF02500 Atu C6 | 12.0 (3/25) | 11.6 | 75.0 (3/4) | 12.3 | 68.3 | 32 |
| 561 | RF00209 IRES Pesti | 12.0 (9/75) | 7.1 | 100.0 (9/9) | 7.4 | 88.4 | 25 |
| 562 | RF03038 nhaA-II | 12.0 (3/25) | 5.2 | 75.0 (3/4) | 6.2 | 77.8 | 16 |
| 563 | RF01751 potC | 11.8 (2/17) | 40.6 | 66.7 (2/3) | 48.9 | 81.2 | 136 |
| 564 | RF01845 enod40 | 11.8 (2/17) | 8.8 | 100.0 (2/2) | 11.1 | 55.8 | 35 |
| 565 | RF03105 RAGATH-7-assoc | 11.8 (4/34) | 5.0 | 80.0 (4/5) | 7.4 | 74.9 | 26 |
| 566 | RF01248 snR8 | 11.8 (4/34) | 4.1 | 80.0 (4/5) | 5.7 | 69.2 | 11 |
| 567 | RF00063 SscA | 11.8 (2/17) | 2.4 | 100.0 (2/2) | 3.4 | 71.2 | 5 |
| 568 | RF01537 TB11Cs2H1 | 11.8 (2/17) | 0.6 | 100.0 (2/2) | 2.0 | 73.7 | 7 |
| 569 | RF01686 Acido-1 | 11.8 (2/17) | 0.6 | 100.0 (2/2) | 2.6 | 68.8 | 4 |
| 570 | RF02362 Yfr10 | 11.8 (2/17) | 0.0 | 100.0 (2/2) | 1.5 | 72.6 | 6 |
| 571 | RF01830 StyR-44 | 11.5 (3/26) | 14.2 | 100.0 (3/3) | 15.1 | 76.8 | 34 |
| 572 | RF01591 snoR15 | 11.5 (3/26) | 0.0 | 100.0 (3/3) | 1.6 | 81.3 | 4 |
| 573 | RF00706 mir-263 | 11.1 (3/27) | 21.5 | 100.0 (3/3) | 20.3 | 58.4 | 23 |
| 574 | RF03031 ssnA | 11.1 (1/9) | 15.6 | 100.0 (1/1) | 15.9 | 69.0 | 118 |
| 575 | RF00377 snoU6-53 | 11.1 (1/9) | 13.3 | 100.0 (1/1) | 14.3 | 75.2 | 25 |
| 576 | RF03115 KDPG-aldolase | 11.1 (2/18) | 6.7 | 100.0 (2/2) | 8.6 | 73.8 | 43 |
| 577 | RF00271 SNORD60 | 11.1 (1/9) | 3.3 | 50.0 (1/2) | 4.8 | 79.6 | 22 |

Continued on next page

| | RNA family (seed alignment) | Sensitivity annotated bpairs that covary % (cov_bps/bps) | Power average power % | Positive Predictive Value covarying pairs in structure % (cov_bps/cov_pairs) | average substitutions per bpair | avg pairwise identity % | number of sequences |
|---|---|---|---|---|---|---|---|
| 578 | RF01322 CRISPR-DR9 | 11.1 (1/9) | 3.3 | 100.0 (1/1) | 4.3 | 77.7 | 7 |
| 579 | RF02971 emrB-Lactobacillus | 11.1 (2/18) | 2.8 | 100.0 (2/2) | 5.2 | 82.9 | 39 |
| 580 | RF01744 livK | 11.1 (3/27) | 2.2 | 100.0 (3/3) | 3.6 | 69.1 | 8 |
| 581 | RF00359 snoZ102 R77 | 11.1 (1/9) | 2.2 | 100.0 (1/1) | 3.3 | 87.2 | 9 |
| 582 | RF00369 sroC | 11.1 (4/36) | 1.7 | 100.0 (4/4) | 2.9 | 75.0 | 6 |
| 583 | RF01713 Moco-II | 11.1 (1/9) | 1.1 | 100.0 (1/1) | 3.1 | 70.2 | 7 |
| 584 | RF03035 nqrA-Marinomonas | 11.1 (4/36) | 0.8 | 100.0 (4/4) | 2.6 | 72.3 | 5 |
| 585 | RF02595 EBv-sisRNA-1 | 11.1 (1/9) | 0.0 | 100.0 (1/1) | 1.1 | 85.8 | 8 |
| 586 | RF01258 snR10 | 10.9 (7/64) | 7.3 | 100.0 (7/7) | 8.2 | 74.1 | 15 |
| 587 | RF00021 Spot 42 | 10.8 (4/37) | 3.2 | 100.0 (4/4) | 4.6 | 79.3 | 19 |
| 588 | RF03089 cold-seep-1 | 10.8 (4/37) | 2.4 | 66.7 (4/6) | 4.9 | 71.8 | 15 |
| 589 | RF00411 SNORA9 | 10.7 (3/28) | 10.7 | 100.0 (3/3) | 11.8 | 77.6 | 36 |
| 590 | RF00996 mir-631 | 10.7 (3/28) | 1.4 | 100.0 (3/3) | 2.9 | 71.0 | 7 |
| 591 | RF03090 lysM-Actino | 10.6 (5/47) | 6.6 | 100.0 (5/5) | 8.1 | 79.4 | 56 |
| 592 | RF01412 BsrG | 10.6 (7/66) | 2.0 | 100.0 (7/7) | 5.0 | 53.8 | 6 |
| 593 | RF00782 MIR480 | 10.5 (2/19) | 54.7 | 100.0 (2/2) | 57.1 | 68.3 | 69 |
| 594 | RF01694 Bacteroides-1 | 10.5 (2/19) | 17.9 | 100.0 (2/2) | 17.8 | 75.9 | 40 |
| 595 | RF01497 ALIL | 10.5 (2/19) | 7.4 | 100.0 (2/2) | 8.0 | 67.8 | 26 |
| 596 | RF02544 mt-tmRNA | 10.5 (2/19) | 1.6 | 100.0 (2/2) | 3.5 | 79.0 | 11 |
| 597 | RF02910 Coronavirus 5p sl 1 2 | 10.5 (2/19) | 0.0 | 100.0 (2/2) | 1.5 | 73.8 | 4 |
| 598 | RF00757 mir-275 | 10.3 (3/29) | 5.9 | 100.0 (3/3) | 6.9 | 68.5 | 15 |
| 599 | RF00045 SNORA73 | 10.0 (5/50) | 33.4 | 100.0 (5/5) | 32.9 | 72.9 | 66 |
| 600 | RF01738 gabT | 10.0 (2/20) | 12.5 | 100.0 (2/2) | 12.4 | 67.9 | 17 |
| 601 | RF01191 SNORD121A | 10.0 (1/10) | 9.0 | 100.0 (1/1) | 9.9 | 80.9 | 26 |
| 602 | RF03112 Staphylococcus-1 | 10.0 (4/40) | 8.5 | 100.0 (4/4) | 9.2 | 75.1 | 21 |
| 603 | RF00079 OmrA-B | 10.0 (2/20) | 7.5 | 100.0 (2/2) | 7.8 | 76.6 | 23 |
| 604 | RF00200 snoZ199 | 10.0 (1/10) | 7.0 | 100.0 (1/1) | 8.7 | 82.4 | 17 |
| 605 | RF01776 RatA | 10.0 (1/10) | 6.0 | 100.0 (1/1) | 6.7 | 67.8 | 16 |
| 606 | RF02523 ROSE 3 | 10.0 (2/20) | 5.0 | 100.0 (2/2) | 5.6 | 71.4 | 16 |
| 607 | RF00831 mir-73 | 10.0 (3/30) | 2.0 | 100.0 (3/3) | 3.2 | 68.6 | 7 |
| 608 | RF01535 TB10Cs5H2 | 10.0 (1/10) | 1.0 | 100.0 (1/1) | 2.3 | 73.2 | 5 |
| 609 | RF01774 rpsL ricks | 10.0 (3/30) | 0.3 | 100.0 (3/3) | 1.7 | 87.4 | 7 |
| 610 | RF00436 UnaL2 | 10.0 (1/10) | 0.0 | 100.0 (1/1) | 1.8 | 76.8 | 30 |
| 611 | RF02889 Sr006 | 10.0 (3/30) | 0.0 | 75.0 (3/4) | 1.2 | 64.9 | 3 |
| 612 | RF01700 Collinsella-1 | 9.6 (5/52) | 2.9 | 71.4 (5/7) | 4.8 | 71.3 | 6 |
| 613 | RF00667 mir-33 | 9.5 (2/21) | 17.1 | 100.0 (2/2) | 17.7 | 69.5 | 50 |
| 614 | RF00513 Trp leader | 9.5 (2/21) | 13.3 | 100.0 (2/2) | 13.9 | 60.0 | 22 |
| 615 | RF01684 mascRNA-menRNA | 9.5 (2/21) | 10.5 | 100.0 (2/2) | 11.0 | 79.1 | 53 |
| 616 | RF03113 Poribacteria-1 | 9.5 (2/21) | 1.4 | 100.0 (2/2) | 3.3 | 68.0 | 6 |
| 617 | RF00161 nos TCE | 9.5 (2/21) | 1.4 | 100.0 (2/2) | 2.4 | 80.3 | 9 |
| 618 | RF01698 Chloroflexi-1 | 9.5 (2/21) | 0.0 | 100.0 (2/2) | 0.8 | 82.1 | 3 |
| 619 | RF02357 RNaseP-T | 9.4 (5/53) | 1.5 | 100.0 (5/5) | 3.3 | 82.0 | 9 |
| 620 | RF02414 rli60 | 9.3 (4/43) | 4.9 | 100.0 (4/4) | 6.9 | 63.7 | 11 |
| 621 | RF02410 snoR136 | 9.3 (4/43) | 3.2 | 100.0 (4/4) | 5.2 | 63.6 | 8 |
| 622 | RF02452 ncr1575 | 9.2 (5/54) | 8.3 | 100.0 (5/5) | 9.1 | 71.2 | 29 |
| 623 | RF00175 HIV-1 DIS | 9.1 (1/11) | 7.3 | 100.0 (1/1) | 8.5 | 87.1 | 70 |
| 624 | RF01415 Flavivirus SLIV | 9.1 (2/22) | 5.0 | 100.0 (2/2) | 5.8 | 80.5 | 20 |
| 625 | RF03056 RAGATH-35 | 9.1 (1/11) | 1.8 | 100.0 (1/1) | 4.4 | 78.2 | 16 |
| 626 | RF01508 BYDV 5 UTR | 9.1 (3/33) | 1.2 | 100.0 (3/3) | 2.8 | 82.3 | 4 |
| 627 | RF01257 snR31 | 9.1 (5/55) | 0.7 | 100.0 (5/5) | 2.6 | 74.0 | 8 |
| 628 | RF01823 rspL | 9.1 (2/22) | 0.5 | 100.0 (2/2) | 1.1 | 88.7 | 6 |
| 629 | RF02860 Arrc14 | 9.1 (4/44) | 0.0 | 100.0 (4/4) | 1.5 | 72.9 | 4 |
| 630 | RF00516 ylbH | 9.1 (3/33) | 0.0 | 100.0 (3/3) | 2.6 | 55.7 | 3 |
| 631 | RF01080 UPD-PKib | 9.1 (1/11) | 0.0 | 100.0 (1/1) | 1.9 | 83.3 | 4 |
| 632 | RF00365 mir-BHRF1-1 | 9.1 (2/22) | 0.0 | 100.0 (2/2) | 0.8 | 85.7 | 5 |
| 633 | RF02909 rli117 | 9.0 (7/78) | 0.1 | 100.0 (7/7) | 1.7 | 59.4 | 3 |
| 634 | RF00102 VA | 8.9 (4/45) | 30.0 | 100.0 (4/4) | 31.0 | 70.2 | 54 |
| 635 | RF02346 ar35 | 8.9 (4/45) | 3.8 | 100.0 (4/4) | 5.3 | 72.3 | 13 |
| 636 | RF01296 snoU85 | 8.8 (5/57) | 24.2 | 100.0 (5/5) | 22.8 | 63.6 | 21 |
| 637 | RF02520 mir-965 | 8.8 (3/34) | 19.1 | 100.0 (3/3) | 19.0 | 64.6 | 32 |
| 638 | RF00754 mir-279 | 8.8 (3/34) | 5.6 | 100.0 (3/3) | 7.5 | 66.5 | 12 |
| 639 | RF01796 frnS | 8.8 (3/34) | 5.0 | 100.0 (3/3) | 6.0 | 77.8 | 16 |
| 640 | RF03096 RAGATH-19 | 8.8 (3/34) | 3.2 | 100.0 (3/3) | 5.9 | 67.7 | 11 |
| 641 | RF00019 Y RNA | 8.7 (2/23) | 28.7 | 66.7 (2/3) | 29.1 | 65.4 | 104 |
| 642 | RF00648 MIR396 | 8.7 (2/23) | 13.0 | 100.0 (2/2) | 12.7 | 58.6 | 26 |
| 643 | RF02981 FTHFS | 8.7 (2/23) | 5.7 | 100.0 (2/2) | 8.7 | 72.9 | 27 |
| 644 | RF00291 snoR639 | 8.7 (2/23) | 0.9 | 100.0 (2/2) | 2.5 | 80.5 | 9 |
| 645 | RF01292 snoR2 | 8.6 (3/35) | 3.4 | 100.0 (3/3) | 4.9 | 73.7 | 11 |
| 646 | RF02076 STnc100 | 8.5 (4/47) | 13.0 | 100.0 (4/4) | 13.5 | 62.4 | 24 |
| 647 | RF01726 SAM-II long loops | 8.3 (1/12) | 20.0 | 50.0 (1/2) | 27.0 | 75.3 | 126 |
| 648 | RF02354 BjrC80 | 8.3 (4/48) | 9.2 | 80.0 (4/5) | 10.2 | 67.4 | 15 |
| 649 | RF00113 SIB RNA | 8.3 (2/24) | 4.6 | 100.0 (2/2) | 4.7 | 72.7 | 17 |
| 650 | RF00713 mir-239 | 8.3 (2/24) | 3.8 | 100.0 (2/2) | 4.8 | 59.0 | 9 |
| 651 | RF00543 snopsi18S-1377 | 8.3 (2/24) | 2.5 | 100.0 (2/2) | 3.9 | 72.5 | 15 |
| 652 | RF01536 TB10Cs5H3 | 8.3 (1/12) | 1.7 | 100.0 (1/1) | 3.7 | 68.6 | 7 |
| 653 | RF01504 Afu 254 | 8.3 (1/12) | 1.7 | 50.0 (1/2) | 3.8 | 55.9 | 6 |
| 654 | RF00954 MIR1446 | 8.3 (3/36) | 1.7 | 75.0 (3/4) | 3.5 | 69.7 | 7 |
| 655 | RF02945 Corio-PBP | 8.3 (2/24) | 1.2 | 100.0 (2/2) | 2.3 | 86.8 | 14 |
| 656 | RF01838 sobemo FSE | 8.3 (1/12) | 0.8 | 100.0 (1/1) | 1.4 | 68.2 | 5 |
| 657 | RF00172 CAESAR | 8.3 (1/12) | 0.8 | 100.0 (1/1) | 2.2 | 79.4 | 9 |
| 658 | RF02796 Pab160 | 8.3 (2/24) | 0.0 | 100.0 (2/2) | 1.1 | 86.2 | 4 |
| 659 | RF01855 Plant SRP | 8.1 (10/123) | 51.7 | 100.0 (10/10) | 55.0 | 64.6 | 63 |
| 660 | RF01470 rli38 | 8.1 (3/37) | 17.3 | 100.0 (3/3) | 17.3 | 63.2 | 20 |
| 661 | RF01610 ceN104 | 8.1 (3/37) | 0.8 | 75.0 (3/4) | 2.8 | 68.3 | 5 |

Continued on next page

| | RNA family (seed alignment) | Sensitivity annotated bpairs that covary % (cov_bps/bps) | Power average power % | Positive Predictive Value covarying pairs in structure % (cov_bps/cov_pairs) | average substitutions per bpair | avg pairwise identity % | number of sequences |
|---|---|---|---|---|---|---|---|
| 662 | RF00378 Qrr | 8.0 (2/25) | 8.4 | 100.0 (2/2) | 8.7 | 80.2 | 30 |
| 663 | RF00813 mir-11 | 8.0 (2/25) | 4.8 | 100.0 (2/2) | 6.2 | 70.6 | 15 |
| 664 | RF00051 mir-17 | 8.0 (2/25) | 2.0 | 100.0 (2/2) | 3.6 | 70.1 | 54 |
| 665 | RF01047 HBV epsilon | 8.0 (2/25) | 2.0 | 100.0 (2/2) | 2.4 | 97.0 | 32 |
| 666 | RF02364 Yfr13 | 8.0 (2/25) | 0.8 | 100.0 (2/2) | 2.0 | 82.6 | 7 |
| 667 | RF03107 saliva-tongue-1 | 8.0 (2/25) | 0.4 | 100.0 (2/2) | 2.9 | 77.8 | 10 |
| 668 | RF01758 sucA-II | 7.9 (5/63) | 8.1 | 100.0 (5/5) | 9.0 | 73.4 | 17 |
| 669 | RF02263 MAT2A D | 7.7 (1/13) | 18.5 | 100.0 (1/1) | 18.6 | 78.1 | 35 |
| 670 | RF02934 caiA | 7.7 (1/13) | 4.6 | 100.0 (1/1) | 7.0 | 73.0 | 14 |
| 671 | RF00947 mir-929 | 7.7 (2/26) | 3.5 | 66.7 (2/3) | 3.8 | 74.3 | 15 |
| 672 | RF03094 LAGLIDADG-2 | 7.7 (3/39) | 2.8 | 75.0 (3/4) | 4.6 | 82.9 | 25 |
| 673 | RF00488 U1 yeast | 7.7 (6/78) | 1.4 | 100.0 (6/6) | 3.4 | 63.1 | 5 |
| 674 | RF02343 ar9 | 7.5 (3/40) | 8.5 | 100.0 (3/3) | 8.1 | 76.6 | 28 |
| 675 | RF01254 snR34 | 7.5 (3/40) | 0.0 | 100.0 (3/3) | 0.7 | 89.8 | 4 |
| 676 | RF01740 gyrA | 7.4 (2/27) | 15.9 | 100.0 (2/2) | 15.7 | 64.5 | 18 |
| 677 | RF02037 IMES-4 | 7.4 (2/27) | 9.6 | 100.0 (2/2) | 10.2 | 87.3 | 30 |
| 678 | RF00722 mir-451 | 7.4 (2/27) | 7.0 | 100.0 (2/2) | 7.3 | 77.7 | 19 |
| 679 | RF00637 mir-276 | 7.4 (2/27) | 3.3 | 100.0 (2/2) | 3.4 | 73.1 | 16 |
| 680 | RF00437 HLE | 7.4 (2/27) | 1.1 | 100.0 (2/2) | 2.6 | 83.1 | 5 |
| 681 | RF02384 FasX | 7.3 (5/68) | 2.2 | 100.0 (5/5) | 3.9 | 74.2 | 8 |
| 682 | RF02411 snoR138 | 7.3 (3/41) | 1.2 | 100.0 (3/3) | 3.5 | 68.4 | 7 |
| 683 | RF02809 RsmW | 7.3 (3/41) | 0.0 | 100.0 (3/3) | 1.6 | 57.0 | 3 |
| 684 | RF00199 SL2 | 7.1 (2/28) | 17.1 | 100.0 (2/2) | 16.6 | 76.9 | 30 |
| 685 | RF00260 HepC CRE | 7.1 (1/14) | 10.7 | 100.0 (1/1) | 10.3 | 86.9 | 52 |
| 686 | RF00078 MicA | 7.1 (1/14) | 5.7 | 100.0 (1/1) | 5.9 | 82.1 | 17 |
| 687 | RF01722 Pyrobac-1 | 7.1 (1/14) | 5.0 | 100.0 (1/1) | 5.8 | 65.4 | 8 |
| 688 | RF02807 PyrR209 | 7.1 (1/14) | 0.7 | 100.0 (1/1) | 2.1 | 74.7 | 4 |
| 689 | RF01539 TB11Cs4H1 | 7.1 (1/14) | 0.7 | 100.0 (1/1) | 1.6 | 77.2 | 5 |
| 690 | RF00823 lsy-6 | 7.1 (2/28) | 0.3 | 100.0 (2/2) | 1.2 | 77.3 | 4 |
| 691 | RF01643 ceN56 | 7.1 (2/28) | 0.0 | 100.0 (2/2) | 0.9 | 83.9 | 4 |
| 692 | RF02803 PyrR205 | 7.1 (1/14) | 0.0 | 100.0 (1/1) | 0.7 | 87.3 | 5 |
| 693 | RF00370 sroD | 7.1 (1/14) | 0.0 | 100.0 (1/1) | 0.4 | 85.4 | 5 |
| 694 | RF00545 snopsi18S-841 | 7.0 (3/43) | 15.8 | 100.0 (3/3) | 15.3 | 69.0 | 24 |
| 695 | RF01756 rne-II | 6.9 (5/72) | 7.1 | 100.0 (5/5) | 7.4 | 75.6 | 16 |
| 696 | RF01809 symR | 6.9 (2/29) | 3.8 | 100.0 (2/2) | 4.8 | 78.0 | 20 |
| 697 | RF00422 SCARNA24 | 6.9 (2/29) | 3.1 | 100.0 (2/2) | 5.0 | 83.1 | 23 |
| 698 | RF03043 Prevotella-2 | 6.9 (2/29) | 2.1 | 100.0 (2/2) | 5.2 | 74.5 | 20 |
| 699 | RF00014 DsrA | 6.9 (2/29) | 0.7 | 100.0 (2/2) | 2.1 | 83.3 | 5 |
| 700 | RF00949 mir-983 | 6.9 (2/29) | 0.0 | 100.0 (2/2) | 0.9 | 77.2 | 3 |
| 701 | RF01247 snR32 | 6.8 (3/44) | 4.8 | 100.0 (3/3) | 6.8 | 66.9 | 11 |
| 702 | RF00534 SgrS | 6.8 (4/59) | 4.1 | 100.0 (4/4) | 7.3 | 48.0 | 8 |
| 703 | RF01782 ASpks | 6.7 (1/15) | 25.3 | 100.0 (1/1) | 25.6 | 78.5 | 77 |
| 704 | RF00112 CyaR RyeE | 6.7 (1/15) | 6.0 | 100.0 (1/1) | 6.3 | 77.6 | 17 |
| 705 | RF01213 snoR103 | 6.7 (1/15) | 3.3 | 100.0 (1/1) | 4.3 | 66.8 | 11 |
| 706 | RF00170 msr | 6.7 (1/15) | 1.3 | 100.0 (1/1) | 4.1 | 63.0 | 9 |
| 707 | RF02437 SpF44 sRNA | 6.7 (2/30) | 1.3 | 100.0 (2/2) | 3.1 | 83.7 | 12 |
| 708 | RF01557 TB9Cs4H1 | 6.7 (1/15) | 0.7 | 100.0 (1/1) | 1.8 | 72.2 | 4 |
| 709 | RF01251 snR3 | 6.7 (3/45) | 0.2 | 100.0 (3/3) | 1.1 | 86.8 | 4 |
| 710 | RF02806 PyrR208 | 6.7 (1/15) | 0.0 | 100.0 (1/1) | 1.1 | 72.4 | 4 |
| 711 | RF01073 GP knot1 | 6.7 (1/15) | 0.0 | 100.0 (1/1) | 1.9 | 76.6 | 7 |
| 712 | RF00341 snoZ39 | 6.7 (1/15) | 0.0 | 100.0 (1/1) | 0.7 | 88.4 | 5 |
| 713 | RF01548 TB6Cs1H4 | 6.7 (1/15) | 0.0 | 100.0 (1/1) | 1.4 | 79.2 | 4 |
| 714 | RF01554 TB9Cs2H1 | 6.7 (1/15) | 0.0 | 100.0 (1/1) | 0.5 | 84.6 | 4 |
| 715 | RF00094 HDV ribozyme | 6.5 (2/31) | 7.7 | 100.0 (2/2) | 8.8 | 74.0 | 33 |
| 716 | RF00789 mir-286 | 6.5 (2/31) | 7.4 | 100.0 (2/2) | 7.8 | 69.2 | 14 |
| 717 | RF00540 snopsi18S-1854 | 6.5 (2/31) | 3.2 | 100.0 (2/2) | 4.1 | 78.5 | 16 |
| 718 | RF01608 ceN102 | 6.5 (2/31) | 0.3 | 100.0 (2/2) | 1.0 | 75.3 | 4 |
| 719 | RF00820 mir-248 | 6.5 (2/31) | 0.3 | 100.0 (2/2) | 1.4 | 70.0 | 4 |
| 720 | RF00810 mir-85 | 6.5 (2/31) | 0.3 | 100.0 (2/2) | 1.3 | 67.3 | 5 |
| 721 | RF02905 sRNA41 | 6.5 (2/31) | 0.0 | 100.0 (2/2) | 1.7 | 67.3 | 3 |
| 722 | RF02997 HOLDH | 6.3 (6/95) | 2.3 | 100.0 (6/6) | 4.6 | 74.2 | 13 |
| 723 | RF02502 Atu C8 | 6.2 (2/32) | 11.6 | 66.7 (2/3) | 12.0 | 69.1 | 27 |
| 724 | RF01456 VrrA | 6.2 (2/32) | 4.7 | 100.0 (2/2) | 6.3 | 86.2 | 11 |
| 725 | RF00720 mir-317 | 6.2 (2/32) | 3.8 | 100.0 (2/2) | 5.3 | 72.2 | 12 |
| 726 | RF03011 Methylophilales-1 | 6.2 (2/32) | 3.4 | 100.0 (2/2) | 4.5 | 86.0 | 14 |
| 727 | RF01835 HIV FS2 | 6.2 (1/16) | 3.1 | 100.0 (1/1) | 4.2 | 86.4 | 19 |
| 728 | RF00374 Gammaretro CES | 6.2 (2/32) | 2.2 | 100.0 (2/2) | 2.9 | 89.3 | 22 |
| 729 | RF01555 TB9Cs3H1 | 6.2 (1/16) | 1.9 | 100.0 (1/1) | 3.5 | 69.8 | 6 |
| 730 | RF00624 P9 | 6.2 (1/16) | 1.9 | 100.0 (1/1) | 2.4 | 77.6 | 17 |
| 731 | RF02846 Ref84 | 6.2 (1/16) | 0.6 | 100.0 (1/1) | 2.3 | 81.0 | 5 |
| 732 | RF01802 HSUR | 6.2 (1/16) | 0.6 | 100.0 (1/1) | 2.8 | 74.2 | 4 |
| 733 | RF02716 sno ncR3 | 6.2 (1/16) | 0.6 | 100.0 (1/1) | 1.9 | 82.1 | 5 |
| 734 | RF01532 TB10Cs3H2 | 6.2 (1/16) | 0.6 | 100.0 (1/1) | 1.8 | 77.8 | 6 |
| 735 | RF02790 sodF sRNA | 6.2 (2/32) | 0.3 | 100.0 (2/2) | 1.5 | 82.8 | 6 |
| 736 | RF02848 RefB11 | 6.2 (3/48) | 0.2 | 100.0 (3/3) | 2.0 | 62.8 | 4 |
| 737 | RF00800 mir-353 | 6.2 (2/32) | 0.0 | 100.0 (2/2) | 1.4 | 77.6 | 4 |
| 738 | RF00033 MicF | 6.2 (1/16) | 0.0 | 100.0 (1/1) | 1.9 | 84.0 | 4 |
| 739 | RF01489 sbrA | 6.2 (1/16) | 0.0 | 100.0 (1/1) | 0.9 | 91.3 | 6 |
| 740 | RF02096 mir-2973 | 6.2 (2/32) | 0.0 | 100.0 (2/2) | 1.5 | 80.6 | 7 |
| 741 | RF00797 mir-355 | 6.2 (2/32) | 0.0 | 100.0 (2/2) | 1.0 | 72.4 | 4 |
| 742 | RF01227 snoR83 | 6.1 (2/33) | 1.8 | 66.7 (2/3) | 3.2 | 72.6 | 7 |
| 743 | RF03104 RAGATH-36 | 6.1 (2/33) | 1.5 | 100.0 (2/2) | 4.2 | 80.2 | 14 |
| 744 | RF03088 Parabacteroides-1 | 6.1 (2/33) | 1.5 | 100.0 (2/2) | 3.9 | 78.1 | 15 |
| 745 | RF02834 VqmR | 6.1 (2/33) | 0.3 | 100.0 (2/2) | 1.6 | 72.4 | 4 |

Continued on next page

| | RNA family (seed alignment) | Sensitivity annotated bpairs that covary % (cov_bps/bps) | Power average power % | Positive Predictive Value covarying pairs in structure % (cov_bps/cov_pairs) | average substitutions per bpair | avg pairwise identity % | number of sequences |
|---|---|---|---|---|---|---|---|
| 746 | RF00083 GlmZ SraJ | 6.0 (3/50) | 3.6 | 100.0 (3/3) | 4.0 | 79.0 | 21 |
| 747 | RF01715 Pedo-repair | 5.9 (4/68) | 0.6 | 100.0 (4/4) | 2.2 | 79.2 | 6 |
| 748 | RF01046 WLE3 | 5.9 (1/17) | 0.6 | 100.0 (1/1) | 2.4 | 84.3 | 10 |
| 749 | RF03102 RAGATH-33 | 5.9 (1/17) | 0.6 | 100.0 (1/1) | 2.6 | 82.5 | 7 |
| 750 | RF01543 TB11Cs5H2 | 5.9 (1/17) | 0.6 | 100.0 (1/1) | 1.8 | 71.7 | 5 |
| 751 | RF00794 mir-42 | 5.9 (2/34) | 0.3 | 100.0 (2/2) | 1.1 | 78.2 | 5 |
| 752 | RF02900 AaHKsRNA82 | 5.9 (1/17) | 0.0 | 100.0 (1/1) | 1.9 | 66.8 | 4 |
| 753 | RF02431 SpF22 sRNA | 5.7 (2/35) | 7.4 | 100.0 (2/2) | 8.4 | 75.3 | 22 |
| 754 | RF02413 snoR145 | 5.7 (2/35) | 6.3 | 100.0 (2/2) | 7.4 | 69.1 | 19 |
| 755 | RF02469 Ms IGR-7 | 5.7 (2/35) | 3.1 | 100.0 (2/2) | 5.0 | 72.5 | 14 |
| 756 | RF00263 SNORA68 | 5.5 (2/36) | 15.3 | 100.0 (2/2) | 15.6 | 70.9 | 26 |
| 757 | RF00668 mir-302 | 5.5 (1/18) | 14.4 | 100.0 (1/1) | 14.9 | 72.7 | 44 |
| 758 | RF00643 MIR171 1 | 5.5 (1/18) | 12.2 | 100.0 (1/1) | 14.0 | 59.8 | 57 |
| 759 | RF01696 Chlorobi-1 | 5.5 (1/18) | 7.8 | 100.0 (1/1) | 9.1 | 74.6 | 15 |
| 760 | RF02919 ilvB-OMG | 5.5 (3/55) | 7.3 | 100.0 (3/3) | 8.1 | 79.5 | 28 |
| 761 | RF00591 SNORD77 | 5.5 (1/18) | 6.7 | 100.0 (1/1) | 8.0 | 82.4 | 21 |
| 762 | RF02509 PYLIS 2 | 5.5 (1/18) | 5.5 | 100.0 (1/1) | 6.5 | 75.1 | 23 |
| 763 | RF02144 rsmX | 5.5 (2/36) | 4.7 | 66.7 (2/3) | 6.1 | 79.0 | 16 |
| 764 | RF01295 SCARNA7 | 5.5 (1/18) | 2.8 | 100.0 (1/1) | 5.0 | 81.3 | 18 |
| 765 | RF00165 Corona pk3 | 5.5 (1/18) | 2.2 | 100.0 (1/1) | 4.9 | 69.3 | 14 |
| 766 | RF02349 psRNA2 | 5.5 (3/55) | 2.2 | 100.0 (3/3) | 4.1 | 64.0 | 7 |
| 767 | RF02266 XIST intron | 5.5 (1/18) | 2.2 | 100.0 (1/1) | 2.7 | 91.3 | 20 |
| 768 | RF02275 Hammerhead HH9 | 5.5 (1/18) | 1.7 | 100.0 (1/1) | 2.4 | 89.3 | 33 |
| 769 | RF00520 ybhL | 5.5 (1/18) | 1.1 | 100.0 (1/1) | 2.8 | 74.5 | 6 |
| 770 | RF00688 MIR394 | 5.5 (1/18) | 1.1 | 100.0 (1/1) | 1.6 | 70.7 | 19 |
| 771 | RF00481 HCV X3 | 5.5 (2/36) | 1.1 | 100.0 (2/2) | 1.7 | 95.8 | 22 |
| 772 | RF01768 flavi FSE | 5.5 (1/18) | 0.6 | 100.0 (1/1) | 1.6 | 87.0 | 7 |
| 773 | RF01620 ceN125 | 5.5 (2/36) | 0.6 | 50.0 (2/4) | 1.2 | 82.6 | 4 |
| 774 | RF01430 snoR134 | 5.5 (2/36) | 0.3 | 100.0 (2/2) | 1.8 | 68.3 | 5 |
| 775 | RF00795 mir-43 | 5.5 (2/36) | 0.0 | 100.0 (2/2) | 0.9 | 73.2 | 4 |
| 776 | RF00293 snoM1 | 5.5 (1/18) | 0.0 | 100.0 (1/1) | 1.4 | 89.1 | 7 |
| 777 | RF00389 satBaMV CRE | 5.3 (2/38) | 6.3 | 100.0 (2/2) | 6.5 | 93.1 | 42 |
| 778 | RF00303 snoR86 | 5.3 (1/19) | 3.7 | 100.0 (1/1) | 4.3 | 76.6 | 11 |
| 779 | RF02353 BjrC68 | 5.3 (1/19) | 3.7 | 100.0 (1/1) | 4.8 | 80.7 | 12 |
| 780 | RF03101 RAGATH-31 | 5.3 (1/19) | 1.6 | 100.0 (1/1) | 2.7 | 87.8 | 26 |
| 781 | RF01807 GIR1 | 5.3 (3/56) | 1.6 | 100.0 (3/3) | 3.8 | 76.5 | 12 |
| 782 | RF00138 SNORD16 | 5.3 (1/19) | 1.1 | 100.0 (1/1) | 2.2 | 79.5 | 5 |
| 783 | RF03004 Lacto-phage-1 | 5.3 (1/19) | 0.5 | 100.0 (1/1) | 2.6 | 77.3 | 9 |
| 784 | RF02823 LPR69 | 5.3 (1/19) | 0.5 | 100.0 (1/1) | 2.3 | 75.8 | 5 |
| 785 | RF02530 URE2 IRES | 5.3 (1/19) | 0.0 | 100.0 (1/1) | 1.2 | 86.7 | 5 |
| 786 | RF00056 SNORA71 | 5.1 (2/39) | 13.6 | 100.0 (2/2) | 14.6 | 78.6 | 40 |
| 787 | RF00791 mir-357 | 5.1 (2/39) | 1.0 | 100.0 (2/2) | 2.9 | 66.7 | 5 |
| 788 | RF02029 sraA | 5.0 (1/20) | 5.5 | 50.0 (1/2) | 7.0 | 73.9 | 20 |
| 789 | RF02269 HPnc0580 | 5.0 (1/20) | 0.5 | 100.0 (1/1) | 2.2 | 85.5 | 7 |
| 790 | RF02681 Twister-sister | 5.0 (1/20) | 0.5 | 100.0 (1/1) | 1.4 | 78.4 | 4 |
| 791 | RF00906 MIR1122 | 4.9 (2/41) | 2.2 | 100.0 (2/2) | 3.7 | 71.6 | 12 |
| 792 | RF01612 ceN105 | 4.9 (2/41) | 0.2 | 100.0 (2/2) | 1.6 | 66.4 | 3 |
| 793 | RF00657 mir-184 | 4.8 (1/21) | 6.2 | 100.0 (1/1) | 6.8 | 68.4 | 32 |
| 794 | RF01252 snR5 | 4.8 (2/42) | 1.9 | 100.0 (2/2) | 4.1 | 76.2 | 11 |
| 795 | RF00220 Rhino CRE | 4.8 (1/21) | 1.9 | 50.0 (1/2) | 3.8 | 81.4 | 12 |
| 796 | RF00043 Plasmid R1162 | 4.8 (1/21) | 1.4 | 100.0 (1/1) | 2.4 | 75.9 | 8 |
| 797 | RF01819 RsaD | 4.8 (2/42) | 1.2 | 100.0 (2/2) | 2.1 | 84.7 | 8 |
| 798 | RF01815 rpsB | 4.8 (1/21) | 0.5 | 100.0 (1/1) | 3.7 | 41.4 | 4 |
| 799 | RF02832 Scr5676 | 4.8 (2/42) | 0.2 | 100.0 (2/2) | 1.0 | 86.5 | 5 |
| 800 | RF01014 mir-1306 | 4.8 (1/21) | 0.0 | 100.0 (1/1) | 0.7 | 89.5 | 8 |
| 801 | RF00512 Leu leader | 4.7 (2/43) | 1.2 | 100.0 (2/2) | 3.4 | 64.1 | 6 |
| 802 | RF00647 MIR164 | 4.5 (1/22) | 9.5 | 100.0 (1/1) | 10.4 | 61.1 | 39 |
| 803 | RF00407 SNORA50 | 4.5 (1/22) | 3.2 | 100.0 (1/1) | 5.5 | 72.0 | 14 |
| 804 | RF01719 Pseudomon-1 | 4.5 (1/22) | 3.2 | 100.0 (1/1) | 4.4 | 89.0 | 19 |
| 805 | RF00052 lin-4 | 4.5 (1/22) | 2.7 | 100.0 (1/1) | 4.4 | 69.1 | 12 |
| 806 | RF00518 speF | 4.5 (1/22) | 1.8 | 100.0 (1/1) | 3.7 | 59.3 | 6 |
| 807 | RF02464 Ms AS-4 | 4.5 (1/22) | 0.9 | 100.0 (1/1) | 3.0 | 79.1 | 9 |
| 808 | RF02885 SAM VI | 4.5 (1/22) | 0.9 | 100.0 (1/1) | 2.0 | 85.5 | 7 |
| 809 | RF01077 TLS-PK2 | 4.5 (1/22) | 0.5 | 100.0 (1/1) | 1.0 | 82.8 | 4 |
| 810 | RF01020 mir-572 | 4.5 (1/22) | 0.0 | 100.0 (1/1) | 0.1 | 90.2 | 4 |
| 811 | RF02772 AilA thermometer | 4.5 (1/22) | 0.0 | 100.0 (1/1) | 0.7 | 84.9 | 5 |
| 812 | RF01413 miR-430 | 4.3 (1/23) | 22.2 | 100.0 (1/1) | 21.5 | 59.8 | 46 |
| 813 | RF02935 che1 | 4.3 (1/23) | 16.9 | 100.0 (1/1) | 16.5 | 82.1 | 98 |
| 814 | RF00103 mir-1 | 4.3 (1/23) | 16.5 | 100.0 (1/1) | 16.9 | 68.6 | 66 |
| 815 | RF00640 MIR167 1 | 4.3 (1/23) | 15.7 | 100.0 (1/1) | 15.7 | 57.5 | 42 |
| 816 | RF00649 mir-128 | 4.3 (1/23) | 3.9 | 100.0 (1/1) | 4.5 | 73.3 | 18 |
| 817 | RF02688 RAGATH-13 | 4.3 (1/23) | 2.2 | 50.0 (1/2) | 4.2 | 74.1 | 20 |
| 818 | RF01042 mir-891 | 4.3 (1/23) | 0.4 | 100.0 (1/1) | 1.3 | 84.6 | 7 |
| 819 | RF02989 gntR-DTE | 4.3 (1/23) | 0.4 | 100.0 (1/1) | 2.5 | 75.0 | 7 |
| 820 | RF01246 snR81 | 4.3 (2/46) | 0.0 | 100.0 (2/2) | 0.2 | 88.5 | 3 |
| 821 | RF00250 mir-TAR | 4.2 (1/24) | 10.8 | 100.0 (1/1) | 12.7 | 84.7 | 45 |
| 822 | RF00420 SNORA61 | 4.2 (1/24) | 6.2 | 100.0 (1/1) | 8.2 | 73.8 | 25 |
| 823 | RF00724 mir-282 | 4.2 (1/24) | 2.5 | 100.0 (1/1) | 3.5 | 67.9 | 15 |
| 824 | RF01224 snoR80 | 4.2 (1/24) | 2.5 | 50.0 (1/2) | 3.6 | 70.9 | 9 |
| 825 | RF00073 mir-156 | 4.2 (1/24) | 1.7 | 100.0 (1/1) | 2.2 | 73.1 | 10 |
| 826 | RF01260 snR11 | 4.2 (3/71) | 0.6 | 100.0 (3/3) | 2.8 | 76.4 | 6 |
| 827 | RF01239 snR49 | 4.2 (2/47) | 0.6 | 100.0 (2/2) | 2.3 | 76.0 | 6 |
| 828 | RF02869 ncS025 | 4.2 (1/24) | 0.0 | 100.0 (1/1) | 0.5 | 93.5 | 4 |
| 829 | RF02082 STnc540 | 4.2 (2/47) | 0.0 | 100.0 (2/2) | 0.4 | 90.9 | 3 |

Continued on next page

| | RNA family (seed alignment) | Sensitivity annotated bpairs that covary % (cov_bps/bps) | Power average power % | Positive Predictive Value covarying pairs in structure % (cov_bps/cov_pairs) | average substitutions per bpair | avg pairwise identity % | number of sequences |
|---|---|---|---|---|---|---|---|
| 830 | RF00115 McaS | 4.2 (1/24) | 0.0 | 50.0 (1/2) | 1.1 | 83.7 | 4 |
| 831 | RF01808 MicX | 4.1 (2/49) | 1.6 | 100.0 (2/2) | 3.3 | 74.7 | 10 |
| 832 | RF00061 IRES HCV | 4.0 (3/75) | 33.6 | 100.0 (3/3) | 37.0 | 87.3 | 79 |
| 833 | RF00042 CopA | 4.0 (1/25) | 7.2 | 100.0 (1/1) | 7.1 | 80.4 | 37 |
| 834 | RF01899 mir-2241 | 4.0 (1/25) | 0.8 | 100.0 (1/1) | 2.7 | 73.9 | 10 |
| 835 | RF00846 mir-64 | 4.0 (1/25) | 0.4 | 100.0 (1/1) | 1.5 | 58.1 | 4 |
| 836 | RF00830 mir-74 | 4.0 (1/25) | 0.0 | 100.0 (1/1) | 1.0 | 66.4 | 4 |
| 837 | RF01242 snR36 | 3.9 (2/51) | 3.1 | 100.0 (2/2) | 4.9 | 70.4 | 10 |
| 838 | RF02842 RefA1 | 3.9 (2/51) | 0.0 | 100.0 (2/2) | 0.9 | 91.2 | 4 |
| 839 | RF01769 greA | 3.8 (1/26) | 9.2 | 100.0 (1/1) | 10.2 | 72.8 | 25 |
| 840 | RF00413 SNORA19 | 3.8 (1/26) | 6.5 | 100.0 (1/1) | 7.6 | 79.0 | 34 |
| 841 | RF00655 mir-28 | 3.8 (1/26) | 5.4 | 100.0 (1/1) | 6.5 | 77.0 | 29 |
| 842 | RF00716 mir-3 | 3.8 (1/26) | 3.8 | 50.0 (1/2) | 5.4 | 70.0 | 11 |
| 843 | RF00248 mir-148 | 3.8 (1/26) | 0.8 | 100.0 (1/1) | 1.8 | 72.5 | 5 |
| 844 | RF00750 mir-458 | 3.8 (1/26) | 0.4 | 100.0 (1/1) | 1.2 | 81.0 | 7 |
| 845 | RF00890 mir-668 | 3.8 (1/26) | 0.4 | 100.0 (1/1) | 1.4 | 89.2 | 6 |
| 846 | RF02489 GlsR26 | 3.8 (1/26) | 0.0 | 100.0 (1/1) | 1.0 | 84.0 | 3 |
| 847 | RF02725 sno ZL8 | 3.8 (1/26) | 0.0 | 100.0 (1/1) | 0.7 | 91.0 | 6 |
| 848 | RF00392 SNORA5 | 3.8 (1/26) | 0.0 | 100.0 (1/1) | 1.6 | 75.8 | 6 |
| 849 | RF00225 IRES Tobamo | 3.8 (1/26) | 0.0 | 100.0 (1/1) | 0.9 | 87.5 | 7 |
| 850 | RF01675 CrcZ | 3.7 (2/54) | 9.4 | 100.0 (2/2) | 9.7 | 68.5 | 19 |
| 851 | RF00403 SNORA41 | 3.7 (1/27) | 9.2 | 100.0 (1/1) | 10.3 | 77.7 | 31 |
| 852 | RF00510 Tombus IRE | 3.7 (1/27) | 7.8 | 100.0 (1/1) | 9.3 | 84.3 | 23 |
| 853 | RF01255 snR35 | 3.7 (2/54) | 4.1 | 100.0 (2/2) | 5.7 | 76.4 | 11 |
| 854 | RF02370 Trp leader 2 | 3.7 (1/27) | 1.9 | 100.0 (1/1) | 3.5 | 75.4 | 9 |
| 855 | RF00062 HgcC | 3.7 (1/27) | 1.5 | 25.0 (1/4) | 3.3 | 69.7 | 5 |
| 856 | RF00256 mir-196 | 3.7 (1/27) | 1.5 | 100.0 (1/1) | 3.4 | 73.7 | 14 |
| 857 | RF00725 mir-iab-4 | 3.7 (1/27) | 0.7 | 100.0 (1/1) | 1.6 | 85.0 | 8 |
| 858 | RF02378 SurC | 3.7 (2/54) | 0.6 | 100.0 (2/2) | 1.4 | 85.3 | 4 |
| 859 | RF02366 Yfr19 | 3.7 (1/27) | 0.0 | 100.0 (1/1) | 0.7 | 86.8 | 6 |
| 860 | RF01919 mir-1419 | 3.7 (1/27) | 0.0 | 100.0 (1/1) | 1.1 | 79.3 | 5 |
| 861 | RF01492 rli28 | 3.6 (1/28) | 18.9 | 100.0 (1/1) | 18.9 | 55.5 | 21 |
| 862 | RF00130 mir-192 | 3.6 (1/28) | 11.1 | 100.0 (1/1) | 11.7 | 68.0 | 41 |
| 863 | RF00950 mir-927 | 3.6 (1/28) | 5.3 | 100.0 (1/1) | 5.7 | 77.2 | 14 |
| 864 | RF00286 SCARNA8 | 3.6 (1/28) | 3.9 | 100.0 (1/1) | 5.8 | 79.5 | 22 |
| 865 | RF01233 snoU109 | 3.6 (1/28) | 3.6 | 100.0 (1/1) | 4.9 | 83.2 | 26 |
| 866 | RF00717 mir-315 | 3.6 (1/28) | 2.9 | 50.0 (1/2) | 4.6 | 75.9 | 16 |
| 867 | RF01850 beta tmRNA | 3.6 (2/55) | 2.5 | 100.0 (2/2) | 4.8 | 71.1 | 7 |
| 868 | RF00418 SNORA58 | 3.6 (1/28) | 2.5 | 100.0 (1/1) | 4.3 | 80.9 | 20 |
| 869 | RF00082 SraG | 3.6 (1/28) | 2.1 | 100.0 (1/1) | 2.9 | 71.7 | 7 |
| 870 | RF01039 mir-937 | 3.6 (1/28) | 1.1 | 100.0 (1/1) | 3.0 | 73.8 | 9 |
| 871 | RF02547 mtPerm-5S | 3.6 (1/28) | 0.3 | 100.0 (1/1) | 1.3 | 84.4 | 6 |
| 872 | RF01087 PK-repZ | 3.6 (1/28) | 0.3 | 100.0 (1/1) | 1.1 | 89.5 | 6 |
| 873 | RF01788 drz-agam-2-2 | 3.6 (2/56) | 0.2 | 100.0 (2/2) | 1.6 | 71.8 | 5 |
| 874 | RF00927 mir-582 | 3.6 (1/28) | 0.0 | 100.0 (1/1) | 0.4 | 91.6 | 7 |
| 875 | RF00064 HgcG | 3.5 (2/57) | 0.2 | 66.7 (2/3) | 2.2 | 68.4 | 5 |
| 876 | RF00434 BTE | 3.4 (1/29) | 12.4 | 100.0 (1/1) | 12.4 | 72.9 | 17 |
| 877 | RF01496 Afu 182 | 3.4 (1/29) | 8.6 | 100.0 (1/1) | 9.5 | 65.3 | 19 |
| 878 | RF00842 MIR403 | 3.4 (1/29) | 3.1 | 100.0 (1/1) | 4.6 | 67.3 | 13 |
| 879 | RF02728 HrrF | 3.4 (1/29) | 1.4 | 100.0 (1/1) | 2.2 | 82.0 | 7 |
| 880 | RF02779 PepN thermometer | 3.4 (1/29) | 0.3 | 100.0 (1/1) | 1.4 | 81.8 | 5 |
| 881 | RF00765 mir-337 | 3.4 (1/29) | 0.3 | 100.0 (1/1) | 1.1 | 83.8 | 6 |
| 882 | RF01056 Mg sensor | 3.4 (1/29) | 0.0 | 100.0 (1/1) | 0.9 | 75.7 | 4 |
| 883 | RF00809 mir-241 | 3.4 (1/29) | 0.0 | 50.0 (1/2) | 0.8 | 76.0 | 4 |
| 884 | RF01517 iscRS | 3.4 (1/29) | 0.0 | 100.0 (1/1) | 0.9 | 87.3 | 4 |
| 885 | RF01853 mtDNA ssA | 3.3 (1/30) | 22.0 | 100.0 (1/1) | 21.6 | 67.6 | 53 |
| 886 | RF00892 mir-551 | 3.3 (1/30) | 8.0 | 100.0 (1/1) | 8.5 | 73.3 | 20 |
| 887 | RF00650 mir-153 | 3.3 (1/30) | 3.0 | 100.0 (1/1) | 4.4 | 78.6 | 18 |
| 888 | RF00730 mir-277 | 3.3 (1/30) | 2.7 | 100.0 (1/1) | 4.2 | 69.1 | 12 |
| 889 | RF01997 mir-969 | 3.3 (1/30) | 1.3 | 100.0 (1/1) | 2.4 | 76.1 | 8 |
| 890 | RF00768 MIR405 | 3.3 (1/30) | 0.7 | 100.0 (1/1) | 2.2 | 78.1 | 13 |
| 891 | RF00760 mir-342 | 3.3 (1/30) | 0.7 | 100.0 (1/1) | 1.1 | 85.9 | 10 |
| 892 | RF02450 ncr1175 | 3.3 (1/30) | 0.0 | 100.0 (1/1) | 0.6 | 86.3 | 4 |
| 893 | RF02092 mir-2970 | 3.3 (1/30) | 0.0 | 100.0 (1/1) | 0.6 | 89.5 | 5 |
| 894 | RF00774 mir-360 | 3.3 (1/30) | 0.0 | 100.0 (1/1) | 0.9 | 67.4 | 5 |
| 895 | RF00672 mir-190 | 3.2 (1/31) | 16.1 | 50.0 (1/2) | 15.9 | 68.0 | 29 |
| 896 | RF00628 RgsA | 3.2 (1/31) | 11.3 | 100.0 (1/1) | 11.9 | 73.3 | 27 |
| 897 | RF02994 IMPDH | 3.2 (1/31) | 7.7 | 50.0 (1/2) | 9.0 | 81.6 | 81 |
| 898 | RF00629 P24 | 3.2 (2/63) | 5.1 | 100.0 (2/2) | 6.1 | 75.5 | 14 |
| 899 | RF01916 mir-988 | 3.2 (1/31) | 0.3 | 50.0 (1/2) | 1.7 | 77.2 | 4 |
| 900 | RF00826 mir-55 | 3.2 (1/31) | 0.3 | 100.0 (1/1) | 1.4 | 72.6 | 5 |
| 901 | RF00936 mir-744 | 3.2 (1/31) | 0.0 | 100.0 (1/1) | 0.4 | 94.2 | 5 |
| 902 | RF00833 mir-70 | 3.2 (1/31) | 0.0 | 100.0 (1/1) | 1.2 | 58.0 | 4 |
| 903 | RF00825 mir-344 | 3.1 (1/32) | 20.3 | 50.0 (1/2) | 20.6 | 67.2 | 35 |
| 904 | RF01267 snR37 | 3.1 (4/128) | 2.8 | 100.0 (4/4) | 4.8 | 72.0 | 9 |
| 905 | RF00302 SNORA65 | 3.1 (1/32) | 2.5 | 50.0 (1/2) | 5.2 | 72.8 | 14 |
| 906 | RF00698 mir-489 | 3.1 (1/32) | 1.9 | 100.0 (1/1) | 4.2 | 79.1 | 19 |
| 907 | RF02094 mir-1803 | 3.1 (1/32) | 0.9 | 100.0 (1/1) | 2.6 | 66.3 | 6 |
| 908 | RF01230 snoR77 | 3.1 (1/32) | 0.6 | 100.0 (1/1) | 2.1 | 73.9 | 5 |
| 909 | RF00828 mir-75 | 3.1 (1/32) | 0.3 | 100.0 (1/1) | 0.9 | 71.6 | 4 |
| 910 | RF00568 SNORA26 | 3.0 (1/33) | 24.8 | 100.0 (1/1) | 24.5 | 78.2 | 76 |
| 911 | RF01851 cyano tmRNA | 3.0 (2/67) | 10.4 | 100.0 (2/2) | 11.2 | 83.2 | 27 |
| 912 | RF00503 RNAIII | 3.0 (4/134) | 6.6 | 100.0 (4/4) | 8.8 | 62.2 | 10 |
| 913 | RF00625 P11 | 3.0 (1/33) | 4.8 | 100.0 (1/1) | 5.7 | 69.0 | 15 |

Continued on next page

| | RNA family (seed alignment) | Sensitivity annotated bpairs that covary % (cov_bps/bps) | Power average power % | Positive Predictive Value covarying pairs in structure % (cov_bps/cov_pairs) | average substitutions per bpair | avg pairwise identity % | number of sequences |
|---|---|---|---|---|---|---|---|
| 914 | RF00957 mir-663 | 3.0 (1/33) | 3.6 | 100.0 (1/1) | 5.4 | 74.5 | 12 |
| 915 | RF00767 mir-150 | 3.0 (1/33) | 3.0 | 100.0 (1/1) | 4.6 | 80.5 | 17 |
| 916 | RF02535 ODC IRES | 3.0 (1/33) | 3.0 | 100.0 (1/1) | 5.0 | 79.7 | 13 |
| 917 | RF00858 mir-306 | 3.0 (1/33) | 1.2 | 100.0 (1/1) | 3.0 | 70.2 | 7 |
| 918 | RF00653 mir-22 | 3.0 (1/33) | 0.9 | 100.0 (1/1) | 2.5 | 76.9 | 12 |
| 919 | RF02899 AaHKsRNA54 | 3.0 (1/33) | 0.3 | 100.0 (1/1) | 1.7 | 76.9 | 6 |
| 920 | RF01633 ceN43 | 3.0 (1/33) | 0.3 | 100.0 (1/1) | 1.2 | 86.7 | 4 |
| 921 | RF00773 mir-298 | 3.0 (1/33) | 0.3 | 100.0 (1/1) | 2.7 | 71.3 | 5 |
| 922 | RF01910 mir-506 | 3.0 (1/33) | 0.3 | 100.0 (1/1) | 0.8 | 92.5 | 8 |
| 923 | RF00786 mir-289 | 3.0 (1/33) | 0.0 | 100.0 (1/1) | 1.0 | 85.7 | 3 |
| 924 | RF02748 FtrB | 3.0 (1/33) | 0.0 | 100.0 (1/1) | 1.3 | 87.8 | 5 |
| 925 | RF01418 HIV POL-1 SL | 2.9 (1/34) | 10.0 | 100.0 (1/1) | 11.0 | 81.6 | 29 |
| 926 | RF00425 SNORA18 | 2.9 (1/34) | 9.1 | 100.0 (1/1) | 10.5 | 76.7 | 29 |
| 927 | RF00746 mir-454 | 2.9 (1/34) | 5.6 | 100.0 (1/1) | 6.4 | 66.7 | 17 |
| 928 | RF00398 SNORA15 | 2.9 (1/35) | 5.4 | 100.0 (1/1) | 7.0 | 81.1 | 22 |
| 929 | RF00565 SCARNA3 | 2.9 (1/34) | 3.8 | 100.0 (1/1) | 5.7 | 75.9 | 23 |
| 930 | RF00544 snopsi28S-3327 | 2.9 (1/35) | 3.4 | 100.0 (1/1) | 4.5 | 78.4 | 14 |
| 931 | RF00562 SNORA49 | 2.9 (1/34) | 3.2 | 100.0 (1/1) | 4.6 | 81.3 | 21 |
| 932 | RF00627 P15 | 2.9 (1/35) | 3.1 | 100.0 (1/1) | 4.3 | 79.4 | 22 |
| 933 | RF01029 mir-649 | 2.9 (1/35) | 0.8 | 100.0 (1/1) | 2.9 | 74.1 | 5 |
| 934 | RF01914 mir-932 | 2.9 (1/34) | 0.6 | 100.0 (1/1) | 2.4 | 64.9 | 5 |
| 935 | RF00448 IRES EBNA | 2.9 (1/35) | 0.3 | 100.0 (1/1) | 0.8 | 83.2 | 6 |
| 936 | RF00124 IS102 | 2.9 (1/35) | 0.3 | 100.0 (1/1) | 1.2 | 88.9 | 5 |
| 937 | RF00948 mir-996 | 2.9 (1/35) | 0.3 | 100.0 (1/1) | 1.4 | 77.7 | 5 |
| 938 | RF01269 snR80 | 2.9 (1/34) | 0.3 | 100.0 (1/1) | 1.1 | 87.1 | 6 |
| 939 | RF00764 mir-191 | 2.9 (1/34) | 0.0 | 100.0 (1/1) | 0.6 | 88.3 | 5 |
| 940 | RF00340 SNORA36 | 2.9 (1/34) | 0.0 | 100.0 (1/1) | 1.2 | 85.1 | 6 |
| 941 | RF03106 RT-11 | 2.9 (1/34) | 0.0 | 100.0 (1/1) | 1.3 | 70.5 | 3 |
| 942 | RF03114 RT-1 | 2.9 (1/35) | 0.0 | 100.0 (1/1) | 1.0 | 87.7 | 5 |
| 943 | RF00995 mir-616 | 2.9 (1/34) | 0.0 | 100.0 (1/1) | 0.8 | 79.0 | 3 |
| 944 | RF00388 Anti-Q RNA | 2.9 (1/34) | 0.0 | 100.0 (1/1) | 1.9 | 72.7 | 6 |
| 945 | RF00761 mir-340 | 2.8 (1/36) | 22.5 | 100.0 (1/1) | 21.9 | 64.0 | 31 |
| 946 | RF00614 SNORA11 | 2.8 (1/36) | 8.1 | 100.0 (1/1) | 9.6 | 82.8 | 35 |
| 947 | RF00106 RNAI | 2.8 (1/36) | 2.5 | 100.0 (1/1) | 3.8 | 74.9 | 10 |
| 948 | RF01655 ceN84 | 2.8 (1/36) | 1.4 | 100.0 (1/1) | 3.7 | 74.5 | 5 |
| 949 | RF00827 mir-77 | 2.8 (1/36) | 0.3 | 100.0 (1/1) | 1.2 | 78.5 | 5 |
| 950 | RF00785 mir-90 | 2.8 (1/36) | 0.3 | 100.0 (1/1) | 1.2 | 76.3 | 5 |
| 951 | RF00449 IRES HIF1 | 2.7 (1/37) | 6.2 | 50.0 (1/2) | 8.0 | 75.9 | 17 |
| 952 | RF01656 ceN72-3 ceN74-2 | 2.7 (1/37) | 5.7 | 100.0 (1/1) | 7.4 | 70.9 | 13 |
| 953 | RF00397 SNORA14 | 2.7 (1/37) | 4.9 | 100.0 (1/1) | 6.5 | 77.8 | 18 |
| 954 | RF00409 SNORA7 | 2.7 (1/37) | 3.8 | 50.0 (1/2) | 5.5 | 78.6 | 31 |
| 955 | RF00415 SNORA30 | 2.7 (1/37) | 3.0 | 100.0 (1/1) | 5.2 | 72.6 | 11 |
| 956 | RF00077 SraB | 2.7 (1/37) | 0.8 | 100.0 (1/1) | 1.8 | 85.6 | 5 |
| 957 | RF00822 mir-274 | 2.7 (1/37) | 0.8 | 100.0 (1/1) | 1.7 | 81.6 | 7 |
| 958 | RF02714 snopsi28S-3378 | 2.7 (1/37) | 0.0 | 100.0 (1/1) | 1.1 | 85.9 | 8 |
| 959 | RF01649 ceN67 | 2.7 (1/37) | 0.0 | 100.0 (1/1) | 0.9 | 65.1 | 3 |
| 960 | RF02356 BjrC1505 | 2.6 (1/39) | 14.6 | 100.0 (1/1) | 13.9 | 76.0 | 25 |
| 961 | RF01229 SNORA84 | 2.6 (1/39) | 8.7 | 100.0 (1/1) | 9.5 | 75.4 | 28 |
| 962 | RF02471 Ms IGR-5 | 2.6 (1/39) | 8.2 | 100.0 (1/1) | 8.8 | 73.6 | 21 |
| 963 | RF00600 SNORA79 | 2.6 (1/38) | 5.0 | 100.0 (1/1) | 6.9 | 78.9 | 25 |
| 964 | RF00561 SNORA40 | 2.6 (1/39) | 4.1 | 100.0 (1/1) | 5.9 | 83.6 | 28 |
| 965 | RF02235 asX1 | 2.6 (1/38) | 3.1 | 100.0 (1/1) | 5.0 | 81.1 | 15 |
| 966 | RF02079 STnc180 | 2.6 (1/39) | 1.3 | 100.0 (1/1) | 3.5 | 64.0 | 10 |
| 967 | RF00128 GlmY tke1 | 2.6 (1/38) | 1.1 | 100.0 (1/1) | 2.3 | 76.8 | 17 |
| 968 | RF01664 ceN101 | 2.6 (1/39) | 0.8 | 50.0 (1/2) | 1.4 | 87.7 | 4 |
| 969 | RF00401 SNORA20 | 2.6 (1/39) | 0.5 | 100.0 (1/1) | 2.3 | 87.1 | 17 |
| 970 | RF02774 KatA thermometer | 2.6 (1/38) | 0.5 | 100.0 (1/1) | 1.5 | 84.8 | 5 |
| 971 | RF01653 ceN80 | 2.6 (1/39) | 0.2 | 100.0 (1/1) | 0.9 | 80.5 | 5 |
| 972 | RF01408 sraL | 2.6 (1/38) | 0.0 | 100.0 (1/1) | 1.3 | 81.8 | 6 |
| 973 | RF01440 S pombe snR42 | 2.6 (1/38) | 0.0 | 100.0 (1/1) | 1.4 | 73.7 | 3 |
| 974 | RF02666 PsiU2-35.45 | 2.6 (1/38) | 0.0 | 100.0 (1/1) | 0.8 | 84.3 | 3 |
| 975 | RF01925 MIR1428 | 2.6 (1/39) | 0.0 | 100.0 (1/1) | 0.5 | 90.3 | 5 |
| 976 | RF00334 SNORA3 | 2.5 (1/40) | 0.8 | 50.0 (1/2) | 2.1 | 74.4 | 6 |
| 977 | RF01259 snR63 | 2.5 (2/80) | 0.2 | 100.0 (2/2) | 1.0 | 93.1 | 6 |
| 978 | RF00888 mir-770 | 2.5 (1/40) | 0.0 | 100.0 (1/1) | 1.4 | 71.2 | 4 |
| 979 | RF02485 GlsR22 | 2.5 (1/40) | 0.0 | 100.0 (1/1) | 0.5 | 86.4 | 3 |
| 980 | RF02036 IMES-3 | 2.4 (1/42) | 15.7 | 100.0 (1/1) | 14.8 | 81.6 | 32 |
| 981 | RF00779 MIR474 | 2.4 (1/41) | 8.3 | 100.0 (1/1) | 10.3 | 55.9 | 11 |
| 982 | RF01766 cspA | 2.4 (3/124) | 5.4 | 100.0 (3/3) | 7.3 | 74.5 | 15 |
| 983 | RF00470 Toga 5 CRE | 2.4 (1/42) | 5.2 | 100.0 (1/1) | 5.9 | 81.0 | 33 |
| 984 | RF01405 STnc490 | 2.4 (1/41) | 3.6 | 100.0 (1/1) | 4.8 | 94.0 | 76 |
| 985 | RF01671 P18 | 2.4 (1/41) | 0.2 | 50.0 (1/2) | 1.7 | 74.6 | 4 |
| 986 | RF01663 ceN93 | 2.4 (1/41) | 0.0 | 100.0 (1/1) | 0.6 | 84.0 | 4 |
| 987 | RF02363 Yfr11 | 2.4 (1/41) | 0.0 | 50.0 (1/2) | 2.0 | 66.7 | 4 |
| 988 | RF00235 Plasmid RNAIII | 2.4 (1/41) | 0.0 | 100.0 (1/1) | 0.6 | 94.0 | 7 |
| 989 | RF00560 SNORA17 | 2.3 (1/44) | 16.8 | 100.0 (1/1) | 17.0 | 79.6 | 53 |
| 990 | RF00100 7SK | 2.3 (2/87) | 11.6 | 100.0 (2/2) | 12.6 | 81.7 | 45 |
| 991 | RF02355 BjrC174 | 2.3 (1/43) | 10.9 | 100.0 (1/1) | 11.3 | 75.6 | 15 |
| 992 | RF02936 Chloroflexus-1 | 2.3 (1/43) | 0.0 | 33.3 (1/3) | 0.9 | 79.0 | 4 |
| 993 | RF01434 S pombe snR3 | 2.3 (1/43) | 0.0 | 100.0 (1/1) | 1.5 | 78.2 | 3 |
| 994 | RF02462 Telomerase Asco | 2.2 (2/89) | 0.8 | 50.0 (2/4) | 2.1 | 69.9 | 4 |
| 995 | RF02376 SR1 | 2.2 (1/45) | 0.4 | 100.0 (1/1) | 2.7 | 74.2 | 6 |
| 996 | RF00552 rncO | 2.1 (1/47) | 4.2 | 100.0 (1/1) | 5.9 | 83.7 | 18 |
| 997 | RF02970 EGFOA-assoc-1 | 2.1 (1/47) | 0.6 | 100.0 (1/1) | 3.0 | 80.7 | 10 |

Continued on next page

| | RNA family (seed alignment) | Sensitivity annotated bpairs that covary % (cov_bps/bps) | Power average power % | Positive Predictive Value covarying pairs in structure % (cov_bps/cov_pairs) | average substitutions per bpair | avg pairwise identity % | number of sequences |
|---|---|---|---|---|---|---|---|
| 998 | RF01476 rliF | 2.1 (1/47) | 0.4 | 100.0 (1/1) | 1.4 | 76.0 | 5 |
| 999 | RF01005 MIR530 | 2.1 (1/47) | 0.0 | 100.0 (1/1) | 0.4 | 73.6 | 3 |
| 1000 | RF01581 RUF6-5 | 2.0 (1/51) | 7.6 | 100.0 (1/1) | 9.1 | 82.1 | 20 |
| 1001 | RF01452 S pombe snR99 | 2.0 (1/50) | 0.6 | 100.0 (1/1) | 1.9 | 84.5 | 5 |
| 1002 | RF02792 sRNA162 | 2.0 (1/50) | 0.0 | 50.0 (1/2) | 1.0 | 67.4 | 3 |
| 1003 | RF01789 EBER1 | 2.0 (1/51) | 0.0 | 100.0 (1/1) | 1.0 | 76.5 | 3 |
| 1004 | RF02870 ncS035 | 2.0 (1/51) | 0.0 | 100.0 (1/1) | 0.8 | 77.4 | 3 |
| 1005 | RF02081 STnc550 | 1.9 (1/53) | 1.3 | 50.0 (1/2) | 3.3 | 79.0 | 11 |
| 1006 | RF00125 IS128 | 1.9 (1/52) | 1.1 | 100.0 (1/1) | 2.0 | 87.7 | 5 |
| 1007 | RF02695 NrsZ | 1.9 (1/53) | 0.9 | 100.0 (1/1) | 2.4 | 80.1 | 4 |
| 1008 | RF01253 snR46 | 1.8 (1/55) | 0.2 | 100.0 (1/1) | 1.5 | 79.9 | 6 |
| 1009 | RF02876 AS-pc02 | 1.8 (1/57) | 0.0 | 100.0 (1/1) | 0.5 | 92.5 | 4 |
| 1010 | RF00231 SCARNA13 | 1.6 (1/62) | 3.7 | 100.0 (1/1) | 5.5 | 78.9 | 24 |
| 1011 | RF02687 RAGATH-8 | 1.6 (1/63) | 0.0 | 100.0 (1/1) | 0.4 | 91.6 | 3 |
| 1012 | RF00090 SNORA74 | 1.5 (1/67) | 7.5 | 100.0 (1/1) | 9.0 | 77.8 | 24 |
| 1013 | RF02423 Bp1 781 | 1.5 (1/67) | 3.3 | 100.0 (1/1) | 4.7 | 74.8 | 15 |
| 1014 | RF02442 SpF66 sRNA | 1.4 (1/71) | 1.0 | 100.0 (1/1) | 2.9 | 84.5 | 9 |
| 1015 | RF02532 MNV 3UTR | 1.4 (1/74) | 0.3 | 50.0 (1/2) | 0.9 | 91.9 | 10 |
| 1016 | RF02691 BTH s19 | 1.4 (1/70) | 0.1 | 100.0 (1/1) | 1.6 | 81.5 | 7 |
| 1017 | RF02757 Erse | 1.4 (1/74) | 0.1 | 100.0 (1/1) | 1.2 | 88.2 | 6 |
| 1018 | RF00563 SNORA53 | 1.3 (1/75) | 6.5 | 100.0 (1/1) | 8.6 | 80.1 | 28 |
| 1019 | RF00551 bicoid 3 | 1.3 (2/156) | 6.0 | 100.0 (2/2) | 7.3 | 77.7 | 15 |
| 1020 | RF00040 rne5 | 1.2 (1/84) | 0.8 | 100.0 (1/1) | 2.9 | 65.6 | 6 |
| 1021 | RF01390 isrG | 1.1 (1/90) | 0.4 | 100.0 (1/1) | 1.7 | 85.6 | 5 |
| 1022 | RF02858 Arrc08 | 1.1 (1/95) | 0.1 | 100.0 (1/1) | 1.1 | 76.6 | 3 |
| 1023 | RF01570 Dictyostelium SRP | 1.1 (1/92) | 0.1 | 100.0 (1/1) | 1.3 | 71.9 | 3 |
| 1024 | RF01670 P17 | 1.1 (1/88) | 0.0 | 100.0 (1/1) | 0.4 | 92.0 | 3 |
| 1025 | RF00885 MIR821 | 0.9 (1/113) | 0.1 | 100.0 (1/1) | 0.4 | 88.6 | 5 |
| 1026 | RF02737 Rev13 | 0.9 (1/112) | 0.0 | 50.0 (1/2) | 0.6 | 79.3 | 3 |
| 1027 | RF01473 rli41 | 0.8 (1/128) | 0.1 | 50.0 (1/2) | 0.4 | 93.1 | 6 |
| 1028 | RF01491 rli54 | 0.7 (1/146) | 0.0 | 100.0 (1/1) | 0.2 | 95.5 | 5 |
| 1029 | RF01270 snR84 | 0.6 (1/157) | 0.7 | 100.0 (1/1) | 2.2 | 79.6 | 6 |
| 1030 | RF01050 Sacc telomerase | 0.5 (1/191) | 4.0 | 100.0 (1/1) | 5.3 | 73.9 | 13 |
| 1031 | RF02576 tsr1 | 0.5 (1/204) | 0.1 | 100.0 (1/1) | 1.2 | 94.3 | 3 |
| 1032 | RF01419 IsrR | 0.0 (0/11) | 77.3 | 0.0 (0/3) | 157.2 | 66.9 | 308 |
| 1033 | RF01942 mir-1937 | 0.0 (0/11) | 66.4 | 0.0 (0/0) | 84.6 | 65.8 | 171 |
| 1034 | RF00523 Prion pknot | 0.0 (0/6) | 50.0 | 0.0 (0/0) | 53.0 | 85.0 | 148 |
| 1035 | RF00485 K chan RES | 0.0 (0/24) | 41.2 | 0.0 (0/3) | 45.1 | 68.7 | 85 |
| 1036 | RF00468 HCV SLVII | 0.0 (0/17) | 38.8 | 0.0 (0/0) | 42.9 | 74.3 | 110 |
| 1037 | RF00693 mir-147 | 0.0 (0/15) | 29.3 | 0.0 (0/0) | 28.5 | 68.5 | 64 |
| 1038 | RF00480 HIV FE | 0.0 (0/10) | 27.0 | 0.0 (0/0) | 32.2 | 84.0 | 145 |
| 1039 | RF00093 SNORD18 | 0.0 (0/4) | 22.5 | 0.0 (0/0) | 21.5 | 69.9 | 16 |
| 1040 | RF00376 HIV GSL3 | 0.0 (0/8) | 21.2 | 0.0 (0/0) | 22.0 | 81.4 | 72 |
| 1041 | RF01753 psbNH | 0.0 (0/12) | 20.8 | 0.0 (0/0) | 21.4 | 76.6 | 39 |
| 1042 | RF00469 HCV SLIV | 0.0 (0/15) | 20.7 | 0.0 (0/0) | 21.3 | 86.3 | 110 |
| 1043 | RF00047 mir-2 | 0.0 (0/21) | 20.0 | 0.0 (0/0) | 20.4 | 66.6 | 56 |
| 1044 | RF00550 HepE CRE | 0.0 (0/41) | 17.1 | 0.0 (0/0) | 16.7 | 84.2 | 46 |
| 1045 | RF00535 snoMe28S-Am982 | 0.0 (0/3) | 16.7 | 0.0 (0/0) | 16.3 | 76.2 | 13 |
| 1046 | RF00104 mir-10 | 0.0 (0/24) | 16.7 | 0.0 (0/0) | 16.6 | 68.1 | 36 |
| 1047 | RF00736 mir-320 | 0.0 (0/19) | 16.3 | 0.0 (0/0) | 17.1 | 68.0 | 55 |
| 1048 | RF00654 mir-216 | 0.0 (0/18) | 16.1 | 0.0 (0/0) | 15.8 | 61.5 | 33 |
| 1049 | RF00134 snoZ196 | 0.0 (0/7) | 15.7 | 0.0 (0/0) | 16.7 | 68.4 | 22 |
| 1050 | RF00490 S-element | 0.0 (0/22) | 15.4 | 0.0 (0/0) | 15.7 | 75.8 | 29 |
| 1051 | RF02510 PYLIS 3 | 0.0 (0/8) | 15.0 | 0.0 (0/0) | 16.5 | 63.0 | 23 |
| 1052 | RF02002 mir-720 | 0.0 (0/26) | 15.0 | 0.0 (0/0) | 15.1 | 81.2 | 35 |
| 1053 | RF02027 MIR2907 | 0.0 (0/16) | 15.0 | 0.0 (0/0) | 15.6 | 76.9 | 52 |
| 1054 | RF00651 mir-221 | 0.0 (0/21) | 14.8 | 0.0 (0/0) | 14.7 | 73.9 | 47 |
| 1055 | RF00041 Entero OriR | 0.0 (0/35) | 14.0 | 0.0 (0/0) | 14.1 | 88.0 | 60 |
| 1056 | RF01803 GABA3 | 0.0 (0/21) | 13.8 | 0.0 (0/1) | 14.0 | 84.5 | 52 |
| 1057 | RF00665 mir-290 | 0.0 (0/25) | 13.6 | 0.0 (0/0) | 13.7 | 65.9 | 27 |
| 1058 | RF01518 pRNA | 0.0 (0/22) | 13.6 | 0.0 (0/0) | 14.0 | 57.2 | 23 |
| 1059 | RF00424 SCARNA16 | 0.0 (0/54) | 13.0 | 0.0 (0/0) | 14.3 | 75.9 | 37 |
| 1060 | RF00451 mir-395 | 0.0 (0/30) | 12.7 | 0.0 (0/0) | 12.9 | 65.0 | 25 |
| 1061 | RF00679 mir-210 | 0.0 (0/27) | 12.6 | 0.0 (0/0) | 12.4 | 61.6 | 26 |
| 1062 | RF00670 mir-105 | 0.0 (0/29) | 12.4 | 0.0 (0/0) | 13.3 | 67.3 | 20 |
| 1063 | RF00357 snoR44 J54 | 0.0 (0/5) | 12.0 | 0.0 (0/0) | 11.2 | 72.1 | 29 |
| 1064 | RF02447 SpR19 sRNA | 0.0 (0/30) | 12.0 | 0.0 (0/1) | 13.0 | 74.8 | 23 |
| 1065 | RF01982 PYLIS 1 | 0.0 (0/15) | 11.3 | 0.0 (0/0) | 10.9 | 71.7 | 20 |
| 1066 | RF00639 mir-515 | 0.0 (0/19) | 11.1 | 0.0 (0/0) | 12.4 | 80.2 | 40 |
| 1067 | RF02516 mir-393 | 0.0 (0/29) | 11.0 | 0.0 (0/1) | 10.9 | 63.8 | 27 |
| 1068 | RF00691 mir-146 | 0.0 (0/17) | 10.6 | 0.0 (0/0) | 10.4 | 63.1 | 33 |
| 1069 | RF00034 RprA | 0.0 (0/18) | 10.6 | 0.0 (0/0) | 10.4 | 66.8 | 13 |
| 1070 | RF02031 tpke11 | 0.0 (0/16) | 10.6 | 0.0 (0/0) | 10.3 | 68.9 | 28 |
| 1071 | RF00446 mir-133 | 0.0 (0/20) | 10.5 | 0.0 (0/0) | 10.7 | 67.6 | 46 |
| 1072 | RF00311 snoZ188 | 0.0 (0/6) | 10.0 | 0.0 (0/0) | 12.3 | 78.8 | 21 |
| 1073 | RF01304 sR5 | 0.0 (0/3) | 10.0 | 0.0 (0/0) | 12.3 | 62.1 | 11 |
| 1074 | RF01214 snR51 | 0.0 (0/3) | 10.0 | 0.0 (0/0) | 12.3 | 76.1 | 17 |
| 1075 | RF01772 rnk pseudo | 0.0 (0/23) | 9.6 | 0.0 (0/0) | 10.2 | 57.3 | 15 |
| 1076 | RF02446 SpR18 sRNA | 0.0 (0/28) | 9.6 | 0.0 (0/0) | 10.8 | 77.6 | 23 |
| 1077 | RF02271 uc 338 | 0.0 (0/50) | 9.2 | 0.0 (0/0) | 10.5 | 76.1 | 34 |
| 1078 | RF00426 SCARNA15 | 0.0 (0/36) | 8.9 | 0.0 (0/0) | 10.4 | 72.1 | 22 |
| 1079 | RF02162 XIST A REPEAT | 0.0 (0/7) | 8.6 | 0.0 (0/0) | 8.7 | 84.1 | 54 |
| 1080 | RF02025 mir-3017 | 0.0 (0/23) | 8.3 | 0.0 (0/0) | 9.6 | 84.6 | 36 |
| 1081 | RF00246 mir-135 | 0.0 (0/28) | 8.2 | 0.0 (0/0) | 8.5 | 72.1 | 32 |

Continued on next page

| | RNA family (seed alignment) | Sensitivity annotated bpairs that covary % (cov_bps/bps) | Power average power % | Positive Predictive Value covarying pairs in structure % (cov_bps/cov_pairs) | average substitutions per bpair | avg pairwise identity % | number of sequences |
|---|---|---|---|---|---|---|---|
| 1082 | RF01697 Chlorobi-RRM | 0.0 (0/22) | 8.2 | 0.0 (0/0) | 9.5 | 77.5 | 18 |
| 1083 | RF01016 mir-584 | 0.0 (0/37) | 8.1 | 0.0 (0/0) | 9.4 | 63.3 | 16 |
| 1084 | RF01059 mir-598 | 0.0 (0/30) | 8.0 | 0.0 (0/0) | 9.4 | 68.9 | 24 |
| 1085 | RF00494 snoU2 19 | 0.0 (0/9) | 7.8 | 0.0 (0/0) | 9.8 | 79.8 | 29 |
| 1086 | RF00245 mir-19 | 0.0 (0/22) | 7.7 | 0.0 (0/0) | 8.1 | 81.8 | 46 |
| 1087 | RF00602 SCARNA21 | 0.0 (0/31) | 7.7 | 0.0 (0/0) | 9.3 | 75.0 | 24 |
| 1088 | RF01396 isrN | 0.0 (0/25) | 7.6 | 0.0 (0/0) | 9.9 | 61.4 | 17 |
| 1089 | RF01421 snoR114 | 0.0 (0/4) | 7.5 | 0.0 (0/0) | 8.8 | 64.9 | 24 |
| 1090 | RF00157 SNORD39 | 0.0 (0/4) | 7.5 | 0.0 (0/0) | 8.2 | 82.6 | 20 |
| 1091 | RF01791 F6 | 0.0 (0/20) | 7.5 | 0.0 (0/0) | 8.5 | 71.3 | 16 |
| 1092 | RF00582 SCARNA14 | 0.0 (0/38) | 7.4 | 0.0 (0/0) | 8.8 | 76.9 | 22 |
| 1093 | RF01268 SCARNA2 | 0.0 (0/88) | 7.4 | 0.0 (0/0) | 9.4 | 77.7 | 20 |
| 1094 | RF01797 fstAT | 0.0 (0/18) | 7.2 | 0.0 (0/1) | 9.1 | 75.9 | 22 |
| 1095 | RF01294 snoU89 | 0.0 (0/72) | 7.1 | 0.0 (0/0) | 8.4 | 75.3 | 16 |
| 1096 | RF00131 mir-30 | 0.0 (0/21) | 7.1 | 0.0 (0/0) | 7.4 | 75.2 | 49 |
| 1097 | RF00697 mir-186 | 0.0 (0/34) | 7.0 | 0.0 (0/0) | 8.6 | 75.0 | 19 |
| 1098 | RF00464 mir-92 | 0.0 (0/26) | 6.9 | 0.0 (0/0) | 7.2 | 76.2 | 23 |
| 1099 | RF02534 Noro CRE | 0.0 (0/13) | 6.9 | 0.0 (0/0) | 8.1 | 83.1 | 21 |
| 1100 | RF00402 SNORA25 | 0.0 (0/34) | 6.8 | 0.0 (0/0) | 8.7 | 77.6 | 30 |
| 1101 | RF00419 SNORA52 | 0.0 (0/31) | 6.8 | 0.0 (0/0) | 7.9 | 72.5 | 21 |
| 1102 | RF00506 Thr leader | 0.0 (0/35) | 6.8 | 0.0 (0/0) | 8.1 | 77.1 | 25 |
| 1103 | RF00723 mir-448 | 0.0 (0/30) | 6.7 | 0.0 (0/0) | 7.6 | 74.9 | 19 |
| 1104 | RF00144 mir-199 | 0.0 (0/30) | 6.7 | 0.0 (0/0) | 7.4 | 74.8 | 40 |
| 1105 | RF00749 mir-208 | 0.0 (0/26) | 6.5 | 0.0 (0/0) | 7.5 | 71.9 | 17 |
| 1106 | RF00599 SNORA77 | 0.0 (0/34) | 6.5 | 0.0 (0/0) | 7.7 | 75.1 | 19 |
| 1107 | RF00460 U1A PIE | 0.0 (0/14) | 6.4 | 0.0 (0/0) | 8.0 | 74.3 | 39 |
| 1108 | RF00237 mir-9 | 0.0 (0/19) | 6.3 | 0.0 (0/0) | 7.1 | 78.4 | 43 |
| 1109 | RF01512 Afu 309 | 0.0 (0/90) | 6.3 | 0.0 (0/0) | 8.0 | 64.3 | 10 |
| 1110 | RF00547 IRES TrkB | 0.0 (0/117) | 6.3 | 0.0 (0/0) | 8.5 | 74.4 | 16 |
| 1111 | RF00139 SNORA72 | 0.0 (0/30) | 6.3 | 0.0 (0/0) | 8.0 | 82.8 | 29 |
| 1112 | RF01673 PhrS | 0.0 (0/21) | 6.2 | 0.0 (0/0) | 8.3 | 64.8 | 13 |
| 1113 | RF00304 snoZ279 R105 R108 | 0.0 (0/26) | 6.2 | 0.0 (0/0) | 7.6 | 73.7 | 20 |
| 1114 | RF00567 SNORD17 | 0.0 (0/85) | 6.1 | 0.0 (0/0) | 7.9 | 78.3 | 19 |
| 1115 | RF00620 HCV ARF SL | 0.0 (0/46) | 6.1 | 0.0 (0/0) | 6.7 | 88.0 | 36 |
| 1116 | RF02084 STnc130 | 0.0 (0/10) | 6.0 | 0.0 (0/1) | 5.8 | 66.3 | 13 |
| 1117 | RF00206 U54 | 0.0 (0/5) | 6.0 | 0.0 (0/1) | 5.0 | 70.1 | 21 |
| 1118 | RF00586 SNORA12 | 0.0 (0/30) | 6.0 | 0.0 (0/0) | 7.3 | 77.1 | 23 |
| 1119 | RF00593 snoU83B | 0.0 (0/5) | 6.0 | 0.0 (0/0) | 9.6 | 81.6 | 21 |
| 1120 | RF01234 SNORA47 | 0.0 (0/31) | 5.8 | 0.0 (0/0) | 7.1 | 76.0 | 20 |
| 1121 | RF00478 SCARNA6 | 0.0 (0/57) | 5.6 | 0.0 (0/0) | 7.3 | 73.3 | 17 |
| 1122 | RF02000 MIR1846 | 0.0 (0/32) | 5.6 | 0.0 (0/0) | 7.5 | 77.3 | 21 |
| 1123 | RF00564 SCARNA11 | 0.0 (0/41) | 5.6 | 0.0 (0/0) | 7.4 | 79.8 | 24 |
| 1124 | RF00445 mir-399 | 0.0 (0/27) | 5.5 | 0.0 (0/0) | 6.5 | 59.4 | 13 |
| 1125 | RF00675 mir-145 | 0.0 (0/31) | 5.5 | 0.0 (0/0) | 5.6 | 79.3 | 13 |
| 1126 | RF00671 mir-138 | 0.0 (0/24) | 5.4 | 0.0 (0/0) | 5.1 | 79.6 | 24 |
| 1127 | RF01813 rdlD | 0.0 (0/17) | 5.3 | 0.0 (0/0) | 6.0 | 83.7 | 52 |
| 1128 | RF00400 SNORA28 | 0.0 (0/31) | 5.2 | 0.0 (0/0) | 7.0 | 79.7 | 26 |
| 1129 | RF00253 mir-101 | 0.0 (0/23) | 5.2 | 0.0 (0/0) | 5.6 | 82.9 | 24 |
| 1130 | RF00414 SNORA22 | 0.0 (0/39) | 5.1 | 0.0 (0/0) | 7.3 | 79.4 | 31 |
| 1131 | RF00440 SNORD37 | 0.0 (0/2) | 5.0 | 0.0 (0/0) | 7.0 | 78.9 | 6 |
| 1132 | RF01241 SNORA81 | 0.0 (0/36) | 5.0 | 0.0 (0/0) | 7.3 | 83.6 | 28 |
| 1133 | RF00656 mir-205 | 0.0 (0/24) | 5.0 | 0.0 (0/0) | 5.9 | 73.3 | 29 |
| 1134 | RF02419 Spd-sr37 | 0.0 (0/12) | 5.0 | 0.0 (0/1) | 6.1 | 79.6 | 25 |
| 1135 | RF00105 SNORD115 | 0.0 (0/8) | 5.0 | 0.0 (0/0) | 7.4 | 89.1 | 32 |
| 1136 | RF00429 SNORA29 | 0.0 (0/16) | 5.0 | 0.0 (0/0) | 6.2 | 79.4 | 26 |
| 1137 | RF00255 mir-218 | 0.0 (0/28) | 5.0 | 0.0 (0/0) | 5.1 | 77.3 | 18 |
| 1138 | RF01140 sR20 | 0.0 (0/2) | 5.0 | 0.0 (0/0) | 9.5 | 64.9 | 6 |
| 1139 | RF00455 mir-15 | 0.0 (0/22) | 5.0 | 0.0 (0/0) | 5.9 | 73.3 | 17 |
| 1140 | RF00097 snoR71 | 0.0 (0/8) | 5.0 | 0.0 (0/0) | 6.6 | 82.0 | 25 |
| 1141 | RF01781 ASdes | 0.0 (0/21) | 4.8 | 0.0 (0/0) | 7.0 | 80.1 | 26 |
| 1142 | RF00406 SNORA42 | 0.0 (0/35) | 4.8 | 0.0 (0/0) | 6.7 | 79.4 | 21 |
| 1143 | RF00998 mir-562 | 0.0 (0/34) | 4.7 | 0.0 (0/0) | 6.8 | 72.8 | 14 |
| 1144 | RF02375 Aar | 0.0 (0/49) | 4.7 | 0.0 (0/0) | 6.3 | 72.9 | 13 |
| 1145 | RF01743 leu-phe leader | 0.0 (0/34) | 4.7 | 0.0 (0/0) | 6.3 | 72.5 | 9 |
| 1146 | RF00143 mir-6 | 0.0 (0/24) | 4.6 | 0.0 (0/0) | 5.7 | 70.8 | 24 |
| 1147 | RF00416 SNORA43 | 0.0 (0/26) | 4.6 | 0.0 (0/0) | 6.0 | 75.8 | 21 |
| 1148 | RF01099 PK-IAV | 0.0 (0/11) | 4.5 | 0.0 (0/0) | 5.7 | 88.2 | 32 |
| 1149 | RF00708 mir-450 | 0.0 (0/29) | 4.5 | 0.0 (0/0) | 6.4 | 78.5 | 21 |
| 1150 | RF00732 mir-305 | 0.0 (0/31) | 4.5 | 0.0 (0/0) | 5.3 | 75.2 | 14 |
| 1151 | RF00267 snoR64 | 0.0 (0/11) | 4.5 | 0.0 (0/1) | 6.2 | 69.7 | 14 |
| 1152 | RF02432 SpF25 sRNA | 0.0 (0/34) | 4.4 | 0.0 (0/0) | 5.3 | 83.4 | 20 |
| 1153 | RF00644 mir-27 | 0.0 (0/18) | 4.4 | 0.0 (0/0) | 5.2 | 67.3 | 32 |
| 1154 | RF02506 Atu Ti1 | 0.0 (0/27) | 4.4 | 0.0 (0/0) | 6.1 | 72.6 | 14 |
| 1155 | RF00129 mir-103 | 0.0 (0/18) | 4.4 | 0.0 (0/0) | 4.8 | 84.9 | 28 |
| 1156 | RF02449 ncr1015 | 0.0 (0/23) | 4.3 | 0.0 (0/0) | 5.4 | 71.7 | 16 |
| 1157 | RF00328 snoZ161 228 | 0.0 (0/7) | 4.3 | 0.0 (0/0) | 7.3 | 67.6 | 11 |
| 1158 | RF00152 SNORD79 | 0.0 (0/7) | 4.3 | 0.0 (0/0) | 6.1 | 76.2 | 28 |
| 1159 | RF01045 mir-544 | 0.0 (0/30) | 4.3 | 0.0 (0/0) | 6.3 | 73.1 | 24 |
| 1160 | RF00241 mir-8 | 0.0 (0/21) | 4.3 | 0.0 (0/0) | 5.7 | 60.3 | 13 |
| 1161 | RF00527 snoMe28S-G3255 | 0.0 (0/7) | 4.3 | 0.0 (0/0) | 4.7 | 78.8 | 10 |
| 1162 | RF01380 HIV-1 SD | 0.0 (0/7) | 4.3 | 0.0 (0/0) | 4.3 | 93.6 | 22 |
| 1163 | RF00886 MIR807 | 0.0 (0/64) | 4.2 | 0.0 (0/0) | 5.9 | 85.0 | 30 |
| 1164 | RF00431 SNORA55 | 0.0 (0/31) | 4.2 | 0.0 (0/0) | 5.3 | 81.3 | 29 |
| 1165 | RF00076 mir-181 | 0.0 (0/24) | 4.2 | 0.0 (0/0) | 4.5 | 81.1 | 19 |

Continued on next page

| | RNA family (seed alignment) | Sensitivity annotated bpairs that covary % (cov_bps/bps) | Power average power % | Positive Predictive Value covarying pairs in structure % (cov_bps/cov_pairs) | average substitutions per bpair | avg pairwise identity % | number of sequences |
|---|---|---|---|---|---|---|---|
| 1166 | RF02526 SSRC34 1 | 0.0 (0/36) | 4.2 | 0.0 (0/1) | 5.7 | 83.4 | 16 |
| 1167 | RF00048 Entero CRE | 0.0 (0/12) | 4.2 | 0.0 (0/1) | 5.2 | 81.7 | 56 |
| 1168 | RF00495 IRES Hsp70 | 0.0 (0/54) | 4.1 | 0.0 (0/1) | 6.5 | 81.1 | 14 |
| 1169 | RF00332 snoZ266 | 0.0 (0/10) | 4.0 | 0.0 (0/0) | 6.9 | 74.8 | 11 |
| 1170 | RF00358 snoZ101 | 0.0 (0/10) | 4.0 | 0.0 (0/0) | 5.9 | 76.5 | 13 |
| 1171 | RF00296 snoR16 | 0.0 (0/5) | 4.0 | 0.0 (0/0) | 4.8 | 74.7 | 18 |
| 1172 | RF00554 SNORA48 | 0.0 (0/38) | 3.9 | 0.0 (0/0) | 6.8 | 78.5 | 24 |
| 1173 | RF00646 mir-204 | 0.0 (0/28) | 3.9 | 0.0 (0/0) | 4.7 | 72.5 | 32 |
| 1174 | RF00763 mir-339 | 0.0 (0/33) | 3.9 | 0.0 (0/0) | 4.9 | 75.1 | 14 |
| 1175 | RF00917 mir-708 | 0.0 (0/31) | 3.9 | 0.0 (0/0) | 4.7 | 79.9 | 21 |
| 1176 | RF00405 SNORA44 | 0.0 (0/21) | 3.8 | 0.0 (0/0) | 6.4 | 81.5 | 27 |
| 1177 | RF01811 PtaRNA1 | 0.0 (0/21) | 3.8 | 0.0 (0/0) | 4.5 | 78.3 | 16 |
| 1178 | RF02514 5 ureB sRNA | 0.0 (0/90) | 3.7 | 0.0 (0/0) | 5.9 | 76.3 | 13 |
| 1179 | RF00404 SNORA46 | 0.0 (0/41) | 3.6 | 0.0 (0/0) | 5.3 | 82.6 | 22 |
| 1180 | RF00215 Tombus 3 III | 0.0 (0/14) | 3.6 | 0.0 (0/1) | 6.6 | 79.5 | 28 |
| 1181 | RF00493 snoU2-30 | 0.0 (0/11) | 3.6 | 0.0 (0/0) | 4.3 | 87.9 | 21 |
| 1182 | RF00661 mir-31 | 0.0 (0/14) | 3.6 | 0.0 (0/0) | 4.6 | 63.6 | 28 |
| 1183 | RF00623 P1 | 0.0 (0/28) | 3.6 | 0.0 (0/1) | 4.8 | 67.7 | 14 |
| 1184 | RF00252 Alfamo CPB | 0.0 (0/56) | 3.6 | 0.0 (0/0) | 4.0 | 89.6 | 18 |
| 1185 | RF02060 STnc410 | 0.0 (0/17) | 3.5 | 0.0 (0/0) | 6.4 | 70.7 | 14 |
| 1186 | RF00183 G-CSF SLDE | 0.0 (0/26) | 3.5 | 0.0 (0/0) | 4.9 | 80.6 | 16 |
| 1187 | RF00728 mir-81 | 0.0 (0/34) | 3.5 | 0.0 (0/0) | 5.0 | 64.5 | 9 |
| 1188 | RF02548 Oskar OES | 0.0 (0/21) | 3.3 | 0.0 (0/0) | 5.0 | 80.3 | 14 |
| 1189 | RF02061 mir-301 | 0.0 (0/24) | 3.3 | 0.0 (0/0) | 4.2 | 77.5 | 17 |
| 1190 | RF00581 SNORD12 | 0.0 (0/9) | 3.3 | 0.0 (0/0) | 7.2 | 71.1 | 8 |
| 1191 | RF01203 snR47 | 0.0 (0/3) | 3.3 | 0.0 (0/0) | 4.0 | 73.9 | 17 |
| 1192 | RF01237 snR161 | 0.0 (0/36) | 3.3 | 0.0 (0/0) | 5.2 | 65.2 | 10 |
| 1193 | RF00427 SCARNA23 | 0.0 (0/36) | 3.3 | 0.0 (0/0) | 5.3 | 79.0 | 18 |
| 1194 | RF00278 SNORD50 | 0.0 (0/3) | 3.3 | 0.0 (0/0) | 6.3 | 81.7 | 26 |
| 1195 | RF00745 mir-499 | 0.0 (0/25) | 3.2 | 0.0 (0/0) | 3.5 | 77.9 | 19 |
| 1196 | RF00662 mir-132 | 0.0 (0/16) | 3.1 | 0.0 (0/1) | 3.1 | 66.4 | 22 |
| 1197 | RF01392 isrI | 0.0 (0/13) | 3.1 | 0.0 (0/0) | 4.0 | 82.4 | 30 |
| 1198 | RF01019 mir-922 | 0.0 (0/27) | 3.0 | 0.0 (0/0) | 4.8 | 80.2 | 12 |
| 1199 | RF00456 mir-34 | 0.0 (0/30) | 3.0 | 0.0 (0/0) | 4.8 | 74.0 | 18 |
| 1200 | RF00408 SNORA1 | 0.0 (0/24) | 2.9 | 0.0 (0/0) | 5.2 | 76.6 | 29 |
| 1201 | RF00683 mir-143 | 0.0 (0/28) | 2.9 | 0.0 (0/1) | 3.0 | 78.4 | 18 |
| 1202 | RF02434 SpF39 sRNA | 0.0 (0/28) | 2.9 | 0.0 (0/0) | 4.9 | 83.2 | 13 |
| 1203 | RF01879 TUSC7 | 0.0 (0/24) | 2.9 | 0.0 (0/0) | 5.2 | 87.5 | 25 |
| 1204 | RF00265 SNORA69 | 0.0 (0/38) | 2.9 | 0.0 (0/0) | 5.1 | 83.9 | 16 |
| 1205 | RF00316 snoR43 | 0.0 (0/7) | 2.9 | 0.0 (0/0) | 5.4 | 86.9 | 16 |
| 1206 | RF02261 MAT2A B | 0.0 (0/21) | 2.9 | 0.0 (0/0) | 4.1 | 94.0 | 28 |
| 1207 | RF00598 SNORA76 | 0.0 (0/39) | 2.8 | 0.0 (0/0) | 4.5 | 83.7 | 22 |
| 1208 | RF02095 mir-2985-2 | 0.0 (0/32) | 2.8 | 0.0 (0/0) | 4.0 | 83.8 | 20 |
| 1209 | RF01010 mir-632 | 0.0 (0/32) | 2.8 | 0.0 (0/0) | 5.1 | 82.9 | 16 |
| 1210 | RF00841 mir-384 | 0.0 (0/26) | 2.7 | 0.0 (0/0) | 4.8 | 72.2 | 16 |
| 1211 | RF00423 SCARNA4 | 0.0 (0/33) | 2.7 | 0.0 (0/0) | 4.8 | 82.9 | 24 |
| 1212 | RF01249 snR190 | 0.0 (0/62) | 2.7 | 0.0 (0/0) | 5.1 | 80.6 | 10 |
| 1213 | RF00578 SNORD89 | 0.0 (0/33) | 2.7 | 0.0 (0/0) | 4.3 | 88.5 | 18 |
| 1214 | RF02260 MAT2A A | 0.0 (0/19) | 2.6 | 0.0 (0/0) | 4.7 | 89.7 | 20 |
| 1215 | RF00410 SNORA2 | 0.0 (0/23) | 2.6 | 0.0 (0/0) | 4.4 | 75.8 | 18 |
| 1216 | RF00876 mir-684 | 0.0 (0/27) | 2.6 | 0.0 (0/0) | 4.4 | 88.3 | 28 |
| 1217 | RF02531 NRF2 IRES | 0.0 (0/27) | 2.6 | 0.0 (0/0) | 5.1 | 82.3 | 20 |
| 1218 | RF00682 mir-144 | 0.0 (0/27) | 2.6 | 0.0 (0/0) | 3.4 | 84.0 | 22 |
| 1219 | RF01033 mir-767 | 0.0 (0/39) | 2.6 | 0.0 (0/0) | 4.0 | 82.7 | 14 |
| 1220 | RF02701 PsrR1 | 0.0 (0/8) | 2.5 | 0.0 (0/1) | 5.1 | 65.9 | 13 |
| 1221 | RF01182 SNORD11 | 0.0 (0/4) | 2.5 | 0.0 (0/0) | 6.8 | 85.2 | 19 |
| 1222 | RF00279 SNORD45 | 0.0 (0/4) | 2.5 | 0.0 (0/0) | 4.5 | 78.8 | 11 |
| 1223 | RF00339 snoR60 | 0.0 (0/4) | 2.5 | 0.0 (0/0) | 2.5 | 84.5 | 10 |
| 1224 | RF02234 sX15 | 0.0 (0/48) | 2.5 | 0.0 (0/0) | 3.2 | 82.6 | 14 |
| 1225 | RF01832 ROSE 2 | 0.0 (0/16) | 2.5 | 0.0 (0/0) | 3.1 | 83.7 | 14 |
| 1226 | RF00438 SNORA33 | 0.0 (0/24) | 2.5 | 0.0 (0/0) | 4.7 | 77.8 | 28 |
| 1227 | RF00607 SNORD98 | 0.0 (0/4) | 2.5 | 0.0 (0/0) | 4.8 | 81.6 | 10 |
| 1228 | RF00733 mir-296 | 0.0 (0/33) | 2.4 | 0.0 (0/0) | 3.7 | 83.2 | 12 |
| 1229 | RF00840 mir-374 | 0.0 (0/29) | 2.4 | 0.0 (0/0) | 3.8 | 81.7 | 13 |
| 1230 | RF01770 rimP | 0.0 (0/13) | 2.3 | 0.0 (0/0) | 2.8 | 71.2 | 46 |
| 1231 | RF02100 tfoR | 0.0 (0/30) | 2.3 | 0.0 (0/0) | 3.9 | 81.8 | 10 |
| 1232 | RF00108 SNORD116 | 0.0 (0/13) | 2.3 | 0.0 (0/0) | 3.7 | 82.0 | 48 |
| 1233 | RF00214 Retro dr1 | 0.0 (0/22) | 2.3 | 0.0 (0/0) | 3.4 | 89.2 | 26 |
| 1234 | RF00983 mir-662 | 0.0 (0/35) | 2.3 | 0.0 (0/0) | 3.9 | 84.8 | 14 |
| 1235 | RF00875 mir-692 | 0.0 (0/22) | 2.3 | 0.0 (0/0) | 4.5 | 78.0 | 11 |
| 1236 | RF03024 Rothia-sucC | 0.0 (0/9) | 2.2 | 0.0 (0/0) | 4.6 | 82.5 | 28 |
| 1237 | RF00155 SNORA66 | 0.0 (0/23) | 2.2 | 0.0 (0/0) | 3.7 | 81.8 | 25 |
| 1238 | RF00362 Pospi RY | 0.0 (0/23) | 2.2 | 0.0 (0/0) | 2.7 | 92.2 | 16 |
| 1239 | RF00333 snoZ157 | 0.0 (0/18) | 2.2 | 0.0 (0/0) | 5.3 | 76.3 | 10 |
| 1240 | RF01200 SNORD125 | 0.0 (0/9) | 2.2 | 0.0 (0/1) | 4.0 | 81.4 | 16 |
| 1241 | RF00465 JEV hairpin | 0.0 (0/18) | 2.2 | 0.0 (0/0) | 3.3 | 86.5 | 20 |
| 1242 | RF01911 MIR2118 | 0.0 (0/43) | 2.1 | 0.0 (0/0) | 4.3 | 65.3 | 7 |
| 1243 | RF01173 snoU105B | 0.0 (0/14) | 2.1 | 0.0 (0/0) | 3.9 | 78.2 | 13 |
| 1244 | RF00542 snopsi28S-1192 | 0.0 (0/30) | 2.0 | 0.0 (0/0) | 3.0 | 81.5 | 15 |
| 1245 | RF01414 class I RNA | 0.0 (0/10) | 2.0 | 0.0 (0/0) | 3.6 | 77.4 | 21 |
| 1246 | RF01320 CRISPR-DR7 | 0.0 (0/5) | 2.0 | 0.0 (0/0) | 2.2 | 76.9 | 10 |
| 1247 | RF02227 sX8 | 0.0 (0/25) | 2.0 | 0.0 (0/0) | 4.2 | 79.2 | 12 |
| 1248 | RF00673 mir-217 | 0.0 (0/30) | 2.0 | 0.0 (0/0) | 3.6 | 78.2 | 27 |
| 1249 | RF00579 SNORD90 | 0.0 (0/30) | 2.0 | 0.0 (0/0) | 3.4 | 86.7 | 18 |

Continued on next page

| | RNA family (seed alignment) | **Sensitivity** annotated bpairs that covary % (cov_bps/bps) | **Power** average power % | **Positive Predictive Value** covarying pairs in structure % (cov_bps/cov_pairs) | average substitutions per bpair | avg pairwise identity % | number of sequences |
|---|---|---|---|---|---|---|---|
| 1250 | RF01231 snoR74 | 0.0 (0/25) | 2.0 | 0.0 (0/0) | 4.0 | 68.0 | 9 |
| 1251 | RF03002 lysM-Prevotella | 0.0 (0/10) | 2.0 | 0.0 (0/0) | 4.3 | 79.6 | 47 |
| 1252 | RF00951 mir-1302 | 0.0 (0/15) | 2.0 | 0.0 (0/0) | 4.6 | 82.8 | 24 |
| 1253 | RF00428 SNORA38 | 0.0 (0/25) | 2.0 | 0.0 (0/0) | 3.6 | 83.9 | 25 |
| 1254 | RF00570 SNORD64 | 0.0 (0/15) | 2.0 | 0.0 (0/0) | 4.9 | 80.5 | 17 |
| 1255 | RF00074 mir-29 | 0.0 (0/21) | 1.9 | 0.0 (0/0) | 2.4 | 74.5 | 10 |
| 1256 | RF01540 TB11Cs4H2 | 0.0 (0/16) | 1.9 | 0.0 (0/0) | 3.2 | 75.0 | 6 |
| 1257 | RF02064 STnc370 | 0.0 (0/16) | 1.9 | 0.0 (0/0) | 3.0 | 78.9 | 10 |
| 1258 | RF00228 IRES HepA | 0.0 (0/102) | 1.9 | 0.0 (0/0) | 2.4 | 96.4 | 23 |
| 1259 | RF00482 snoF1 F2 | 0.0 (0/32) | 1.9 | 0.0 (0/0) | 2.8 | 86.5 | 8 |
| 1260 | RF00664 mir-223 | 0.0 (0/31) | 1.9 | 0.0 (0/0) | 2.6 | 77.5 | 19 |
| 1261 | RF02057 STnc40 | 0.0 (0/16) | 1.9 | 0.0 (0/0) | 3.2 | 68.5 | 17 |
| 1262 | RF02222 sX2 | 0.0 (0/31) | 1.9 | 0.0 (0/0) | 2.9 | 88.8 | 9 |
| 1263 | RF01668 P10 | 0.0 (0/16) | 1.9 | 0.0 (0/0) | 3.1 | 71.6 | 8 |
| 1264 | RF01840 ovine lenti FSE | 0.0 (0/17) | 1.8 | 0.0 (0/0) | 2.6 | 88.1 | 14 |
| 1265 | RF00126 ryfA | 0.0 (0/83) | 1.8 | 0.0 (0/0) | 3.5 | 74.1 | 9 |
| 1266 | RF03029 RT-8 | 0.0 (0/22) | 1.8 | 0.0 (0/0) | 4.6 | 81.0 | 9 |
| 1267 | RF02228 sX9 | 0.0 (0/17) | 1.8 | 0.0 (0/0) | 4.1 | 76.8 | 16 |
| 1268 | RF02422 Bp1 738 | 0.0 (0/33) | 1.8 | 0.0 (0/0) | 3.3 | 75.0 | 21 |
| 1269 | RF00753 mir-503 | 0.0 (0/28) | 1.8 | 0.0 (0/0) | 2.6 | 87.8 | 14 |
| 1270 | RF01757 sbcD | 0.0 (0/51) | 1.8 | 0.0 (0/0) | 3.2 | 77.5 | 6 |
| 1271 | RF00576 SNORD71 | 0.0 (0/22) | 1.8 | 0.0 (0/0) | 2.5 | 83.3 | 18 |
| 1272 | RF00783 mir-484 | 0.0 (0/23) | 1.7 | 0.0 (0/0) | 4.8 | 83.1 | 15 |
| 1273 | RF00180 REN-SRE | 0.0 (0/6) | 1.7 | 0.0 (0/0) | 4.2 | 89.4 | 13 |
| 1274 | RF00109 Vimentin3 | 0.0 (0/18) | 1.7 | 0.0 (0/0) | 4.1 | 75.5 | 19 |
| 1275 | RF02265 MAT2A F | 0.0 (0/23) | 1.7 | 0.0 (0/0) | 2.7 | 94.3 | 26 |
| 1276 | RF00731 mir-155 | 0.0 (0/23) | 1.7 | 0.0 (0/0) | 3.3 | 77.7 | 17 |
| 1277 | RF02495 ohsC RNA | 0.0 (0/24) | 1.7 | 0.0 (0/0) | 3.8 | 90.2 | 34 |
| 1278 | RF02264 MAT2A E | 0.0 (0/24) | 1.7 | 0.0 (0/0) | 3.0 | 93.0 | 28 |
| 1279 | RF00674 mir-187 | 0.0 (0/23) | 1.7 | 0.0 (0/0) | 2.2 | 76.5 | 20 |
| 1280 | RF00443 SNORA27 | 0.0 (0/24) | 1.7 | 0.0 (0/0) | 4.2 | 84.3 | 24 |
| 1281 | RF00727 bantam | 0.0 (0/25) | 1.6 | 0.0 (0/0) | 3.1 | 71.2 | 11 |
| 1282 | RF00928 mir-590 | 0.0 (0/32) | 1.6 | 0.0 (0/0) | 3.8 | 76.6 | 10 |
| 1283 | RF01018 mir-569 | 0.0 (0/31) | 1.6 | 0.0 (0/0) | 3.4 | 85.8 | 13 |
| 1284 | RF00747 mir-283 | 0.0 (0/25) | 1.6 | 0.0 (0/0) | 3.4 | 69.7 | 11 |
| 1285 | RF00669 mir-96 | 0.0 (0/26) | 1.5 | 0.0 (0/0) | 2.5 | 89.2 | 25 |
| 1286 | RF00457 IRES mnt | 0.0 (0/33) | 1.5 | 0.0 (0/0) | 3.2 | 91.6 | 21 |
| 1287 | RF00222 IRES Bag1 | 0.0 (0/52) | 1.5 | 0.0 (0/0) | 3.5 | 78.8 | 15 |
| 1288 | RF01238 snR70 | 0.0 (0/52) | 1.5 | 0.0 (0/0) | 3.7 | 81.6 | 9 |
| 1289 | RF00459 MPMV package | 0.0 (0/57) | 1.4 | 0.0 (0/0) | 3.6 | 71.9 | 9 |
| 1290 | RF01787 drz-agam-1 | 0.0 (0/22) | 1.4 | 0.0 (0/0) | 4.3 | 72.6 | 7 |
| 1291 | RF00660 mir-214 | 0.0 (0/36) | 1.4 | 0.0 (0/0) | 2.3 | 77.9 | 8 |
| 1292 | RF00145 snoZ105 | 0.0 (0/7) | 1.4 | 0.0 (0/0) | 4.0 | 65.3 | 11 |
| 1293 | RF02808 PyrR210 | 0.0 (0/14) | 1.4 | 0.0 (0/0) | 2.7 | 81.0 | 6 |
| 1294 | RF00621 CoTC ribozyme | 0.0 (0/49) | 1.4 | 0.0 (0/0) | 3.2 | 89.7 | 10 |
| 1295 | RF00390 UPSK | 0.0 (0/7) | 1.4 | 0.0 (0/0) | 1.6 | 93.6 | 6 |
| 1296 | RF02465 Ms AS-5 | 0.0 (0/7) | 1.4 | 0.0 (0/0) | 2.6 | 84.6 | 9 |
| 1297 | RF00712 mir-460 | 0.0 (0/21) | 1.4 | 0.0 (0/0) | 1.7 | 77.5 | 8 |
| 1298 | RF03042 porB | 0.0 (0/7) | 1.4 | 0.0 (0/0) | 4.6 | 82.7 | 85 |
| 1299 | RF00952 mir-650 | 0.0 (0/29) | 1.4 | 0.0 (0/0) | 3.7 | 83.8 | 11 |
| 1300 | RF01036 mir-567 | 0.0 (0/28) | 1.4 | 0.0 (0/0) | 3.9 | 72.3 | 9 |
| 1301 | RF00319 SNORA23 | 0.0 (0/45) | 1.3 | 0.0 (0/0) | 3.0 | 83.0 | 9 |
| 1302 | RF00081 ArcZ | 0.0 (0/15) | 1.3 | 0.0 (0/0) | 2.5 | 77.2 | 9 |
| 1303 | RF02052 STnc630 | 0.0 (0/31) | 1.3 | 0.0 (0/1) | 2.5 | 76.8 | 10 |
| 1304 | RF01927 MIR1222 | 0.0 (0/39) | 1.3 | 0.0 (0/0) | 3.5 | 57.8 | 5 |
| 1305 | RF00705 mir-202 | 0.0 (0/30) | 1.3 | 0.0 (0/0) | 2.1 | 81.2 | 13 |
| 1306 | RF00622 CPEB3 ribozyme | 0.0 (0/23) | 1.3 | 0.0 (0/0) | 2.7 | 84.5 | 12 |
| 1307 | RF01093 RF site5 | 0.0 (0/16) | 1.2 | 0.0 (0/0) | 2.5 | 76.4 | 12 |
| 1308 | RF00989 mir-492 | 0.0 (0/32) | 1.2 | 0.0 (0/0) | 4.0 | 82.0 | 14 |
| 1309 | RF00198 SL1 | 0.0 (0/24) | 1.2 | 0.0 (0/0) | 1.7 | 92.1 | 28 |
| 1310 | RF01990 SECIS 4 | 0.0 (0/8) | 1.2 | 0.0 (0/0) | 1.9 | 88.4 | 24 |
| 1311 | RF02230 sX11 | 0.0 (0/43) | 1.2 | 0.0 (0/0) | 2.5 | 84.9 | 10 |
| 1312 | RF00696 mir-203 | 0.0 (0/25) | 1.2 | 0.0 (0/0) | 2.0 | 79.6 | 10 |
| 1313 | RF00247 mir-160 | 0.0 (0/25) | 1.2 | 0.0 (0/0) | 2.6 | 65.8 | 7 |
| 1314 | RF00704 MIR397 | 0.0 (0/25) | 1.2 | 0.0 (0/0) | 2.5 | 60.1 | 7 |
| 1315 | RF01546 TB6Cs1H1 | 0.0 (0/16) | 1.2 | 0.0 (0/1) | 1.6 | 79.5 | 6 |
| 1316 | RF01558 TB9Cs4H2 | 0.0 (0/16) | 1.2 | 0.0 (0/0) | 2.4 | 83.7 | 5 |
| 1317 | RF01903 mir-500 | 0.0 (0/25) | 1.2 | 0.0 (0/0) | 2.9 | 78.5 | 26 |
| 1318 | RF00487 IRES Cx43 | 0.0 (0/56) | 1.1 | 0.0 (0/0) | 3.2 | 87.2 | 14 |
| 1319 | RF00226 IRES n-myc | 0.0 (0/36) | 1.1 | 0.0 (0/0) | 2.5 | 72.2 | 6 |
| 1320 | RF02552 RcsR1 | 0.0 (0/28) | 1.1 | 0.0 (0/1) | 2.6 | 81.9 | 10 |
| 1321 | RF01387 isrC | 0.0 (0/37) | 1.1 | 0.0 (0/0) | 3.1 | 90.9 | 16 |
| 1322 | RF00381 Antizyme FSE | 0.0 (0/18) | 1.1 | 0.0 (0/0) | 1.6 | 84.2 | 13 |
| 1323 | RF00751 mir-12 | 0.0 (0/27) | 1.1 | 0.0 (0/0) | 2.1 | 77.5 | 7 |
| 1324 | RF00658 mir-21 | 0.0 (0/27) | 1.1 | 0.0 (0/0) | 2.6 | 80.3 | 11 |
| 1325 | RF00375 HIV PBS | 0.0 (0/18) | 1.1 | 0.0 (0/0) | 2.7 | 90.9 | 130 |
| 1326 | RF02518 mir-2494 | 0.0 (0/38) | 1.1 | 0.0 (0/0) | 3.0 | 81.8 | 10 |
| 1327 | RF00541 snopsi28S-2876 | 0.0 (0/28) | 1.1 | 0.0 (0/0) | 3.1 | 80.5 | 8 |
| 1328 | RF01775 RsaOG | 0.0 (0/36) | 1.1 | 0.0 (0/1) | 2.5 | 84.7 | 7 |
| 1329 | RF01549 TB8Cs2H1 | 0.0 (0/18) | 1.1 | 0.0 (0/0) | 3.5 | 69.9 | 6 |
| 1330 | RF00195 RsmY | 0.0 (0/19) | 1.1 | 0.0 (0/1) | 2.1 | 77.0 | 11 |
| 1331 | RF00240 RNA-OUT | 0.0 (0/20) | 1.0 | 0.0 (0/0) | 3.0 | 86.8 | 16 |
| 1332 | RF01031 mir-639 | 0.0 (0/31) | 1.0 | 0.0 (0/0) | 2.3 | 76.2 | 6 |
| 1333 | RF01061 mir-548 | 0.0 (0/40) | 1.0 | 0.0 (0/0) | 2.6 | 84.8 | 11 |

Continued on next page

| | RNA family (seed alignment) | Sensitivity annotated bpairs that covary % (cov_bps/bps) | Power average power % | Positive Predictive Value covarying pairs in structure % (cov_bps/cov_pairs) | average substitutions per bpair | avg pairwise identity % | number of sequences |
|---|---|---|---|---|---|---|---|
| 1334 | RF02420 Bp1 162 | 0.0 (0/20) | 1.0 | 0.0 (0/0) | 2.5 | 82.9 | 11 |
| 1335 | RF01228 snoR111 | 0.0 (0/31) | 1.0 | 0.0 (0/1) | 2.5 | 66.7 | 6 |
| 1336 | RF00242 ctRNA pT181 | 0.0 (0/20) | 1.0 | 0.0 (0/0) | 2.3 | 80.9 | 16 |
| 1337 | RF02238 asX4 | 0.0 (0/87) | 1.0 | 0.0 (0/0) | 2.3 | 86.0 | 8 |
| 1338 | RF02911 Baculoviridae NAE | 0.0 (0/29) | 1.0 | 0.0 (0/0) | 2.9 | 72.3 | 6 |
| 1339 | RF02438 SpF51 sRNA | 0.0 (0/10) | 1.0 | 0.0 (0/0) | 1.5 | 90.1 | 8 |
| 1340 | RF00452 mir-172 | 0.0 (0/23) | 0.9 | 0.0 (0/0) | 2.0 | 60.6 | 11 |
| 1341 | RF02503 Atu C9 | 0.0 (0/21) | 0.9 | 0.0 (0/0) | 2.3 | 74.1 | 9 |
| 1342 | RF01836 weev FSE | 0.0 (0/11) | 0.9 | 0.0 (0/0) | 1.5 | 87.3 | 6 |
| 1343 | RF00686 mir-338 | 0.0 (0/21) | 0.9 | 0.0 (0/0) | 1.9 | 73.0 | 21 |
| 1344 | RF02786 snoTBR2 | 0.0 (0/33) | 0.9 | 0.0 (0/1) | 3.0 | 72.5 | 5 |
| 1345 | RF01402 STnc150 | 0.0 (0/22) | 0.9 | 0.0 (0/22) | 3.0 | 91.2 | 9 |
| 1346 | RF00417 SNORA56 | 0.0 (0/33) | 0.9 | 0.0 (0/0) | 3.5 | 82.6 | 17 |
| 1347 | RF02424 Bp2 287 | 0.0 (0/44) | 0.9 | 0.0 (0/0) | 1.6 | 83.5 | 14 |
| 1348 | RF00492 SCARNA17 | 0.0 (0/32) | 0.9 | 0.0 (0/0) | 3.2 | 76.4 | 6 |
| 1349 | RF02053 STnc430 | 0.0 (0/33) | 0.9 | 0.0 (0/0) | 3.1 | 68.6 | 7 |
| 1350 | RF01784 bablM | 0.0 (0/11) | 0.9 | 0.0 (0/0) | 1.5 | 89.9 | 12 |
| 1351 | RF01897 mir-188 | 0.0 (0/21) | 0.9 | 0.0 (0/0) | 3.0 | 71.2 | 14 |
| 1352 | RF00036 RRE | 0.0 (0/116) | 0.9 | 0.0 (0/0) | 1.6 | 97.0 | 65 |
| 1353 | RF00994 mir-1255 | 0.0 (0/21) | 0.9 | 0.0 (0/0) | 2.5 | 83.1 | 10 |
| 1354 | RF00121 MicC | 0.0 (0/23) | 0.9 | 0.0 (0/0) | 2.0 | 71.5 | 8 |
| 1355 | RF00663 mir-183 | 0.0 (0/23) | 0.9 | 0.0 (0/0) | 2.3 | 78.1 | 16 |
| 1356 | RF02891 AgrB | 0.0 (0/21) | 0.9 | 0.0 (0/0) | 1.7 | 77.4 | 4 |
| 1357 | RF02546 LSU trypano mito | 0.0 (0/58) | 0.9 | 0.0 (0/0) | 4.1 | 76.9 | 6 |
| 1358 | RF01895 mir-193 | 0.0 (0/23) | 0.9 | 0.0 (0/0) | 2.1 | 72.1 | 17 |
| 1359 | RF01824 RUF20 | 0.0 (0/76) | 0.9 | 0.0 (0/0) | 3.8 | 73.8 | 6 |
| 1360 | RF02512 PYLIS 5 | 0.0 (0/22) | 0.9 | 0.0 (0/0) | 1.2 | 94.0 | 17 |
| 1361 | RF00176 Tombus 3 IV | 0.0 (0/25) | 0.8 | 0.0 (0/0) | 1.2 | 93.3 | 18 |
| 1362 | RF01941 MIR1223 | 0.0 (0/25) | 0.8 | 0.0 (0/0) | 2.4 | 59.5 | 8 |
| 1363 | RF01291 SNORD97 | 0.0 (0/13) | 0.8 | 0.0 (0/1) | 2.1 | 86.5 | 20 |
| 1364 | RF01220 snoR104 | 0.0 (0/24) | 0.8 | 0.0 (0/0) | 3.4 | 66.4 | 8 |
| 1365 | RF02371 PyrG leader | 0.0 (0/12) | 0.8 | 0.0 (0/0) | 1.9 | 73.9 | 10 |
| 1366 | RF02232 sX13 | 0.0 (0/37) | 0.8 | 0.0 (0/0) | 1.9 | 90.4 | 12 |
| 1367 | RF01401 rseX | 0.0 (0/13) | 0.8 | 0.0 (0/0) | 2.3 | 76.1 | 12 |
| 1368 | RF00865 MIR169 5 | 0.0 (0/26) | 0.8 | 0.0 (0/0) | 2.6 | 65.1 | 6 |
| 1369 | RF00636 NRON | 0.0 (0/64) | 0.8 | 0.0 (0/0) | 2.0 | 91.9 | 24 |
| 1370 | RF01052 Arthropod 7SK | 0.0 (0/39) | 0.8 | 0.0 (0/0) | 2.5 | 67.0 | 19 |
| 1371 | RF00735 mir-367 | 0.0 (0/25) | 0.8 | 0.0 (0/0) | 1.5 | 85.9 | 13 |
| 1372 | RF00489 ctRNA p42d | 0.0 (0/12) | 0.8 | 0.0 (0/0) | 2.2 | 86.2 | 10 |
| 1373 | RF00394 SNORA4 | 0.0 (0/40) | 0.8 | 0.0 (0/0) | 2.5 | 76.4 | 7 |
| 1374 | RF00142 snoZ118 | 0.0 (0/13) | 0.8 | 0.0 (0/0) | 2.8 | 84.2 | 7 |
| 1375 | RF00467 RSV PBS | 0.0 (0/15) | 0.7 | 0.0 (0/0) | 0.9 | 92.7 | 22 |
| 1376 | RF00999 mir-924 | 0.0 (0/14) | 0.7 | 0.0 (0/0) | 2.1 | 81.5 | 4 |
| 1377 | RF00772 mir-328 | 0.0 (0/28) | 0.7 | 0.0 (0/0) | 2.3 | 79.3 | 13 |
| 1378 | RF00659 mir-365 | 0.0 (0/29) | 0.7 | 0.0 (0/1) | 2.2 | 80.4 | 9 |
| 1379 | RF01996 mir-995 | 0.0 (0/30) | 0.7 | 0.0 (0/0) | 1.8 | 77.8 | 9 |
| 1380 | RF00361 snoZ119 | 0.0 (0/15) | 0.7 | 0.0 (0/0) | 1.1 | 94.0 | 6 |
| 1381 | RF00792 mir-490 | 0.0 (0/28) | 0.7 | 0.0 (0/0) | 2.9 | 78.5 | 19 |
| 1382 | RF00678 mir-140 | 0.0 (0/30) | 0.7 | 0.0 (0/0) | 1.2 | 87.0 | 14 |
| 1383 | RF01449 S pombe snR100 | 0.0 (0/57) | 0.7 | 0.0 (0/0) | 3.0 | 68.0 | 3 |
| 1384 | RF02802 PyrR204 | 0.0 (0/15) | 0.7 | 0.0 (0/0) | 1.5 | 76.7 | 5 |
| 1385 | RF01393 isrJ | 0.0 (0/15) | 0.7 | 0.0 (0/0) | 3.0 | 76.3 | 5 |
| 1386 | RF00756 mir-299 | 0.0 (0/28) | 0.7 | 0.0 (0/0) | 1.0 | 87.1 | 6 |
| 1387 | RF01208 snoR99 | 0.0 (0/14) | 0.7 | 0.0 (0/0) | 2.2 | 70.0 | 9 |
| 1388 | RF02466 Ms AS-8 | 0.0 (0/15) | 0.7 | 0.0 (0/0) | 2.1 | 78.9 | 10 |
| 1389 | RF01790 htlv FSE | 0.0 (0/14) | 0.7 | 0.0 (0/0) | 1.3 | 84.5 | 8 |
| 1390 | RF01219 snoR100 | 0.0 (0/15) | 0.7 | 0.0 (0/0) | 2.3 | 71.8 | 9 |
| 1391 | RF01703 Dictyoglomi-1 | 0.0 (0/81) | 0.7 | 0.0 (0/0) | 2.0 | 78.5 | 4 |
| 1392 | RF00072 SNORA75 | 0.0 (0/28) | 0.7 | 0.0 (0/0) | 1.5 | 73.3 | 6 |
| 1393 | RF00832 mir-71 | 0.0 (0/17) | 0.6 | 0.0 (0/0) | 2.2 | 74.2 | 12 |
| 1394 | RF00801 mir-280 | 0.0 (0/35) | 0.6 | 0.0 (0/0) | 1.6 | 86.5 | 8 |
| 1395 | RF01455 DPB | 0.0 (0/18) | 0.6 | 0.0 (0/0) | 2.0 | 85.0 | 19 |
| 1396 | RF00505 RydC | 0.0 (0/16) | 0.6 | 0.0 (0/0) | 1.7 | 86.4 | 5 |
| 1397 | RF00461 IRES VEGF A | 0.0 (0/95) | 0.6 | 0.0 (0/0) | 1.7 | 90.7 | 7 |
| 1398 | RF00737 mir-322 | 0.0 (0/31) | 0.6 | 0.0 (0/0) | 1.8 | 88.9 | 11 |
| 1399 | RF01008 mir-636 | 0.0 (0/31) | 0.6 | 0.0 (0/0) | 1.9 | 78.3 | 5 |
| 1400 | RF00959 mir-612 | 0.0 (0/35) | 0.6 | 0.0 (0/0) | 2.1 | 76.1 | 6 |
| 1401 | RF01773 rpsL pseudo | 0.0 (0/35) | 0.6 | 0.0 (0/1) | 2.5 | 80.6 | 9 |
| 1402 | RF02374 YenS | 0.0 (0/54) | 0.6 | 0.0 (0/0) | 1.9 | 82.1 | 7 |
| 1403 | RF00700 mir-375 | 0.0 (0/17) | 0.6 | 0.0 (0/0) | 1.1 | 78.9 | 9 |
| 1404 | RF02425 SpF01 sRNA | 0.0 (0/16) | 0.6 | 0.0 (0/0) | 2.8 | 83.1 | 13 |
| 1405 | RF02421 Bp1 684 | 0.0 (0/16) | 0.6 | 0.0 (0/1) | 1.1 | 86.3 | 14 |
| 1406 | RF01012 mir-628 | 0.0 (0/32) | 0.6 | 0.0 (0/1) | 2.9 | 78.2 | 9 |
| 1407 | RF02262 MAT2A C | 0.0 (0/18) | 0.6 | 0.0 (0/0) | 2.1 | 90.9 | 18 |
| 1408 | RF02551 DapZ | 0.0 (0/18) | 0.6 | 0.0 (0/0) | 1.1 | 79.9 | 6 |
| 1409 | RF00501 Rota CRE | 0.0 (0/17) | 0.6 | 0.0 (0/1) | 1.6 | 86.6 | 14 |
| 1410 | RF02819 V AS7 | 0.0 (0/53) | 0.6 | 0.0 (0/0) | 2.6 | 78.1 | 7 |
| 1411 | RF00223 IRES Bip | 0.0 (0/33) | 0.6 | 0.0 (0/2) | 2.4 | 81.1 | 9 |
| 1412 | RF01524 TB10Cs1H3 | 0.0 (0/17) | 0.6 | 0.0 (0/0) | 1.4 | 80.8 | 5 |
| 1413 | RF01023 mir-940 | 0.0 (0/34) | 0.6 | 0.0 (0/0) | 1.7 | 89.3 | 8 |
| 1414 | RF00872 mir-652 | 0.0 (0/31) | 0.6 | 0.0 (0/0) | 2.5 | 76.9 | 13 |
| 1415 | RF00065 snoR9 | 0.0 (0/18) | 0.6 | 0.0 (0/1) | 1.3 | 83.7 | 5 |
| 1416 | RF00251 mir-219 | 0.0 (0/17) | 0.6 | 0.0 (0/0) | 1.9 | 83.4 | 13 |
| 1417 | RF01215 snoR97 | 0.0 (0/17) | 0.6 | 0.0 (0/0) | 1.4 | 72.2 | 7 |

Continued on next page

| | RNA family (seed alignment) | Sensitivity annotated bpairs that covary % (cov_bps/bps) | Power average power % | Positive Predictive Value covarying pairs in structure % (cov_bps/cov_pairs) | average substitutions per bpair | avg pairwise identity % | number of sequences |
|---|---|---|---|---|---|---|---|
| 1418 | RF00991 mir-599 | 0.0 (0/35) | 0.6 | 0.0 (0/0) | 2.3 | 86.2 | 11 |
| 1419 | RF00912 mir-877 | 0.0 (0/18) | 0.6 | 0.0 (0/0) | 2.6 | 81.1 | 13 |
| 1420 | RF01385 isrA | 0.0 (0/20) | 0.5 | 0.0 (0/0) | 2.2 | 87.6 | 10 |
| 1421 | RF00194 Rubella 3 | 0.0 (0/20) | 0.5 | 0.0 (0/0) | 2.4 | 95.3 | 25 |
| 1422 | RF00844 mir-67 | 0.0 (0/19) | 0.5 | 0.0 (0/0) | 1.7 | 77.7 | 14 |
| 1423 | RF01523 TB10Cs1H2 | 0.0 (0/19) | 0.5 | 0.0 (0/0) | 1.8 | 79.8 | 5 |
| 1424 | RF00261 IRES L-myc | 0.0 (0/75) | 0.5 | 0.0 (0/0) | 1.9 | 88.6 | 11 |
| 1425 | RF02722 sca ncR27 | 0.0 (0/22) | 0.5 | 0.0 (0/0) | 1.5 | 80.3 | 4 |
| 1426 | RF02524 sagA | 0.0 (0/41) | 0.5 | 0.0 (0/0) | 2.1 | 69.4 | 6 |
| 1427 | RF02270 nse sRNA | 0.0 (0/19) | 0.5 | 0.0 (0/0) | 0.9 | 90.1 | 7 |
| 1428 | RF00778 MIR473 | 0.0 (0/20) | 0.5 | 0.0 (0/0) | 1.8 | 59.0 | 6 |
| 1429 | RF01940 hvt-mir-H | 0.0 (0/20) | 0.5 | 0.0 (0/0) | 2.5 | 67.2 | 4 |
| 1430 | RF02093 mir-2968 | 0.0 (0/39) | 0.5 | 0.0 (0/0) | 2.2 | 87.8 | 12 |
| 1431 | RF02231 sX12 | 0.0 (0/21) | 0.5 | 0.0 (0/0) | 0.8 | 94.4 | 8 |
| 1432 | RF00254 mir-16 | 0.0 (0/19) | 0.5 | 0.0 (0/0) | 1.3 | 72.9 | 12 |
| 1433 | RF00689 MIR390 | 0.0 (0/20) | 0.5 | 0.0 (0/0) | 1.1 | 73.8 | 16 |
| 1434 | RF02883 BcKCsr2 | 0.0 (0/38) | 0.5 | 0.0 (0/0) | 2.3 | 72.7 | 5 |
| 1435 | RF01771 rnk leader | 0.0 (0/19) | 0.5 | 0.0 (0/0) | 2.7 | 74.4 | 13 |
| 1436 | RF02453 ncr952 | 0.0 (0/38) | 0.5 | 0.0 (0/0) | 2.0 | 83.3 | 6 |
| 1437 | RF00264 SNORA64 | 0.0 (0/37) | 0.5 | 0.0 (0/0) | 2.5 | 70.8 | 9 |
| 1438 | RF00942 mir-1224 | 0.0 (0/22) | 0.5 | 0.0 (0/0) | 0.7 | 85.9 | 11 |
| 1439 | RF02711 TeloSII ncR49 | 0.0 (0/24) | 0.4 | 0.0 (0/0) | 1.5 | 92.5 | 3 |
| 1440 | RF00897 mir-675 | 0.0 (0/26) | 0.4 | 0.0 (0/0) | 1.8 | 85.2 | 11 |
| 1441 | RF00703 mir-139 | 0.0 (0/24) | 0.4 | 0.0 (0/0) | 1.0 | 81.0 | 5 |
| 1442 | RF00035 OxyS | 0.0 (0/26) | 0.4 | 0.0 (0/0) | 1.8 | 90.0 | 5 |
| 1443 | RF00692 MIR171 2 | 0.0 (0/26) | 0.4 | 0.0 (0/0) | 1.3 | 70.5 | 7 |
| 1444 | RF00549 IRES c-sis | 0.0 (0/201) | 0.4 | 0.0 (0/1) | 1.9 | 93.9 | 10 |
| 1445 | RF00690 MIR408 | 0.0 (0/24) | 0.4 | 0.0 (0/0) | 1.4 | 68.4 | 8 |
| 1446 | RF00968 mir-626 | 0.0 (0/26) | 0.4 | 0.0 (0/0) | 2.0 | 73.8 | 6 |
| 1447 | RF02111 IS009 | 0.0 (0/23) | 0.4 | 0.0 (0/0) | 2.2 | 76.0 | 11 |
| 1448 | RF00677 MIR168 | 0.0 (0/23) | 0.4 | 0.0 (0/0) | 1.8 | 66.8 | 10 |
| 1449 | RF00943 MIR824 | 0.0 (0/53) | 0.4 | 0.0 (0/0) | 0.8 | 90.7 | 12 |
| 1450 | RF00882 MIR811 | 0.0 (0/81) | 0.4 | 0.0 (0/1) | 1.2 | 81.2 | 5 |
| 1451 | RF00710 mir-44 | 0.0 (0/27) | 0.4 | 0.0 (0/0) | 1.1 | 75.2 | 6 |
| 1452 | RF02225 sX6 | 0.0 (0/68) | 0.4 | 0.0 (0/2) | 1.3 | 82.8 | 8 |
| 1453 | RF00684 mir-122 | 0.0 (0/24) | 0.4 | 0.0 (0/0) | 0.8 | 92.2 | 26 |
| 1454 | RF01044 mir-345 | 0.0 (0/27) | 0.4 | 0.0 (0/0) | 1.6 | 85.7 | 10 |
| 1455 | RF00243 traJ 5 | 0.0 (0/27) | 0.4 | 0.0 (0/0) | 1.0 | 86.2 | 6 |
| 1456 | RF01043 MIR1023 | 0.0 (0/46) | 0.4 | 0.0 (0/0) | 1.9 | 59.8 | 4 |
| 1457 | RF01827 SAR11 0636 | 0.0 (0/23) | 0.4 | 0.0 (0/0) | 2.1 | 91.3 | 13 |
| 1458 | RF02773 TrxA thermometer | 0.0 (0/25) | 0.4 | 0.0 (0/0) | 1.4 | 87.5 | 8 |
| 1459 | RF00258 mir-130 | 0.0 (0/26) | 0.4 | 0.0 (0/0) | 1.4 | 86.7 | 9 |
| 1460 | RF00486 mir-129 | 0.0 (0/25) | 0.4 | 0.0 (0/1) | 1.0 | 83.0 | 6 |
| 1461 | RF02605 scr5239 | 0.0 (0/46) | 0.4 | 0.0 (0/0) | 1.5 | 82.7 | 6 |
| 1462 | RF02405 P34 | 0.0 (0/99) | 0.4 | 0.0 (0/0) | 2.9 | 68.0 | 5 |
| 1463 | RF00307 snoR98 | 0.0 (0/24) | 0.4 | 0.0 (0/0) | 1.7 | 86.7 | 4 |
| 1464 | RF02528 SSRC41 | 0.0 (0/25) | 0.4 | 0.0 (0/0) | 2.0 | 81.0 | 8 |
| 1465 | RF02895 S414 | 0.0 (0/36) | 0.3 | 0.0 (0/0) | 1.6 | 71.5 | 5 |
| 1466 | RF00878 mir-456 | 0.0 (0/32) | 0.3 | 0.0 (0/0) | 1.8 | 68.8 | 5 |
| 1467 | RF00777 mir-541 | 0.0 (0/30) | 0.3 | 0.0 (0/1) | 2.4 | 82.6 | 10 |
| 1468 | RF01635 ceN45 | 0.0 (0/37) | 0.3 | 0.0 (0/0) | 1.6 | 78.5 | 4 |
| 1469 | RF02268 MtlS | 0.0 (0/38) | 0.3 | 0.0 (0/0) | 1.9 | 76.4 | 6 |
| 1470 | RF01436 S pombe snR33 | 0.0 (0/33) | 0.3 | 0.0 (0/0) | 1.5 | 68.7 | 3 |
| 1471 | RF02673 scr4677 | 0.0 (0/36) | 0.3 | 0.0 (0/1) | 0.6 | 74.8 | 12 |
| 1472 | RF00807 mir-314 | 0.0 (0/32) | 0.3 | 0.0 (0/0) | 1.2 | 84.1 | 6 |
| 1473 | RF02569 IhtA | 0.0 (0/34) | 0.3 | 0.0 (0/0) | 1.7 | 83.6 | 5 |
| 1474 | RF01661 ceN92 | 0.0 (0/30) | 0.3 | 0.0 (0/0) | 1.3 | 75.6 | 4 |
| 1475 | RF00740 mir-370 | 0.0 (0/29) | 0.3 | 0.0 (0/0) | 1.2 | 91.3 | 6 |
| 1476 | RF00432 SNORA51 | 0.0 (0/29) | 0.3 | 0.0 (0/0) | 2.6 | 82.3 | 9 |
| 1477 | RF00594 SNORD86 | 0.0 (0/30) | 0.3 | 0.0 (0/0) | 2.1 | 77.2 | 6 |
| 1478 | RF00911 mir-672 | 0.0 (0/30) | 0.3 | 0.0 (0/0) | 2.4 | 84.2 | 7 |
| 1479 | RF02692 BTH s39 | 0.0 (0/34) | 0.3 | 0.0 (0/0) | 0.8 | 89.4 | 7 |
| 1480 | RF00272 SNORA67 | 0.0 (0/33) | 0.3 | 0.0 (0/1) | 1.9 | 80.2 | 12 |
| 1481 | RF00755 mir-542 | 0.0 (0/29) | 0.3 | 0.0 (0/0) | 0.9 | 89.9 | 5 |
| 1482 | RF01435 S pombe snR5 | 0.0 (0/34) | 0.3 | 0.0 (0/0) | 1.3 | 81.4 | 3 |
| 1483 | RF01444 S pombe snR92 | 0.0 (0/36) | 0.3 | 0.0 (0/0) | 1.5 | 70.4 | 3 |
| 1484 | RF01943 mir-999 | 0.0 (0/32) | 0.3 | 0.0 (0/0) | 2.0 | 65.0 | 4 |
| 1485 | RF00937 mir-653 | 0.0 (0/32) | 0.3 | 0.0 (0/0) | 1.4 | 84.9 | 12 |
| 1486 | RF01431 snoR135 | 0.0 (0/28) | 0.3 | 0.0 (0/0) | 1.3 | 73.5 | 6 |
| 1487 | RF00808 mir-86 | 0.0 (0/37) | 0.3 | 0.0 (0/0) | 1.6 | 79.5 | 5 |
| 1488 | RF00775 mir-432 | 0.0 (0/28) | 0.3 | 0.0 (0/0) | 1.9 | 73.4 | 6 |
| 1489 | RF00766 mir-335 | 0.0 (0/32) | 0.3 | 0.0 (0/0) | 0.9 | 90.2 | 9 |
| 1490 | RF00848 mir-61 | 0.0 (0/32) | 0.3 | 0.0 (0/0) | 1.5 | 73.1 | 4 |
| 1491 | RF00726 mir-87 | 0.0 (0/30) | 0.3 | 0.0 (0/0) | 0.8 | 85.9 | 8 |
| 1492 | RF00884 MIR815 | 0.0 (0/32) | 0.3 | 0.0 (0/0) | 1.6 | 79.5 | 3 |
| 1493 | RF00680 mir-224 | 0.0 (0/32) | 0.3 | 0.0 (0/0) | 1.2 | 86.7 | 5 |
| 1494 | RF00788 mir-287 | 0.0 (0/29) | 0.3 | 0.0 (0/0) | 2.0 | 82.1 | 9 |
| 1495 | RF02850 Ysr276 | 0.0 (0/56) | 0.3 | 0.0 (0/0) | 1.6 | 83.6 | 5 |
| 1496 | RF00837 mir-251 | 0.0 (0/34) | 0.3 | 0.0 (0/0) | 0.7 | 80.1 | 4 |
| 1497 | RF01607 ceN100 | 0.0 (0/35) | 0.3 | 0.0 (0/3) | 1.3 | 79.8 | 4 |
| 1498 | RF02409 snoR125 | 0.0 (0/65) | 0.3 | 0.0 (0/0) | 1.3 | 86.4 | 5 |
| 1499 | RF01669 P14 | 0.0 (0/36) | 0.3 | 0.0 (0/4) | 1.8 | 71.8 | 4 |
| 1500 | RF02245 mir-788 | 0.0 (0/28) | 0.3 | 0.0 (0/0) | 1.1 | 75.4 | 4 |
| 1501 | RF00850 mir-259 | 0.0 (0/30) | 0.3 | 0.0 (0/0) | 0.8 | 75.3 | 4 |

Continued on next page

| | RNA family (seed alignment) | Sensitivity annotated bpairs that covary % (cov_bps/bps) | Power average power % | Positive Predictive Value covarying pairs in structure % (cov_bps/cov_pairs) | average substitutions per bpair | avg pairwise identity % | number of sequences |
|---|---|---|---|---|---|---|---|
| 1502 | RF01261 snR82 | 0.0 (0/72) | 0.3 | 0.0 (0/0) | 1.8 | 78.6 | 5 |
| 1503 | RF00787 mir-288 | 0.0 (0/34) | 0.3 | 0.0 (0/0) | 1.4 | 90.5 | 7 |
| 1504 | RF01240 snR85 | 0.0 (0/36) | 0.3 | 0.0 (0/0) | 0.9 | 80.6 | 5 |
| 1505 | RF02820 V AS9 | 0.0 (0/33) | 0.3 | 0.0 (0/0) | 1.2 | 73.5 | 3 |
| 1506 | RF00855 mir-254 | 0.0 (0/34) | 0.3 | 0.0 (0/0) | 1.0 | 74.9 | 4 |
| 1507 | RF01629 ceN39 | 0.0 (0/39) | 0.2 | 0.0 (0/0) | 0.6 | 90.3 | 4 |
| 1508 | RF02855 Ysr251 | 0.0 (0/52) | 0.2 | 0.0 (0/0) | 1.4 | 77.1 | 5 |
| 1509 | RF02404 P33 | 0.0 (0/45) | 0.2 | 0.0 (0/0) | 0.9 | 91.4 | 3 |
| 1510 | RF02718 sca ncR14 | 0.0 (0/41) | 0.2 | 0.0 (0/0) | 1.0 | 88.8 | 4 |
| 1511 | RF00232 Spi-1 | 0.0 (0/53) | 0.2 | 0.0 (0/0) | 1.2 | 90.9 | 5 |
| 1512 | RF02830 Scr1601 | 0.0 (0/61) | 0.2 | 0.0 (0/0) | 0.8 | 80.1 | 6 |
| 1513 | RF00499 Parecho CRE | 0.0 (0/43) | 0.2 | 0.0 (0/0) | 1.4 | 87.2 | 5 |
| 1514 | RF02872 SrbA | 0.0 (0/55) | 0.2 | 0.0 (0/0) | 1.6 | 62.8 | 3 |
| 1515 | RF02836 Bcj14 | 0.0 (0/55) | 0.2 | 0.0 (0/0) | 0.8 | 82.3 | 3 |
| 1516 | RF02566 Ms1 | 0.0 (0/96) | 0.2 | 0.0 (0/0) | 0.9 | 85.3 | 4 |
| 1517 | RF02223 sX4 | 0.0 (0/42) | 0.2 | 0.0 (0/0) | 2.0 | 85.0 | 5 |
| 1518 | RF01674 P27 | 0.0 (0/56) | 0.2 | 0.0 (0/0) | 0.8 | 85.3 | 4 |
| 1519 | RF01400 istR | 0.0 (0/44) | 0.2 | 0.0 (0/1) | 2.0 | 84.1 | 7 |
| 1520 | RF02868 ncS037 | 0.0 (0/49) | 0.2 | 0.0 (0/0) | 1.2 | 74.0 | 3 |
| 1521 | RF01417 RSV RNA | 0.0 (0/53) | 0.2 | 0.0 (0/0) | 1.3 | 88.5 | 8 |
| 1522 | RF02690 BTH s1 | 0.0 (0/62) | 0.2 | 0.0 (0/0) | 0.9 | 84.3 | 5 |
| 1523 | RF02829 Scr4115 | 0.0 (0/39) | 0.2 | 0.0 (0/0) | 1.0 | 87.8 | 6 |
| 1524 | RF00958 mir-498 | 0.0 (0/43) | 0.2 | 0.0 (0/0) | 2.2 | 65.6 | 5 |
| 1525 | RF01865 Vg1 ribozyme | 0.0 (0/41) | 0.2 | 0.0 (0/3) | 2.3 | 72.7 | 4 |
| 1526 | RF02768 Ysr155 RyfD | 0.0 (0/47) | 0.2 | 0.0 (0/0) | 1.5 | 85.1 | 9 |
| 1527 | RF01262 snR44 | 0.0 (0/55) | 0.2 | 0.0 (0/0) | 1.7 | 86.0 | 7 |
| 1528 | RF02675 Ysr141 | 0.0 (0/80) | 0.1 | 0.0 (0/0) | 1.2 | 84.5 | 4 |
| 1529 | RF01271 snR30 | 0.0 (0/184) | 0.1 | 0.0 (0/0) | 0.7 | 90.8 | 5 |
| 1530 | RF02746 AsrC | 0.0 (0/266) | 0.1 | 0.0 (0/3) | 1.4 | 85.4 | 7 |
| 1531 | RF02897 S808 | 0.0 (0/72) | 0.1 | 0.0 (0/0) | 1.4 | 82.4 | 4 |
| 1532 | RF01825 RUF21 | 0.0 (0/74) | 0.1 | 0.0 (0/0) | 1.5 | 67.6 | 5 |
| 1533 | RF00216 IRES c-myc | 0.0 (0/73) | 0.1 | 0.0 (0/0) | 1.1 | 98.0 | 23 |
| 1534 | RF02400 Hsp83 3 UTR | 0.0 (0/143) | 0.1 | 0.0 (0/0) | 0.6 | 91.2 | 5 |
| 1535 | RF01494 rliD | 0.0 (0/83) | 0.1 | 0.0 (0/0) | 0.5 | 96.9 | 9 |
| 1536 | RF01272 snR86 | 0.0 (0/334) | 0.1 | 0.0 (0/0) | 1.5 | 73.9 | 5 |
| 1537 | RF01395 isrL | 0.0 (0/94) | 0.1 | 0.0 (0/0) | 1.1 | 84.9 | 4 |
| 1538 | RF01264 snR83 | 0.0 (0/85) | 0.1 | 0.0 (0/1) | 1.1 | 87.1 | 5 |
| 1539 | RF02574 Ricks sRNA10 | 0.0 (0/89) | 0.1 | 0.0 (0/1) | 1.0 | 73.2 | 3 |
| 1540 | RF01265 snR42 | 0.0 (0/107) | 0.1 | 0.0 (0/0) | 1.0 | 84.2 | 5 |
| 1541 | RF00224 IRES FGF2 | 0.0 (0/96) | 0.1 | 0.0 (0/1) | 0.9 | 89.5 | 6 |
| 1542 | RF01397 isrO | 0.0 (0/64) | 0.1 | 0.0 (0/0) | 2.2 | 84.7 | 6 |
| 1543 | RF00908 MIR529 | 0.0 (0/28) | 0.0 | 0.0 (0/0) | 0.4 | 76.5 | 3 |
| 1544 | RF02074 STnc240 | 0.0 (0/11) | 0.0 | 0.0 (0/0) | 1.9 | 76.5 | 15 |
| 1545 | RF01003 mir-563 | 0.0 (0/28) | 0.0 | 0.0 (0/0) | 2.6 | 75.7 | 7 |
| 1546 | RF00285 snoZ6 | 0.0 (0/4) | 0.0 | 0.0 (0/0) | 2.0 | 91.2 | 7 |
| 1547 | RF00804 mir-240 | 0.0 (0/30) | 0.0 | 0.0 (0/1) | 0.7 | 69.7 | 3 |
| 1548 | RF02159 PART1 1 | 0.0 (0/0) | 0.0 | 0.0 (0/0) | 0.0 | 78.8 | 22 |
| 1549 | RF01441 S pombe snR46 | 0.0 (0/39) | 0.0 | 0.0 (0/0) | 0.0 | 100.0 | 2 |
| 1550 | RF02167 PVT1 4 | 0.0 (0/0) | 0.0 | 0.0 (0/0) | 0.0 | 70.4 | 10 |
| 1551 | RF01886 HSR-omega 2 | 0.0 (0/0) | 0.0 | 0.0 (0/0) | 0.0 | 85.1 | 10 |
| 1552 | RF02314 TtnuHACA7 | 0.0 (0/32) | 0.0 | 0.0 (0/0) | 0.3 | 76.6 | 2 |
| 1553 | RF01983 Pinc | 0.0 (0/0) | 0.0 | 0.0 (0/0) | 0.0 | 84.4 | 16 |
| 1554 | RF01946 KCNQ1OT1 1 | 0.0 (0/0) | 0.0 | 0.0 (0/0) | 0.0 | 84.0 | 8 |
| 1555 | RF01863 TB10Cs2H2 | 0.0 (0/16) | 0.0 | 0.0 (0/0) | 0.0 | 100.0 | 3 |
| 1556 | RF01164 SNORD107 | 0.0 (0/4) | 0.0 | 0.0 (0/0) | 1.2 | 86.2 | 13 |
| 1557 | RF02056 STnc390 | 0.0 (0/15) | 0.0 | 0.0 (0/0) | 0.0 | 96.6 | 2 |
| 1558 | RF02158 NPPA-AS1 3 | 0.0 (0/0) | 0.0 | 0.0 (0/0) | 0.0 | 74.7 | 24 |
| 1559 | RF01377 CRISPR-DR64 | 0.0 (0/4) | 0.0 | 0.0 (0/0) | 0.0 | 97.3 | 2 |
| 1560 | RF02028 mir-1827 | 0.0 (0/21) | 0.0 | 0.0 (0/0) | 0.9 | 86.3 | 6 |
| 1561 | RF02623 BSnc140 | 0.0 (0/21) | 0.0 | 0.0 (0/0) | 0.0 | 98.9 | 2 |
| 1562 | RF02492 Gl U2 | 0.0 (0/27) | 0.0 | 0.0 (0/0) | 0.0 | 95.6 | 2 |
| 1563 | RF00729 mir-278 | 0.0 (0/27) | 0.0 | 0.0 (0/0) | 1.1 | 78.6 | 9 |
| 1564 | RF01923 mir-711 | 0.0 (0/24) | 0.0 | 0.0 (0/0) | 0.7 | 82.8 | 5 |
| 1565 | RF00502 TCV Pr | 0.0 (0/8) | 0.0 | 0.0 (0/0) | 0.2 | 91.9 | 4 |
| 1566 | RF02742 Rev72 | 0.0 (0/122) | 0.0 | 0.0 (0/0) | 0.5 | 86.8 | 3 |
| 1567 | RF02874 AS-traG | 0.0 (0/24) | 0.0 | 0.0 (0/0) | 1.3 | 76.4 | 4 |
| 1568 | RF00635 HAR1A | 0.0 (0/0) | 0.0 | 0.0 (0/0) | 0.0 | 92.0 | 13 |
| 1569 | RF01041 mir-604 | 0.0 (0/30) | 0.0 | 0.0 (0/0) | 0.1 | 94.7 | 2 |
| 1570 | RF00956 MIR1444 | 0.0 (0/31) | 0.0 | 0.0 (0/0) | 0.4 | 82.3 | 3 |
| 1571 | RF00201 snoZ278 | 0.0 (0/8) | 0.0 | 0.0 (0/0) | 0.8 | 94.1 | 7 |
| 1572 | RF01486 rli62 | 0.0 (0/33) | 0.0 | 0.0 (0/0) | 0.3 | 58.8 | 2 |
| 1573 | RF02046 Sphinx 1 | 0.0 (0/0) | 0.0 | 0.0 (0/0) | 0.0 | 88.0 | 3 |
| 1574 | RF00652 MIR478 | 0.0 (0/0) | 0.0 | 0.0 (0/0) | 0.0 | 89.6 | 17 |
| 1575 | RF01987 ZEB2 AS1 4 | 0.0 (0/0) | 0.0 | 0.0 (0/1) | 0.0 | 90.6 | 13 |
| 1576 | RF01842 mycoplasma FSE | 0.0 (0/11) | 0.0 | 0.0 (0/0) | 0.0 | 94.6 | 3 |
| 1577 | RF02240 Xoo1 | 0.0 (0/24) | 0.0 | 0.0 (0/0) | 1.0 | 85.2 | 5 |
| 1578 | RF02098 DGCR5 | 0.0 (0/0) | 0.0 | 0.0 (0/0) | 0.0 | 87.8 | 4 |
| 1579 | RF01795 FourU | 0.0 (0/21) | 0.0 | 0.0 (0/0) | 0.0 | 92.3 | 3 |
| 1580 | RF00771 mir-185 | 0.0 (0/34) | 0.0 | 0.0 (0/0) | 0.6 | 91.7 | 5 |
| 1581 | RF02555 hveRNA | 0.0 (0/31) | 0.0 | 0.0 (0/0) | 0.7 | 100.0 | 7 |
| 1582 | RF01531 TB10Cs3H1 | 0.0 (0/18) | 0.0 | 0.0 (0/0) | 0.8 | 80.7 | 4 |
| 1583 | RF00384 Pox AX element | 0.0 (0/22) | 0.0 | 0.0 (0/0) | 0.4 | 93.4 | 10 |
| 1584 | RF01279 snoR53Y | 0.0 (0/5) | 0.0 | 0.0 (0/0) | 1.2 | 71.8 | 7 |
| 1585 | RF00762 mir-412 | 0.0 (0/34) | 0.0 | 0.0 (0/0) | 0.8 | 90.7 | 9 |

Continued on next page

| | RNA family (seed alignment) | Sensitivity annotated bpairs that covary % (cov_bps/bps) | Power average power % | Positive Predictive Value covarying pairs in structure % (cov_bps/cov_pairs) | average substitutions per bpair | avg pairwise identity % | number of sequences |
|---|---|---|---|---|---|---|---|
| 1586 | RF02019 mir-1265 | 0.0 (0/35) | 0.0 | 0.0 (0/0) | 0.1 | 95.5 | 3 |
| 1587 | RF00046 snoR30 | 0.0 (0/4) | 0.0 | 0.0 (0/0) | 1.5 | 87.0 | 6 |
| 1588 | RF02247 Six3os1 2 | 0.0 (0/0) | 0.0 | 0.0 (0/0) | 0.0 | 85.6 | 15 |
| 1589 | RF01330 CRISPR-DR16 | 0.0 (0/7) | 0.0 | 0.0 (0/0) | 0.7 | 87.0 | 6 |
| 1590 | RF02439 SpF56 sRNA | 0.0 (0/24) | 0.0 | 0.0 (0/0) | 0.0 | 94.9 | 2 |
| 1591 | RF01601 plasmodium snoR26 | 0.0 (0/8) | 0.0 | 0.0 (0/0) | 0.0 | 92.8 | 3 |
| 1592 | RF02499 Atu C4 | 0.0 (0/21) | 0.0 | 0.0 (0/0) | 0.0 | 98.6 | 2 |
| 1593 | RF02782 CpoB ybgF thermometer | 0.0 (0/38) | 0.0 | 0.0 (0/0) | 0.9 | 91.5 | 8 |
| 1594 | RF01626 ceN30 | 0.0 (0/3) | 0.0 | 0.0 (0/0) | 0.0 | 89.8 | 4 |
| 1595 | RF02766 Ysr49 | 0.0 (0/30) | 0.0 | 0.0 (0/0) | 0.4 | 93.5 | 3 |
| 1596 | RF01007 mir-624 | 0.0 (0/38) | 0.0 | 0.0 (0/0) | 0.0 | 96.9 | 2 |
| 1597 | RF02762 C1 109596F thermometer | 0.0 (0/60) | 0.0 | 0.0 (0/0) | 0.0 | 98.7 | 2 |
| 1598 | RF01423 snoR117 | 0.0 (0/4) | 0.0 | 0.0 (0/0) | 1.5 | 69.3 | 6 |
| 1599 | RF02669 PsiU6-40 | 0.0 (0/46) | 0.0 | 0.0 (0/0) | 0.1 | 96.2 | 3 |
| 1600 | RF01585 snoR07 | 0.0 (0/3) | 0.0 | 0.0 (0/0) | 0.0 | 85.9 | 4 |
| 1601 | RF02783 IscS1 thermometer | 0.0 (0/46) | 0.0 | 0.0 (0/0) | 1.4 | 78.6 | 5 |
| 1602 | RF01355 CRISPR-DR26 | 0.0 (0/3) | 0.0 | 0.0 (0/0) | 0.0 | 93.3 | 4 |
| 1603 | RF01125 sR4 | 0.0 (0/0) | 0.0 | 0.0 (0/0) | 0.0 | 86.9 | 3 |
| 1604 | RF02859 Arrc11 | 0.0 (0/53) | 0.0 | 0.0 (0/0) | 0.2 | 77.2 | 2 |
| 1605 | RF01132 sR48 | 0.0 (0/0) | 0.0 | 0.0 (0/0) | 0.0 | 76.4 | 2 |
| 1606 | RF02233 sX14 | 0.0 (0/27) | 0.0 | 0.0 (0/0) | 0.1 | 96.0 | 8 |
| 1607 | RF01551 TB8Cs4H2 | 0.0 (0/15) | 0.0 | 0.0 (0/0) | 0.1 | 69.1 | 2 |
| 1608 | RF02292 TtnuCD16 | 0.0 (0/0) | 0.0 | 0.0 (0/0) | 0.0 | 96.5 | 2 |
| 1609 | RF01298 snoU25 | 0.0 (0/9) | 0.0 | 0.0 (0/0) | 0.0 | 98.1 | 3 |
| 1610 | RF02724 sno ZL2 | 0.0 (0/25) | 0.0 | 0.0 (0/0) | 1.2 | 86.5 | 6 |
| 1611 | RF02058 STnc400 | 0.0 (0/43) | 0.0 | 0.0 (0/0) | 0.5 | 65.3 | 2 |
| 1612 | RF02482 GlsR19 | 0.0 (0/21) | 0.0 | 0.0 (0/0) | 0.8 | 83.5 | 3 |
| 1613 | RF01159 snoU18 | 0.0 (0/0) | 0.0 | 0.0 (0/0) | 0.0 | 82.1 | 16 |
| 1614 | RF00990 mir-552 | 0.0 (0/34) | 0.0 | 0.0 (0/0) | 0.0 | 92.6 | 2 |
| 1615 | RF02256 adapt33 2 | 0.0 (0/0) | 0.0 | 0.0 (0/0) | 0.0 | 82.6 | 2 |
| 1616 | RF01283 snoU30 | 0.0 (0/2) | 0.0 | 0.0 (0/0) | 0.0 | 74.8 | 5 |
| 1617 | RF00571 SNORD65 | 0.0 (0/0) | 0.0 | 0.0 (0/0) | 0.0 | 82.6 | 26 |
| 1618 | RF02567 Va-907 | 0.0 (0/100) | 0.0 | 0.0 (0/0) | 0.2 | 95.9 | 3 |
| 1619 | RF02048 STnc30 | 0.0 (0/33) | 0.0 | 0.0 (0/0) | 0.1 | 94.7 | 2 |
| 1620 | RF02710 TeloSII ncR45 | 0.0 (0/26) | 0.0 | 0.0 (0/0) | 0.0 | 98.5 | 2 |
| 1621 | RF01425 snoR121 | 0.0 (0/3) | 0.0 | 0.0 (0/0) | 0.0 | 88.2 | 2 |
| 1622 | RF02181 ST7-AS2 1 | 0.0 (0/0) | 0.0 | 0.0 (0/0) | 0.0 | 89.9 | 4 |
| 1623 | RF00610 SNORD110 | 0.0 (0/4) | 0.0 | 0.0 (0/0) | 1.0 | 87.7 | 16 |
| 1624 | RF01477 rli43 | 0.0 (0/69) | 0.0 | 0.0 (0/0) | 0.4 | 93.5 | 4 |
| 1625 | RF02587 HCV package-SL4629 | 0.0 (0/7) | 0.0 | 0.0 (0/0) | 0.0 | 95.7 | 2 |
| 1626 | RF01483 rli56 | 0.0 (0/31) | 0.0 | 0.0 (0/0) | 0.0 | 97.5 | 6 |
| 1627 | RF02129 GNAS-AS1 3 | 0.0 (0/0) | 0.0 | 0.0 (0/0) | 0.0 | 83.8 | 9 |
| 1628 | RF01083 UPD-PK2 | 0.0 (0/8) | 0.0 | 0.0 (0/0) | 0.6 | 95.2 | 4 |
| 1629 | RF01988 SECIS 2 | 0.0 (0/20) | 0.0 | 0.0 (0/0) | 0.8 | 89.1 | 4 |
| 1630 | RF00039 DicF | 0.0 (0/11) | 0.0 | 0.0 (0/0) | 0.5 | 74.6 | 5 |
| 1631 | RF02839 Ref66 | 0.0 (0/34) | 0.0 | 0.0 (0/0) | 0.1 | 80.2 | 2 |
| 1632 | RF02657 sot2652 | 0.0 (0/58) | 0.0 | 0.0 (0/0) | 0.2 | 86.0 | 2 |
| 1633 | RF02763 IsrM | 0.0 (0/81) | 0.0 | 0.0 (0/0) | 0.1 | 100.0 | 2 |
| 1634 | RF02660 icaR 3p UTR | 0.0 (0/126) | 0.0 | 0.0 (0/0) | 0.0 | 99.7 | 3 |
| 1635 | RF01514 Afu 513 | 0.0 (0/0) | 0.0 | 0.0 (0/0) | 0.0 | 77.4 | 7 |
| 1636 | RF02372 PyrC leader | 0.0 (0/5) | 0.0 | 0.0 (0/0) | 0.2 | 81.7 | 9 |
| 1637 | RF02863 sRNA 2410 | 0.0 (0/57) | 0.0 | 0.0 (0/0) | 0.7 | 74.7 | 3 |
| 1638 | RF01210 snoU13 | 0.0 (0/0) | 0.0 | 0.0 (0/0) | 0.0 | 78.1 | 34 |
| 1639 | RF02568 UptR | 0.0 (0/20) | 0.0 | 0.0 (0/0) | 0.2 | 96.3 | 3 |
| 1640 | RF01896 mir-142 | 0.0 (0/31) | 0.0 | 0.0 (0/0) | 0.1 | 93.4 | 13 |
| 1641 | RF02556 snaR-A | 0.0 (0/38) | 0.0 | 0.0 (0/0) | 0.3 | 95.8 | 5 |
| 1642 | RF02328 TtnuHACA21 | 0.0 (0/34) | 0.0 | 0.0 (0/0) | 0.1 | 95.3 | 2 |
| 1643 | RF01367 CRISPR-DR54 | 0.0 (0/6) | 0.0 | 0.0 (0/0) | 0.0 | 97.3 | 2 |
| 1644 | RF02062 STnc361 | 0.0 (0/35) | 0.0 | 0.0 (0/0) | 0.2 | 79.9 | 2 |
| 1645 | RF02072 STnc590 | 0.0 (0/24) | 0.0 | 0.0 (0/0) | 0.4 | 96.2 | 3 |
| 1646 | RF01944 mir-2518 | 0.0 (0/29) | 0.0 | 0.0 (0/0) | 0.2 | 97.2 | 4 |
| 1647 | RF01579 RUF2 | 0.0 (0/42) | 0.0 | 0.0 (0/0) | 0.4 | 79.1 | 3 |
| 1648 | RF00758 mir-346 | 0.0 (0/27) | 0.0 | 0.0 (0/0) | 0.2 | 93.2 | 5 |
| 1649 | RF02316 TtnuHACA9 | 0.0 (0/37) | 0.0 | 0.0 (0/2) | 0.5 | 79.3 | 3 |
| 1650 | RF02291 TtnuCD15 | 0.0 (0/0) | 0.0 | 0.0 (0/0) | 0.0 | 88.5 | 2 |
| 1651 | RF01166 sn3071 | 0.0 (0/4) | 0.0 | 0.0 (0/0) | 0.0 | 94.7 | 2 |
| 1652 | RF01500 Afu 198 | 0.0 (0/0) | 0.0 | 0.0 (0/0) | 0.0 | 57.3 | 6 |
| 1653 | RF00259 IFN gamma | 0.0 (0/56) | 0.0 | 0.0 (0/0) | 0.6 | 90.4 | 5 |
| 1654 | RF02018 mir-1207 | 0.0 (0/28) | 0.0 | 0.0 (0/0) | 0.1 | 97.7 | 3 |
| 1655 | RF01286 snoR26 | 0.0 (0/7) | 0.0 | 0.0 (0/0) | 0.4 | 74.1 | 8 |
| 1656 | RF01442 S pombe snR90 | 0.0 (0/36) | 0.0 | 0.0 (0/0) | 1.2 | 80.8 | 3 |
| 1657 | RF02406 snoR120 | 0.0 (0/0) | 0.0 | 0.0 (0/0) | 0.0 | 87.7 | 4 |
| 1658 | RF00268 snoZ7 | 0.0 (0/2) | 0.0 | 0.0 (0/0) | 0.0 | 90.5 | 5 |
| 1659 | RF02390 sau-41 | 0.0 (0/35) | 0.0 | 0.0 (0/0) | 0.0 | 94.2 | 2 |
| 1660 | RF00862 mir-491 | 0.0 (0/33) | 0.0 | 0.0 (0/0) | 1.0 | 88.8 | 5 |
| 1661 | RF01303 sR49 | 0.0 (0/11) | 0.0 | 0.0 (0/0) | 0.0 | 87.5 | 3 |
| 1662 | RF02351 psRNA14 | 0.0 (0/35) | 0.0 | 0.0 (0/0) | 1.7 | 58.2 | 3 |
| 1663 | RF01172 sn2343 | 0.0 (0/4) | 0.0 | 0.0 (0/0) | 0.0 | 88.5 | 3 |
| 1664 | RF00681 mir-198 | 0.0 (0/22) | 0.0 | 0.0 (0/0) | 0.3 | 90.3 | 3 |
| 1665 | RF01281 snoR35 | 0.0 (0/5) | 0.0 | 0.0 (0/0) | 2.4 | 69.9 | 12 |
| 1666 | RF02582 tsr33 | 0.0 (0/20) | 0.0 | 0.0 (0/0) | 0.1 | 95.3 | 2 |
| 1667 | RF02815 Lig thermometer | 0.0 (0/57) | 0.0 | 0.0 (0/0) | 0.0 | 99.5 | 2 |
| 1668 | RF00877 mir-592 | 0.0 (0/34) | 0.0 | 0.0 (0/0) | 0.7 | 89.1 | 5 |
| 1669 | RF01307 sR55 | 0.0 (0/2) | 0.0 | 0.0 (0/0) | 0.0 | 87.1 | 4 |

Continued on next page

| | RNA family (seed alignment) | Sensitivity annotated bpairs that covary % (cov_bps/bps) | Power average power % | Positive Predictive Value covarying pairs in structure % (cov_bps/cov_pairs) | average substitutions per bpair | avg pairwise identity % | number of sequences |
|---|---|---|---|---|---|---|---|
| 1670 | RF02321 TtnuHACA14 | 0.0 (0/36) | 0.0 | 0.0 (0/0) | 0.2 | 92.0 | 3 |
| 1671 | RF01665 P13 | 0.0 (0/26) | 0.0 | 0.0 (0/0) | 0.0 | 88.2 | 2 |
| 1672 | RF02563 CbSR3 | 0.0 (0/51) | 0.0 | 0.0 (0/0) | 0.0 | 99.5 | 2 |
| 1673 | RF00383 IS1222 FSE | 0.0 (0/17) | 0.0 | 0.0 (0/0) | 0.8 | 92.7 | 5 |
| 1674 | RF02202 UCA1 | 0.0 (0/0) | 0.0 | 0.0 (0/0) | 0.0 | 94.3 | 5 |
| 1675 | RF02170 PVT1 7 | 0.0 (0/0) | 0.0 | 0.0 (0/0) | 0.0 | 65.0 | 8 |
| 1676 | RF02394 sau-63 | 0.0 (0/26) | 0.0 | 0.0 (0/0) | 0.0 | 98.6 | 3 |
| 1677 | RF00122 GadY | 0.0 (0/28) | 0.0 | 0.0 (0/0) | 0.1 | 99.0 | 5 |
| 1678 | RF01277 snoU54 | 0.0 (0/3) | 0.0 | 0.0 (0/0) | 0.0 | 77.6 | 28 |
| 1679 | RF02184 ST7-OT3 2 | 0.0 (0/0) | 0.0 | 0.0 (0/0) | 0.0 | 91.6 | 5 |
| 1680 | RF01568 DdR18 | 0.0 (0/37) | 0.0 | 0.0 (0/0) | 1.2 | 80.0 | 3 |
| 1681 | RF02255 adapt33 1 | 0.0 (0/0) | 0.0 | 0.0 (0/0) | 0.0 | 77.3 | 3 |
| 1682 | RF00932 mir-471 | 0.0 (0/25) | 0.0 | 0.0 (0/0) | 0.0 | 97.4 | 2 |
| 1683 | RF00980 mir-643 | 0.0 (0/31) | 0.0 | 0.0 (0/0) | 0.0 | 90.7 | 2 |
| 1684 | RF00931 mir-879 | 0.0 (0/27) | 0.0 | 0.0 (0/0) | 0.0 | 96.0 | 2 |
| 1685 | RF02589 Spy779816 | 0.0 (0/26) | 0.0 | 0.0 (0/0) | 1.1 | 79.3 | 4 |
| 1686 | RF01427 snoR127 | 0.0 (0/4) | 0.0 | 0.0 (0/0) | 0.0 | 79.6 | 3 |
| 1687 | RF01562 DdR12 | 0.0 (0/0) | 0.0 | 0.0 (0/0) | 0.0 | 100.0 | 2 |
| 1688 | RF00044 Phage pRNA | 0.0 (0/43) | 0.0 | 0.0 (0/0) | 0.1 | 97.5 | 3 |
| 1689 | RF02241 Xoo2 | 0.0 (0/20) | 0.0 | 0.0 (0/0) | 0.6 | 92.2 | 3 |
| 1690 | RF02884 BcKCsr7 | 0.0 (0/35) | 0.0 | 0.0 (0/0) | 0.8 | 84.6 | 5 |
| 1691 | RF02656 sot0042 | 0.0 (0/53) | 0.0 | 0.0 (0/0) | 0.0 | 92.5 | 2 |
| 1692 | RF01128 sR43 | 0.0 (0/0) | 0.0 | 0.0 (0/0) | 0.0 | 86.0 | 6 |
| 1693 | RF01117 ciona-mir-92 | 0.0 (0/32) | 0.0 | 0.0 (0/0) | 0.2 | 75.3 | 2 |
| 1694 | RF01902 MIR439 | 0.0 (0/37) | 0.0 | 0.0 (0/0) | 0.5 | 95.8 | 5 |
| 1695 | RF02248 Six3os1 3 | 0.0 (0/0) | 0.0 | 0.0 (0/0) | 0.0 | 82.8 | 10 |
| 1696 | RF00606 SNORD93 | 0.0 (0/4) | 0.0 | 0.0 (0/0) | 0.0 | 88.7 | 17 |
| 1697 | RF01464 rliA | 0.0 (0/63) | 0.0 | 0.0 (0/0) | 0.2 | 94.6 | 4 |
| 1698 | RF00058 HgcF | 0.0 (0/38) | 0.0 | 0.0 (0/0) | 0.3 | 87.8 | 4 |
| 1699 | RF02667 PsiU2-38.40.42 | 0.0 (0/34) | 0.0 | 0.0 (0/0) | 0.2 | 86.0 | 3 |
| 1700 | RF01828 SprD | 0.0 (0/47) | 0.0 | 0.0 (0/0) | 0.0 | 100.0 | 2 |
| 1701 | RF01321 CRISPR-DR8 | 0.0 (0/7) | 0.0 | 0.0 (0/0) | 0.9 | 70.3 | 5 |
| 1702 | RF00843 mir-228 | 0.0 (0/35) | 0.0 | 0.0 (0/0) | 0.5 | 84.1 | 4 |
| 1703 | RF01175 snoU83D | 0.0 (0/2) | 0.0 | 0.0 (0/0) | 3.5 | 85.9 | 8 |
| 1704 | RF01979 HOTAIRM1 5 | 0.0 (0/0) | 0.0 | 0.0 (0/0) | 0.0 | 88.9 | 10 |
| 1705 | RF00834 mir-268 | 0.0 (0/28) | 0.0 | 0.0 (0/0) | 2.2 | 80.3 | 4 |
| 1706 | RF02216 ZNFX1-AS1 2 | 0.0 (0/0) | 0.0 | 0.0 (0/0) | 0.0 | 65.8 | 18 |
| 1707 | RF02050 STnc470 | 0.0 (0/43) | 0.0 | 0.0 (0/0) | 0.2 | 78.2 | 2 |
| 1708 | RF01468 rli32 | 0.0 (0/47) | 0.0 | 0.0 (0/0) | 0.7 | 93.9 | 5 |
| 1709 | RF00084 CsrC | 0.0 (0/59) | 0.0 | 0.0 (0/2) | 1.0 | 82.7 | 4 |
| 1710 | RF02148 MESTIT1 1 | 0.0 (0/0) | 0.0 | 0.0 (0/0) | 0.0 | 71.9 | 16 |
| 1711 | RF01336 CRISPR-DR23 | 0.0 (0/8) | 0.0 | 0.0 (0/0) | 0.0 | 92.6 | 3 |
| 1712 | RF01513 Afu 335 | 0.0 (0/0) | 0.0 | 0.0 (0/0) | 0.0 | 76.5 | 5 |
| 1713 | RF01662 ceN89 | 0.0 (0/4) | 0.0 | 0.0 (0/1) | 1.5 | 84.7 | 4 |
| 1714 | RF02627 Ssr1 | 0.0 (0/203) | 0.0 | 0.0 (0/0) | 0.0 | 100.0 | 2 |
| 1715 | RF01659 ceN86 | 0.0 (0/31) | 0.0 | 0.0 (0/0) | 0.1 | 77.9 | 2 |
| 1716 | RF00915 mir-760 | 0.0 (0/26) | 0.0 | 0.0 (0/0) | 1.3 | 89.7 | 7 |
| 1717 | RF02751 ES036 | 0.0 (0/8) | 0.0 | 0.0 (0/0) | 0.8 | 90.7 | 4 |
| 1718 | RF02726 sno ZL63 | 0.0 (0/11) | 0.0 | 0.0 (0/0) | 0.5 | 93.8 | 3 |
| 1719 | RF01596 plasmodium snoR20 | 0.0 (0/0) | 0.0 | 0.0 (0/0) | 0.0 | 86.2 | 2 |
| 1720 | RF01163 snoR64a | 0.0 (0/0) | 0.0 | 0.0 (0/0) | 0.0 | 76.2 | 4 |
| 1721 | RF00821 mir-249 | 0.0 (0/38) | 0.0 | 0.0 (0/0) | 0.8 | 70.9 | 4 |
| 1722 | RF01389 isrF | 0.0 (0/39) | 0.0 | 0.0 (0/0) | 0.8 | 85.4 | 3 |
| 1723 | RF01582 RUF4 | 0.0 (0/90) | 0.0 | 0.0 (0/0) | 0.0 | 96.0 | 2 |
| 1724 | RF01378 CRISPR-DR65 | 0.0 (0/7) | 0.0 | 0.0 (0/0) | 0.0 | 86.5 | 2 |
| 1725 | RF00196 AMV RNA1 SL | 0.0 (0/12) | 0.0 | 0.0 (0/0) | 0.8 | 95.8 | 6 |
| 1726 | RF01037 mir-644 | 0.0 (0/28) | 0.0 | 0.0 (0/0) | 0.1 | 92.5 | 3 |
| 1727 | RF00854 mir-5 | 0.0 (0/26) | 0.0 | 0.0 (0/0) | 0.2 | 91.2 | 6 |
| 1728 | RF01614 ceN110 | 0.0 (0/29) | 0.0 | 0.0 (0/0) | 0.4 | 87.2 | 3 |
| 1729 | RF02557 CbSR1 | 0.0 (0/29) | 0.0 | 0.0 (0/0) | 0.0 | 100.0 | 2 |
| 1730 | RF00336 snoJ26 | 0.0 (0/14) | 0.0 | 0.0 (0/0) | 0.1 | 95.1 | 5 |
| 1731 | RF02168 PVT1 5 | 0.0 (0/0) | 0.0 | 0.0 (0/0) | 0.0 | 69.2 | 23 |
| 1732 | RF00317 snoZ163 | 0.0 (0/5) | 0.0 | 0.0 (0/0) | 0.2 | 81.7 | 7 |
| 1733 | RF02099 rivX | 0.0 (0/66) | 0.0 | 0.0 (0/0) | 0.0 | 97.2 | 2 |
| 1734 | RF00366 mir-BHRF1-2 | 0.0 (0/25) | 0.0 | 0.0 (0/0) | 1.3 | 84.2 | 5 |
| 1735 | RF01484 rli59 | 0.0 (0/44) | 0.0 | 0.0 (0/0) | 0.2 | 96.2 | 5 |
| 1736 | RF01814 rhtB | 0.0 (0/13) | 0.0 | 0.0 (0/0) | 0.8 | 76.0 | 14 |
| 1737 | RF01275 sR22 | 0.0 (0/12) | 0.0 | 0.0 (0/0) | 1.2 | 92.0 | 6 |
| 1738 | RF02106 DLEU2 2 | 0.0 (0/0) | 0.0 | 0.0 (0/0) | 0.0 | 78.2 | 18 |
| 1739 | RF00060 HgcE | 0.0 (0/20) | 0.0 | 0.0 (0/0) | 0.8 | 79.0 | 4 |
| 1740 | RF01969 RMST 8 | 0.0 (0/0) | 0.0 | 0.0 (0/0) | 0.0 | 81.7 | 22 |
| 1741 | RF00511 IRES KSHV | 0.0 (0/61) | 0.0 | 0.0 (0/0) | 0.0 | 99.3 | 5 |
| 1742 | RF02607 AbsR25 | 0.0 (0/47) | 0.0 | 0.0 (0/0) | 0.0 | 99.4 | 2 |
| 1743 | RF00969 mir-556 | 0.0 (0/29) | 0.0 | 0.0 (0/0) | 0.1 | 83.2 | 2 |
| 1744 | RF02177 SMCR2 1 | 0.0 (0/0) | 0.0 | 0.0 (0/0) | 0.0 | 68.5 | 6 |
| 1745 | RF01345 CRISPR-DR35 | 0.0 (0/8) | 0.0 | 0.0 (0/0) | 0.1 | 97.2 | 2 |
| 1746 | RF02787 snoTBR4 | 0.0 (0/26) | 0.0 | 0.0 (0/2) | 1.2 | 77.3 | 3 |
| 1747 | RF02577 tsr24 | 0.0 (0/68) | 0.0 | 0.0 (0/0) | 0.1 | 98.2 | 4 |
| 1748 | RF02448 SpR20 sRNA | 0.0 (0/11) | 0.0 | 0.0 (0/0) | 0.7 | 89.8 | 7 |
| 1749 | RF00421 SNORA32 | 0.0 (0/23) | 0.0 | 0.0 (0/0) | 1.4 | 83.7 | 9 |
| 1750 | RF01088 TLS-PK5 | 0.0 (0/23) | 0.0 | 0.0 (0/0) | 0.7 | 80.6 | 3 |
| 1751 | RF01552 TB9Cs1H2 | 0.0 (0/8) | 0.0 | 0.0 (0/0) | 0.1 | 78.8 | 5 |
| 1752 | RF01034 mir-618 | 0.0 (0/36) | 0.0 | 0.0 (0/0) | 0.4 | 94.7 | 5 |
| 1753 | RF02600 BASRCI27 | 0.0 (0/57) | 0.0 | 0.0 (0/0) | 0.1 | 98.8 | 3 |

Continued on next page

| | RNA family (seed alignment) | Sensitivity annotated bpairs that covary % (cov_bps/bps) | Power average power % | Positive Predictive Value covarying pairs in structure % (cov_bps/cov_pairs) | average substitutions per bpair | avg pairwise identity % | number of sequences |
|---|---|---|---|---|---|---|---|
| 1754 | RF01885 HSR-omega 1 | 0.0 (0/0) | 0.0 | 0.0 (0/0) | 0.0 | 92.6 | 13 |
| 1755 | RF02598 EBv-sisRNA-2 | 0.0 (0/31) | 0.0 | 0.0 (0/0) | 0.1 | 94.5 | 2 |
| 1756 | RF01098 RF site9 | 0.0 (0/12) | 0.0 | 0.0 (0/0) | 0.1 | 91.8 | 2 |
| 1757 | RF01092 GP knot2 | 0.0 (0/14) | 0.0 | 0.0 (0/0) | 0.0 | 100.0 | 2 |
| 1758 | RF02305 TtnuCD31 | 0.0 (0/0) | 0.0 | 0.0 (0/0) | 0.0 | 97.0 | 2 |
| 1759 | RF02360 Yfr8 | 0.0 (0/63) | 0.0 | 0.0 (0/0) | 0.7 | 92.0 | 5 |
| 1760 | RF00780 MIR477 | 0.0 (0/29) | 0.0 | 0.0 (0/0) | 1.0 | 61.9 | 3 |
| 1761 | RF02498 Atu C3 | 0.0 (0/53) | 0.0 | 0.0 (0/0) | 0.0 | 92.9 | 2 |
| 1762 | RF01113 BMV3 UPD-PK3 | 0.0 (0/7) | 0.0 | 0.0 (0/0) | 0.0 | 95.7 | 2 |
| 1763 | RF00976 mir-583 | 0.0 (0/26) | 0.0 | 0.0 (0/0) | 0.2 | 96.4 | 3 |
| 1764 | RF01030 mir-422 | 0.0 (0/33) | 0.0 | 0.0 (0/0) | 1.0 | 78.0 | 3 |
| 1765 | RF02297 TtnuCD21 | 0.0 (0/0) | 0.0 | 0.0 (0/0) | 0.0 | 82.5 | 2 |
| 1766 | RF01900 mir-2024 | 0.0 (0/30) | 0.0 | 0.0 (0/0) | 0.3 | 88.8 | 7 |
| 1767 | RF01459 rliE | 0.0 (0/48) | 0.0 | 0.0 (0/1) | 0.5 | 82.7 | 4 |
| 1768 | RF02369 h2cR | 0.0 (0/51) | 0.0 | 0.0 (0/0) | 1.5 | 87.9 | 7 |
| 1769 | RF01487 rliI | 0.0 (0/81) | 0.0 | 0.0 (0/0) | 0.1 | 98.3 | 5 |
| 1770 | RF01679 P36 | 0.0 (0/21) | 0.0 | 0.0 (0/0) | 0.0 | 98.4 | 2 |
| 1771 | RF02777 OppA thermometer | 0.0 (0/82) | 0.0 | 0.0 (0/0) | 0.0 | 99.6 | 3 |
| 1772 | RF01107 SBRMV1 UPD-PKf | 0.0 (0/8) | 0.0 | 0.0 (0/0) | 0.0 | 100.0 | 2 |
| 1773 | RF02073 STnc260 | 0.0 (0/36) | 0.0 | 0.0 (0/0) | 0.1 | 92.6 | 2 |
| 1774 | RF02085 Yar 1 | 0.0 (0/0) | 0.0 | 0.0 (0/0) | 0.0 | 77.6 | 10 |
| 1775 | RF00988 mir-657 | 0.0 (0/19) | 0.0 | 0.0 (0/0) | 0.1 | 89.8 | 2 |
| 1776 | RF00817 mir-80 | 0.0 (0/34) | 0.0 | 0.0 (0/0) | 0.1 | 89.5 | 4 |
| 1777 | RF02138 HOXA11-AS1 2 | 0.0 (0/0) | 0.0 | 0.0 (0/0) | 0.0 | 83.0 | 21 |
| 1778 | RF02199 TTC28-AS1 2 | 0.0 (0/0) | 0.0 | 0.0 (0/0) | 0.0 | 71.4 | 8 |
| 1779 | RF01379 CRISPR-DR66 | 0.0 (0/7) | 0.0 | 0.0 (0/0) | 0.4 | 94.1 | 4 |
| 1780 | RF02901 AaHKsRNA41 | 0.0 (0/51) | 0.0 | 0.0 (0/0) | 0.4 | 46.4 | 2 |
| 1781 | RF02293 TtnuCD17 | 0.0 (0/0) | 0.0 | 0.0 (0/0) | 0.0 | 97.0 | 2 |
| 1782 | RF02258 adapt33 4 | 0.0 (0/0) | 0.0 | 0.0 (0/0) | 0.0 | 74.8 | 2 |
| 1783 | RF02712 EBER2 | 0.0 (0/47) | 0.0 | 0.0 (0/0) | 0.0 | 97.3 | 3 |
| 1784 | RF01216 snR87 | 0.0 (0/2) | 0.0 | 0.0 (0/0) | 0.5 | 89.9 | 4 |
| 1785 | RF02640 MOSES4 | 0.0 (0/46) | 0.0 | 0.0 (0/0) | 0.0 | 99.2 | 3 |
| 1786 | RF01091 PK-SPCSV | 0.0 (0/17) | 0.0 | 0.0 (0/0) | 0.0 | 100.0 | 2 |
| 1787 | RF02290 TtnuCD14 | 0.0 (0/0) | 0.0 | 0.0 (0/0) | 0.0 | 93.2 | 2 |
| 1788 | RF01222 sn2417 | 0.0 (0/2) | 0.0 | 0.0 (0/0) | 0.5 | 92.3 | 3 |
| 1789 | RF01308 sR58 | 0.0 (0/2) | 0.0 | 0.0 (0/0) | 0.5 | 87.1 | 5 |
| 1790 | RF01129 sR44 | 0.0 (0/0) | 0.0 | 0.0 (0/0) | 0.0 | 84.8 | 3 |
| 1791 | RF02218 ZNRD1-AS1 1 | 0.0 (0/0) | 0.0 | 0.0 (0/0) | 0.0 | 83.7 | 8 |
| 1792 | RF01632 ceN42 | 0.0 (0/35) | 0.0 | 0.0 (0/0) | 0.4 | 79.1 | 3 |
| 1793 | RF01359 CRISPR-DR46 | 0.0 (0/7) | 0.0 | 0.0 (0/0) | 0.0 | 84.2 | 2 |
| 1794 | RF02717 sno ncR4 | 0.0 (0/24) | 0.0 | 0.0 (0/0) | 0.3 | 88.6 | 3 |
| 1795 | RF02415 rliG | 0.0 (0/56) | 0.0 | 0.0 (0/0) | 1.3 | 72.8 | 5 |
| 1796 | RF02539 SNOR75 | 0.0 (0/4) | 0.0 | 0.0 (0/0) | 0.8 | 86.9 | 6 |
| 1797 | RF01001 mir-609 | 0.0 (0/34) | 0.0 | 0.0 (0/0) | 0.1 | 94.7 | 2 |
| 1798 | RF02795 Pab91 | 0.0 (0/19) | 0.0 | 0.0 (0/0) | 0.2 | 91.2 | 3 |
| 1799 | RF01409 STnc250 | 0.0 (0/25) | 0.0 | 0.0 (0/0) | 0.1 | 97.9 | 3 |
| 1800 | RF00603 SNORD23 | 0.0 (0/5) | 0.0 | 0.0 (0/0) | 1.6 | 83.8 | 15 |
| 1801 | RF02281 TtnuCD4 | 0.0 (0/0) | 0.0 | 0.0 (0/0) | 0.0 | 82.2 | 3 |
| 1802 | RF02087 Yar 3 | 0.0 (0/0) | 0.0 | 0.0 (0/0) | 0.0 | 87.8 | 4 |
| 1803 | RF02368 Yfr21 | 0.0 (0/43) | 0.0 | 0.0 (0/0) | 0.2 | 88.7 | 2 |
| 1804 | RF01109 SBRMV1 UPD-PKd | 0.0 (0/8) | 0.0 | 0.0 (0/0) | 0.0 | 100.0 | 2 |
| 1805 | RF02244 mir-785 | 0.0 (0/31) | 0.0 | 0.0 (0/0) | 0.9 | 70.8 | 4 |
| 1806 | RF02284 TtnuCD7 | 0.0 (0/0) | 0.0 | 0.0 (0/0) | 0.0 | 94.4 | 2 |
| 1807 | RF01028 mir-633 | 0.0 (0/19) | 0.0 | 0.0 (0/0) | 1.1 | 76.0 | 3 |
| 1808 | RF00922 mir-673 | 0.0 (0/26) | 0.0 | 0.0 (0/0) | 0.1 | 97.4 | 2 |
| 1809 | RF02639 EF0869 EF0870 | 0.0 (0/140) | 0.0 | 0.0 (0/0) | 0.0 | 99.5 | 4 |
| 1810 | RF02811 FHbp thermometer | 0.0 (0/18) | 0.0 | 0.0 (0/0) | 0.1 | 94.5 | 2 |
| 1811 | RF02283 TtnuCD6 | 0.0 (0/0) | 0.0 | 0.0 (0/0) | 0.0 | 78.6 | 2 |
| 1812 | RF02185 ST7-OT3 3 | 0.0 (0/0) | 0.0 | 0.0 (0/0) | 0.0 | 71.2 | 21 |
| 1813 | RF01467 rli36 | 0.0 (0/23) | 0.0 | 0.0 (0/0) | 0.3 | 95.2 | 6 |
| 1814 | RF01889 lincRNA-p21 1 | 0.0 (0/0) | 0.0 | 0.0 (0/0) | 0.0 | 88.1 | 2 |
| 1815 | RF02179 ST7-AS1 1 | 0.0 (0/0) | 0.0 | 0.0 (0/0) | 0.0 | 76.6 | 25 |
| 1816 | RF00212 SNORD38 | 0.0 (0/8) | 0.0 | 0.0 (0/1) | 1.2 | 80.8 | 7 |
| 1817 | RF01201 snR40 | 0.0 (0/0) | 0.0 | 0.0 (0/0) | 0.0 | 76.9 | 18 |
| 1818 | RF02642 Spy491311c | 0.0 (0/41) | 0.0 | 0.0 (0/0) | 0.0 | 100.0 | 2 |
| 1819 | RF01136 sR28 | 0.0 (0/0) | 0.0 | 0.0 (0/0) | 0.0 | 71.0 | 6 |
| 1820 | RF00981 mir-939 | 0.0 (0/31) | 0.0 | 0.0 (0/0) | 0.0 | 99.2 | 3 |
| 1821 | RF00874 mir-BART12 | 0.0 (0/31) | 0.0 | 0.0 (0/0) | 0.2 | 71.6 | 2 |
| 1822 | RF01981 PCGEM1 | 0.0 (0/0) | 0.0 | 0.0 (0/0) | 0.0 | 86.1 | 26 |
| 1823 | RF02110 DLEU2 6 | 0.0 (0/0) | 0.0 | 0.0 (0/0) | 0.0 | 71.5 | 32 |
| 1824 | RF01818 RsaC | 0.0 (0/116) | 0.0 | 0.0 (0/0) | 0.0 | 99.8 | 3 |
| 1825 | RF00182 Corona package | 0.0 (0/30) | 0.0 | 0.0 (0/0) | 1.0 | 78.3 | 3 |
| 1826 | RF00277 SNORD49 | 0.0 (0/0) | 0.0 | 0.0 (0/0) | 0.0 | 76.9 | 28 |
| 1827 | RF00192 BLV package | 0.0 (0/12) | 0.0 | 0.0 (0/0) | 0.1 | 96.4 | 5 |
| 1828 | RF02009 mir-987 | 0.0 (0/31) | 0.0 | 0.0 (0/0) | 0.0 | 100.0 | 3 |
| 1829 | RF01938 mir-1251 | 0.0 (0/22) | 0.0 | 0.0 (0/0) | 0.0 | 99.4 | 4 |
| 1830 | RF01606 plasmodium snoR31 | 0.0 (0/36) | 0.0 | 0.0 (0/0) | 0.4 | 80.9 | 3 |
| 1831 | RF02644 Spy490380c | 0.0 (0/23) | 0.0 | 0.0 (0/0) | 0.7 | 88.5 | 4 |
| 1832 | RF02257 adapt33 3 | 0.0 (0/0) | 0.0 | 0.0 (0/0) | 0.0 | 73.7 | 3 |
| 1833 | RF00363 mir-BART1 | 0.0 (0/24) | 0.0 | 0.0 (0/0) | 0.2 | 94.3 | 5 |
| 1834 | RF02068 STnc480 | 0.0 (0/8) | 0.0 | 0.0 (0/0) | 0.1 | 82.4 | 6 |
| 1835 | RF01332 CRISPR-DR19 | 0.0 (0/8) | 0.0 | 0.0 (0/0) | 0.2 | 77.6 | 4 |
| 1836 | RF00945 mir-1226 | 0.0 (0/28) | 0.0 | 0.0 (0/0) | 0.1 | 86.7 | 2 |
| 1837 | RF00301 snoZ256 | 0.0 (0/5) | 0.0 | 0.0 (0/0) | 1.4 | 83.2 | 5 |

Continued on next page

| | RNA family (seed alignment) | Sensitivity annotated bpairs that covary % (cov_bps/bps) | Power average power % | Positive Predictive Value covarying pairs in structure % (cov_bps/cov_pairs) | average substitutions per bpair | avg pairwise identity % | number of sequences |
|---|---|---|---|---|---|---|---|
| 1838 | RF02609 BSR0602 | 0.0 (0/53) | 0.0 | 0.0 (0/0) | 0.0 | 98.8 | 2 |
| 1839 | RF00721 MIR475 | 0.0 (0/41) | 0.0 | 0.0 (0/0) | 1.1 | 87.7 | 4 |
| 1840 | RF02501 Atu C7 | 0.0 (0/26) | 0.0 | 0.0 (0/0) | 0.0 | 90.5 | 3 |
| 1841 | RF01199 snR58 | 0.0 (0/0) | 0.0 | 0.0 (0/0) | 0.0 | 81.3 | 3 |
| 1842 | RF02392 sau-53 | 0.0 (0/56) | 0.0 | 0.0 (0/0) | 0.7 | 89.5 | 4 |
| 1843 | RF02606 AbsR28 | 0.0 (0/54) | 0.0 | 0.0 (0/0) | 0.1 | 98.3 | 2 |
| 1844 | RF02104 DLEU1 2 | 0.0 (0/0) | 0.0 | 0.0 (0/0) | 0.0 | 73.3 | 16 |
| 1845 | RF00814 mir-316 | 0.0 (0/35) | 0.0 | 0.0 (0/0) | 0.3 | 92.4 | 4 |
| 1846 | RF02124 JPX 1 | 0.0 (0/0) | 0.0 | 0.0 (0/0) | 0.0 | 71.1 | 14 |
| 1847 | RF01493 rli37 | 0.0 (0/57) | 0.0 | 0.0 (0/0) | 0.2 | 96.0 | 4 |
| 1848 | RF01462 rli26 | 0.0 (0/47) | 0.0 | 0.0 (0/0) | 0.4 | 94.1 | 5 |
| 1849 | RF01498 Afu 190 | 0.0 (0/0) | 0.0 | 0.0 (0/0) | 0.0 | 59.8 | 3 |
| 1850 | RF01119 sR32 | 0.0 (0/0) | 0.0 | 0.0 (0/0) | 0.0 | 88.9 | 3 |
| 1851 | RF02752 ES056 | 0.0 (0/28) | 0.0 | 0.0 (0/0) | 0.9 | 94.1 | 6 |
| 1852 | RF01465 rli31 | 0.0 (0/31) | 0.0 | 0.0 (0/0) | 0.8 | 89.0 | 3 |
| 1853 | RF01399 isrQ | 0.0 (0/50) | 0.0 | 0.0 (0/0) | 0.2 | 97.4 | 6 |
| 1854 | RF02904 AaHKsRNA69 | 0.0 (0/21) | 0.0 | 0.0 (0/0) | 0.7 | 71.6 | 3 |
| 1855 | RF02592 Spy392987 | 0.0 (0/21) | 0.0 | 0.0 (0/0) | 0.0 | 95.7 | 3 |
| 1856 | RF01822 RsaJ | 0.0 (0/79) | 0.0 | 0.0 (0/0) | 0.1 | 97.2 | 3 |
| 1857 | RF01105 SBWMV2 UPD-PKl | 0.0 (0/9) | 0.0 | 0.0 (0/0) | 0.0 | 100.0 | 2 |
| 1858 | RF02418 Spd-sr07 | 0.0 (0/16) | 0.0 | 0.0 (0/0) | 0.2 | 98.9 | 3 |
| 1859 | RF00926 MIR1151 | 0.0 (0/61) | 0.0 | 0.0 (0/0) | 0.1 | 88.2 | 2 |
| 1860 | RF01157 sn1185 | 0.0 (0/3) | 0.0 | 0.0 (0/0) | 0.0 | 92.1 | 5 |
| 1861 | RF01457 rli22 | 0.0 (0/23) | 0.0 | 0.0 (0/0) | 0.0 | 98.0 | 6 |
| 1862 | RF02172 RFPL3-AS1 2 | 0.0 (0/0) | 0.0 | 0.0 (0/0) | 0.0 | 81.7 | 3 |
| 1863 | RF01888 DLX6-AS1 2 | 0.0 (0/0) | 0.0 | 0.0 (0/0) | 0.0 | 87.9 | 19 |
| 1864 | RF01858 RsaF | 0.0 (0/38) | 0.0 | 0.0 (0/0) | 0.1 | 98.6 | 4 |
| 1865 | RF01587 snoR10 | 0.0 (0/4) | 0.0 | 0.0 (0/0) | 0.2 | 91.2 | 4 |
| 1866 | RF01992 VIS1 | 0.0 (0/0) | 0.0 | 0.0 (0/0) | 0.0 | 86.3 | 27 |
| 1867 | RF00741 mir-378 | 0.0 (0/24) | 0.0 | 0.0 (0/0) | 0.1 | 99.3 | 8 |
| 1868 | RF01347 CRISPR-DR37 | 0.0 (0/7) | 0.0 | 0.0 (0/0) | 0.1 | 78.1 | 3 |
| 1869 | RF02436 SpF43 sRNA | 0.0 (0/34) | 0.0 | 0.0 (0/0) | 1.1 | 85.5 | 5 |
| 1870 | RF02017 mir-1912 | 0.0 (0/26) | 0.0 | 0.0 (0/0) | 0.7 | 91.4 | 13 |
| 1871 | RF00847 mir-62 | 0.0 (0/22) | 0.0 | 0.0 (0/0) | 0.3 | 92.2 | 4 |
| 1872 | RF02123 FTX 4 | 0.0 (0/0) | 0.0 | 0.0 (0/0) | 0.0 | 79.4 | 13 |
| 1873 | RF02086 Yar 2 | 0.0 (0/0) | 0.0 | 0.0 (0/0) | 0.0 | 87.7 | 7 |
| 1874 | RF02788 snoTBR6 | 0.0 (0/13) | 0.0 | 0.0 (0/0) | 0.2 | 72.1 | 3 |
| 1875 | RF02628 Hrs1 | 0.0 (0/20) | 0.0 | 0.0 (0/0) | 0.1 | 86.5 | 2 |
| 1876 | RF01120 sR33 | 0.0 (0/0) | 0.0 | 0.0 (0/0) | 0.0 | 75.5 | 3 |
| 1877 | RF01211 SNORD124 | 0.0 (0/7) | 0.0 | 0.0 (0/0) | 0.3 | 80.8 | 18 |
| 1878 | RF02203 WT1-AS 1 | 0.0 (0/0) | 0.0 | 0.0 (0/0) | 0.0 | 86.6 | 10 |
| 1879 | RF01506 Afu 294 | 0.0 (0/0) | 0.0 | 0.0 (0/0) | 0.0 | 71.8 | 10 |
| 1880 | RF00953 mir-1497 | 0.0 (0/16) | 0.0 | 0.0 (0/0) | 0.0 | 82.2 | 2 |
| 1881 | RF01424 snoR118 | 0.0 (0/0) | 0.0 | 0.0 (0/0) | 0.0 | 75.0 | 8 |
| 1882 | RF02614 BSR1073 | 0.0 (0/67) | 0.0 | 0.0 (0/0) | 0.0 | 99.5 | 2 |
| 1883 | RF02736 Rev11 | 0.0 (0/111) | 0.0 | 0.0 (0/0) | 0.2 | 89.0 | 3 |
| 1884 | RF00327 snoZ194 | 0.0 (0/19) | 0.0 | 0.0 (0/0) | 0.0 | 98.8 | 5 |
| 1885 | RF01373 CRISPR-DR60 | 0.0 (0/3) | 0.0 | 0.0 (0/0) | 0.0 | 83.8 | 2 |
| 1886 | RF02416 TCV H4 | 0.0 (0/9) | 0.0 | 0.0 (0/0) | 0.1 | 84.0 | 2 |
| 1887 | RF02759 LasI thermometer | 0.0 (0/31) | 0.0 | 0.0 (0/0) | 0.0 | 97.0 | 3 |
| 1888 | RF00856 mir-232 | 0.0 (0/29) | 0.0 | 0.0 (0/0) | 0.5 | 77.9 | 3 |
| 1889 | RF00699 mir-134 | 0.0 (0/24) | 0.0 | 0.0 (0/0) | 1.1 | 87.8 | 6 |
| 1890 | RF02713 MCS4 | 0.0 (0/31) | 0.0 | 0.0 (0/0) | 0.1 | 92.6 | 5 |
| 1891 | RF02473 GlsR4 | 0.0 (0/0) | 0.0 | 0.0 (0/0) | 0.0 | 90.2 | 2 |
| 1892 | RF00208 snoR72 | 0.0 (0/9) | 0.0 | 0.0 (0/0) | 0.0 | 97.2 | 5 |
| 1893 | RF01904 HOTAIR 1 | 0.0 (0/0) | 0.0 | 0.0 (0/0) | 0.0 | 91.8 | 9 |
| 1894 | RF00367 mir-BHRF1-3 | 0.0 (0/22) | 0.0 | 0.0 (0/0) | 0.0 | 100.0 | 4 |
| 1895 | RF02852 Ysr206 | 0.0 (0/87) | 0.0 | 0.0 (0/0) | 0.2 | 72.7 | 2 |
| 1896 | RF01276 sR53 | 0.0 (0/0) | 0.0 | 0.0 (0/0) | 0.0 | 90.3 | 3 |
| 1897 | RF02497 Atu C10 | 0.0 (0/25) | 0.0 | 0.0 (0/0) | 0.0 | 98.9 | 2 |
| 1898 | RF00798 mir-49 | 0.0 (0/34) | 0.0 | 0.0 (0/0) | 0.7 | 76.2 | 4 |
| 1899 | RF02077 STnc220 | 0.0 (0/11) | 0.0 | 0.0 (0/0) | 0.3 | 90.9 | 4 |
| 1900 | RF01779 AS1726 | 0.0 (0/23) | 0.0 | 0.0 (0/0) | 0.0 | 100.0 | 2 |
| 1901 | RF01169 SNORD112 | 0.0 (0/6) | 0.0 | 0.0 (0/0) | 2.0 | 81.0 | 13 |
| 1902 | RF02754 ES205 | 0.0 (0/15) | 0.0 | 0.0 (0/0) | 0.1 | 95.7 | 5 |
| 1903 | RF02658 IRES RhPV | 0.0 (0/179) | 0.0 | 0.0 (0/0) | 0.0 | 98.3 | 2 |
| 1904 | RF01926 mir-981 | 0.0 (0/26) | 0.0 | 0.0 (0/0) | 1.0 | 57.7 | 3 |
| 1905 | RF02139 HOXA11-AS1 3 | 0.0 (0/0) | 0.0 | 0.0 (0/0) | 0.0 | 83.7 | 24 |
| 1906 | RF02267 Vax2os1 2 | 0.0 (0/0) | 0.0 | 0.0 (0/0) | 0.0 | 92.9 | 3 |
| 1907 | RF02324 TtnuHACA17 | 0.0 (0/27) | 0.0 | 0.0 (0/0) | 0.4 | 86.2 | 3 |
| 1908 | RF02871 ncS54 | 0.0 (0/22) | 0.0 | 0.0 (0/0) | 0.0 | 99.2 | 4 |
| 1909 | RF01954 SOX2OT exon4 | 0.0 (0/0) | 0.0 | 0.0 (0/0) | 0.0 | 81.4 | 10 |
| 1910 | RF01526 ceN106 | 0.0 (0/2) | 0.0 | 0.0 (0/0) | 0.0 | 93.5 | 3 |
| 1911 | RF02013 mir-1280 | 0.0 (0/33) | 0.0 | 0.0 (0/0) | 0.0 | 100.0 | 3 |
| 1912 | RF02393 sau-59 | 0.0 (0/27) | 0.0 | 0.0 (0/0) | 0.6 | 87.2 | 3 |
| 1913 | RF01185 snR75 | 0.0 (0/0) | 0.0 | 0.0 (0/0) | 0.0 | 68.9 | 61 |
| 1914 | RF00484 IRES Cx32 | 0.0 (0/32) | 0.0 | 0.0 (0/0) | 1.0 | 93.7 | 6 |
| 1915 | RF01137 sR21 | 0.0 (0/0) | 0.0 | 0.0 (0/0) | 0.0 | 53.0 | 2 |
| 1916 | RF02709 TeloSII ncR43 | 0.0 (0/70) | 0.0 | 0.0 (0/0) | 0.6 | 91.7 | 4 |
| 1917 | RF01892 TUG1 4 | 0.0 (0/0) | 0.0 | 0.0 (0/0) | 0.0 | 87.1 | 25 |
| 1918 | RF01038 mir-1307 | 0.0 (0/32) | 0.0 | 0.0 (0/0) | 0.2 | 96.4 | 7 |
| 1919 | RF00934 mir-463 | 0.0 (0/23) | 0.0 | 0.0 (0/0) | 0.2 | 87.7 | 2 |
| 1920 | RF01189 sn1502 | 0.0 (0/5) | 0.0 | 0.0 (0/0) | 1.0 | 94.4 | 4 |
| 1921 | RF02114 FAM13A-AS1 1 | 0.0 (0/0) | 0.0 | 0.0 (0/1) | 0.0 | 67.3 | 10 |

Continued on next page

| | RNA family (seed alignment) | Sensitivity annotated bpairs that covary % (cov_bps/bps) | Power average power % | Positive Predictive Value covarying pairs in structure % (cov_bps/cov_pairs) | average substitutions per bpair | avg pairwise identity % | number of sequences |
|---|---|---|---|---|---|---|---|
| 1922 | RF02689 hilD 3p UTR | 0.0 (0/86) | 0.0 | 0.0 (0/0) | 0.8 | 88.1 | 5 |
| 1923 | RF02214 mir-56 | 0.0 (0/0) | 0.0 | 0.0 (0/0) | 0.0 | 77.3 | 4 |
| 1924 | RF01469 rli33 | 0.0 (0/143) | 0.0 | 0.0 (0/0) | 0.1 | 94.8 | 3 |
| 1925 | RF01507 Afu 298 | 0.0 (0/0) | 0.0 | 0.0 (0/0) | 0.0 | 79.6 | 3 |
| 1926 | RF01197 snR39 | 0.0 (0/2) | 0.0 | 0.0 (0/0) | 1.0 | 86.0 | 4 |
| 1927 | RF01578 RUF1 | 0.0 (0/67) | 0.0 | 0.0 (0/0) | 0.1 | 98.6 | 3 |
| 1928 | RF02120 FTX 2 | 0.0 (0/0) | 0.0 | 0.0 (0/0) | 0.0 | 74.3 | 18 |
| 1929 | RF00509 snosnR64 | 0.0 (0/0) | 0.0 | 0.0 (0/1) | 0.0 | 81.4 | 10 |
| 1930 | RF02398 sau-6072 | 0.0 (0/17) | 0.0 | 0.0 (0/0) | 0.0 | 92.5 | 2 |
| 1931 | RF02224 sX5 | 0.0 (0/10) | 0.0 | 0.0 (0/0) | 0.9 | 94.8 | 10 |
| 1932 | RF01945 mir-1388 | 0.0 (0/28) | 0.0 | 0.0 (0/0) | 1.2 | 70.5 | 4 |
| 1933 | RF02020 mir-25 | 0.0 (0/27) | 0.0 | 0.0 (0/0) | 0.2 | 95.2 | 3 |
| 1934 | RF02747 FtrA | 0.0 (0/22) | 0.0 | 0.0 (0/0) | 0.2 | 97.3 | 5 |
| 1935 | RF02133 HOXB13-AS1 2 | 0.0 (0/0) | 0.0 | 0.0 (0/0) | 0.0 | 71.9 | 14 |
| 1936 | RF00905 mir-789 | 0.0 (0/36) | 0.0 | 0.0 (0/0) | 0.1 | 91.8 | 2 |
| 1937 | RF02847 Ref64 | 0.0 (0/23) | 0.0 | 0.0 (0/0) | 0.8 | 85.5 | 3 |
| 1938 | RF02560 CbSR9 | 0.0 (0/48) | 0.0 | 0.0 (0/0) | 0.0 | 99.5 | 2 |
| 1939 | RF02192 TCL6 2 | 0.0 (0/0) | 0.0 | 0.0 (0/0) | 0.0 | 69.8 | 12 |
| 1940 | RF02826 Scr6925 | 0.0 (0/43) | 0.0 | 0.0 (0/0) | 0.9 | 82.6 | 4 |
| 1941 | RF02513 PYLIS 6 | 0.0 (0/11) | 0.0 | 0.0 (0/0) | 0.0 | 100.0 | 3 |
| 1942 | RF00323 snoR79 | 0.0 (0/5) | 0.0 | 0.0 (0/0) | 0.8 | 99.4 | 5 |
| 1943 | RF00973 mir-597 | 0.0 (0/32) | 0.0 | 0.0 (0/0) | 0.9 | 81.2 | 4 |
| 1944 | RF00986 mir-920 | 0.0 (0/23) | 0.0 | 0.0 (0/0) | 1.2 | 77.1 | 4 |
| 1945 | RF02140 HOXA11-AS1 4 | 0.0 (0/0) | 0.0 | 0.0 (0/0) | 0.0 | 86.7 | 20 |
| 1946 | RF01622 ceN22 | 0.0 (0/4) | 0.0 | 0.0 (0/0) | 0.2 | 94.7 | 2 |
| 1947 | RF00963 mir-642 | 0.0 (0/36) | 0.0 | 0.0 (0/0) | 0.0 | 91.8 | 2 |
| 1948 | RF00193 CTV rep sig | 0.0 (0/73) | 0.0 | 0.0 (0/0) | 0.4 | 98.1 | 9 |
| 1949 | RF02426 SpF03 sRNA | 0.0 (0/31) | 0.0 | 0.0 (0/0) | 0.4 | 93.2 | 3 |
| 1950 | RF02625 WsnRNA46 | 0.0 (0/36) | 0.0 | 0.0 (0/0) | 0.4 | 73.8 | 3 |
| 1951 | RF01883 TUG1 2 | 0.0 (0/0) | 0.0 | 0.0 (0/0) | 0.0 | 88.8 | 17 |
| 1952 | RF02197 TP73-AS1 | 0.0 (0/0) | 0.0 | 0.0 (0/0) | 0.0 | 71.7 | 4 |
| 1953 | RF00977 mir-600 | 0.0 (0/25) | 0.0 | 0.0 (0/0) | 0.9 | 83.5 | 3 |
| 1954 | RF02789 snoTBR12 | 0.0 (0/32) | 0.0 | 0.0 (0/0) | 0.9 | 73.4 | 3 |
| 1955 | RF01574 DdR6 | 0.0 (0/0) | 0.0 | 0.0 (0/0) | 0.0 | 91.4 | 3 |
| 1956 | RF01516 v-snoRNA-1 | 0.0 (0/0) | 0.0 | 0.0 (0/0) | 0.0 | 83.1 | 2 |
| 1957 | RF01324 CRISPR-DR11 | 0.0 (0/7) | 0.0 | 0.0 (0/0) | 1.4 | 91.1 | 5 |
| 1958 | RF02288 TtnuCD11 | 0.0 (0/0) | 0.0 | 0.0 (0/0) | 0.0 | 76.1 | 3 |
| 1959 | RF00894 mir-790 | 0.0 (0/23) | 0.0 | 0.0 (0/0) | 0.1 | 85.1 | 2 |
| 1960 | RF02590 Spy1186876 | 0.0 (0/59) | 0.0 | 0.0 (0/0) | 0.2 | 95.3 | 3 |
| 1961 | RF02864 sRNA 0030 | 0.0 (0/58) | 0.0 | 0.0 (0/0) | 0.3 | 97.6 | 3 |
| 1962 | RF01388 isrD | 0.0 (0/13) | 0.0 | 0.0 (0/0) | 0.0 | 98.0 | 2 |
| 1963 | RF01022 mir-611 | 0.0 (0/17) | 0.0 | 0.0 (0/0) | 2.0 | 75.1 | 8 |
| 1964 | RF02119 FTX 1 | 0.0 (0/0) | 0.0 | 0.0 (0/0) | 0.0 | 79.3 | 11 |
| 1965 | RF00707 mir-197 | 0.0 (0/31) | 0.0 | 0.0 (0/1) | 1.0 | 85.0 | 5 |
| 1966 | RF02303 TtnuCD28 | 0.0 (0/0) | 0.0 | 0.0 (0/0) | 0.0 | 95.2 | 2 |
| 1967 | RF00867 mir-BART5 | 0.0 (0/29) | 0.0 | 0.0 (0/0) | 0.1 | 78.7 | 2 |
| 1968 | RF01575 DdR7 | 0.0 (0/0) | 0.0 | 0.0 (0/0) | 0.0 | 85.0 | 3 |
| 1969 | RF01603 snoR29 | 0.0 (0/4) | 0.0 | 0.0 (0/0) | 0.0 | 84.7 | 3 |
| 1970 | RF01510 MFR | 0.0 (0/20) | 0.0 | 0.0 (0/0) | 0.8 | 74.1 | 3 |
| 1971 | RF01429 snoR130 | 0.0 (0/3) | 0.0 | 0.0 (0/0) | 0.0 | 85.7 | 2 |
| 1972 | RF02191 TCL6 1 | 0.0 (0/0) | 0.0 | 0.0 (0/0) | 0.0 | 77.8 | 9 |
| 1973 | RF01027 mir-765 | 0.0 (0/37) | 0.0 | 0.0 (0/0) | 0.1 | 92.1 | 2 |
| 1974 | RF01250 snR189 | 0.0 (0/44) | 0.0 | 0.0 (0/1) | 0.3 | 88.7 | 4 |
| 1975 | RF02812 asponA | 0.0 (0/88) | 0.0 | 0.0 (0/0) | 0.6 | 82.2 | 3 |
| 1976 | RF01647 ceN61 | 0.0 (0/3) | 0.0 | 0.0 (0/0) | 0.0 | 93.1 | 4 |
| 1977 | RF01860 Afu 455 | 0.0 (0/0) | 0.0 | 0.0 (0/0) | 0.0 | 78.1 | 6 |
| 1978 | RF01082 SBWMV1 UPD-PKh | 0.0 (0/10) | 0.0 | 0.0 (0/0) | 0.2 | 80.0 | 2 |
| 1979 | RF01563 DdR13 | 0.0 (0/0) | 0.0 | 0.0 (0/0) | 0.0 | 81.5 | 3 |
| 1980 | RF01309 sR60 | 0.0 (0/2) | 0.0 | 0.0 (0/0) | 0.0 | 87.6 | 4 |
| 1981 | RF01913 mir-2778 | 0.0 (0/29) | 0.0 | 0.0 (0/0) | 0.4 | 93.8 | 7 |
| 1982 | RF00290 BaMV CRE | 0.0 (0/30) | 0.0 | 0.0 (0/0) | 0.1 | 99.1 | 5 |
| 1983 | RF01505 Afu 264 | 0.0 (0/0) | 0.0 | 0.0 (0/0) | 0.0 | 69.2 | 5 |
| 1984 | RF01346 CRISPR-DR36 | 0.0 (0/6) | 0.0 | 0.0 (0/0) | 0.2 | 94.4 | 3 |
| 1985 | RF01085 TLS-PK4 | 0.0 (0/38) | 0.0 | 0.0 (0/0) | 0.1 | 89.7 | 2 |
| 1986 | RF02201 TTC28-AS1 4 | 0.0 (0/0) | 0.0 | 0.0 (0/0) | 0.0 | 71.9 | 18 |
| 1987 | RF01232 snoR442 | 0.0 (0/8) | 0.0 | 0.0 (0/0) | 1.6 | 88.2 | 10 |
| 1988 | RF02441 SpF61 sRNA | 0.0 (0/12) | 0.0 | 0.0 (0/0) | 0.2 | 88.5 | 3 |
| 1989 | RF02389 sau-31 | 0.0 (0/11) | 0.0 | 0.0 (0/0) | 0.2 | 97.0 | 3 |
| 1990 | RF02097 mir-1662 | 0.0 (0/22) | 0.0 | 0.0 (0/0) | 0.0 | 86.4 | 2 |
| 1991 | RF01009 mir-M7 | 0.0 (0/23) | 0.0 | 0.0 (0/0) | 0.3 | 74.0 | 2 |
| 1992 | RF02674 AsdA | 0.0 (0/23) | 0.0 | 0.0 (0/0) | 0.5 | 94.0 | 4 |
| 1993 | RF02395 sau-66 | 0.0 (0/20) | 0.0 | 0.0 (0/0) | 0.1 | 97.5 | 3 |
| 1994 | RF01026 MIR828 | 0.0 (0/19) | 0.0 | 0.0 (0/0) | 0.8 | 70.8 | 4 |
| 1995 | RF01205 snR62 | 0.0 (0/3) | 0.0 | 0.0 (0/0) | 0.0 | 95.0 | 2 |
| 1996 | RF00904 mir-392 | 0.0 (0/26) | 0.0 | 0.0 (0/0) | 0.6 | 70.0 | 3 |
| 1997 | RF01206 snoR109 | 0.0 (0/19) | 0.0 | 0.0 (0/0) | 1.3 | 77.6 | 6 |
| 1998 | RF01837 toga FSE | 0.0 (0/11) | 0.0 | 0.0 (0/0) | 0.1 | 95.4 | 3 |
| 1999 | RF02472 Gl RNase MRP | 0.0 (0/22) | 0.0 | 0.0 (0/0) | 0.1 | 90.8 | 2 |
| 2000 | RF01646 ceN63 | 0.0 (0/3) | 0.0 | 0.0 (0/0) | 0.3 | 97.0 | 3 |
| 2001 | RF00107 FinP | 0.0 (0/24) | 0.0 | 0.0 (0/0) | 0.2 | 90.7 | 6 |
| 2002 | RF01968 RMST 7 | 0.0 (0/0) | 0.0 | 0.0 (0/0) | 0.0 | 84.8 | 22 |
| 2003 | RF02189 ST7-OT4 3 | 0.0 (0/0) | 0.0 | 0.0 (0/0) | 0.0 | 73.6 | 20 |
| 2004 | RF02888 BtsR1 | 0.0 (0/9) | 0.0 | 0.0 (0/0) | 0.9 | 89.5 | 3 |
| 2005 | RF01344 CRISPR-DR34 | 0.0 (0/8) | 0.0 | 0.0 (0/0) | 1.5 | 89.2 | 5 |

Continued on next page

| | RNA family (seed alignment) | Sensitivity annotated bpairs that covary % (cov_bps/bps) | Power average power % | Positive Predictive Value covarying pairs in structure % (cov_bps/cov_pairs) | average substitutions per bpair | avg pairwise identity % | number of sequences |
|---|---|---|---|---|---|---|---|
| 2006 | RF02428 SpF11 sRNA | 0.0 (0/20) | 0.0 | 0.0 (0/0) | 0.2 | 90.9 | 3 |
| 2007 | RF02237 asX3 | 0.0 (0/50) | 0.0 | 0.0 (0/0) | 0.9 | 90.7 | 4 |
| 2008 | RF01306 sR52 | 0.0 (0/4) | 0.0 | 0.0 (0/0) | 0.0 | 86.2 | 2 |
| 2009 | RF02740 Fwd6 3p UTR | 0.0 (0/28) | 0.0 | 0.0 (0/0) | 0.1 | 97.7 | 3 |
| 2010 | RF01188 snR56 | 0.0 (0/0) | 0.0 | 0.0 (0/0) | 0.0 | 70.5 | 16 |
| 2011 | RF00835 mir-58 | 0.0 (0/25) | 0.0 | 0.0 (0/1) | 0.0 | 87.6 | 4 |
| 2012 | RF00907 mir-941 | 0.0 (0/33) | 0.0 | 0.0 (0/0) | 0.1 | 94.4 | 2 |
| 2013 | RF00920 MIR444 | 0.0 (0/48) | 0.0 | 0.0 (0/1) | 0.9 | 74.8 | 4 |
| 2014 | RF01644 ceN59 | 0.0 (0/26) | 0.0 | 0.0 (0/2) | 0.2 | 72.1 | 3 |
| 2015 | RF02892 SR6 antitoxin | 0.0 (0/30) | 0.0 | 0.0 (0/0) | 0.4 | 88.4 | 3 |
| 2016 | RF01871 MALAT1 | 0.0 (0/0) | 0.0 | 0.0 (0/0) | 0.0 | 84.2 | 17 |
| 2017 | RF00836 mir-250 | 0.0 (0/27) | 0.0 | 0.0 (0/0) | 0.4 | 82.4 | 4 |
| 2018 | RF02101 HULC | 0.0 (0/0) | 0.0 | 0.0 (0/0) | 0.0 | 66.9 | 19 |
| 2019 | RF01921 mir-1296 | 0.0 (0/32) | 0.0 | 0.0 (0/0) | 0.1 | 96.7 | 6 |
| 2020 | RF00702 mir-182 | 0.0 (0/20) | 0.0 | 0.0 (0/0) | 0.2 | 85.0 | 17 |
| 2021 | RF01984 ZEB2 AS1 1 | 0.0 (0/0) | 0.0 | 0.0 (0/0) | 0.0 | 88.4 | 11 |
| 2022 | RF01337 CRISPR-DR24 | 0.0 (0/5) | 0.0 | 0.0 (0/0) | 0.4 | 85.6 | 4 |
| 2023 | RF01515 Afu 514 | 0.0 (0/0) | 0.0 | 0.0 (0/0) | 0.0 | 72.0 | 5 |
| 2024 | RF01617 ceN113 | 0.0 (0/3) | 0.0 | 0.0 (0/0) | 0.0 | 81.9 | 2 |
| 2025 | RF00918 mir-872 | 0.0 (0/25) | 0.0 | 0.0 (0/0) | 0.6 | 85.7 | 6 |
| 2026 | RF02083 OrzO-P | 0.0 (0/17) | 0.0 | 0.0 (0/0) | 1.0 | 85.5 | 7 |
| 2027 | RF02650 CncR1 | 0.0 (0/62) | 0.0 | 0.0 (0/0) | 0.0 | 98.4 | 3 |
| 2028 | RF01341 CRISPR-DR30 | 0.0 (0/6) | 0.0 | 0.0 (0/0) | 0.0 | 80.7 | 3 |
| 2029 | RF01433 snoR137 | 0.0 (0/31) | 0.0 | 0.0 (0/1) | 0.5 | 76.7 | 4 |
| 2030 | RF02023 mir-1208 | 0.0 (0/24) | 0.0 | 0.0 (0/0) | 0.0 | 99.1 | 3 |
| 2031 | RF01437 S pombe snR10 | 0.0 (0/34) | 0.0 | 0.0 (0/0) | 0.0 | 100.0 | 2 |
| 2032 | RF02570 BM-sr0117 | 0.0 (0/25) | 0.0 | 0.0 (0/0) | 0.1 | 98.4 | 3 |
| 2033 | RF02486 GlsR23 | 0.0 (0/17) | 0.0 | 0.0 (0/0) | 0.1 | 83.0 | 2 |
| 2034 | RF02734 Cgb105 | 0.0 (0/43) | 0.0 | 0.0 (0/0) | 0.0 | 97.0 | 2 |
| 2035 | RF02090 DAOA-AS1 1 | 0.0 (0/0) | 0.0 | 0.0 (0/0) | 0.0 | 76.2 | 24 |
| 2036 | RF02699 Avashort thermometer | 0.0 (0/13) | 0.0 | 0.0 (0/0) | 0.0 | 91.4 | 2 |
| 2037 | RF01553 TB9Cs1H3 | 0.0 (0/11) | 0.0 | 0.0 (0/0) | 0.7 | 73.8 | 3 |
| 2038 | RF01906 HOTAIR 3 | 0.0 (0/0) | 0.0 | 0.0 (0/0) | 0.0 | 87.0 | 17 |
| 2039 | RF01876 MIAT exon5 2 | 0.0 (0/0) | 0.0 | 0.0 (0/0) | 0.0 | 89.1 | 10 |
| 2040 | RF02659 ncRv12659 | 0.0 (0/50) | 0.0 | 0.0 (0/0) | 0.0 | 99.4 | 2 |
| 2041 | RF00734 mir-52 | 0.0 (0/34) | 0.0 | 0.0 (0/0) | 0.8 | 80.1 | 5 |
| 2042 | RF01370 CRISPR-DR57 | 0.0 (0/7) | 0.0 | 0.0 (0/0) | 0.0 | 80.2 | 3 |
| 2043 | RF02586 HCV package-SL733 | 0.0 (0/7) | 0.0 | 0.0 (0/0) | 0.0 | 94.9 | 4 |
| 2044 | RF02730 JA02 | 0.0 (0/35) | 0.0 | 0.0 (0/0) | 0.4 | 89.9 | 4 |
| 2045 | RF02841 Ref70 | 0.0 (0/29) | 0.0 | 0.0 (0/0) | 0.9 | 80.9 | 3 |
| 2046 | RF02341 ncrMT1302 | 0.0 (0/20) | 0.0 | 0.0 (0/0) | 1.4 | 78.1 | 7 |
| 2047 | RF01297 sR40 | 0.0 (0/12) | 0.0 | 0.0 (0/0) | 0.0 | 98.4 | 2 |
| 2048 | RF00853 mir-304 | 0.0 (0/33) | 0.0 | 0.0 (0/0) | 0.4 | 91.7 | 5 |
| 2049 | RF01874 MIAT exon1 | 0.0 (0/0) | 0.0 | 0.0 (0/0) | 0.0 | 87.5 | 10 |
| 2050 | RF02295 TtnuCD19 | 0.0 (0/0) | 0.0 | 0.0 (0/0) | 0.0 | 83.8 | 2 |
| 2051 | RF01360 CRISPR-DR47 | 0.0 (0/4) | 0.0 | 0.0 (0/0) | 0.0 | 97.3 | 2 |
| 2052 | RF02195 TP53TG1 1 | 0.0 (0/0) | 0.0 | 0.0 (0/0) | 0.0 | 69.3 | 16 |
| 2053 | RF02117 FMR1-AS1 1 | 0.0 (0/0) | 0.0 | 0.0 (0/0) | 0.0 | 86.1 | 20 |
| 2054 | RF02641 Spy490483c | 0.0 (0/27) | 0.0 | 0.0 (0/0) | 0.1 | 98.6 | 3 |
| 2055 | RF01676 P31 | 0.0 (0/18) | 0.0 | 0.0 (0/0) | 1.1 | 87.2 | 4 |
| 2056 | RF01604 plasmodium snoR28 | 0.0 (0/4) | 0.0 | 0.0 (0/0) | 0.0 | 85.5 | 3 |
| 2057 | RF01550 TB8Cs3H1 | 0.0 (0/16) | 0.0 | 0.0 (0/0) | 1.8 | 78.3 | 7 |
| 2058 | RF01151 snoU82P | 0.0 (0/0) | 0.0 | 0.0 (0/0) | 0.0 | 86.3 | 14 |
| 2059 | RF01898 mir-363 | 0.0 (0/26) | 0.0 | 0.0 (0/0) | 0.4 | 95.9 | 7 |
| 2060 | RF01955 NEAT1 1 | 0.0 (0/0) | 0.0 | 0.0 (0/0) | 0.0 | 83.7 | 17 |
| 2061 | RF01605 plasmodium snoR30 | 0.0 (0/3) | 0.0 | 0.0 (0/0) | 0.0 | 86.5 | 3 |
| 2062 | RF01152 sR1 | 0.0 (0/0) | 0.0 | 0.0 (0/1) | 0.0 | 64.5 | 12 |
| 2063 | RF02135 HTT-AS1 2 | 0.0 (0/0) | 0.0 | 0.0 (0/0) | 0.0 | 90.4 | 3 |
| 2064 | RF00364 mir-BART2 | 0.0 (0/21) | 0.0 | 0.0 (0/0) | 1.4 | 92.9 | 8 |
| 2065 | RF01131 sR47 | 0.0 (0/0) | 0.0 | 0.0 (0/0) | 0.0 | 72.1 | 8 |
| 2066 | RF02163 sR-tMet | 0.0 (0/17) | 0.0 | 0.0 (0/0) | 0.1 | 81.4 | 2 |
| 2067 | RF00483 IRES IGF2 | 0.0 (0/28) | 0.0 | 0.0 (0/0) | 0.6 | 94.0 | 9 |
| 2068 | RF01138 sR23 | 0.0 (0/0) | 0.0 | 0.0 (0/0) | 0.0 | 78.9 | 6 |
| 2069 | RF02770 Ysr224 | 0.0 (0/28) | 0.0 | 0.0 (0/0) | 0.7 | 84.3 | 5 |
| 2070 | RF02296 TtnuCD20 | 0.0 (0/0) | 0.0 | 0.0 (0/0) | 0.0 | 97.0 | 2 |
| 2071 | RF01520 CC0734 | 0.0 (0/17) | 0.0 | 0.0 (0/0) | 0.1 | 83.2 | 2 |
| 2072 | RF02250 Six3os1 5 | 0.0 (0/0) | 0.0 | 0.0 (0/0) | 0.0 | 83.7 | 15 |
| 2073 | RF01130 sR46 | 0.0 (0/5) | 0.0 | 0.0 (0/0) | 1.0 | 83.4 | 7 |
| 2074 | RF02671 Ysr35 | 0.0 (0/95) | 0.0 | 0.0 (0/0) | 0.0 | 99.2 | 3 |
| 2075 | RF01407 STnc560 | 0.0 (0/39) | 0.0 | 0.0 (0/0) | 1.1 | 95.4 | 12 |
| 2076 | RF00921 mir-665 | 0.0 (0/27) | 0.0 | 0.0 (0/0) | 0.7 | 86.8 | 6 |
| 2077 | RF02507 Atu Ti3 | 0.0 (0/7) | 0.0 | 0.0 (0/0) | 1.0 | 78.0 | 4 |
| 2078 | RF02024 mir-1180 | 0.0 (0/27) | 0.0 | 0.0 (0/0) | 0.4 | 89.4 | 3 |
| 2079 | RF02335 GlsR2 miR4 | 0.0 (0/32) | 0.0 | 0.0 (0/0) | 0.1 | 99.3 | 3 |
| 2080 | RF01353 CRISPR-DR44 | 0.0 (0/4) | 0.0 | 0.0 (0/0) | 0.0 | 83.3 | 3 |
| 2081 | RF02867 ncS011 | 0.0 (0/62) | 0.0 | 0.0 (0/0) | 0.4 | 89.0 | 4 |
| 2082 | RF00715 mir-383 | 0.0 (0/25) | 0.0 | 0.0 (0/0) | 0.5 | 91.1 | 6 |
| 2083 | RF00372 sroH | 0.0 (0/6) | 0.0 | 0.0 (0/0) | 0.0 | 76.5 | 2 |
| 2084 | RF01641 ceN53 | 0.0 (0/4) | 0.0 | 0.0 (0/0) | 0.5 | 90.4 | 3 |
| 2085 | RF01623 ceN23-1 | 0.0 (0/25) | 0.0 | 0.0 (0/0) | 0.0 | 93.7 | 3 |
| 2086 | RF00887 mir-802 | 0.0 (0/33) | 0.0 | 0.0 (0/0) | 1.3 | 87.3 | 13 |
| 2087 | RF00601 SCARNA20 | 0.0 (0/33) | 0.0 | 0.0 (0/0) | 1.0 | 79.0 | 5 |
| 2088 | RF01905 HOTAIR 2 | 0.0 (0/0) | 0.0 | 0.0 (0/0) | 0.0 | 88.7 | 11 |
| 2089 | RF01576 DdR8 | 0.0 (0/0) | 0.0 | 0.0 (0/0) | 0.0 | 100.0 | 2 |

Continued on next page

| | RNA family (seed alignment) | Sensitivity annotated bpairs that covary % (cov_bps/bps) | Power average power % | Positive Predictive Value covarying pairs in structure % (cov_bps/cov_pairs) | average substitutions per bpair | avg pairwise identity % | number of sequences |
|---|---|---|---|---|---|---|---|
| 2090 | RF01985 ZEB2 AS1 2 | 0.0 (0/0) | 0.0 | 0.0 (0/0) | 0.0 | 88.2 | 11 |
| 2091 | RF00979 mir-553 | 0.0 (0/19) | 0.0 | 0.0 (0/0) | 1.1 | 79.8 | 3 |
| 2092 | RF02213 ZFAT-AS1 3 | 0.0 (0/0) | 0.0 | 0.0 (0/0) | 0.0 | 83.9 | 5 |
| 2093 | RF01474 rli42 | 0.0 (0/58) | 0.0 | 0.0 (0/0) | 0.2 | 95.0 | 4 |
| 2094 | RF01589 plasmodium snoR11 | 0.0 (0/53) | 0.0 | 0.0 (0/0) | 0.1 | 96.5 | 2 |
| 2095 | RF00744 mir-361 | 0.0 (0/24) | 0.0 | 0.0 (0/0) | 0.2 | 93.3 | 5 |
| 2096 | RF01078 PK-PYVV | 0.0 (0/15) | 0.0 | 0.0 (0/0) | 0.3 | 84.1 | 3 |
| 2097 | RF00806 mir-350 | 0.0 (0/29) | 0.0 | 0.0 (0/0) | 1.6 | 87.9 | 11 |
| 2098 | RF01821 RsaH | 0.0 (0/29) | 0.0 | 0.0 (0/0) | 1.8 | 70.9 | 5 |
| 2099 | RF02386 sau-19 | 0.0 (0/27) | 0.0 | 0.0 (0/0) | 0.2 | 92.4 | 4 |
| 2100 | RF01861 TB9Cs1H1 | 0.0 (0/11) | 0.0 | 0.0 (0/0) | 1.0 | 70.2 | 4 |
| 2101 | RF02207 WT1-AS 5 | 0.0 (0/0) | 0.0 | 0.0 (0/0) | 0.0 | 98.2 | 4 |
| 2102 | RF02602 BASRCI414 | 0.0 (0/74) | 0.0 | 0.0 (0/0) | 0.2 | 96.5 | 3 |
| 2103 | RF00781 mir-505 | 0.0 (0/32) | 0.0 | 0.0 (0/0) | 0.3 | 93.5 | 5 |
| 2104 | RF02274 AniS | 0.0 (0/22) | 0.0 | 0.0 (0/0) | 0.3 | 95.5 | 5 |
| 2105 | RF01032 mir-554 | 0.0 (0/18) | 0.0 | 0.0 (0/0) | 1.1 | 79.4 | 4 |
| 2106 | RF02801 PyrR201 | 0.0 (0/21) | 0.0 | 0.0 (0/0) | 0.5 | 86.5 | 5 |
| 2107 | RF02677 NsiR4 | 0.0 (0/12) | 0.0 | 0.0 (0/0) | 0.8 | 86.6 | 4 |
| 2108 | RF01887 DLX6-AS1 1 | 0.0 (0/0) | 0.0 | 0.0 (0/0) | 0.0 | 90.8 | 15 |
| 2109 | RF02647 Cis52 sRNA | 0.0 (0/57) | 0.0 | 0.0 (0/0) | 0.1 | 88.3 | 2 |
| 2110 | RF02339 GlsR17 miR2 | 0.0 (0/21) | 0.0 | 0.0 (0/0) | 0.6 | 93.4 | 4 |
| 2111 | RF02326 TtnuHACA19 | 0.0 (0/27) | 0.0 | 0.0 (0/0) | 0.0 | 95.9 | 2 |
| 2112 | RF00179 GAIT | 0.0 (0/18) | 0.0 | 0.0 (0/0) | 0.7 | 98.8 | 8 |
| 2113 | RF02226 sX7 | 0.0 (0/26) | 0.0 | 0.0 (0/0) | 0.5 | 95.9 | 8 |
| 2114 | RF02042 HOTTIP 3 | 0.0 (0/0) | 0.0 | 0.0 (0/0) | 0.0 | 80.0 | 23 |
| 2115 | RF01677 P35 | 0.0 (0/23) | 0.0 | 0.0 (0/0) | 0.0 | 98.4 | 2 |
| 2116 | RF02578 tsr25 | 0.0 (0/53) | 0.0 | 0.0 (0/0) | 0.0 | 99.6 | 3 |
| 2117 | RF01650 ceN68 | 0.0 (0/29) | 0.0 | 0.0 (0/0) | 0.0 | 100.0 | 2 |
| 2118 | RF01556 TB9Cs3H2 | 0.0 (0/14) | 0.0 | 0.0 (0/0) | 1.0 | 74.9 | 3 |
| 2119 | RF00812 mir-83 | 0.0 (0/33) | 0.0 | 0.0 (0/1) | 0.5 | 85.2 | 4 |
| 2120 | RF01108 BMV3 UPD-PK1 | 0.0 (0/10) | 0.0 | 0.0 (0/0) | 0.0 | 100.0 | 2 |
| 2121 | RF01841 veev FSE | 0.0 (0/9) | 0.0 | 0.0 (0/1) | 1.7 | 88.5 | 10 |
| 2122 | RF02821 V IGR5 | 0.0 (0/29) | 0.0 | 0.0 (0/0) | 0.6 | 61.3 | 2 |
| 2123 | RF02136 HTT-AS1 3 | 0.0 (0/0) | 0.0 | 0.0 (0/0) | 0.0 | 82.2 | 4 |
| 2124 | RF02252 Six3os1 7 | 0.0 (0/0) | 0.0 | 0.0 (0/2) | 0.0 | 83.4 | 5 |
| 2125 | RF02828 Scr3920 | 0.0 (0/39) | 0.0 | 0.0 (0/0) | 0.3 | 93.7 | 6 |
| 2126 | RF01948 KCNQ1OT1 3 | 0.0 (0/0) | 0.0 | 0.0 (0/0) | 0.0 | 89.8 | 9 |
| 2127 | RF00961 mir-581 | 0.0 (0/34) | 0.0 | 0.0 (0/0) | 0.1 | 94.8 | 2 |
| 2128 | RF00439 SNORD87 | 0.0 (0/7) | 0.0 | 0.0 (0/0) | 4.3 | 90.7 | 10 |
| 2129 | RF02128 GNAS-AS1 2 | 0.0 (0/0) | 0.0 | 0.0 (0/0) | 0.0 | 69.3 | 16 |
| 2130 | RF00776 mir-540 | 0.0 (0/28) | 0.0 | 0.0 (0/0) | 0.1 | 85.5 | 2 |
| 2131 | RF02173 SMAD5-AS1 1 | 0.0 (0/0) | 0.0 | 0.0 (0/0) | 0.0 | 71.5 | 18 |
| 2132 | RF02049 STnc460 | 0.0 (0/47) | 0.0 | 0.0 (0/0) | 0.1 | 79.0 | 2 |
| 2133 | RF00385 IBV D-RNA | 0.0 (0/15) | 0.0 | 0.0 (0/0) | 0.2 | 93.9 | 10 |
| 2134 | RF02468 Ms IGR-4 | 0.0 (0/10) | 0.0 | 0.0 (0/0) | 0.2 | 94.7 | 2 |
| 2135 | RF02878 MH s7 | 0.0 (0/60) | 0.0 | 0.0 (0/2) | 0.9 | 77.0 | 3 |
| 2136 | RF00399 SNORA24 | 0.0 (0/33) | 0.0 | 0.0 (0/0) | 0.7 | 88.8 | 5 |
| 2137 | RF02703 AsxR | 0.0 (0/9) | 0.0 | 0.0 (0/0) | 0.1 | 98.1 | 2 |
| 2138 | RF01342 CRISPR-DR32 | 0.0 (0/4) | 0.0 | 0.0 (0/0) | 0.0 | 81.1 | 2 |
| 2139 | RF00447 IRES Kv1 4 | 0.0 (0/51) | 0.0 | 0.0 (0/0) | 0.5 | 89.1 | 6 |
| 2140 | RF02596 cyano het sRNA | 0.0 (0/15) | 0.0 | 0.0 (0/1) | 0.4 | 90.9 | 5 |
| 2141 | RF02315 TtnuHACA8 | 0.0 (0/38) | 0.0 | 0.0 (0/0) | 0.4 | 88.0 | 3 |
| 2142 | RF02571 Mcr7 | 0.0 (0/124) | 0.0 | 0.0 (0/0) | 0.0 | 96.3 | 2 |
| 2143 | RF02427 SpF10 sRNA | 0.0 (0/17) | 0.0 | 0.0 (0/0) | 0.1 | 98.2 | 2 |
| 2144 | RF02454 ncr982 | 0.0 (0/14) | 0.0 | 0.0 (0/0) | 1.3 | 79.7 | 6 |
| 2145 | RF01127 sR42 | 0.0 (0/0) | 0.0 | 0.0 (0/0) | 0.0 | 83.1 | 3 |
| 2146 | RF02775 SodB thermometer | 0.0 (0/24) | 0.0 | 0.0 (0/1) | 1.3 | 85.6 | 5 |
| 2147 | RF02727 sno ZL116 | 0.0 (0/20) | 0.0 | 0.0 (0/0) | 0.4 | 95.0 | 5 |
| 2148 | RF02893 RpsF leader | 0.0 (0/16) | 0.0 | 0.0 (0/0) | 0.2 | 89.8 | 10 |
| 2149 | RF02282 TtnuCD5 | 0.0 (0/0) | 0.0 | 0.0 (0/0) | 0.0 | 97.4 | 2 |
| 2150 | RF02317 TtnuHACA10 | 0.0 (0/23) | 0.0 | 0.0 (0/0) | 0.4 | 86.5 | 3 |
| 2151 | RF01278 SNORD109A | 0.0 (0/5) | 0.0 | 0.0 (0/0) | 0.6 | 87.2 | 7 |
| 2152 | RF02538 TarB | 0.0 (0/18) | 0.0 | 0.0 (0/0) | 0.0 | 94.9 | 2 |
| 2153 | RF00985 mir-640 | 0.0 (0/29) | 0.0 | 0.0 (0/0) | 0.0 | 97.2 | 3 |
| 2154 | RF02865 ncS01 | 0.0 (0/39) | 0.0 | 0.0 (0/0) | 0.5 | 89.2 | 4 |
| 2155 | RF02837 Bcj7 | 0.0 (0/50) | 0.0 | 0.0 (0/0) | 0.5 | 88.7 | 3 |
| 2156 | RF00752 mir-14 | 0.0 (0/22) | 0.0 | 0.0 (0/0) | 1.3 | 80.8 | 6 |
| 2157 | RF00769 mir-331 | 0.0 (0/26) | 0.0 | 0.0 (0/0) | 0.1 | 93.8 | 5 |
| 2158 | RF01963 RMST 2 | 0.0 (0/0) | 0.0 | 0.0 (0/0) | 0.0 | 82.8 | 16 |
| 2159 | RF02488 GlsR25 | 0.0 (0/29) | 0.0 | 0.0 (0/0) | 0.7 | 85.8 | 3 |
| 2160 | RF01681 P4 | 0.0 (0/70) | 0.0 | 0.0 (0/0) | 0.0 | 96.7 | 2 |
| 2161 | RF01300 snoU49 | 0.0 (0/8) | 0.0 | 0.0 (0/0) | 0.8 | 80.4 | 5 |
| 2162 | RF02662 BLi r0872 | 0.0 (0/48) | 0.0 | 0.0 (0/0) | 0.0 | 98.6 | 2 |
| 2163 | RF01357 CRISPR-DR31 | 0.0 (0/10) | 0.0 | 0.0 (0/0) | 0.4 | 91.0 | 3 |
| 2164 | RF02309 TtnuHACA1 | 0.0 (0/37) | 0.0 | 0.0 (0/0) | 0.0 | 96.5 | 2 |
| 2165 | RF00217 SNORD20 | 0.0 (0/4) | 0.0 | 0.0 (0/0) | 0.8 | 88.3 | 13 |
| 2166 | RF02251 Six3os1 6 | 0.0 (0/0) | 0.0 | 0.0 (0/0) | 0.0 | 83.6 | 4 |
| 2167 | RF02403 Afu 328 | 0.0 (0/0) | 0.0 | 0.0 (0/0) | 0.0 | 82.1 | 4 |
| 2168 | RF01095 PK-CuYV BPYV | 0.0 (0/15) | 0.0 | 0.0 (0/0) | 0.0 | 100.0 | 2 |
| 2169 | RF02708 TeloSII ncR40 | 0.0 (0/47) | 0.0 | 0.0 (0/0) | 0.1 | 93.8 | 2 |
| 2170 | RF01973 H19 2 | 0.0 (0/0) | 0.0 | 0.0 (0/0) | 0.0 | 89.8 | 23 |
| 2171 | RF02254 mir-35 | 0.0 (0/40) | 0.0 | 0.0 (0/0) | 0.1 | 76.0 | 2 |
| 2172 | RF02537 TarA | 0.0 (0/22) | 0.0 | 0.0 (0/0) | 0.0 | 96.9 | 2 |
| 2173 | RF01528 CC3513 | 0.0 (0/53) | 0.0 | 0.0 (0/0) | 0.1 | 81.3 | 2 |

Continued on next page

| | RNA family (seed alignment) | Sensitivity annotated bpairs that covary % (cov_bps/bps) | Power average power % | Positive Predictive Value covarying pairs in structure % (cov_bps/cov_pairs) | average substitutions per bpair | avg pairwise identity % | number of sequences |
|---|---|---|---|---|---|---|---|
| 2174 | RF00709 mir-455 | 0.0 (0/29) | 0.0 | 0.0 (0/0) | 0.3 | 93.0 | 5 |
| 2175 | RF02026 mir-2833 | 0.0 (0/23) | 0.0 | 0.0 (0/0) | 0.5 | 94.6 | 5 |
| 2176 | RF01967 RMST 6 | 0.0 (0/0) | 0.0 | 0.0 (0/0) | 0.0 | 84.5 | 18 |
| 2177 | RF02220 ZNRD1-AS1 3 | 0.0 (0/0) | 0.0 | 0.0 (0/0) | 0.0 | 92.3 | 9 |
| 2178 | RF01375 CRISPR-DR62 | 0.0 (0/4) | 0.0 | 0.0 (0/0) | 0.0 | 91.9 | 2 |
| 2179 | RF02880 MH s15 | 0.0 (0/91) | 0.0 | 0.0 (0/0) | 0.2 | 70.2 | 2 |
| 2180 | RF00306 snoZ178 | 0.0 (0/25) | 0.0 | 0.0 (0/0) | 0.0 | 99.7 | 5 |
| 2181 | RF02165 PVT1 2 | 0.0 (0/0) | 0.0 | 0.0 (0/0) | 0.0 | 69.0 | 17 |
| 2182 | RF02903 AaHKsRNA96 | 0.0 (0/38) | 0.0 | 0.0 (0/0) | 0.7 | 74.4 | 3 |
| 2183 | RF01445 S pombe snR94 | 0.0 (0/50) | 0.0 | 0.0 (0/0) | 1.2 | 74.0 | 3 |
| 2184 | RF02581 tsr32 | 0.0 (0/51) | 0.0 | 0.0 (0/0) | 0.5 | 95.7 | 4 |
| 2185 | RF00770 mir-330 | 0.0 (0/32) | 0.0 | 0.0 (0/0) | 0.6 | 88.5 | 6 |
| 2186 | RF02102 DISC2 | 0.0 (0/0) | 0.0 | 0.0 (0/0) | 0.0 | 86.8 | 5 |
| 2187 | RF02851 Ysr283 | 0.0 (0/60) | 0.0 | 0.0 (0/1) | 0.7 | 79.7 | 3 |
| 2188 | RF01864 plasmodium snoR21 | 0.0 (0/4) | 0.0 | 0.0 (0/3) | 0.0 | 83.5 | 4 |
| 2189 | RF00857 mir-233 | 0.0 (0/31) | 0.0 | 0.0 (0/0) | 0.6 | 82.3 | 4 |
| 2190 | RF02697 LDH1 5p UTR | 0.0 (0/22) | 0.0 | 0.0 (0/0) | 0.1 | 97.4 | 4 |
| 2191 | RF00935 mir-876 | 0.0 (0/32) | 0.0 | 0.0 (0/0) | 0.6 | 91.8 | 3 |
| 2192 | RF01103 UPD-PKc | 0.0 (0/11) | 0.0 | 0.0 (0/0) | 0.1 | 96.5 | 2 |
| 2193 | RF01382 HIV-1 SL4 | 0.0 (0/5) | 0.0 | 0.0 (0/0) | 0.0 | 91.8 | 16 |
| 2194 | RF01566 DdR16 | 0.0 (0/0) | 0.0 | 0.0 (0/0) | 0.0 | 85.4 | 4 |
| 2195 | RF02778 FdoG1 thermometer | 0.0 (0/48) | 0.0 | 0.0 (0/0) | 0.6 | 87.3 | 4 |
| 2196 | RF02157 NPPA-AS1 2 | 0.0 (0/0) | 0.0 | 0.0 (0/0) | 0.0 | 77.0 | 16 |
| 2197 | RF01366 CRISPR-DR53 | 0.0 (0/9) | 0.0 | 0.0 (0/0) | 0.0 | 97.3 | 2 |
| 2198 | RF02849 Ysr197 | 0.0 (0/60) | 0.0 | 0.0 (0/0) | 1.1 | 80.0 | 3 |
| 2199 | RF02643 Spy491738 | 0.0 (0/22) | 0.0 | 0.0 (0/0) | 0.3 | 90.9 | 4 |
| 2200 | RF00845 MIR158 | 0.0 (0/33) | 0.0 | 0.0 (0/0) | 0.0 | 90.0 | 2 |
| 2201 | RF01236 snoU19 | 0.0 (0/35) | 0.0 | 0.0 (0/0) | 1.1 | 67.7 | 3 |
| 2202 | RF02561 CbSR12 | 0.0 (0/31) | 0.0 | 0.0 (0/0) | 0.0 | 99.3 | 2 |
| 2203 | RF01318 CRISPR-DR5 | 0.0 (0/7) | 0.0 | 0.0 (0/0) | 0.0 | 78.0 | 12 |
| 2204 | RF02006 mir-1253 | 0.0 (0/38) | 0.0 | 0.0 (0/0) | 0.1 | 95.2 | 2 |
| 2205 | RF01567 DdR17 | 0.0 (0/0) | 0.0 | 0.0 (0/0) | 0.0 | 100.0 | 2 |
| 2206 | RF02646 Cis8 sRNA | 0.0 (0/40) | 0.0 | 0.0 (0/0) | 0.2 | 75.8 | 2 |
| 2207 | RF00869 mir-BART7 | 0.0 (0/33) | 0.0 | 0.0 (0/0) | 0.1 | 84.5 | 2 |
| 2208 | RF02721 sca ncR26 | 0.0 (0/18) | 0.0 | 0.0 (0/0) | 0.3 | 85.1 | 2 |
| 2209 | RF01453 RCNMV TE DR1 | 0.0 (0/43) | 0.0 | 0.0 (0/0) | 0.1 | 88.7 | 3 |
| 2210 | RF02580 tsr31 | 0.0 (0/18) | 0.0 | 0.0 (0/0) | 0.1 | 94.7 | 4 |
| 2211 | RF01381 HIV-1 SL3 | 0.0 (0/5) | 0.0 | 0.0 (0/0) | 0.6 | 90.3 | 19 |
| 2212 | RF01096 PK-HAV | 0.0 (0/17) | 0.0 | 0.0 (0/0) | 0.0 | 87.3 | 2 |
| 2213 | RF02287 TtnuCD10 | 0.0 (0/0) | 0.0 | 0.0 (0/0) | 0.0 | 79.5 | 4 |
| 2214 | RF01124 sR36 | 0.0 (0/0) | 0.0 | 0.0 (0/0) | 0.0 | 84.2 | 5 |
| 2215 | RF02890 SprC | 0.0 (0/36) | 0.0 | 0.0 (0/0) | 0.3 | 64.3 | 2 |
| 2216 | RF01986 ZEB2 AS1 3 | 0.0 (0/0) | 0.0 | 0.0 (0/0) | 0.0 | 90.2 | 11 |
| 2217 | RF02833 Scr3202 | 0.0 (0/28) | 0.0 | 0.0 (0/0) | 0.8 | 82.5 | 4 |
| 2218 | RF01450 S pombe snR96 | 0.0 (0/50) | 0.0 | 0.0 (0/0) | 0.0 | 99.5 | 2 |
| 2219 | RF02781 ManX thermometer | 0.0 (0/33) | 0.0 | 0.0 (0/0) | 0.7 | 82.3 | 4 |
| 2220 | RF02621 BSnc120 | 0.0 (0/23) | 0.0 | 0.0 (0/0) | 0.2 | 98.5 | 3 |
| 2221 | RF01542 TB11Cs5H1 | 0.0 (0/17) | 0.0 | 0.0 (0/0) | 0.1 | 74.3 | 2 |
| 2222 | RF01184 snR79 | 0.0 (0/0) | 0.0 | 0.0 (0/0) | 0.0 | 75.5 | 18 |
| 2223 | RF00067 SNORD15 | 0.0 (0/4) | 0.0 | 0.0 (0/0) | 0.2 | 60.9 | 11 |
| 2224 | RF01873 PISRT1 | 0.0 (0/0) | 0.0 | 0.0 (0/0) | 0.0 | 88.1 | 18 |
| 2225 | RF02476 GlsR7 | 0.0 (0/3) | 0.0 | 0.0 (0/0) | 0.0 | 88.1 | 3 |
| 2226 | RF02715 sno ncR1 | 0.0 (0/32) | 0.0 | 0.0 (0/0) | 0.8 | 87.1 | 5 |
| 2227 | RF02527 SSRC38 | 0.0 (0/44) | 0.0 | 0.0 (0/0) | 0.0 | 97.6 | 3 |
| 2228 | RF02705 B rapa snoR775 | 0.0 (0/29) | 0.0 | 0.0 (0/0) | 0.0 | 96.1 | 2 |
| 2229 | RF00925 MIR1027 | 0.0 (0/43) | 0.0 | 0.0 (0/0) | 0.0 | 100.0 | 2 |
| 2230 | RF02007 mir-1237 | 0.0 (0/38) | 0.0 | 0.0 (0/0) | 0.1 | 96.1 | 3 |
| 2231 | RF02112 DLG2-AS1 1 | 0.0 (0/0) | 0.0 | 0.0 (0/0) | 0.0 | 80.9 | 21 |
| 2232 | RF00266 snoZ17 | 0.0 (0/6) | 0.0 | 0.0 (0/0) | 0.8 | 76.9 | 26 |
| 2233 | RF00941 mir-434 | 0.0 (0/31) | 0.0 | 0.0 (0/0) | 0.1 | 89.4 | 2 |
| 2234 | RF01962 RMST 1 | 0.0 (0/0) | 0.0 | 0.0 (0/0) | 0.0 | 83.8 | 14 |
| 2235 | RF02622 BSnc121 | 0.0 (0/31) | 0.0 | 0.0 (0/0) | 0.0 | 99.1 | 2 |
| 2236 | RF01075 TLS-PK1 | 0.0 (0/31) | 0.0 | 0.0 (0/0) | 0.0 | 87.5 | 2 |
| 2237 | RF01177 snR67 | 0.0 (0/0) | 0.0 | 0.0 (0/0) | 0.0 | 73.7 | 16 |
| 2238 | RF00305 snoZ248 | 0.0 (0/26) | 0.0 | 0.0 (0/0) | 0.0 | 98.1 | 5 |
| 2239 | RF01915 mir-2238 | 0.0 (0/31) | 0.0 | 0.0 (0/0) | 1.2 | 79.5 | 4 |
| 2240 | RF02055 STnc380 | 0.0 (0/28) | 0.0 | 0.0 (0/0) | 0.9 | 79.8 | 5 |
| 2241 | RF01365 CRISPR-DR52 | 0.0 (0/7) | 0.0 | 0.0 (0/0) | 0.0 | 100.0 | 2 |
| 2242 | RF02075 STnc230 | 0.0 (0/12) | 0.0 | 0.0 (0/0) | 1.1 | 67.4 | 11 |
| 2243 | RF01126 sR41 | 0.0 (0/0) | 0.0 | 0.0 (0/0) | 0.0 | 73.3 | 7 |
| 2244 | RF01461 rli24 | 0.0 (0/35) | 0.0 | 0.0 (0/0) | 0.0 | 94.3 | 6 |
| 2245 | RF01618 ceN114 | 0.0 (0/3) | 0.0 | 0.0 (0/0) | 0.0 | 97.8 | 3 |
| 2246 | RF02301 TtnuCD25 | 0.0 (0/0) | 0.0 | 0.0 (0/0) | 0.0 | 83.3 | 3 |
| 2247 | RF01503 Afu 203 | 0.0 (0/35) | 0.0 | 0.0 (0/0) | 0.2 | 92.1 | 3 |
| 2248 | RF01235 snR68 | 0.0 (0/38) | 0.0 | 0.0 (0/0) | 0.2 | 95.2 | 5 |
| 2249 | RF01352 CRISPR-DR43 | 0.0 (0/0) | 0.0 | 0.0 (0/0) | 0.0 | 86.2 | 3 |
| 2250 | RF01021 mir-558 | 0.0 (0/34) | 0.0 | 0.0 (0/0) | 0.4 | 95.7 | 3 |
| 2251 | RF02626 WsnRNA59 | 0.0 (0/23) | 0.0 | 0.0 (0/0) | 0.3 | 76.1 | 2 |
| 2252 | RF01654 ceN82 | 0.0 (0/30) | 0.0 | 0.0 (0/0) | 0.7 | 87.4 | 6 |
| 2253 | RF01584 snoR03 | 0.0 (0/38) | 0.0 | 0.0 (0/0) | 0.2 | 75.7 | 2 |
| 2254 | RF01149 sR10 | 0.0 (0/0) | 0.0 | 0.0 (0/0) | 0.0 | 85.5 | 4 |
| 2255 | RF01586 snoR09 | 0.0 (0/3) | 0.0 | 0.0 (0/0) | 0.0 | 93.7 | 3 |
| 2256 | RF02653 StyR-143 | 0.0 (0/44) | 0.0 | 0.0 (0/0) | 0.0 | 96.5 | 2 |
| 2257 | RF02461 Virus CITE 6 | 0.0 (0/34) | 0.0 | 0.0 (0/0) | 1.3 | 85.1 | 3 |

Continued on next page

| | RNA family (seed alignment) | Sensitivity annotated bpairs that covary % (cov_bps/bps) | Power average power % | Positive Predictive Value covarying pairs in structure % (cov_bps/cov_pairs) | average substitutions per bpair | avg pairwise identity % | number of sequences |
|---|---|---|---|---|---|---|---|
| 2258 | RF00326 snoZ155 | 0.0 (0/6) | 0.0 | 0.0 (0/0) | 4.2 | 78.8 | 8 |
| 2259 | RF01495 ACAT | 0.0 (0/32) | 0.0 | 0.0 (0/0) | 1.5 | 80.8 | 8 |
| 2260 | RF02205 WT1-AS 3 | 0.0 (0/0) | 0.0 | 0.0 (0/0) | 0.0 | 71.5 | 19 |
| 2261 | RF01839 eeev FSE | 0.0 (0/10) | 0.0 | 0.0 (0/0) | 0.5 | 93.2 | 6 |
| 2262 | RF02155 NCRUPAR 2 | 0.0 (0/0) | 0.0 | 0.0 (0/0) | 0.0 | 92.5 | 3 |
| 2263 | RF02845 Ref1C | 0.0 (0/34) | 0.0 | 0.0 (0/0) | 0.6 | 59.5 | 2 |
| 2264 | RF02825 LPR17 | 0.0 (0/69) | 0.0 | 0.0 (0/0) | 0.2 | 87.7 | 2 |
| 2265 | RF02616 SSRC8 2 | 0.0 (0/87) | 0.0 | 0.0 (0/0) | 0.0 | 97.2 | 2 |
| 2266 | RF02460 Virus CITE 5 | 0.0 (0/32) | 0.0 | 0.0 (0/0) | 0.1 | 83.7 | 2 |
| 2267 | RF01572 DdR4 | 0.0 (0/0) | 0.0 | 0.0 (0/0) | 0.0 | 86.2 | 3 |
| 2268 | RF00898 mir-242 | 0.0 (0/23) | 0.0 | 0.0 (0/0) | 0.0 | 85.7 | 2 |
| 2269 | RF02137 HOXA11-AS1 1 | 0.0 (0/0) | 0.0 | 0.0 (0/0) | 0.0 | 88.5 | 15 |
| 2270 | RF01509 Afu 300 | 0.0 (0/0) | 0.0 | 0.0 (0/0) | 0.0 | 73.3 | 10 |
| 2271 | RF02039 SPRY4-IT1 2 | 0.0 (0/0) | 0.0 | 0.0 (0/0) | 0.0 | 70.0 | 24 |
| 2272 | RF01204 snR65 | 0.0 (0/0) | 0.0 | 0.0 (0/0) | 0.0 | 92.7 | 5 |
| 2273 | RF02193 TCL6 3 | 0.0 (0/0) | 0.0 | 0.0 (0/0) | 0.0 | 76.6 | 26 |
| 2274 | RF00331 snoZ169 | 0.0 (0/6) | 0.0 | 0.0 (0/0) | 1.5 | 87.4 | 3 |
| 2275 | RF01338 CRISPR-DR25 | 0.0 (0/6) | 0.0 | 0.0 (0/0) | 0.7 | 92.8 | 5 |
| 2276 | RF00816 mir-245 | 0.0 (0/18) | 0.0 | 0.0 (0/0) | 0.2 | 87.3 | 3 |
| 2277 | RF02767 Ysr186 sR026 CsrC | 0.0 (0/87) | 0.0 | 0.0 (0/0) | 0.7 | 87.3 | 4 |
| 2278 | RF02132 HOXB13-AS1 1 | 0.0 (0/0) | 0.0 | 0.0 (0/0) | 0.0 | 73.4 | 12 |
| 2279 | RF01833 astro FSE | 0.0 (0/12) | 0.0 | 0.0 (0/0) | 0.3 | 90.3 | 4 |
| 2280 | RF00861 mir-488 | 0.0 (0/30) | 0.0 | 0.0 (0/0) | 0.3 | 92.9 | 5 |
| 2281 | RF02209 WT1-AS 7 | 0.0 (0/0) | 0.0 | 0.0 (0/0) | 0.0 | 84.8 | 21 |
| 2282 | RF00087 SNORD26 | 0.0 (0/4) | 0.0 | 0.0 (0/1) | 2.2 | 79.2 | 17 |
| 2283 | RF02126 GHRLOS | 0.0 (0/0) | 0.0 | 0.0 (0/0) | 0.0 | 85.5 | 16 |
| 2284 | RF02131 GNAS-AS1 5 | 0.0 (0/0) | 0.0 | 0.0 (0/0) | 0.0 | 81.8 | 22 |
| 2285 | RF01428 snoR128 | 0.0 (0/3) | 0.0 | 0.0 (0/0) | 0.0 | 74.1 | 5 |
| 2286 | RF01590 plasmodium snoR14 | 0.0 (0/3) | 0.0 | 0.0 (0/0) | 0.0 | 86.4 | 4 |
| 2287 | RF01534 TB10Cs4H4 | 0.0 (0/18) | 0.0 | 0.0 (0/0) | 0.2 | 77.8 | 2 |
| 2288 | RF02696 Teg49 | 0.0 (0/49) | 0.0 | 0.0 (0/0) | 0.5 | 93.8 | 6 |
| 2289 | RF01100 PK-BYV | 0.0 (0/11) | 0.0 | 0.0 (0/0) | 0.0 | 100.0 | 2 |
| 2290 | RF00974 mir-607 | 0.0 (0/41) | 0.0 | 0.0 (0/0) | 0.3 | 94.4 | 3 |
| 2291 | RF02217 ZNFX1-AS1 3 | 0.0 (0/0) | 0.0 | 0.0 (0/0) | 0.0 | 92.9 | 3 |
| 2292 | RF02854 Ysr100 | 0.0 (0/30) | 0.0 | 0.0 (0/0) | 0.6 | 86.4 | 3 |
| 2293 | RF01244 snR4 | 0.0 (0/54) | 0.0 | 0.0 (0/1) | 1.5 | 81.1 | 5 |
| 2294 | RF00978 mir-638 | 0.0 (0/22) | 0.0 | 0.0 (0/0) | 0.4 | 90.7 | 6 |
| 2295 | RF00294 snoTBR17 | 0.0 (0/2) | 0.0 | 0.0 (0/0) | 2.5 | 76.8 | 6 |
| 2296 | RF01964 RMST 3 | 0.0 (0/0) | 0.0 | 0.0 (0/0) | 0.0 | 89.1 | 16 |
| 2297 | RF00900 mir-255 | 0.0 (0/25) | 0.0 | 0.0 (0/0) | 0.0 | 69.9 | 2 |
| 2298 | RF02327 TtnuHACA20 | 0.0 (0/38) | 0.0 | 0.0 (0/0) | 0.0 | 94.3 | 2 |
| 2299 | RF01363 CRISPR-DR50 | 0.0 (0/13) | 0.0 | 0.0 (0/0) | 0.2 | 82.6 | 2 |
| 2300 | RF00496 Corona SL-III | 0.0 (0/8) | 0.0 | 0.0 (0/0) | 0.5 | 93.3 | 5 |
| 2301 | RF01141 sR18 | 0.0 (0/0) | 0.0 | 0.0 (0/0) | 0.0 | 71.5 | 5 |
| 2302 | RF01350 CRISPR-DR41 | 0.0 (0/8) | 0.0 | 0.0 (0/0) | 0.0 | 93.1 | 2 |
| 2303 | RF01223 snR13 | 0.0 (0/2) | 0.0 | 0.0 (0/0) | 0.5 | 95.1 | 2 |
| 2304 | RF01053 Deinococcus Y RNA | 0.0 (0/35) | 0.0 | 0.0 (0/0) | 0.0 | 100.0 | 2 |
| 2305 | RF01953 SOX2OT exon3 | 0.0 (0/0) | 0.0 | 0.0 (0/1) | 0.0 | 83.2 | 11 |
| 2306 | RF02793 asR3 | 0.0 (0/17) | 0.0 | 0.0 (0/0) | 0.2 | 79.8 | 3 |
| 2307 | RF02014 mir-1178 | 0.0 (0/28) | 0.0 | 0.0 (0/0) | 0.0 | 94.0 | 3 |
| 2308 | RF02686 RAGATH-6 | 0.0 (0/35) | 0.0 | 0.0 (0/0) | 0.0 | 87.3 | 2 |
| 2309 | RF01829 sR6 | 0.0 (0/8) | 0.0 | 0.0 (0/0) | 0.0 | 100.0 | 2 |
| 2310 | RF02208 WT1-AS 6 | 0.0 (0/0) | 0.0 | 0.0 (0/0) | 0.0 | 78.9 | 19 |
| 2311 | RF02493 Gl U4 | 0.0 (0/13) | 0.0 | 0.0 (0/0) | 0.0 | 92.6 | 2 |
| 2312 | RF02183 ST7-OT3 1 | 0.0 (0/0) | 0.0 | 0.0 (0/0) | 0.0 | 79.3 | 28 |
| 2313 | RF02385 sau-13 | 0.0 (0/25) | 0.0 | 0.0 (0/1) | 1.8 | 79.5 | 6 |
| 2314 | RF01142 sR19 | 0.0 (0/0) | 0.0 | 0.0 (0/0) | 0.0 | 91.5 | 3 |
| 2315 | RF00118 rydB | 0.0 (0/12) | 0.0 | 0.0 (0/0) | 0.2 | 78.9 | 7 |
| 2316 | RF02533 HAV CRE | 0.0 (0/35) | 0.0 | 0.0 (0/0) | 0.1 | 91.2 | 2 |
| 2317 | RF02601 BASRCI337 | 0.0 (0/57) | 0.0 | 0.0 (0/0) | 0.1 | 98.4 | 4 |
| 2318 | RF02367 Yfr20 | 0.0 (0/25) | 0.0 | 0.0 (0/0) | 1.2 | 79.5 | 7 |
| 2319 | RF02794 Pab19 | 0.0 (0/31) | 0.0 | 0.0 (0/0) | 0.9 | 80.9 | 3 |
| 2320 | RF02835 Bcj11 | 0.0 (0/39) | 0.0 | 0.0 (0/0) | 0.9 | 80.7 | 4 |
| 2321 | RF02814 Sso133 | 0.0 (0/19) | 0.0 | 0.0 (0/0) | 0.6 | 80.4 | 3 |
| 2322 | RF02088 STnc510 | 0.0 (0/230) | 0.0 | 0.0 (0/0) | 0.6 | 87.7 | 5 |
| 2323 | RF01683 P6 | 0.0 (0/151) | 0.0 | 0.0 (0/0) | 0.1 | 95.4 | 3 |
| 2324 | RF01305 sR51 | 0.0 (0/12) | 0.0 | 0.0 (0/0) | 1.3 | 80.4 | 4 |
| 2325 | RF02164 PVT1 1 | 0.0 (0/0) | 0.0 | 0.0 (0/0) | 0.0 | 74.0 | 17 |
| 2326 | RF01933 bxd 5 | 0.0 (0/0) | 0.0 | 0.0 (0/0) | 0.0 | 96.1 | 5 |
| 2327 | RF02065 STnc340 | 0.0 (0/15) | 0.0 | 0.0 (0/0) | 0.4 | 78.2 | 4 |
| 2328 | RF02178 SMCR2 2 | 0.0 (0/0) | 0.0 | 0.0 (0/0) | 0.0 | 69.7 | 5 |
| 2329 | RF01934 bxd 6 | 0.0 (0/0) | 0.0 | 0.0 (0/0) | 0.0 | 87.3 | 6 |
| 2330 | RF00676 mir-127 | 0.0 (0/31) | 0.0 | 0.0 (0/0) | 0.0 | 98.8 | 5 |
| 2331 | RF00829 mir-149 | 0.0 (0/34) | 0.0 | 0.0 (0/0) | 0.3 | 91.0 | 3 |
| 2332 | RF02008 mir-621 | 0.0 (0/33) | 0.0 | 0.0 (0/0) | 0.2 | 95.8 | 3 |
| 2333 | RF01672 P2 | 0.0 (0/47) | 0.0 | 0.0 (0/0) | 0.0 | 99.2 | 7 |
| 2334 | RF01094 RF site6 | 0.0 (0/16) | 0.0 | 0.0 (0/0) | 0.0 | 87.3 | 2 |
| 2335 | RF02611 BSR0653 | 0.0 (0/181) | 0.0 | 0.0 (0/0) | 0.0 | 99.6 | 4 |
| 2336 | RF00119 C0299 | 0.0 (0/20) | 0.0 | 0.0 (0/0) | 0.2 | 96.7 | 5 |
| 2337 | RF02804 PyrR206 | 0.0 (0/14) | 0.0 | 0.0 (0/0) | 0.6 | 88.7 | 4 |
| 2338 | RF00207 K10 TLS | 0.0 (0/17) | 0.0 | 0.0 (0/0) | 0.0 | 100.0 | 5 |
| 2339 | RF01932 bxd 4 | 0.0 (0/0) | 0.0 | 0.0 (0/0) | 0.0 | 78.0 | 5 |
| 2340 | RF01615 ceN111 | 0.0 (0/3) | 0.0 | 0.0 (0/0) | 0.0 | 87.5 | 3 |
| 2341 | RF02733 ToxT thermometer | 0.0 (0/20) | 0.0 | 0.0 (0/0) | 0.2 | 94.4 | 3 |

Continued on next page

| | RNA family (seed alignment) | Sensitivity annotated bpairs that covary % (cov_bps/bps) | Power average power % | Positive Predictive Value covarying pairs in structure % (cov_bps/cov_pairs) | average substitutions per bpair | avg pairwise identity % | number of sequences |
|---|---|---|---|---|---|---|---|
| 2342 | RF00433 Hsp90 CRE | 0.0 (0/49) | 0.0 | 0.0 (0/0) | 0.5 | 95.6 | 6 |
| 2343 | RF02652 StyR-3 | 0.0 (0/40) | 0.0 | 0.0 (0/0) | 0.0 | 98.8 | 5 |
| 2344 | RF02635 EF0820 EF0821 | 0.0 (0/117) | 0.0 | 0.0 (0/0) | 0.0 | 99.7 | 2 |
| 2345 | RF00790 mir-358 | 0.0 (0/30) | 0.0 | 0.0 (0/1) | 0.7 | 68.3 | 3 |
| 2346 | RF00868 mir-BART15 | 0.0 (0/27) | 0.0 | 0.0 (0/0) | 0.1 | 79.5 | 2 |
| 2347 | RF01190 snR50 | 0.0 (0/2) | 0.0 | 0.0 (0/0) | 0.0 | 91.3 | 3 |
| 2348 | RF01466 rli34 | 0.0 (0/10) | 0.0 | 0.0 (0/0) | 0.4 | 87.0 | 5 |
| 2349 | RF02831 Scr2736 | 0.0 (0/18) | 0.0 | 0.0 (0/0) | 0.0 | 96.8 | 4 |
| 2350 | RF02307 TtnuCD33 | 0.0 (0/0) | 0.0 | 0.0 (0/0) | 0.0 | 93.2 | 2 |
| 2351 | RF02107 DLEU2 3 | 0.0 (0/0) | 0.0 | 0.0 (0/0) | 0.0 | 82.4 | 24 |
| 2352 | RF01040 mir-573 | 0.0 (0/29) | 0.0 | 0.0 (0/0) | 0.1 | 90.9 | 2 |
| 2353 | RF02298 TtnuCD22 | 0.0 (0/0) | 0.0 | 0.0 (0/0) | 0.0 | 97.4 | 2 |
| 2354 | RF02091 DAOA-AS1 2 | 0.0 (0/0) | 0.0 | 0.0 (0/0) | 0.0 | 70.8 | 24 |
| 2355 | RF02182 ST7-AS2 2 | 0.0 (0/0) | 0.0 | 0.0 (0/0) | 0.0 | 81.7 | 8 |
| 2356 | RF01817 RsaB | 0.0 (0/15) | 0.0 | 0.0 (0/0) | 0.1 | 98.2 | 2 |
| 2357 | RF02648 Cis90 sRNA | 0.0 (0/73) | 0.0 | 0.0 (0/0) | 0.4 | 65.7 | 2 |
| 2358 | RF02744 Rev39 5p UTR | 0.0 (0/93) | 0.0 | 0.0 (0/0) | 0.3 | 92.7 | 4 |
| 2359 | RF02249 Six3os1 4 | 0.0 (0/0) | 0.0 | 0.0 (0/0) | 0.0 | 81.8 | 7 |
| 2360 | RF01101 TLS-PK6 | 0.0 (0/8) | 0.0 | 0.0 (0/0) | 0.0 | 88.3 | 3 |
| 2361 | RF01625 ceN28 | 0.0 (0/4) | 0.0 | 0.0 (0/0) | 0.0 | 90.8 | 4 |
| 2362 | RF01972 H19 1 | 0.0 (0/0) | 0.0 | 0.0 (0/0) | 0.0 | 91.5 | 35 |
| 2363 | RF02800 Rp sR47 | 0.0 (0/88) | 0.0 | 0.0 (0/0) | 0.1 | 92.4 | 2 |
| 2364 | RF02632 Hrs10 | 0.0 (0/42) | 0.0 | 0.0 (0/0) | 0.1 | 94.6 | 2 |
| 2365 | RF02333 TtnuHACA27 | 0.0 (0/17) | 0.0 | 0.0 (0/0) | 0.0 | 95.3 | 2 |
| 2366 | RF00992 mir-593 | 0.0 (0/33) | 0.0 | 0.0 (0/0) | 0.0 | 95.0 | 2 |
| 2367 | RF01956 NEAT1 2 | 0.0 (0/0) | 0.0 | 0.0 (0/0) | 0.0 | 84.1 | 13 |
| 2368 | RF00852 mir-231 | 0.0 (0/26) | 0.0 | 0.0 (0/0) | 0.4 | 83.8 | 4 |
| 2369 | RF01511 Afu 304 | 0.0 (0/0) | 0.0 | 0.0 (0/1) | 0.0 | 73.6 | 5 |
| 2370 | RF01595 snoR19 | 0.0 (0/4) | 0.0 | 0.0 (0/0) | 0.0 | 79.2 | 4 |
| 2371 | RF00839 mir-452 | 0.0 (0/33) | 0.0 | 0.0 (0/0) | 0.7 | 88.8 | 4 |
| 2372 | RF02470 Ms IGR-8 | 0.0 (0/15) | 0.0 | 0.0 (0/0) | 1.2 | 64.5 | 3 |
| 2373 | RF01560 DdR10 | 0.0 (0/0) | 0.0 | 0.0 (0/0) | 0.0 | 84.2 | 4 |
| 2374 | RF01245 snR9 | 0.0 (0/38) | 0.0 | 0.0 (0/0) | 0.6 | 90.5 | 5 |
| 2375 | RF01451 S pombe snR97 | 0.0 (0/26) | 0.0 | 0.0 (0/0) | 0.0 | 100.0 | 3 |
| 2376 | RF02429 SpF14 sRNA | 0.0 (0/36) | 0.0 | 0.0 (0/0) | 1.8 | 80.1 | 4 |
| 2377 | RF02320 TtnuHACA13 | 0.0 (0/37) | 0.0 | 0.0 (0/0) | 0.1 | 92.5 | 2 |
| 2378 | RF02279 TtnuCD1 | 0.0 (0/0) | 0.0 | 0.0 (0/0) | 0.0 | 96.8 | 2 |
| 2379 | RF00803 mir-425 | 0.0 (0/20) | 0.0 | 0.0 (0/0) | 0.7 | 79.8 | 5 |
| 2380 | RF01565 DdR15 | 0.0 (0/0) | 0.0 | 0.0 (0/0) | 0.0 | 84.7 | 3 |
| 2381 | RF02882 MH s36 | 0.0 (0/28) | 0.0 | 0.0 (0/0) | 0.1 | 82.2 | 2 |
| 2382 | RF01611 ceN108 | 0.0 (0/4) | 0.0 | 0.0 (0/0) | 0.0 | 87.6 | 3 |
| 2383 | RF00742 MIR162 2 | 0.0 (0/25) | 0.0 | 0.0 (0/0) | 0.4 | 78.2 | 10 |
| 2384 | RF01312 sR9 | 0.0 (0/1) | 0.0 | 0.0 (0/0) | 2.0 | 41.8 | 3 |
| 2385 | RF01924 mir-2774 | 0.0 (0/17) | 0.0 | 0.0 (0/0) | 0.5 | 87.3 | 4 |
| 2386 | RF00154 SNORD63 | 0.0 (0/4) | 0.0 | 0.0 (0/0) | 1.2 | 80.7 | 23 |
| 2387 | RF02521 Virus CITE 7 | 0.0 (0/27) | 0.0 | 0.0 (0/0) | 0.0 | 89.4 | 2 |
| 2388 | RF01667 rox1 | 0.0 (0/23) | 0.0 | 0.0 (0/0) | 0.0 | 94.3 | 3 |
| 2389 | RF02340 DENV SLA | 0.0 (0/22) | 0.0 | 0.0 (0/0) | 0.7 | 86.2 | 4 |
| 2390 | RF02765 Ysr209 | 0.0 (0/8) | 0.0 | 0.0 (0/0) | 0.1 | 95.7 | 4 |
| 2391 | RF01875 MIAT exon5 1 | 0.0 (0/0) | 0.0 | 0.0 (0/0) | 0.0 | 91.0 | 9 |
| 2392 | RF02822 Srn266 | 0.0 (0/32) | 0.0 | 0.0 (0/0) | 0.6 | 85.4 | 4 |
| 2393 | RF02484 GlsR21 | 0.0 (0/31) | 0.0 | 0.0 (0/0) | 0.2 | 87.2 | 3 |
| 2394 | RF02618 SSRC34 2 | 0.0 (0/39) | 0.0 | 0.0 (0/0) | 0.0 | 98.9 | 3 |
| 2395 | RF01097 RF site8 | 0.0 (0/12) | 0.0 | 0.0 (0/0) | 0.2 | 90.1 | 4 |
| 2396 | RF01658 ceN81 | 0.0 (0/40) | 0.0 | 0.0 (0/0) | 0.2 | 90.4 | 3 |
| 2397 | RF02204 WT1-AS 2 | 0.0 (0/0) | 0.0 | 0.0 (0/0) | 0.0 | 73.4 | 18 |
| 2398 | RF01600 snoR25 | 0.0 (0/4) | 0.0 | 0.0 (0/0) | 1.0 | 91.0 | 4 |
| 2399 | RF01843 neisseria FSE | 0.0 (0/13) | 0.0 | 0.0 (0/0) | 0.5 | 92.9 | 4 |
| 2400 | RF02188 ST7-OT4 2 | 0.0 (0/0) | 0.0 | 0.0 (0/0) | 0.0 | 70.5 | 29 |
| 2401 | RF02313 TtnuHACA6 | 0.0 (0/36) | 0.0 | 0.0 (0/0) | 0.1 | 89.2 | 2 |
| 2402 | RF01527 CrfA | 0.0 (0/45) | 0.0 | 0.0 (0/0) | 0.1 | 90.4 | 2 |
| 2403 | RF00611 SNORD111 | 0.0 (0/5) | 0.0 | 0.0 (0/0) | 2.0 | 75.5 | 5 |
| 2404 | RF02562 CbSR14 | 0.0 (0/33) | 0.0 | 0.0 (0/0) | 0.2 | 100.0 | 2 |
| 2405 | RF01621 ceN126 | 0.0 (0/35) | 0.0 | 0.0 (0/0) | 0.2 | 89.0 | 3 |
| 2406 | RF02445 SpR14 sRNA | 0.0 (0/17) | 0.0 | 0.0 (0/0) | 1.3 | 86.6 | 5 |
| 2407 | RF01499 Afu 191 | 0.0 (0/0) | 0.0 | 0.0 (0/0) | 0.0 | 76.1 | 7 |
| 2408 | RF02594 NsiR9 | 0.0 (0/41) | 0.0 | 0.0 (0/0) | 0.7 | 86.3 | 3 |
| 2409 | RF01144 sR17 | 0.0 (0/0) | 0.0 | 0.0 (0/0) | 0.0 | 69.8 | 5 |
| 2410 | RF01545 TB3Cs2H1 | 0.0 (0/15) | 0.0 | 0.0 (0/0) | 0.3 | 77.6 | 2 |
| 2411 | RF00312 snoZ206 | 0.0 (0/5) | 0.0 | 0.0 (0/0) | 1.0 | 89.8 | 6 |
| 2412 | RF01121 sR38 | 0.0 (0/0) | 0.0 | 0.0 (0/0) | 0.0 | 93.8 | 3 |
| 2413 | RF02045 CDKN2B-AS 3 | 0.0 (0/0) | 0.0 | 0.0 (0/0) | 0.0 | 70.6 | 18 |
| 2414 | RF01015 mir-885 | 0.0 (0/30) | 0.0 | 0.0 (0/0) | 0.9 | 77.5 | 3 |
| 2415 | RF02180 ST7-AS1 2 | 0.0 (0/0) | 0.0 | 0.0 (0/0) | 0.0 | 76.8 | 24 |
| 2416 | RF01327 CRISPR-DR14 | 0.0 (0/0) | 0.0 | 0.0 (0/0) | 0.0 | 89.0 | 5 |
| 2417 | RF00818 mir-318 | 0.0 (0/26) | 0.0 | 0.0 (0/0) | 0.5 | 89.1 | 5 |
| 2418 | RF01161 SNORD5 | 0.0 (0/0) | 0.0 | 0.0 (0/0) | 0.0 | 81.9 | 20 |
| 2419 | RF01862 TB10Cs4H2 | 0.0 (0/16) | 0.0 | 0.0 (0/2) | 2.2 | 66.9 | 4 |
| 2420 | RF02599 BASRCI408 | 0.0 (0/133) | 0.0 | 0.0 (0/0) | 0.4 | 98.6 | 3 |
| 2421 | RF02776 SodC thermometer | 0.0 (0/18) | 0.0 | 0.0 (0/0) | 0.0 | 98.9 | 2 |
| 2422 | RF01939 mir-761 | 0.0 (0/26) | 0.0 | 0.0 (0/0) | 0.0 | 98.5 | 4 |
| 2423 | RF02285 TtnuCD8 | 0.0 (0/0) | 0.0 | 0.0 (0/0) | 0.0 | 87.9 | 2 |
| 2424 | RF02738 Rev24 | 0.0 (0/140) | 0.0 | 0.0 (0/0) | 1.0 | 82.3 | 4 |
| 2425 | RF02575 DM SisR1 | 0.0 (0/107) | 0.0 | 0.0 (0/0) | 0.3 | 87.6 | 3 |

Continued on next page

| | RNA family (seed alignment) | Sensitivity annotated bpairs that covary % (cov_bps/bps) | Power average power % | Positive Predictive Value covarying pairs in structure % (cov_bps/cov_pairs) | average substitutions per bpair | avg pairwise identity % | number of sequences |
|---|---|---|---|---|---|---|---|
| 2426 | RF01162 sn668 | 0.0 (0/4) | 0.0 | 0.0 (0/0) | 0.0 | 94.4 | 6 |
| 2427 | RF02289 TtnuCD13 | 0.0 (0/0) | 0.0 | 0.0 (0/0) | 0.0 | 98.4 | 2 |
| 2428 | RF01922 mir-654 | 0.0 (0/26) | 0.0 | 0.0 (0/0) | 0.3 | 79.7 | 6 |
| 2429 | RF02243 Xoo8 | 0.0 (0/81) | 0.0 | 0.0 (0/0) | 0.7 | 85.5 | 4 |
| 2430 | RF00314 snoZ182 | 0.0 (0/6) | 0.0 | 0.0 (0/0) | 0.5 | 96.1 | 6 |
| 2431 | RF02661 icaR 5p UTR | 0.0 (0/22) | 0.0 | 0.0 (0/0) | 0.0 | 98.7 | 2 |
| 2432 | RF01226 snoZ5 | 0.0 (0/2) | 0.0 | 0.0 (0/0) | 0.0 | 84.0 | 8 |
| 2433 | RF02522 Virus CITE 8 | 0.0 (0/21) | 0.0 | 0.0 (0/0) | 0.0 | 94.4 | 2 |
| 2434 | RF00322 SNORA31 | 0.0 (0/38) | 0.0 | 0.0 (0/0) | 1.6 | 85.4 | 5 |
| 2435 | RF02558 CbSR2 | 0.0 (0/80) | 0.0 | 0.0 (0/0) | 0.0 | 99.6 | 2 |
| 2436 | RF01384 InvR | 0.0 (0/17) | 0.0 | 0.0 (0/0) | 0.0 | 96.0 | 4 |
| 2437 | RF01371 CRISPR-DR58 | 0.0 (0/5) | 0.0 | 0.0 (0/0) | 0.2 | 94.6 | 2 |
| 2438 | RF01331 CRISPR-DR18 | 0.0 (0/5) | 0.0 | 0.0 (0/0) | 0.0 | 80.5 | 6 |
| 2439 | RF02302 TtnuCD26 | 0.0 (0/0) | 0.0 | 0.0 (0/0) | 0.0 | 86.7 | 2 |
| 2440 | RF02043 HOTTIP 4 | 0.0 (0/0) | 0.0 | 0.0 (0/0) | 0.0 | 81.6 | 19 |
| 2441 | RF02044 CDKN2B-AS 2 | 0.0 (0/0) | 0.0 | 0.0 (0/0) | 0.0 | 79.4 | 6 |
| 2442 | RF01328 CRISPR-DR17 | 0.0 (0/6) | 0.0 | 0.0 (0/0) | 0.0 | 94.7 | 3 |
| 2443 | RF01133 sR3 | 0.0 (0/0) | 0.0 | 0.0 (0/0) | 0.0 | 67.2 | 19 |
| 2444 | RF00914 mir-674 | 0.0 (0/32) | 0.0 | 0.0 (0/0) | 0.0 | 98.0 | 2 |
| 2445 | RF01167 sn2429 | 0.0 (0/4) | 0.0 | 0.0 (0/0) | 0.5 | 90.3 | 4 |
| 2446 | RF02387 sau-27 | 0.0 (0/27) | 0.0 | 0.0 (0/0) | 0.6 | 88.3 | 3 |
| 2447 | RF02755 ES222 | 0.0 (0/36) | 0.0 | 0.0 (0/0) | 0.2 | 96.3 | 5 |
| 2448 | RF00964 mir-938 | 0.0 (0/32) | 0.0 | 0.0 (0/0) | 0.1 | 89.2 | 2 |
| 2449 | RF01134 sR30 | 0.0 (0/0) | 0.0 | 0.0 (0/0) | 0.0 | 88.8 | 3 |
| 2450 | RF02633 Hrs21 | 0.0 (0/31) | 0.0 | 0.0 (0/0) | 0.0 | 94.8 | 2 |
| 2451 | RF01666 rox2 | 0.0 (0/18) | 0.0 | 0.0 (0/0) | 0.4 | 86.4 | 4 |
| 2452 | RF01374 CRISPR-DR61 | 0.0 (0/7) | 0.0 | 0.0 (0/0) | 0.0 | 94.6 | 2 |
| 2453 | RF02149 MESTIT1 2 | 0.0 (0/0) | 0.0 | 0.0 (0/0) | 0.0 | 84.7 | 16 |
| 2454 | RF02565 YrlB | 0.0 (0/24) | 0.0 | 0.0 (0/0) | 0.2 | 75.6 | 2 |
| 2455 | RF01521 CC1840 | 0.0 (0/25) | 0.0 | 0.0 (0/0) | 1.0 | 70.7 | 3 |
| 2456 | RF01597 snoR22 | 0.0 (0/4) | 0.0 | 0.0 (0/0) | 0.0 | 90.7 | 3 |
| 2457 | RF01912 mir-2807 | 0.0 (0/36) | 0.0 | 0.0 (0/0) | 0.9 | 85.8 | 7 |
| 2458 | RF02109 DLEU2 5 | 0.0 (0/0) | 0.0 | 0.0 (0/0) | 0.0 | 81.3 | 30 |
| 2459 | RF02308 TtnuCD34 | 0.0 (0/0) | 0.0 | 0.0 (0/0) | 0.0 | 93.8 | 2 |
| 2460 | RF00933 mir-875 | 0.0 (0/25) | 0.0 | 0.0 (0/1) | 0.9 | 90.1 | 9 |
| 2461 | RF02853 Ysr201 | 0.0 (0/19) | 0.0 | 0.0 (0/0) | 0.2 | 94.3 | 3 |
| 2462 | RF02505 Atu L6 | 0.0 (0/36) | 0.0 | 0.0 (0/0) | 0.3 | 82.7 | 3 |
| 2463 | RF02771 CnfY thermometer | 0.0 (0/26) | 0.0 | 0.0 (0/0) | 0.0 | 94.6 | 3 |
| 2464 | RF01106 SBWMV1 UPD-PKb | 0.0 (0/10) | 0.0 | 0.0 (0/0) | 0.0 | 92.0 | 2 |
| 2465 | RF02059 STnc50 | 0.0 (0/12) | 0.0 | 0.0 (0/0) | 0.0 | 90.2 | 2 |
| 2466 | RF01682 P8 | 0.0 (0/25) | 0.0 | 0.0 (0/0) | 0.1 | 92.3 | 2 |
| 2467 | RF01333 CRISPR-DR20 | 0.0 (0/8) | 0.0 | 0.0 (0/0) | 0.5 | 78.7 | 3 |
| 2468 | RF00962 mir-586 | 0.0 (0/37) | 0.0 | 0.0 (0/0) | 0.1 | 94.8 | 2 |
| 2469 | RF01765 srg1 | 0.0 (0/152) | 0.0 | 0.0 (0/0) | 0.1 | 87.7 | 2 |
| 2470 | RF00805 mir-351 | 0.0 (0/30) | 0.0 | 0.0 (0/0) | 0.0 | 91.8 | 2 |
| 2471 | RF01446 S pombe snR95 | 0.0 (0/60) | 0.0 | 0.0 (0/0) | 0.0 | 100.0 | 2 |
| 2472 | RF02211 ZFAT-AS1 1 | 0.0 (0/0) | 0.0 | 0.0 (0/0) | 0.0 | 96.6 | 3 |
| 2473 | RF02588 HCV package-SL6067 | 0.0 (0/9) | 0.0 | 0.0 (0/0) | 0.6 | 93.3 | 3 |
| 2474 | RF02336 GlsR1 miR6 | 0.0 (0/19) | 0.0 | 0.0 (0/0) | 0.1 | 88.2 | 2 |
| 2475 | RF02761 sR084 | 0.0 (0/18) | 0.0 | 0.0 (0/0) | 0.3 | 87.8 | 3 |
| 2476 | RF01361 CRISPR-DR48 | 0.0 (0/7) | 0.0 | 0.0 (0/0) | 0.0 | 97.3 | 2 |
| 2477 | RF00870 mir-423 | 0.0 (0/31) | 0.0 | 0.0 (0/0) | 1.2 | 88.8 | 4 |
| 2478 | RF01598 snoR23 | 0.0 (0/3) | 0.0 | 0.0 (0/0) | 0.7 | 85.5 | 3 |
| 2479 | RF02735 Sernc350 | 0.0 (0/172) | 0.0 | 0.0 (0/0) | 0.2 | 78.5 | 2 |
| 2480 | RF02491 Gl U1 | 0.0 (0/37) | 0.0 | 0.0 (0/0) | 0.1 | 89.2 | 2 |
| 2481 | RF02161 PART1 3 | 0.0 (0/0) | 0.0 | 0.0 (0/0) | 0.0 | 81.8 | 3 |
| 2482 | RF01362 CRISPR-DR49 | 0.0 (0/6) | 0.0 | 0.0 (0/0) | 0.5 | 90.1 | 3 |
| 2483 | RF01480 rli52 | 0.0 (0/26) | 0.0 | 0.0 (0/0) | 0.6 | 94.8 | 6 |
| 2484 | RF02071 STnc280 | 0.0 (0/11) | 0.0 | 0.0 (0/0) | 0.4 | 83.8 | 3 |
| 2485 | RF01937 mir-2780 | 0.0 (0/28) | 0.0 | 0.0 (0/0) | 0.3 | 96.2 | 4 |
| 2486 | RF01652 ceN70 | 0.0 (0/3) | 0.0 | 0.0 (0/0) | 0.0 | 90.3 | 3 |
| 2487 | RF01011 mir-605 | 0.0 (0/36) | 0.0 | 0.0 (0/0) | 0.0 | 94.0 | 2 |
| 2488 | RF01340 CRISPR-DR29 | 0.0 (0/9) | 0.0 | 0.0 (0/0) | 0.0 | 100.0 | 2 |
| 2489 | RF02813 PA5194 thermometer | 0.0 (0/26) | 0.0 | 0.0 (0/0) | 0.3 | 64.4 | 2 |
| 2490 | RF02572 babR 5UTR | 0.0 (0/37) | 0.0 | 0.0 (0/0) | 0.0 | 99.0 | 3 |
| 2491 | RF00313 snoZ173 | 0.0 (0/7) | 0.0 | 0.0 (0/0) | 0.0 | 99.4 | 4 |
| 2492 | RF01930 bxd 2 | 0.0 (0/0) | 0.0 | 0.0 (0/0) | 0.0 | 84.1 | 5 |
| 2493 | RF01891 TUG1 3 | 0.0 (0/0) | 0.0 | 0.0 (0/0) | 0.0 | 87.0 | 24 |
| 2494 | RF02665 PsiU1-6 | 0.0 (0/44) | 0.0 | 0.0 (0/0) | 0.1 | 94.5 | 3 |
| 2495 | RF01422 snoR116 | 0.0 (0/3) | 0.0 | 0.0 (0/0) | 0.3 | 74.4 | 5 |
| 2496 | RF02490 GlsR27 | 0.0 (0/31) | 0.0 | 0.0 (0/0) | 0.1 | 91.1 | 2 |
| 2497 | RF00743 mir-308 | 0.0 (0/24) | 0.0 | 0.0 (0/0) | 1.0 | 85.9 | 11 |
| 2498 | RF02108 DLEU2 4 | 0.0 (0/0) | 0.0 | 0.0 (0/0) | 0.0 | 80.2 | 21 |
| 2499 | RF01974 H19 3 | 0.0 (0/0) | 0.0 | 0.0 (0/0) | 0.0 | 94.1 | 5 |
| 2500 | RF01448 S pombe snR93 | 0.0 (0/33) | 0.0 | 0.0 (0/0) | 0.0 | 100.0 | 2 |
| 2501 | RF01991 SECIS 5 | 0.0 (0/20) | 0.0 | 0.0 (0/0) | 0.7 | 83.3 | 3 |
| 2502 | RF02160 PART1 2 | 0.0 (0/0) | 0.0 | 0.0 (0/0) | 0.0 | 78.3 | 29 |
| 2503 | RF01339 CRISPR-DR27 | 0.0 (0/4) | 0.0 | 0.0 (0/0) | 0.0 | 94.7 | 3 |
| 2504 | RF00719 mir-326 | 0.0 (0/35) | 0.0 | 0.0 (0/0) | 0.9 | 92.4 | 7 |
| 2505 | RF01150 sR11 | 0.0 (0/0) | 0.0 | 0.0 (0/0) | 0.0 | 57.8 | 8 |
| 2506 | RF01398 isrP | 0.0 (0/44) | 0.0 | 0.0 (0/0) | 0.3 | 96.8 | 7 |
| 2507 | RF02011 mir-575 | 0.0 (0/31) | 0.0 | 0.0 (0/0) | 0.0 | 99.3 | 3 |
| 2508 | RF01463 rli27 | 0.0 (0/10) | 0.0 | 0.0 (0/0) | 0.0 | 93.9 | 3 |
| 2509 | RF02564 naRNA4 | 0.0 (0/24) | 0.0 | 0.0 (0/0) | 0.0 | 99.1 | 3 |

Continued on next page

| | RNA family (seed alignment) | Sensitivity annotated bpairs that covary % (cov_bps/bps) | Power average power % | Positive Predictive Value covarying pairs in structure % (cov_bps/cov_pairs) | average substitutions per bpair | avg pairwise identity % | number of sequences |
|---|---|---|---|---|---|---|---|
| 2510 | RF02475 GlsR6 | 0.0 (0/0) | 0.0 | 0.0 (0/0) | 0.0 | 89.8 | 3 |
| 2511 | RF02805 PyrR207 | 0.0 (0/18) | 0.0 | 0.0 (0/0) | 1.1 | 85.1 | 4 |
| 2512 | RF02723 sno ZL1 | 0.0 (0/60) | 0.0 | 0.0 (0/0) | 0.5 | 89.3 | 5 |
| 2513 | RF02511 PYLIS 4 | 0.0 (0/13) | 0.0 | 0.0 (0/0) | 0.2 | 81.0 | 2 |
| 2514 | RF02517 mir-1017 | 0.0 (0/17) | 0.0 | 0.0 (0/0) | 0.5 | 88.1 | 7 |
| 2515 | RF00178 mir-24 | 0.0 (0/16) | 0.0 | 0.0 (0/0) | 1.2 | 79.6 | 17 |
| 2516 | RF02187 ST7-OT4 1 | 0.0 (0/0) | 0.0 | 0.0 (0/0) | 0.0 | 73.6 | 24 |
| 2517 | RF01778 Evf-2 5p | 0.0 (0/0) | 0.0 | 0.0 (0/0) | 0.0 | 86.5 | 13 |
| 2518 | RF01155 snoR101 | 0.0 (0/0) | 0.0 | 0.0 (0/0) | 0.0 | 83.6 | 5 |
| 2519 | RF01638 ceN48 | 0.0 (0/32) | 0.0 | 0.0 (0/0) | 0.5 | 90.6 | 5 |
| 2520 | RF02444 SpR10 sRNA | 0.0 (0/11) | 0.0 | 0.0 (0/0) | 0.6 | 91.8 | 5 |
| 2521 | RF01391 isrH | 0.0 (0/87) | 0.0 | 0.0 (0/0) | 0.3 | 97.2 | 6 |
| 2522 | RF01573 DdR5 | 0.0 (0/0) | 0.0 | 0.0 (0/0) | 0.0 | 95.1 | 3 |
| 2523 | RF02022 mir-1275 | 0.0 (0/24) | 0.0 | 0.0 (0/0) | 0.9 | 90.4 | 6 |
| 2524 | RF02198 TTC28-AS1 1 | 0.0 (0/0) | 0.0 | 0.0 (0/0) | 0.0 | 67.2 | 14 |
| 2525 | RF02536 AEV CRE | 0.0 (0/43) | 0.0 | 0.0 (0/0) | 0.0 | 97.8 | 2 |
| 2526 | RF00348 snoR9 plant | 0.0 (0/3) | 0.0 | 0.0 (0/0) | 0.0 | 82.5 | 7 |
| 2527 | RF00412 SNORA21 | 0.0 (0/24) | 0.0 | 0.0 (0/0) | 1.0 | 87.9 | 5 |
| 2528 | RF01153 sn3060 | 0.0 (0/5) | 0.0 | 0.0 (0/0) | 0.2 | 98.0 | 3 |
| 2529 | RF01569 DdR2 | 0.0 (0/0) | 0.0 | 0.0 (0/0) | 0.0 | 84.6 | 3 |
| 2530 | RF01472 rli40 | 0.0 (0/81) | 0.0 | 0.0 (0/2) | 0.7 | 86.7 | 4 |
| 2531 | RF02294 TtnuCD18 | 0.0 (0/0) | 0.0 | 0.0 (0/0) | 0.0 | 87.5 | 2 |
| 2532 | RF02559 CbSR4 | 0.0 (0/52) | 0.0 | 0.0 (0/0) | 0.0 | 99.5 | 2 |
| 2533 | RF02707 TeloSII ncR33 | 0.0 (0/68) | 0.0 | 0.0 (0/0) | 0.3 | 92.4 | 3 |
| 2534 | RF01559 DdR1 | 0.0 (0/0) | 0.0 | 0.0 (0/0) | 0.0 | 84.8 | 3 |
| 2535 | RF01971 RMST 10 | 0.0 (0/0) | 0.0 | 0.0 (0/0) | 0.0 | 84.2 | 21 |
| 2536 | RF00642 mir-23 | 0.0 (0/14) | 0.0 | 0.0 (0/0) | 1.3 | 70.6 | 22 |
| 2537 | RF00038 PrfA | 0.0 (0/14) | 0.0 | 0.0 (0/0) | 0.9 | 87.4 | 11 |
| 2538 | RF00101 SraC RyeA | 0.0 (0/23) | 0.0 | 0.0 (0/1) | 0.3 | 86.2 | 13 |
| 2539 | RF02080 STnc170 | 0.0 (0/77) | 0.0 | 0.0 (0/0) | 0.0 | 97.3 | 3 |
| 2540 | RF01207 snR73 | 0.0 (0/0) | 0.0 | 0.0 (0/0) | 0.0 | 80.5 | 5 |
| 2541 | RF01458 rli23 | 0.0 (0/31) | 0.0 | 0.0 (0/0) | 1.7 | 78.9 | 7 |
| 2542 | RF01171 snoR4 | 0.0 (0/0) | 0.0 | 0.0 (0/0) | 0.0 | 61.5 | 2 |
| 2543 | RF02583 Teg23 | 0.0 (0/124) | 0.0 | 0.0 (0/0) | 0.2 | 97.6 | 6 |
| 2544 | RF01785 blv FSE | 0.0 (0/9) | 0.0 | 0.0 (0/0) | 0.0 | 88.9 | 2 |
| 2545 | RF01642 ceN54 | 0.0 (0/2) | 0.0 | 0.0 (0/0) | 0.0 | 91.7 | 3 |
| 2546 | RF01439 S pombe snR36 | 0.0 (0/41) | 0.0 | 0.0 (0/0) | 0.0 | 100.0 | 2 |
| 2547 | RF01147 sR12 | 0.0 (0/0) | 0.0 | 0.0 (0/0) | 0.0 | 69.2 | 5 |
| 2548 | RF02579 tsr26 | 0.0 (0/33) | 0.0 | 0.0 (0/0) | 0.0 | 100.0 | 2 |
| 2549 | RF00498 EAV LTH | 0.0 (0/11) | 0.0 | 0.0 (0/0) | 0.0 | 98.1 | 6 |
| 2550 | RF02408 snoR124 | 0.0 (0/4) | 0.0 | 0.0 (0/0) | 0.2 | 90.8 | 5 |
| 2551 | RF02624 BSnc150 | 0.0 (0/7) | 0.0 | 0.0 (0/0) | 0.0 | 95.5 | 2 |
| 2552 | RF00883 MIR820 | 0.0 (0/76) | 0.0 | 0.0 (0/0) | 0.1 | 93.4 | 2 |
| 2553 | RF02758 RhlA thermometer | 0.0 (0/36) | 0.0 | 0.0 (0/0) | 0.0 | 98.8 | 4 |
| 2554 | RF01447 S pombe snR98 | 0.0 (0/59) | 0.0 | 0.0 (0/0) | 0.0 | 99.5 | 2 |
| 2555 | RF02311 TtnuHACA3 | 0.0 (0/38) | 0.0 | 0.0 (0/0) | 0.1 | 91.5 | 2 |
| 2556 | RF01890 lincRNA-p21 2 | 0.0 (0/0) | 0.0 | 0.0 (0/0) | 0.0 | 96.2 | 2 |
| 2557 | RF01631 ceN41 | 0.0 (0/34) | 0.0 | 0.0 (0/0) | 0.6 | 78.4 | 3 |
| 2558 | RF02210 WT1-AS 8 | 0.0 (0/0) | 0.0 | 0.0 (0/0) | 0.0 | 79.2 | 24 |
| 2559 | RF02272 Vax2os1 3 | 0.0 (0/0) | 0.0 | 0.0 (0/0) | 0.0 | 70.1 | 5 |
| 2560 | RF01354 CRISPR-DR45 | 0.0 (0/6) | 0.0 | 0.0 (0/0) | 0.0 | 94.4 | 3 |
| 2561 | RF02597 shuA thermo | 0.0 (0/7) | 0.0 | 0.0 (0/0) | 0.0 | 100.0 | 5 |
| 2562 | RF00454 p27 CRE | 0.0 (0/14) | 0.0 | 0.0 (0/0) | 1.1 | 93.2 | 12 |
| 2563 | RF00117 C0719 | 0.0 (0/32) | 0.0 | 0.0 (0/0) | 0.2 | 98.5 | 9 |
| 2564 | RF02907 patAB | 0.0 (0/13) | 0.0 | 0.0 (0/0) | 0.0 | 98.2 | 2 |
| 2565 | RF00940 mir-327 | 0.0 (0/20) | 0.0 | 0.0 (0/0) | 0.1 | 86.9 | 2 |
| 2566 | RF02739 Rev41 | 0.0 (0/156) | 0.0 | 0.0 (0/0) | 0.3 | 90.6 | 3 |
| 2567 | RF02365 Yfr17 | 0.0 (0/74) | 0.0 | 0.0 (0/1) | 0.3 | 85.5 | 3 |
| 2568 | RF02331 TtnuHACA24 | 0.0 (0/37) | 0.0 | 0.0 (0/0) | 0.0 | 92.9 | 2 |
| 2569 | RF01243 snR33 | 0.0 (0/39) | 0.0 | 0.0 (0/0) | 0.5 | 90.5 | 5 |
| 2570 | RF02610 BSR0709 | 0.0 (0/96) | 0.0 | 0.0 (0/0) | 0.0 | 99.8 | 4 |
| 2571 | RF01420 snoR113 | 0.0 (0/3) | 0.0 | 0.0 (0/0) | 0.0 | 72.4 | 5 |
| 2572 | RF02719 sca ncR20 | 0.0 (0/30) | 0.0 | 0.0 (0/0) | 0.0 | 100.0 | 2 |
| 2573 | RF02856 30 292 | 0.0 (0/30) | 0.0 | 0.0 (0/0) | 0.6 | 88.6 | 3 |
| 2574 | RF00849 mir-60 | 0.0 (0/27) | 0.0 | 0.0 (0/0) | 0.2 | 88.7 | 4 |
| 2575 | RF01947 KCNQ1OT1 2 | 0.0 (0/0) | 0.0 | 0.0 (0/0) | 0.0 | 83.6 | 18 |
| 2576 | RF02143 HYMAI | 0.0 (0/0) | 0.0 | 0.0 (0/0) | 0.0 | 74.5 | 16 |
| 2577 | RF02554 ppoRNA | 0.0 (0/36) | 0.0 | 0.0 (0/0) | 0.0 | 99.0 | 2 |
| 2578 | RF02487 GlsR24 | 0.0 (0/35) | 0.0 | 0.0 (0/0) | 0.4 | 83.8 | 3 |
| 2579 | RF00815 mir-244 | 0.0 (0/38) | 0.0 | 0.0 (0/0) | 0.7 | 78.3 | 4 |
| 2580 | RF00863 mir-BART17 | 0.0 (0/30) | 0.0 | 0.0 (0/0) | 0.2 | 72.5 | 2 |
| 2581 | RF00910 mir-873 | 0.0 (0/25) | 0.0 | 0.0 (0/0) | 0.2 | 96.5 | 3 |
| 2582 | RF01156 SNORD123 | 0.0 (0/6) | 0.0 | 0.0 (0/0) | 1.0 | 87.9 | 12 |
| 2583 | RF00088 SNORD30 | 0.0 (0/9) | 0.0 | 0.0 (0/0) | 3.0 | 82.0 | 20 |
| 2584 | RF02334 TtnuHACA28 | 0.0 (0/33) | 0.0 | 0.0 (0/0) | 0.0 | 95.5 | 2 |
| 2585 | RF01004 mir-557 | 0.0 (0/23) | 0.0 | 0.0 (0/0) | 0.0 | 93.0 | 2 |
| 2586 | RF01490 rli51 | 0.0 (0/31) | 0.0 | 0.0 (0/0) | 0.9 | 92.6 | 9 |
| 2587 | RF00476 snosnR61 | 0.0 (0/0) | 0.0 | 0.0 (0/0) | 0.0 | 68.7 | 56 |
| 2588 | RF01931 bxd 3 | 0.0 (0/0) | 0.0 | 0.0 (0/1) | 0.0 | 82.5 | 4 |
| 2589 | RF01013 mir-577 | 0.0 (0/36) | 0.0 | 0.0 (0/0) | 0.1 | 95.8 | 2 |
| 2590 | RF01035 mir-887 | 0.0 (0/23) | 0.0 | 0.0 (0/0) | 0.4 | 78.1 | 3 |
| 2591 | RF01146 sR15 | 0.0 (0/0) | 0.0 | 0.0 (0/0) | 0.0 | 52.7 | 2 |
| 2592 | RF00701 mir-126 | 0.0 (0/21) | 0.0 | 0.0 (0/0) | 0.3 | 86.3 | 12 |
| 2593 | RF00310 snoZ165 | 0.0 (0/3) | 0.0 | 0.0 (0/0) | 1.7 | 94.8 | 5 |

Continued on next page

| | RNA family (seed alignment) | Sensitivity annotated bpairs that covary % (cov_bps/bps) | Power average power % | Positive Predictive Value covarying pairs in structure % (cov_bps/cov_pairs) | average substitutions per bpair | avg pairwise identity % | number of sequences |
|---|---|---|---|---|---|---|---|
| 2594 | RF02134 HTT-AS1 1 | 0.0 (0/0) | 0.0 | 0.0 (0/0) | 0.0 | 90.8 | 3 |
| 2595 | RF02430 SpF19 sRNA | 0.0 (0/36) | 0.0 | 0.0 (0/0) | 0.9 | 79.4 | 3 |
| 2596 | RF02908 rli30 | 0.0 (0/38) | 0.0 | 0.0 (0/1) | 0.9 | 75.9 | 3 |
| 2597 | RF01640 ceN51 | 0.0 (0/39) | 0.0 | 0.0 (0/0) | 0.2 | 89.2 | 3 |
| 2598 | RF00666 mir-32 | 0.0 (0/30) | 0.0 | 0.0 (0/0) | 1.3 | 90.3 | 8 |
| 2599 | RF00184 PVX 3 | 0.0 (0/8) | 0.0 | 0.0 (0/0) | 0.0 | 96.0 | 4 |
| 2600 | RF01577 RNase P | 0.0 (0/91) | 0.0 | 0.0 (0/0) | 0.0 | 99.7 | 2 |
| 2601 | RF02463 Ms AS-1 | 0.0 (0/21) | 0.0 | 0.0 (0/0) | 1.7 | 80.0 | 5 |
| 2602 | RF01529 CC3552 | 0.0 (0/28) | 0.0 | 0.0 (0/0) | 0.1 | 71.2 | 2 |
| 2603 | RF02318 TtnuHACA11a | 0.0 (0/36) | 0.0 | 0.0 (0/0) | 0.0 | 94.3 | 2 |
| 2604 | RF01680 P5 | 0.0 (0/20) | 0.0 | 0.0 (0/0) | 0.0 | 93.0 | 2 |
| 2605 | RF02655 BSR0441 | 0.0 (0/83) | 0.0 | 0.0 (0/0) | 0.0 | 99.2 | 5 |
| 2606 | RF02105 DLEU2 1 | 0.0 (0/0) | 0.0 | 0.0 (0/0) | 0.0 | 79.9 | 27 |
| 2607 | RF01404 PinT | 0.0 (0/15) | 0.0 | 0.0 (0/0) | 0.1 | 99.3 | 4 |
| 2608 | RF00819 mir-246 | 0.0 (0/33) | 0.0 | 0.0 (0/1) | 1.3 | 65.4 | 4 |
| 2609 | RF01951 SOX2OT exon1 | 0.0 (0/0) | 0.0 | 0.0 (0/0) | 0.0 | 83.4 | 29 |
| 2610 | RF02215 ZNFX1-AS1 1 | 0.0 (0/0) | 0.0 | 0.0 (0/0) | 0.0 | 69.1 | 15 |
| 2611 | RF02750 ES003 | 0.0 (0/15) | 0.0 | 0.0 (0/0) | 0.4 | 94.7 | 8 |
| 2612 | RF02286 TtnuCD9 | 0.0 (0/0) | 0.0 | 0.0 (0/0) | 0.0 | 96.4 | 2 |
| 2613 | RF02504 Atu L1 | 0.0 (0/34) | 0.0 | 0.0 (0/0) | 0.2 | 92.5 | 4 |
| 2614 | RF01952 SOX2OT exon2 | 0.0 (0/0) | 0.0 | 0.0 (0/0) | 0.0 | 86.0 | 14 |
| 2615 | RF02702 AgvB | 0.0 (0/17) | 0.0 | 0.0 (0/0) | 0.1 | 94.3 | 3 |
| 2616 | RF02604 BASRCI153 | 0.0 (0/73) | 0.0 | 0.0 (0/0) | 0.1 | 92.1 | 2 |
| 2617 | RF02304 TtnuCD29 | 0.0 (0/0) | 0.0 | 0.0 (0/0) | 0.0 | 98.7 | 2 |
| 2618 | RF01935 bxd 7 | 0.0 (0/0) | 0.0 | 0.0 (0/0) | 0.0 | 92.4 | 3 |
| 2619 | RF02455 Virus CITE 1 | 0.0 (0/27) | 0.0 | 0.0 (0/0) | 0.4 | 80.7 | 2 |
| 2620 | RF00966 mir-676 | 0.0 (0/30) | 0.0 | 0.0 (0/0) | 0.6 | 87.7 | 6 |
| 2621 | RF02171 RFPL3-AS1 1 | 0.0 (0/0) | 0.0 | 0.0 (0/0) | 0.0 | 90.7 | 4 |
| 2622 | RF01533 TB10Cs4H3 | 0.0 (0/17) | 0.0 | 0.0 (0/0) | 1.8 | 71.8 | 3 |
| 2623 | RF01335 CRISPR-DR22 | 0.0 (0/7) | 0.0 | 0.0 (0/0) | 2.3 | 79.5 | 9 |
| 2624 | RF02585 HCV package | 0.0 (0/8) | 0.0 | 0.0 (0/0) | 0.1 | 90.8 | 3 |
| 2625 | RF00802 mir-207 | 0.0 (0/31) | 0.0 | 0.0 (0/0) | 1.3 | 83.5 | 6 |
| 2626 | RF01314 mir-1227 | 0.0 (0/29) | 0.0 | 0.0 (0/0) | 0.1 | 95.5 | 3 |
| 2627 | RF00851 mir-230 | 0.0 (0/37) | 0.0 | 0.0 (0/0) | 0.1 | 72.9 | 2 |
| 2628 | RF00292 snoTBR5 | 0.0 (0/2) | 0.0 | 0.0 (0/0) | 0.0 | 81.0 | 6 |
| 2629 | RF01917 MIR2587 | 0.0 (0/35) | 0.0 | 0.0 (0/0) | 0.2 | 94.1 | 4 |
| 2630 | RF02300 TtnuCD24 | 0.0 (0/0) | 0.0 | 0.0 (0/0) | 0.0 | 98.3 | 2 |
| 2631 | RF01583 snoR01 | 0.0 (0/2) | 0.0 | 0.0 (0/0) | 0.0 | 77.6 | 3 |
| 2632 | RF02246 Six3os1 1 | 0.0 (0/0) | 0.0 | 0.0 (0/0) | 0.0 | 80.4 | 9 |
| 2633 | RF00342 snoZ40 | 0.0 (0/5) | 0.0 | 0.0 (0/0) | 0.0 | 94.4 | 5 |
| 2634 | RF01139 sR2 | 0.0 (0/0) | 0.0 | 0.0 (0/0) | 0.0 | 61.7 | 8 |
| 2635 | RF02862 sRNA 1300 | 0.0 (0/77) | 0.0 | 0.0 (0/0) | 0.1 | 95.2 | 3 |
| 2636 | RF01834 fiv FSE | 0.0 (0/11) | 0.0 | 0.0 (0/0) | 0.0 | 93.1 | 5 |
| 2637 | RF02147 MEG8 3 | 0.0 (0/0) | 0.0 | 0.0 (0/0) | 0.0 | 77.6 | 10 |
| 2638 | RF01368 CRISPR-DR55 | 0.0 (0/0) | 0.0 | 0.0 (0/0) | 0.0 | 86.5 | 2 |
| 2639 | RF02676 Esr41 | 0.0 (0/18) | 0.0 | 0.0 (0/0) | 0.1 | 98.2 | 3 |
| 2640 | RF02620 BSnc119 | 0.0 (0/19) | 0.0 | 0.0 (0/0) | 0.1 | 97.0 | 3 |
| 2641 | RF01636 ceN46 | 0.0 (0/30) | 0.0 | 0.0 (0/0) | 0.3 | 90.9 | 3 |
| 2642 | RF02481 GlsR18 | 0.0 (0/30) | 0.0 | 0.0 (0/0) | 0.5 | 84.5 | 3 |
| 2643 | RF01076 RF site2 | 0.0 (0/19) | 0.0 | 0.0 (0/0) | 0.1 | 90.3 | 2 |
| 2644 | RF02529 SSRC8 1 | 0.0 (0/38) | 0.0 | 0.0 (0/0) | 0.2 | 94.7 | 3 |
| 2645 | RF02698 Avalong thermometer | 0.0 (0/36) | 0.0 | 0.0 (0/0) | 0.1 | 86.3 | 2 |
| 2646 | RF02634 EF3314 EF3315 | 0.0 (0/45) | 0.0 | 0.0 (0/0) | 0.0 | 99.3 | 2 |
| 2647 | RF00318 snoZ175 | 0.0 (0/3) | 0.0 | 0.0 (0/0) | 0.0 | 99.1 | 6 |
| 2648 | RF00972 mir-651 | 0.0 (0/34) | 0.0 | 0.0 (0/0) | 0.8 | 81.9 | 4 |
| 2649 | RF02407 snoR122 | 0.0 (0/0) | 0.0 | 0.0 (0/0) | 0.0 | 88.8 | 4 |
| 2650 | RF00866 mir-BART3 | 0.0 (0/29) | 0.0 | 0.0 (0/0) | 0.2 | 81.3 | 2 |
| 2651 | RF00901 MIR845 1 | 0.0 (0/31) | 0.0 | 0.0 (0/0) | 0.1 | 81.2 | 2 |
| 2652 | RF00189 SNORD95 | 0.0 (0/4) | 0.0 | 0.0 (0/0) | 0.8 | 90.2 | 7 |
| 2653 | RF02881 MH s25 | 0.0 (0/36) | 0.0 | 0.0 (0/0) | 0.4 | 67.2 | 2 |
| 2654 | RF01501 Afu 199 | 0.0 (0/0) | 0.0 | 0.0 (0/0) | 0.0 | 71.2 | 3 |
| 2655 | RF02785 snoRNA2 | 0.0 (0/21) | 0.0 | 0.0 (0/0) | 0.6 | 89.6 | 3 |
| 2656 | RF01619 ceN115 | 0.0 (0/40) | 0.0 | 0.0 (0/0) | 0.0 | 100.0 | 2 |
| 2657 | RF02550 RnaG | 0.0 (0/28) | 0.0 | 0.0 (0/0) | 0.0 | 99.0 | 4 |
| 2658 | RF02118 FMR1-AS1 2 | 0.0 (0/0) | 0.0 | 0.0 (0/0) | 0.0 | 79.6 | 25 |
| 2659 | RF01174 snoU43 | 0.0 (0/0) | 0.0 | 0.0 (0/0) | 0.0 | 75.8 | 11 |
| 2660 | RF02146 MEG8 2 | 0.0 (0/0) | 0.0 | 0.0 (0/1) | 0.0 | 73.0 | 12 |
| 2661 | RF01273 sR34 | 0.0 (0/9) | 0.0 | 0.0 (0/0) | 0.0 | 92.6 | 4 |
| 2662 | RF00289 SNORND104 | 0.0 (0/2) | 0.0 | 0.0 (0/0) | 1.0 | 85.6 | 18 |
| 2663 | RF02857 30 255 | 0.0 (0/46) | 0.0 | 0.0 (0/0) | 0.1 | 92.9 | 2 |
| 2664 | RF00462 IRES APC | 0.0 (0/11) | 0.0 | 0.0 (0/0) | 1.6 | 85.4 | 6 |
| 2665 | RF00472 snosnR55 | 0.0 (0/0) | 0.0 | 0.0 (0/0) | 0.0 | 74.8 | 6 |
| 2666 | RF01965 RMST 4 | 0.0 (0/0) | 0.0 | 0.0 (0/0) | 0.0 | 84.4 | 17 |
| 2667 | RF02212 ZFAT-AS1 2 | 0.0 (0/0) | 0.0 | 0.0 (0/0) | 0.0 | 69.0 | 11 |
| 2668 | RF00355 snoR28 | 0.0 (0/3) | 0.0 | 0.0 (0/1) | 1.7 | 81.6 | 9 |
| 2669 | RF01006 mir-601 | 0.0 (0/20) | 0.0 | 0.0 (0/0) | 0.1 | 96.6 | 3 |
| 2670 | RF02116 FAS-AS1 | 0.0 (0/0) | 0.0 | 0.0 (0/0) | 0.0 | 68.0 | 19 |
| 2671 | RF00796 mir-48 | 0.0 (0/30) | 0.0 | 0.0 (0/0) | 1.0 | 70.3 | 4 |
| 2672 | RF00574 SNORD69 | 0.0 (0/10) | 0.0 | 0.0 (0/0) | 2.3 | 88.1 | 11 |
| 2673 | RF02322 TtnuHACA15 | 0.0 (0/32) | 0.0 | 0.0 (0/0) | 0.0 | 96.6 | 2 |
| 2674 | RF02651 SorY | 0.0 (0/24) | 0.0 | 0.0 (0/0) | 0.0 | 97.6 | 2 |
| 2675 | RF00592 SNORD78 | 0.0 (0/0) | 0.0 | 0.0 (0/0) | 0.0 | 82.1 | 19 |
| 2676 | RF02038 SPRY4-IT1 1 | 0.0 (0/0) | 0.0 | 0.0 (0/1) | 0.0 | 78.5 | 23 |
| 2677 | RF00687 mir-136 | 0.0 (0/32) | 0.0 | 0.0 (0/0) | 0.0 | 93.9 | 2 |

Continued on next page

| | RNA family (seed alignment) | Sensitivity annotated bpairs that covary % (cov_bps/bps) | Power average power % | Positive Predictive Value covarying pairs in structure % (cov_bps/cov_pairs) | average substitutions per bpair | avg pairwise identity % | number of sequences |
|---|---|---|---|---|---|---|---|
| 2678 | RF02174 SMAD5-AS1 2 | 0.0 (0/0) | 0.0 | 0.0 (0/0) | 0.0 | 74.6 | 20 |
| 2679 | RF02169 PVT1 6 | 0.0 (0/0) | 0.0 | 0.0 (0/0) | 0.0 | 71.2 | 7 |
| 2680 | RF01485 rli61 | 0.0 (0/32) | 0.0 | 0.0 (0/0) | 0.0 | 99.5 | 4 |
| 2681 | RF00315 snoJ33 | 0.0 (0/6) | 0.0 | 0.0 (0/0) | 2.3 | 79.5 | 5 |
| 2682 | RF02115 FAM13A-AS1 2 | 0.0 (0/0) | 0.0 | 0.0 (0/0) | 0.0 | 69.3 | 10 |
| 2683 | RF01160 sn2841 | 0.0 (0/4) | 0.0 | 0.0 (0/0) | 0.0 | 95.8 | 2 |
| 2684 | RF02151 MIMT1 1 | 0.0 (0/0) | 0.0 | 0.0 (0/0) | 0.0 | 69.7 | 21 |
| 2685 | RF02388 sau-30 | 0.0 (0/12) | 0.0 | 0.0 (0/0) | 0.1 | 85.9 | 2 |
| 2686 | RF02456 Diantho 3 UTR | 0.0 (0/32) | 0.0 | 0.0 (0/0) | 0.2 | 81.2 | 2 |
| 2687 | RF02672 SprX | 0.0 (0/40) | 0.0 | 0.0 (0/0) | 0.0 | 98.7 | 3 |
| 2688 | RF01358 CRISPR-DR39 | 0.0 (0/4) | 0.0 | 0.0 (0/0) | 1.5 | 81.7 | 3 |
| 2689 | RF02753 ES173 | 0.0 (0/13) | 0.0 | 0.0 (0/0) | 0.0 | 95.0 | 4 |
| 2690 | RF01525 TB10Cs2H1 | 0.0 (0/10) | 0.0 | 0.0 (0/2) | 0.1 | 71.7 | 4 |
| 2691 | RF02066 STnc320 | 0.0 (0/12) | 0.0 | 0.0 (0/0) | 0.0 | 93.3 | 3 |
| 2692 | RF00275 SNORD56 | 0.0 (0/7) | 0.0 | 0.0 (0/0) | 2.4 | 80.0 | 7 |
| 2693 | RF00864 mir-BART20 | 0.0 (0/28) | 0.0 | 0.0 (0/0) | 0.1 | 81.2 | 2 |
| 2694 | RF01454 RCNMV 5UTR | 0.0 (0/23) | 0.0 | 0.0 (0/0) | 0.9 | 81.8 | 3 |
| 2695 | RF02154 NCRUPAR 1 | 0.0 (0/0) | 0.0 | 0.0 (0/0) | 0.0 | 79.3 | 9 |
| 2696 | RF02700 HtrA thermometer | 0.0 (0/12) | 0.0 | 0.0 (0/0) | 0.2 | 93.8 | 4 |
| 2697 | RF02477 GlsR9 | 0.0 (0/4) | 0.0 | 0.0 (0/0) | 0.0 | 93.3 | 2 |
| 2698 | RF02358 hsp17 | 0.0 (0/12) | 0.0 | 0.0 (0/0) | 0.8 | 73.3 | 3 |
| 2699 | RF02122 FTX 5 | 0.0 (0/0) | 0.0 | 0.0 (0/0) | 0.0 | 71.4 | 14 |
| 2700 | RF01783 b55 | 0.0 (0/22) | 0.0 | 0.0 (0/0) | 0.0 | 100.0 | 2 |
| 2701 | RF00330 snoZ43 | 0.0 (0/7) | 0.0 | 0.0 (0/0) | 1.0 | 83.0 | 7 |
| 2702 | RF02359 MS2 | 0.0 (0/7) | 0.0 | 0.0 (0/0) | 0.0 | 88.5 | 5 |
| 2703 | RF01894 PCA3 2 | 0.0 (0/0) | 0.0 | 0.0 (0/0) | 0.0 | 89.6 | 20 |
| 2704 | RF02070 STnc300 | 0.0 (0/33) | 0.0 | 0.0 (0/0) | 0.4 | 84.6 | 4 |
| 2705 | RF00218 SNORD46 | 0.0 (0/6) | 0.0 | 0.0 (0/0) | 2.7 | 90.9 | 5 |
| 2706 | RF01369 CRISPR-DR56 | 0.0 (0/9) | 0.0 | 0.0 (0/0) | 0.0 | 94.6 | 2 |
| 2707 | RF00244 mir-26 | 0.0 (0/18) | 0.0 | 0.0 (0/0) | 0.5 | 74.4 | 5 |
| 2708 | RF02103 DLEU1 1 | 0.0 (0/0) | 0.0 | 0.0 (0/0) | 0.0 | 72.4 | 26 |
| 2709 | RF02827 Scr6106 | 0.0 (0/32) | 0.0 | 0.0 (0/0) | 0.5 | 90.0 | 4 |
| 2710 | RF00338 snoR53 | 0.0 (0/11) | 0.0 | 0.0 (0/0) | 1.1 | 94.3 | 5 |
| 2711 | RF01104 SBWMV2 UPD-PKb | 0.0 (0/10) | 0.0 | 0.0 (0/0) | 0.2 | 90.8 | 3 |
| 2712 | RF01628 ceN38 | 0.0 (0/39) | 0.0 | 0.0 (0/0) | 0.6 | 84.7 | 3 |
| 2713 | RF01478 rli47 | 0.0 (0/145) | 0.0 | 0.0 (0/0) | 0.3 | 97.4 | 9 |
| 2714 | RF00859 mir-234 | 0.0 (0/29) | 0.0 | 0.0 (0/0) | 0.8 | 72.6 | 4 |
| 2715 | RF02338 GlsR16 miR3 | 0.0 (0/20) | 0.0 | 0.0 (0/0) | 0.5 | 92.6 | 3 |
| 2716 | RF01519 CC0196 | 0.0 (0/9) | 0.0 | 0.0 (0/0) | 0.0 | 100.0 | 2 |
| 2717 | RF02142 HOXA11-AS1 6 | 0.0 (0/0) | 0.0 | 0.0 (0/0) | 0.0 | 87.9 | 21 |
| 2718 | RF00919 mir-874 | 0.0 (0/21) | 0.0 | 0.0 (0/0) | 0.0 | 99.1 | 4 |
| 2719 | RF01538 TB11Cs3H1 | 0.0 (0/17) | 0.0 | 0.0 (0/0) | 0.8 | 74.8 | 4 |
| 2720 | RF02041 HOTTIP 2 | 0.0 (0/0) | 0.0 | 0.0 (0/0) | 0.0 | 81.6 | 25 |
| 2721 | RF01880 Xist exon1 | 0.0 (0/0) | 0.0 | 0.0 (0/0) | 0.0 | 78.8 | 24 |
| 2722 | RF01122 sR39 | 0.0 (0/0) | 0.0 | 0.0 (0/1) | 0.0 | 82.4 | 5 |
| 2723 | RF02130 GNAS-AS1 4 | 0.0 (0/0) | 0.0 | 0.0 (0/0) | 0.0 | 79.2 | 18 |
| 2724 | RF00473 snosnR54 | 0.0 (0/0) | 0.0 | 0.0 (0/0) | 0.0 | 78.0 | 8 |
| 2725 | RF02791 CcsR1 | 0.0 (0/20) | 0.0 | 0.0 (0/0) | 0.5 | 68.7 | 2 |
| 2726 | RF00891 mir-671 | 0.0 (0/29) | 0.0 | 0.0 (0/0) | 0.0 | 98.2 | 3 |
| 2727 | RF02312 TtnuHACA5 | 0.0 (0/44) | 0.0 | 0.0 (0/0) | 0.0 | 95.4 | 2 |
| 2728 | RF02873 AS-traI | 0.0 (0/22) | 0.0 | 0.0 (0/0) | 0.8 | 89.2 | 3 |
| 2729 | RF02797 PssrA | 0.0 (0/24) | 0.0 | 0.0 (0/0) | 0.5 | 91.5 | 6 |
| 2730 | RF02629 RmaA | 0.0 (0/32) | 0.0 | 0.0 (0/0) | 0.0 | 99.1 | 2 |
| 2731 | RF00873 mir-550 | 0.0 (0/31) | 0.0 | 0.0 (0/0) | 0.1 | 95.9 | 3 |
| 2732 | RF01079 RF site3 | 0.0 (0/8) | 0.0 | 0.0 (0/0) | 0.1 | 94.9 | 2 |
| 2733 | RF01323 CRISPR-DR10 | 0.0 (0/6) | 0.0 | 0.0 (0/0) | 0.0 | 71.1 | 5 |
| 2734 | RF02613 BSR0739 | 0.0 (0/56) | 0.0 | 0.0 (0/0) | 0.0 | 99.1 | 4 |
| 2735 | RF01580 RUF3 | 0.0 (0/46) | 0.0 | 0.0 (0/0) | 0.2 | 92.5 | 3 |
| 2736 | RF01634 ceN44 | 0.0 (0/3) | 0.0 | 0.0 (0/0) | 0.0 | 95.4 | 3 |
| 2737 | RF02458 Virus CITE 3 | 0.0 (0/35) | 0.0 | 0.0 (0/0) | 0.5 | 63.0 | 2 |
| 2738 | RF02153 NBR2 | 0.0 (0/0) | 0.0 | 0.0 (0/0) | 0.0 | 92.7 | 26 |
| 2739 | RF02040 HOTTIP 1 | 0.0 (0/0) | 0.0 | 0.0 (0/1) | 0.0 | 83.4 | 7 |
| 2740 | RF01319 CRISPR-DR6 | 0.0 (0/4) | 0.0 | 0.0 (0/0) | 0.0 | 91.9 | 7 |
| 2741 | RF02206 WT1-AS 4 | 0.0 (0/0) | 0.0 | 0.0 (0/1) | 0.0 | 70.0 | 12 |
| 2742 | RF00939 mir-504 | 0.0 (0/30) | 0.0 | 0.0 (0/0) | 0.3 | 91.7 | 6 |
| 2743 | RF02319 TtnuHACA12 | 0.0 (0/40) | 0.0 | 0.0 (0/0) | 0.5 | 83.0 | 2 |
| 2744 | RF01181 snR77 | 0.0 (0/0) | 0.0 | 0.0 (0/1) | 0.0 | 83.2 | 15 |
| 2745 | RF02912 AAC AAD leader | 0.0 (0/18) | 0.0 | 0.0 (0/0) | 0.8 | 86.0 | 4 |
| 2746 | RF02373 PyrD leader | 0.0 (0/6) | 0.0 | 0.0 (0/0) | 1.3 | 80.9 | 27 |
| 2747 | RF00967 mir-281 | 0.0 (0/29) | 0.0 | 0.0 (0/0) | 0.1 | 78.0 | 2 |
| 2748 | RF01547 TB6Cs1H3 | 0.0 (0/17) | 0.0 | 0.0 (0/0) | 1.9 | 78.5 | 5 |
| 2749 | RF00739 MIR476 | 0.0 (0/36) | 0.0 | 0.0 (0/0) | 0.1 | 88.2 | 2 |
| 2750 | RF02636 EF1368 EF1369 | 0.0 (0/215) | 0.0 | 0.0 (0/0) | 0.0 | 99.1 | 4 |
| 2751 | RF02474 GlsR5 | 0.0 (0/0) | 0.0 | 0.0 (0/0) | 0.0 | 96.5 | 2 |
| 2752 | RF02638 EF0605 EF0606 | 0.0 (0/53) | 0.0 | 0.0 (0/0) | 0.0 | 100.0 | 2 |
| 2753 | RF02417 VR-RNA | 0.0 (0/111) | 0.0 | 0.0 (0/0) | 0.1 | 98.1 | 3 |
| 2754 | RF01804 Lambda thermo | 0.0 (0/20) | 0.0 | 0.0 (0/0) | 1.0 | 89.8 | 5 |
| 2755 | RF01111 SBWMV2 UPD-PKk | 0.0 (0/8) | 0.0 | 0.0 (0/0) | 0.1 | 95.2 | 4 |
| 2756 | RF01024 mir-944 | 0.0 (0/37) | 0.0 | 0.0 (0/0) | 0.0 | 94.3 | 2 |
| 2757 | RF02350 psRNA6 | 0.0 (0/103) | 0.0 | 0.0 (0/0) | 0.2 | 71.7 | 2 |
| 2758 | RF01627 ceN36-1 | 0.0 (0/27) | 0.0 | 0.0 (0/0) | 0.3 | 88.4 | 5 |
| 2759 | RF01432 snoR143 | 0.0 (0/38) | 0.0 | 0.0 (0/0) | 1.5 | 60.6 | 3 |
| 2760 | RF00793 mir-497 | 0.0 (0/34) | 0.0 | 0.0 (0/0) | 0.7 | 89.6 | 8 |
| 2761 | RF02612 BSR1350 | 0.0 (0/52) | 0.0 | 0.0 (0/0) | 0.0 | 98.9 | 3 |

Continued on next page

| | RNA family (seed alignment) | Sensitivity annotated bpairs that covary % (cov_bps/bps) | Power average power % | Positive Predictive Value covarying pairs in structure % (cov_bps/cov_pairs) | average substitutions per bpair | avg pairwise identity % | number of sequences |
|---|---|---|---|---|---|---|---|
| 2762 | RF02545 SSU trypano mito | 0.0 (0/175) | 0.0 | 0.0 (0/0) | 0.7 | 82.9 | 4 |
| 2763 | RF01329 CRISPR-DR15 | 0.0 (0/9) | 0.0 | 0.0 (0/0) | 0.0 | 80.2 | 3 |
| 2764 | RF02141 HOXA11-AS1 5 | 0.0 (0/0) | 0.0 | 0.0 (0/0) | 0.0 | 78.9 | 28 |
| 2765 | RF02631 Hrs13 | 0.0 (0/21) | 0.0 | 0.0 (0/0) | 0.1 | 87.7 | 2 |
| 2766 | RF01630 ceN40 | 0.0 (0/3) | 0.0 | 0.0 (0/0) | 1.7 | 84.7 | 4 |
| 2767 | RF01145 sR14 | 0.0 (0/0) | 0.0 | 0.0 (0/0) | 0.0 | 67.3 | 8 |
| 2768 | RF01810 pntA | 0.0 (0/18) | 0.0 | 0.0 (0/0) | 0.7 | 93.1 | 10 |
| 2769 | RF00337 snoZ112 | 0.0 (0/30) | 0.0 | 0.0 (0/0) | 0.2 | 92.7 | 6 |
| 2770 | RF02508 Atu Ti4 | 0.0 (0/20) | 0.0 | 0.0 (0/0) | 0.9 | 88.0 | 3 |
| 2771 | RF02745 Rev42 5p UTR | 0.0 (0/136) | 0.0 | 0.0 (0/0) | 0.4 | 88.9 | 3 |
| 2772 | RF01460 rliH | 0.0 (0/119) | 0.0 | 0.0 (0/0) | 0.2 | 92.7 | 3 |
| 2773 | RF01438 S pombe snR35 | 0.0 (0/32) | 0.0 | 0.0 (0/0) | 1.1 | 85.6 | 4 |
| 2774 | RF00902 mir-791 | 0.0 (0/25) | 0.0 | 0.0 (0/0) | 0.9 | 70.7 | 3 |
| 2775 | RF02515 AfaR | 0.0 (0/51) | 0.0 | 0.0 (0/0) | 0.0 | 98.7 | 2 |
| 2776 | RF00205 snoR41 | 0.0 (0/6) | 0.0 | 0.0 (0/0) | 1.8 | 90.7 | 7 |
| 2777 | RF01970 RMST 9 | 0.0 (0/0) | 0.0 | 0.0 (0/0) | 0.0 | 89.1 | 28 |
| 2778 | RF01929 bxd 1 | 0.0 (0/0) | 0.0 | 0.0 (0/0) | 0.0 | 87.0 | 6 |
| 2779 | RF00718 mir-431 | 0.0 (0/35) | 0.0 | 0.0 (0/0) | 0.7 | 88.9 | 9 |
| 2780 | RF01351 CRISPR-DR42 | 0.0 (0/0) | 0.0 | 0.0 (0/0) | 0.0 | 92.0 | 3 |
| 2781 | RF02645 Cis2 sRNA | 0.0 (0/52) | 0.0 | 0.0 (0/0) | 0.2 | 80.4 | 2 |
| 2782 | RF01326 CRISPR-DR13 | 0.0 (0/5) | 0.0 | 0.0 (0/0) | 0.0 | 92.8 | 4 |
| 2783 | RF02799 CssrB | 0.0 (0/53) | 0.0 | 0.0 (0/0) | 0.1 | 98.9 | 3 |
| 2784 | RF02259 Vax2os1 1 | 0.0 (0/0) | 0.0 | 0.0 (0/0) | 0.0 | 88.0 | 3 |
| 2785 | RF01081 SBWMV1 UPD-PKe | 0.0 (0/9) | 0.0 | 0.0 (0/0) | 0.8 | 84.6 | 3 |
| 2786 | RF01602 plasmodium snoR27 | 0.0 (0/34) | 0.0 | 0.0 (0/0) | 0.6 | 83.5 | 4 |
| 2787 | RF01158 SNORD108 | 0.0 (0/3) | 0.0 | 0.0 (0/0) | 0.3 | 77.9 | 4 |
| 2788 | RF02325 TtnuHACA18 | 0.0 (0/34) | 0.0 | 0.0 (0/0) | 0.4 | 85.6 | 3 |
| 2789 | RF02824 LPR10 | 0.0 (0/16) | 0.0 | 0.0 (0/0) | 0.9 | 79.4 | 3 |
| 2790 | RF02706 TeloSII ncR30 | 0.0 (0/36) | 0.0 | 0.0 (0/0) | 0.0 | 94.2 | 2 |
| 2791 | RF02150 MESTIT1 3 | 0.0 (0/0) | 0.0 | 0.0 (0/0) | 0.0 | 78.7 | 18 |
| 2792 | RF02166 PVT1 3 | 0.0 (0/0) | 0.0 | 0.0 (0/0) | 0.0 | 67.8 | 25 |
| 2793 | RF01571 DdR35 | 0.0 (0/15) | 0.0 | 0.0 (0/0) | 0.0 | 98.9 | 3 |
| 2794 | RF01135 sR24 | 0.0 (0/2) | 0.0 | 0.0 (0/0) | 0.0 | 95.2 | 3 |
| 2795 | RF02113 DLG2-AS1 2 | 0.0 (0/0) | 0.0 | 0.0 (0/0) | 0.0 | 74.6 | 23 |
| 2796 | RF02467 Ms IGR-2 | 0.0 (0/13) | 0.0 | 0.0 (0/0) | 0.4 | 84.4 | 3 |
| 2797 | RF01475 rli45 | 0.0 (0/14) | 0.0 | 0.0 (0/0) | 1.3 | 84.7 | 6 |
| 2798 | RF00811 mir-84 | 0.0 (0/26) | 0.0 | 0.0 (0/0) | 0.5 | 71.5 | 4 |
| 2799 | RF02277 Hammerhead HH10 | 0.0 (0/34) | 0.0 | 0.0 (0/0) | 0.4 | 90.9 | 18 |
| 2800 | RF01648 ceN65 | 0.0 (0/3) | 0.0 | 0.0 (0/0) | 0.0 | 95.9 | 3 |
| 2801 | RF01957 NEAT1 3 | 0.0 (0/0) | 0.0 | 0.0 (0/0) | 0.0 | 87.5 | 18 |
| 2802 | RF02200 TTC28-AS1 3 | 0.0 (0/0) | 0.0 | 0.0 (0/0) | 0.0 | 67.7 | 6 |
| 2803 | RF00784 mir-486 | 0.0 (0/24) | 0.0 | 0.0 (0/1) | 0.9 | 88.8 | 12 |
| 2804 | RF02010 mir-3180 | 0.0 (0/34) | 0.0 | 0.0 (0/0) | 0.0 | 100.0 | 3 |
| 2805 | RF01212 sn2524 | 0.0 (0/3) | 0.0 | 0.0 (0/0) | 0.7 | 70.8 | 2 |
| 2806 | RF01901 mir-284 | 0.0 (0/34) | 0.0 | 0.0 (0/0) | 0.3 | 92.7 | 10 |
| 2807 | RF02573 LLnc147 | 0.0 (0/30) | 0.0 | 0.0 (0/0) | 0.1 | 94.1 | 2 |
| 2808 | RF02879 MH s10 | 0.0 (0/30) | 0.0 | 0.0 (0/1) | 0.8 | 80.0 | 3 |
| 2809 | RF01613 ceN109 | 0.0 (0/3) | 0.0 | 0.0 (0/1) | 0.0 | 83.1 | 3 |
| 2810 | RF02861 IR 84 | 0.0 (0/54) | 0.0 | 0.0 (0/0) | 1.0 | 75.8 | 3 |
| 2811 | RF01884 LINC00901 | 0.0 (0/0) | 0.0 | 0.0 (0/0) | 0.0 | 88.8 | 30 |
| 2812 | RF00997 mir-942 | 0.0 (0/36) | 0.0 | 0.0 (0/0) | 0.0 | 95.3 | 2 |
| 2813 | RF02397 sau-5971 | 0.0 (0/35) | 0.0 | 0.0 (0/0) | 0.1 | 99.3 | 3 |
| 2814 | RF02078 STnc210 | 0.0 (0/24) | 0.0 | 0.0 (0/0) | 0.1 | 80.5 | 2 |
| 2815 | RF00227 FIE3 | 0.0 (0/9) | 0.0 | 0.0 (0/0) | 1.7 | 65.2 | 5 |
| 2816 | RF02443 SpR08 sRNA | 0.0 (0/17) | 0.0 | 0.0 (0/0) | 0.3 | 97.3 | 3 |
| 2817 | RF02377 SurA | 0.0 (0/71) | 0.0 | 0.0 (0/0) | 0.3 | 94.5 | 4 |
| 2818 | RF02654 MicL | 0.0 (0/105) | 0.0 | 0.0 (0/0) | 0.0 | 99.0 | 2 |
| 2819 | RF00116 C0465 | 0.0 (0/23) | 0.0 | 0.0 (0/0) | 0.4 | 98.9 | 8 |
| 2820 | RF01936 mir-63 | 0.0 (0/48) | 0.0 | 0.0 (0/0) | 1.4 | 85.5 | 5 |
| 2821 | RF02843 RefA5 | 0.0 (0/20) | 0.0 | 0.0 (0/0) | 0.3 | 82.8 | 3 |
| 2822 | RF02584 KRAS 3UTR | 0.0 (0/10) | 0.0 | 0.0 (0/0) | 0.0 | 98.0 | 2 |
| 2823 | RF02780 PutA thermometer | 0.0 (0/15) | 0.0 | 0.0 (0/1) | 1.3 | 83.8 | 4 |
| 2824 | RF01479 rli48 | 0.0 (0/41) | 0.0 | 0.0 (0/0) | 0.2 | 87.0 | 3 |
| 2825 | RF01364 CRISPR-DR51 | 0.0 (0/6) | 0.0 | 0.0 (0/0) | 0.3 | 78.4 | 2 |
| 2826 | RF02637 EF0408 EF0409 | 0.0 (0/127) | 0.0 | 0.0 (0/0) | 0.0 | 98.4 | 3 |
| 2827 | RF00387 IRES FGF1 | 0.0 (0/51) | 0.0 | 0.0 (0/0) | 1.2 | 87.2 | 6 |
| 2828 | RF01443 S pombe snR91 | 0.0 (0/24) | 0.0 | 0.0 (0/0) | 0.0 | 99.3 | 2 |
| 2829 | RF01349 CRISPR-DR40 | 0.0 (0/10) | 0.0 | 0.0 (0/0) | 0.0 | 91.1 | 3 |
| 2830 | RF00799 mir-354 | 0.0 (0/34) | 0.0 | 0.0 (0/0) | 0.1 | 91.3 | 4 |
| 2831 | RF00895 mir-786 | 0.0 (0/32) | 0.0 | 0.0 (0/0) | 0.3 | 64.4 | 2 |
| 2832 | RF02649 Trans44 sRNA | 0.0 (0/83) | 0.0 | 0.0 (0/0) | 0.2 | 85.3 | 2 |
| 2833 | RF02615 BSR0626 | 0.0 (0/30) | 0.0 | 0.0 (0/0) | 0.0 | 99.2 | 2 |
| 2834 | RF02089 CLRN1-AS1 | 0.0 (0/0) | 0.0 | 0.0 (0/0) | 0.0 | 73.6 | 19 |
| 2835 | RF02894 S35 | 0.0 (0/95) | 0.0 | 0.0 (0/0) | 1.2 | 61.4 | 3 |
| 2836 | RF00569 SNORD19 | 0.0 (0/4) | 0.0 | 0.0 (0/0) | 1.8 | 83.7 | 22 |
| 2837 | RF00903 mir-359 | 0.0 (0/35) | 0.0 | 0.0 (0/0) | 0.2 | 74.7 | 2 |
| 2838 | RF02693 psm mec RNA | 0.0 (0/38) | 0.0 | 0.0 (0/0) | 0.5 | 91.8 | 4 |
| 2839 | RF02593 NsiR8 | 0.0 (0/52) | 0.0 | 0.0 (0/0) | 0.1 | 94.8 | 2 |
| 2840 | RF02906 sRNA154 | 0.0 (0/32) | 0.0 | 0.0 (0/0) | 0.6 | 80.5 | 3 |
| 2841 | RF00748 mir-433 | 0.0 (0/22) | 0.0 | 0.0 (0/0) | 0.0 | 94.4 | 5 |
| 2842 | RF01966 RMST 5 | 0.0 (0/0) | 0.0 | 0.0 (0/0) | 0.0 | 87.2 | 21 |
| 2843 | RF02236 asX2 | 0.0 (0/18) | 0.0 | 0.0 (0/0) | 0.2 | 81.2 | 2 |
| 2844 | RF02125 JPX 2 | 0.0 (0/0) | 0.0 | 0.0 (0/1) | 0.0 | 74.8 | 11 |
| 2845 | RF02818 V AS5 | 0.0 (0/38) | 0.0 | 0.0 (0/0) | 0.9 | 69.2 | 3 |

Continued on next page

| | RNA family (seed alignment) | Sensitivity annotated bpairs that covary % (cov_bps/bps) | Power average power % | Positive Predictive Value covarying pairs in structure % (cov_bps/cov_pairs) | average substitutions per bpair | avg pairwise identity % | number of sequences |
|---|---|---|---|---|---|---|---|
| 2846 | RF00159 snoZ168 | 0.0 (0/4) | 0.0 | 0.0 (0/1) | 0.0 | 84.5 | 6 |
| 2847 | RF00987 mir-589 | 0.0 (0/34) | 0.0 | 0.0 (0/0) | 0.1 | 96.0 | 2 |
| 2848 | RF02810 Lst thermometer | 0.0 (0/32) | 0.0 | 0.0 (0/0) | 0.1 | 97.1 | 2 |
| 2849 | RF01176 snR78 | 0.0 (0/2) | 0.0 | 0.0 (0/0) | 4.5 | 72.1 | 8 |
| 2850 | RF02731 JA03 | 0.0 (0/78) | 0.0 | 0.0 (0/0) | 0.1 | 93.7 | 3 |
| 2851 | RF01978 HOTAIRM1 4 | 0.0 (0/0) | 0.0 | 0.0 (0/0) | 0.0 | 86.2 | 10 |
| 2852 | RF00262 sar | 0.0 (0/20) | 0.0 | 0.0 (0/0) | 0.8 | 82.3 | 4 |
| 2853 | RF01376 CRISPR-DR63 | 0.0 (0/8) | 0.0 | 0.0 (0/0) | 0.0 | 97.3 | 2 |
| 2854 | RF00824 mir-50 | 0.0 (0/37) | 0.0 | 0.0 (0/0) | 2.0 | 70.5 | 5 |
| 2855 | RF00975 MIR845 2 | 0.0 (0/30) | 0.0 | 0.0 (0/0) | 0.0 | 91.6 | 2 |
| 2856 | RF01343 CRISPR-DR33 | 0.0 (0/6) | 0.0 | 0.0 (0/0) | 0.2 | 66.7 | 3 |
| 2857 | RF01002 mir-936 | 0.0 (0/30) | 0.0 | 0.0 (0/0) | 1.7 | 78.0 | 4 |
| 2858 | RF02176 SMAD5-AS1 4 | 0.0 (0/0) | 0.0 | 0.0 (0/0) | 0.0 | 82.9 | 19 |
| 2859 | RF00955 mir-1829 | 0.0 (0/23) | 0.0 | 0.0 (0/0) | 0.6 | 87.8 | 3 |
| 2860 | RF02886 npcTB 6715 | 0.0 (0/61) | 0.0 | 0.0 (0/3) | 1.1 | 71.8 | 3 |
| 2861 | RF00351 snoR20 | 0.0 (0/3) | 0.0 | 0.0 (0/0) | 2.0 | 83.7 | 4 |
| 2862 | RF00249 mir-46 | 0.0 (0/22) | 0.0 | 0.0 (0/0) | 1.3 | 73.9 | 11 |
| 2863 | RF00899 mir-235 | 0.0 (0/19) | 0.0 | 0.0 (0/0) | 0.1 | 78.6 | 2 |
| 2864 | RF02145 MEG8 1 | 0.0 (0/0) | 0.0 | 0.0 (0/0) | 0.0 | 74.3 | 23 |
| 2865 | RF01882 TUG1 1 | 0.0 (0/0) | 0.0 | 0.0 (0/0) | 0.0 | 89.9 | 21 |
| 2866 | RF02310 TtnuHACA2 | 0.0 (0/34) | 0.0 | 0.0 (0/0) | 0.1 | 92.4 | 3 |
| 2867 | RF02299 TtnuCD23 | 0.0 (0/0) | 0.0 | 0.0 (0/0) | 0.0 | 93.1 | 2 |
| 2868 | RF00960 mir-661 | 0.0 (0/30) | 0.0 | 0.0 (0/0) | 1.5 | 78.2 | 4 |
| 2869 | RF02478 GlsR10 | 0.0 (0/3) | 0.0 | 0.0 (0/0) | 0.0 | 97.1 | 2 |
| 2870 | RF01624 ceN27 | 0.0 (0/3) | 0.0 | 0.0 (0/0) | 0.3 | 90.0 | 2 |
| 2871 | RF01325 CRISPR-DR12 | 0.0 (0/10) | 0.0 | 0.0 (0/0) | 1.0 | 89.0 | 5 |
| 2872 | RF02816 HBV PRE 1151-1410 | 0.0 (0/82) | 0.0 | 0.0 (0/0) | 0.0 | 94.6 | 2 |
| 2873 | RF02743 Sernc389 | 0.0 (0/210) | 0.0 | 0.0 (0/0) | 0.1 | 86.5 | 2 |
| 2874 | RF00965 mir-549 | 0.0 (0/29) | 0.0 | 0.0 (0/0) | 0.0 | 97.9 | 2 |
| 2875 | RF02433 SpF36 sRNA | 0.0 (0/14) | 0.0 | 0.0 (0/0) | 0.1 | 91.5 | 4 |
| 2876 | RF01180 snoR110 | 0.0 (0/18) | 0.0 | 0.0 (0/0) | 0.0 | 91.6 | 2 |
| 2877 | RF00329 snoZ162 | 0.0 (0/13) | 0.0 | 0.0 (0/0) | 1.2 | 83.1 | 9 |
| 2878 | RF01609 ceN103 | 0.0 (0/3) | 0.0 | 0.0 (0/0) | 0.0 | 78.2 | 4 |
| 2879 | RF01000 mir-580 | 0.0 (0/40) | 0.0 | 0.0 (0/0) | 0.1 | 95.9 | 2 |
| 2880 | RF00893 MIR854 | 0.0 (0/18) | 0.0 | 0.0 (0/0) | 0.0 | 97.9 | 3 |
| 2881 | RF02694 RalA | 0.0 (0/48) | 0.0 | 0.0 (0/2) | 0.5 | 94.6 | 10 |
| 2882 | RF02764 Ysr190 | 0.0 (0/21) | 0.0 | 0.0 (0/0) | 0.3 | 92.0 | 4 |
| 2883 | RF01310 sR7 | 0.0 (0/0) | 0.0 | 0.0 (0/0) | 0.0 | 70.9 | 5 |
| 2884 | RF00929 mir-574 | 0.0 (0/32) | 0.0 | 0.0 (0/0) | 0.2 | 93.5 | 3 |
| 2885 | RF02480 GlsR14 | 0.0 (0/0) | 0.0 | 0.0 (0/0) | 0.0 | 93.1 | 3 |
| 2886 | RF02047 Sphinx 2 | 0.0 (0/0) | 0.0 | 0.0 (0/0) | 0.0 | 79.9 | 37 |
| 2887 | RF02670 scaDm46E3 | 0.0 (0/37) | 0.0 | 0.0 (0/0) | 0.0 | 97.8 | 2 |
| 2888 | RF01471 rliB | 0.0 (0/49) | 0.0 | 0.0 (0/0) | 1.2 | 69.7 | 4 |
| 2889 | RF02756 ES239 | 0.0 (0/34) | 0.0 | 0.0 (0/0) | 0.3 | 96.8 | 5 |
| 2890 | RF01561 DdR11 | 0.0 (0/0) | 0.0 | 0.0 (0/0) | 0.0 | 88.7 | 3 |
| 2891 | RF01881 Xist exon4 | 0.0 (0/0) | 0.0 | 0.0 (0/0) | 0.0 | 87.6 | 25 |
| 2892 | RF00984 mir-576 | 0.0 (0/38) | 0.0 | 0.0 (0/0) | 0.1 | 91.8 | 2 |
| 2893 | RF01102 PK1-TEV CVMV | 0.0 (0/11) | 0.0 | 0.0 (0/0) | 0.2 | 80.6 | 2 |
| 2894 | RF02798 CssrA | 0.0 (0/29) | 0.0 | 0.0 (0/0) | 0.4 | 91.2 | 4 |
| 2895 | RF02016 mir-1183 | 0.0 (0/25) | 0.0 | 0.0 (0/0) | 0.3 | 98.5 | 3 |
| 2896 | RF01165 sn2317 | 0.0 (0/3) | 0.0 | 0.0 (0/0) | 0.0 | 85.0 | 4 |
| 2897 | RF00909 mir-883 | 0.0 (0/28) | 0.0 | 0.0 (0/0) | 0.4 | 85.3 | 3 |
| 2898 | RF01564 DdR14 | 0.0 (0/0) | 0.0 | 0.0 (0/0) | 0.0 | 100.0 | 2 |
| 2899 | RF02015 mir-1287 | 0.0 (0/32) | 0.0 | 0.0 (0/0) | 0.2 | 91.8 | 3 |
| 2900 | RF02273 FsrA | 0.0 (0/20) | 0.0 | 0.0 (0/0) | 0.0 | 96.1 | 6 |
| 2901 | RF02741 Rev44 | 0.0 (0/101) | 0.0 | 0.0 (0/0) | 0.3 | 92.5 | 3 |
| 2902 | RF02242 Xoo5 | 0.0 (0/30) | 0.0 | 0.0 (0/0) | 0.4 | 80.1 | 3 |
| 2903 | RF01541 TB11Cs4H3 | 0.0 (0/13) | 0.0 | 0.0 (0/0) | 0.5 | 76.1 | 3 |
| 2904 | RF01798 g2 | 0.0 (0/21) | 0.0 | 0.0 (0/0) | 0.0 | 100.0 | 2 |
| 2905 | RF00453 Cardiovirus CRE | 0.0 (0/10) | 0.0 | 0.0 (0/0) | 0.8 | 81.3 | 4 |
| 2906 | RF01920 mir-764 | 0.0 (0/30) | 0.0 | 0.0 (0/0) | 0.2 | 91.0 | 6 |
| 2907 | RF01976 HOTAIRM1 2 | 0.0 (0/0) | 0.0 | 0.0 (0/0) | 0.0 | 86.8 | 7 |
| 2908 | RF01950 KCNQ1OT1 5 | 0.0 (0/0) | 0.0 | 0.0 (0/0) | 0.0 | 88.8 | 15 |
| 2909 | RF00158 SNORD82 | 0.0 (0/5) | 0.0 | 0.0 (0/0) | 1.4 | 80.3 | 27 |
| 2910 | RF00393 SNORA8 | 0.0 (0/37) | 0.0 | 0.0 (0/0) | 0.3 | 89.7 | 5 |
| 2911 | RF02494 Gl U6 | 0.0 (0/19) | 0.0 | 0.0 (0/0) | 0.0 | 97.1 | 2 |
| 2912 | RF01800 gadd7 | 0.0 (0/239) | 0.0 | 0.0 (0/0) | 0.0 | 99.7 | 2 |
| 2913 | RF01792 eiav FSE | 0.0 (0/5) | 0.0 | 0.0 (0/0) | 0.0 | 92.6 | 5 |
| 2914 | RF02630 Hrs12 | 0.0 (0/23) | 0.0 | 0.0 (0/0) | 0.1 | 88.5 | 2 |
| 2915 | RF01114 TMV UPD-PK1 | 0.0 (0/7) | 0.0 | 0.0 (0/0) | 0.0 | 100.0 | 2 |
| 2916 | RF01488 rli49 | 0.0 (0/31) | 0.0 | 0.0 (0/0) | 0.5 | 92.3 | 5 |
| 2917 | RF02479 GlsR13 | 0.0 (0/0) | 0.0 | 0.0 (0/0) | 0.0 | 92.7 | 3 |
| 2918 | RF01645 ceN58 | 0.0 (0/40) | 0.0 | 0.0 (0/0) | 0.3 | 88.5 | 3 |
| 2919 | RF00946 mir-1225 | 0.0 (0/13) | 0.0 | 0.0 (0/0) | 0.2 | 94.5 | 7 |
| 2920 | RF00879 mir-615 | 0.0 (0/28) | 0.0 | 0.0 (0/0) | 0.2 | 97.2 | 6 |
| 2921 | RF01975 HOTAIRM1 1 | 0.0 (0/0) | 0.0 | 0.0 (0/0) | 0.0 | 88.7 | 14 |
| 2922 | RF01406 STnc500 | 0.0 (0/67) | 0.0 | 0.0 (0/0) | 0.1 | 98.8 | 8 |
| 2923 | RF02219 ZNRD1-AS1 2 | 0.0 (0/0) | 0.0 | 0.0 (0/0) | 0.0 | 77.2 | 34 |
| 2924 | RF00382 DnaX | 0.0 (0/11) | 0.0 | 0.0 (0/0) | 0.4 | 85.7 | 5 |
| 2925 | RF00971 mir-578 | 0.0 (0/36) | 0.0 | 0.0 (0/0) | 0.2 | 95.6 | 3 |
| 2926 | RF01592 plasmodium snoR17 | 0.0 (0/4) | 0.0 | 0.0 (0/0) | 0.2 | 86.9 | 5 |
| 2927 | RF02875 AS-pc01 | 0.0 (0/28) | 0.0 | 0.0 (0/0) | 0.8 | 72.2 | 3 |
| 2928 | RF02617 SSRC10 | 0.0 (0/34) | 0.0 | 0.0 (0/0) | 0.0 | 97.0 | 3 |
| 2929 | RF02280 TtnuCD3 | 0.0 (0/0) | 0.0 | 0.0 (0/0) | 0.0 | 94.0 | 3 |

Continued on next page

| | RNA family (seed alignment) | Sensitivity annotated bpairs that covary % (cov_bps/bps) | Power average power % | Positive Predictive Value covarying pairs in structure % (cov_bps/cov_pairs) | average substitutions per bpair | avg pairwise identity % | number of sequences |
|---|---|---|---|---|---|---|---|
| 2930 | RF00320 snoZ185 | 0.0 (0/6) | 0.0 | 0.0 (0/0) | 1.8 | 93.8 | 5 |
| 2931 | RF02323 TtnuHACA16 | 0.0 (0/30) | 0.0 | 0.0 (0/0) | 0.0 | 94.9 | 2 |
| 2932 | RF01334 CRISPR-DR21 | 0.0 (0/8) | 0.0 | 0.0 (0/0) | 0.0 | 97.3 | 2 |
| 2933 | RF01274 sR45 | 0.0 (0/14) | 0.0 | 0.0 (0/0) | 0.6 | 82.1 | 7 |
| 2934 | RF01193 snoR20a | 0.0 (0/3) | 0.0 | 0.0 (0/0) | 0.0 | 69.2 | 5 |
| 2935 | RF00371 sroE | 0.0 (0/19) | 0.0 | 0.0 (0/0) | 0.8 | 83.6 | 4 |
| 2936 | RF02549 YFV 3UTR | 0.0 (0/24) | 0.0 | 0.0 (0/0) | 0.0 | 98.3 | 3 |
| 2937 | RF02054 STnc420 | 0.0 (0/13) | 0.0 | 0.0 (0/0) | 0.2 | 95.4 | 3 |
| 2938 | RF01977 HOTAIRM1 3 | 0.0 (0/0) | 0.0 | 0.0 (0/1) | 0.0 | 88.0 | 11 |
| 2939 | RF01678 P37 | 0.0 (0/23) | 0.0 | 0.0 (0/0) | 0.0 | 95.2 | 2 |
| 2940 | RF00068 SNORD21 | 0.0 (0/4) | 0.0 | 0.0 (0/0) | 2.2 | 71.5 | 5 |
| 2941 | RF00352 snoR21 | 0.0 (0/6) | 0.0 | 0.0 (0/0) | 1.3 | 77.6 | 6 |
| 2942 | RF00970 mir-648 | 0.0 (0/27) | 0.0 | 0.0 (0/0) | 0.0 | 92.5 | 2 |
| 2943 | RF02186 ST7-OT3 4 | 0.0 (0/0) | 0.0 | 0.0 (0/0) | 0.0 | 87.3 | 8 |
| 2944 | RF02668 PsiU2-55 | 0.0 (0/52) | 0.0 | 0.0 (0/0) | 0.2 | 96.6 | 3 |
| 2945 | RF01313 AHBV epsilon | 0.0 (0/16) | 0.0 | 0.0 (0/0) | 0.0 | 92.7 | 10 |
| 2946 | RF02330 TtnuHACA23 | 0.0 (0/37) | 0.0 | 0.0 (0/0) | 0.2 | 72.9 | 2 |
| 2947 | RF01812 Pxr | 0.0 (0/37) | 0.0 | 0.0 (0/0) | 0.1 | 93.5 | 2 |
| 2948 | RF02663 GmglnB1 5p UTR | 0.0 (0/26) | 0.0 | 0.0 (0/0) | 0.0 | 98.8 | 2 |
| 2949 | RF00993 mir-1473 | 0.0 (0/29) | 0.0 | 0.0 (0/0) | 0.1 | 76.9 | 2 |
| 2950 | RF02329 TtnuHACA22 | 0.0 (0/30) | 0.0 | 0.0 (0/0) | 0.1 | 93.9 | 2 |
| 2951 | RF01123 sR35 | 0.0 (0/0) | 0.0 | 0.0 (0/0) | 0.0 | 84.6 | 3 |
| 2952 | RF01651 ceN69 | 0.0 (0/5) | 0.0 | 0.0 (0/0) | 0.4 | 91.6 | 4 |
| 2953 | RF01530 CC3664 | 0.0 (0/43) | 0.0 | 0.0 (0/0) | 0.0 | 100.0 | 2 |
| 2954 | RF00463 ApoB 5 CRE | 0.0 (0/41) | 0.0 | 0.0 (0/0) | 0.8 | 90.9 | 5 |
| 2955 | RF01928 Mico1 | 0.0 (0/0) | 0.0 | 0.0 (0/0) | 0.0 | 82.9 | 3 |
| 2956 | RF00632 sxy | 0.0 (0/33) | 0.0 | 0.0 (0/0) | 0.0 | 97.1 | 2 |
| 2957 | RF02175 SMAD5-AS1 3 | 0.0 (0/0) | 0.0 | 0.0 (0/0) | 0.0 | 79.2 | 18 |
| 2958 | RF01481 rli53 | 0.0 (0/53) | 0.0 | 0.0 (0/0) | 0.1 | 95.8 | 5 |
| 2959 | RF01403 STnc290 | 0.0 (0/18) | 0.0 | 0.0 (0/0) | 0.3 | 99.0 | 6 |
| 2960 | RF02664 IRES cyp24a1 | 0.0 (0/96) | 0.0 | 0.0 (0/0) | 0.0 | 100.0 | 2 |
| 2961 | RF02729 JA01 | 0.0 (0/46) | 0.0 | 0.0 (0/0) | 0.0 | 99.6 | 3 |
| 2962 | RF02127 GNAS-AS1 1 | 0.0 (0/0) | 0.0 | 0.0 (0/0) | 0.0 | 80.6 | 21 |
| 2963 | RF01356 CRISPR-DR28 | 0.0 (0/7) | 0.0 | 0.0 (0/0) | 0.1 | 91.7 | 3 |
| 2964 | RF02332 TtnuHACA25 | 0.0 (0/32) | 0.0 | 0.0 (0/0) | 0.0 | 97.3 | 2 |
| 2965 | RF02619 BSnc115 | 0.0 (0/23) | 0.0 | 0.0 (0/0) | 0.1 | 97.1 | 3 |
| 2966 | RF01301 snoR4a | 0.0 (0/8) | 0.0 | 0.0 (0/0) | 0.0 | 97.4 | 2 |
| 2967 | RF02749 Ncrwmel02 | 0.0 (0/78) | 0.0 | 0.0 (0/0) | 0.2 | 97.1 | 3 |
| 2968 | RF00615 LhrA | 0.0 (0/82) | 0.0 | 0.0 (0/0) | 0.1 | 99.0 | 10 |
| 2969 | RF02156 NPPA-AS1 1 | 0.0 (0/0) | 0.0 | 0.0 (0/0) | 0.0 | 74.1 | 19 |
| 2970 | RF02412 snoR144 | 0.0 (0/40) | 0.0 | 0.0 (0/0) | 0.4 | 87.7 | 4 |
| 2971 | RF01386 isrB | 0.0 (0/21) | 0.0 | 0.0 (0/0) | 0.5 | 83.9 | 3 |
| 2972 | RF02121 FTX 3 | 0.0 (0/0) | 0.0 | 0.0 (0/0) | 0.0 | 74.6 | 14 |
| 2973 | RF00092 SNORA63 | 0.0 (0/23) | 0.0 | 0.0 (0/0) | 0.9 | 80.7 | 8 |
| 2974 | RF01872 MEG3 2 | 0.0 (0/0) | 0.0 | 0.0 (0/0) | 0.0 | 87.2 | 16 |
| 2975 | RF02306 TtnuCD32 | 0.0 (0/0) | 0.0 | 0.0 (0/0) | 0.0 | 94.2 | 3 |
| 2976 | RF01639 ceN49 | 0.0 (0/34) | 0.0 | 0.0 (0/0) | 0.4 | 88.6 | 3 |
| 2977 | RF01522 TB10Cs1H1 | 0.0 (0/15) | 0.0 | 0.0 (0/0) | 0.9 | 84.6 | 4 |
| 2978 | RF02483 GlsR20 | 0.0 (0/29) | 0.0 | 0.0 (0/0) | 0.3 | 86.9 | 3 |
| 2979 | RF02190 ST7-OT4 4 | 0.0 (0/0) | 0.0 | 0.0 (0/0) | 0.0 | 77.0 | 26 |
| 2980 | RF01293 ACA59 | 0.0 (0/32) | 0.0 | 0.0 (0/0) | 1.3 | 75.1 | 3 |
| 2981 | RF02239 asX6 | 0.0 (0/31) | 0.0 | 0.0 (0/0) | 0.4 | 94.2 | 3 |
| 2982 | RF01266 snR45 | 0.0 (0/0) | 0.0 | 0.0 (0/1) | 0.0 | 79.3 | 8 |
| 2983 | RF02591 Spy1786666 | 0.0 (0/26) | 0.0 | 0.0 (0/0) | 0.0 | 97.7 | 2 |
| 2984 | RF02196 TP53TG1 2 | 0.0 (0/0) | 0.0 | 0.0 (0/0) | 0.0 | 65.0 | 12 |
| 2985 | RF01202 sn2991 | 0.0 (0/6) | 0.0 | 0.0 (0/0) | 0.3 | 86.7 | 4 |
| 2986 | RF02760 sR035 | 0.0 (0/65) | 0.0 | 0.0 (0/0) | 0.2 | 98.0 | 3 |
| 2987 | RF02608 AbsR11 | 0.0 (0/62) | 0.0 | 0.0 (0/0) | 0.0 | 95.5 | 2 |
| 2988 | RF01909 CDKN2B-AS | 0.0 (0/0) | 0.0 | 0.0 (0/0) | 0.0 | 79.9 | 25 |
| 2989 | RF01780 AS1890 | 0.0 (0/28) | 0.0 | 0.0 (0/0) | 0.0 | 100.0 | 2 |
| 2990 | RF02337 GlsR8 miR10 | 0.0 (0/20) | 0.0 | 0.0 (0/0) | 0.0 | 98.1 | 3 |
| 2991 | RF02603 BASRCII26 | 0.0 (0/61) | 0.0 | 0.0 (0/0) | 0.0 | 100.0 | 2 |
| 2992 | RF01588 snoR13 | 0.0 (0/4) | 0.0 | 0.0 (0/0) | 1.8 | 85.6 | 4 |
| 2993 | RF00714 MIR535 | 0.0 (0/27) | 0.0 | 0.0 (0/0) | 1.0 | 65.9 | 4 |
| 2994 | RF01416 NrrF | 0.0 (0/33) | 0.0 | 0.0 (0/0) | 0.0 | 95.6 | 3 |
| 2995 | RF02063 STnc350 | 0.0 (0/13) | 0.0 | 0.0 (0/0) | 0.0 | 85.1 | 2 |
| 2996 | RF00441 snoZ242 | 0.0 (0/19) | 0.0 | 0.0 (0/0) | 1.0 | 88.2 | 5 |
| 2997 | RF00500 TCV H5 | 0.0 (0/12) | 0.0 | 0.0 (0/0) | 0.7 | 97.1 | 5 |
| 2998 | RF01877 MIAT exon5 3 | 0.0 (0/0) | 0.0 | 0.0 (0/0) | 0.0 | 84.6 | 16 |
| 2999 | RF02866 ncS16 | 0.0 (0/19) | 0.0 | 0.0 (0/0) | 0.7 | 85.7 | 4 |
| 3000 | RF02769 Ysr202 | 0.0 (0/86) | 0.0 | 0.0 (0/0) | 0.1 | 94.6 | 2 |
| 3001 | RF00896 mir-787 | 0.0 (0/31) | 0.0 | 0.0 (0/0) | 0.4 | 63.3 | 2 |
| 3002 | RF02021 mir-3179 | 0.0 (0/37) | 0.0 | 0.0 (0/0) | 0.0 | 100.0 | 3 |
| 3003 | RF01143 sR16 | 0.0 (0/0) | 0.0 | 0.0 (0/0) | 0.0 | 100.0 | 2 |
| 3004 | RF02525 SSRC30 | 0.0 (0/40) | 0.0 | 0.0 (0/0) | 0.1 | 98.4 | 3 |
| 3005 | RF02720 sca ncR21 | 0.0 (0/26) | 0.0 | 0.0 (0/0) | 0.2 | 84.3 | 2 |
| 3006 | RF01383 GRIK4 3p UTR | 0.0 (0/9) | 0.0 | 0.0 (0/0) | 0.2 | 92.4 | 8 |
| 3007 | RF02069 STnc70 | 0.0 (0/37) | 0.0 | 0.0 (0/0) | 0.0 | 93.3 | 2 |
| 3008 | RF01064 mir-253 | 0.0 (0/37) | 0.0 | 0.0 (0/0) | 0.8 | 79.2 | 4 |
| 3009 | RF01195 snR52 | 0.0 (0/0) | 0.0 | 0.0 (0/0) | 0.0 | 96.9 | 5 |
| 3010 | RF02704 LcrF thermometer | 0.0 (0/39) | 0.0 | 0.0 (0/0) | 0.0 | 100.0 | 2 |
| 3011 | RF01090 RF site4 | 0.0 (0/18) | 0.0 | 0.0 (0/0) | 0.8 | 86.5 | 7 |
| 3012 | RF01918 mir-1249 | 0.0 (0/29) | 0.0 | 0.0 (0/0) | 0.0 | 97.5 | 4 |
| 3013 | RF00626 Gurken | 0.0 (0/24) | 0.0 | 0.0 (0/0) | 0.1 | 98.1 | 3 |

Continued on next page

| | RNA family (seed alignment) | **Sensitivity** annotated bpairs that covary % (cov_bps/bps) | **Power** average power % | **Positive Predictive Value** covarying pairs in structure % (cov_bps/cov_pairs) | average substitutions per bpair | avg pairwise identity % | number of sequences |
|---|---|---|---|---|---|---|---|
| 3014 | RF01025 mir-934 | 0.0 (0/36) | 0.0 | 0.0 (0/0) | 0.1 | 94.0 | 2 |
| 3015 | RF01194 sn2903 | 0.0 (0/4) | 0.0 | 0.0 (0/0) | 0.0 | 98.7 | 4 |
| 3016 | RF01289 snoR17 | 0.0 (0/10) | 0.0 | 0.0 (0/0) | 0.2 | 70.4 | 2 |
