## Supplementary material for "Estimating the power of sequence covariation for detecting conserved RNA structure": Table_S2

| | RNA family (seed alignment) | observed bps in structure | power % | expected bps to covary | average substitutions per bpair | avg pairwise identity % | number of sequences |
|---|---|---|---|---|---|---|---|
| 1032 | RF01419 IsrR | 11 | 77.3 | 9 | 157.2 | 66.9 | 308 |
| 1033 | RF01942 mir-1937 | 11 | 66.4 | 7 | 84.6 | 65.8 | 171 |
| 1034 | RF00523 Prion pknot | 6 | 50.0 | 3 | 53.0 | 85.0 | 148 |
| 1035 | RF00485 K chan RES | 24 | 41.2 | 10 | 45.1 | 68.7 | 85 |
| 1036 | RF00468 HCV SLVII | 17 | 38.8 | 7 | 42.9 | 74.3 | 110 |
| 1037 | RF00693 mir-147 | 15 | 29.3 | 4 | 28.5 | 68.5 | 64 |
| 1038 | RF00480 HIV FE | 10 | 27.0 | 3 | 32.2 | 84.0 | 145 |
| 1039 | RF00093 SNORD18 | 4 | 22.5 | 1 | 21.5 | 69.9 | 16 |
| 1040 | RF00376 HIV GSL3 | 8 | 21.2 | 2 | 22.0 | 81.4 | 72 |
| 1041 | RF01753 psbNH | 12 | 20.8 | 3 | 21.4 | 76.6 | 39 |
| 1042 | RF00469 HCV SLIV | 15 | 20.7 | 3 | 21.3 | 86.3 | 110 |
| 1043 | RF00047 mir-2 | 21 | 20.0 | 4 | 20.4 | 66.6 | 56 |
| 1044 | RF00550 HepE CRE | 41 | 17.1 | 7 | 16.7 | 84.2 | 46 |
| 1045 | RF00535 snoMe28S-Am982 | 3 | 16.7 | 1 | 16.3 | 76.2 | 13 |
| 1046 | RF00104 mir-10 | 24 | 16.7 | 4 | 16.6 | 68.1 | 36 |
| 1047 | RF00736 mir-320 | 19 | 16.3 | 3 | 17.1 | 68.0 | 55 |
| 1048 | RF00654 mir-216 | 18 | 16.1 | 3 | 15.8 | 61.5 | 33 |
| 1049 | RF00134 snoZ196 | 7 | 15.7 | 1 | 16.7 | 68.4 | 22 |
| 1050 | RF00490 S-element | 22 | 15.4 | 3 | 15.7 | 75.8 | 29 |
| 1051 | RF02027 MIR2907 | 16 | 15.0 | 2 | 15.6 | 76.9 | 52 |
| 1052 | RF02002 mir-720 | 26 | 15.0 | 4 | 15.1 | 81.2 | 35 |
| 1053 | RF02510 PYLIS 3 | 8 | 15.0 | 1 | 16.5 | 63.0 | 23 |
| 1054 | RF00651 mir-221 | 21 | 14.8 | 3 | 14.7 | 73.9 | 47 |
| 1055 | RF00041 Entero OriR | 35 | 14.0 | 5 | 14.1 | 88.0 | 60 |
| 1056 | RF01803 GABA3 | 21 | 13.8 | 3 | 14.0 | 84.5 | 52 |
| 1057 | RF01518 pRNA | 22 | 13.6 | 3 | 14.0 | 57.2 | 23 |
| 1058 | RF00665 mir-290 | 25 | 13.6 | 3 | 13.7 | 65.9 | 27 |
| 1059 | RF00424 SCARNA16 | 54 | 13.0 | 7 | 14.3 | 75.9 | 37 |
| 1060 | RF00451 mir-395 | 30 | 12.7 | 4 | 12.9 | 65.0 | 25 |
| 1061 | RF00679 mir-210 | 27 | 12.6 | 3 | 12.4 | 61.6 | 26 |
| 1062 | RF00670 mir-105 | 29 | 12.4 | 4 | 13.3 | 67.3 | 20 |
| 1063 | RF00357 snoR44 J54 | 5 | 12.0 | 1 | 11.2 | 72.1 | 29 |
| 1064 | RF02447 SpR19 sRNA | 30 | 12.0 | 4 | 13.0 | 74.8 | 23 |
| 1065 | RF01982 PYLIS 1 | 15 | 11.3 | 2 | 10.9 | 71.7 | 20 |
| 1066 | RF00639 mir-515 | 19 | 11.1 | 2 | 12.4 | 80.2 | 40 |
| 1067 | RF02516 mir-393 | 29 | 11.0 | 3 | 10.9 | 63.8 | 27 |
| 1068 | RF02031 tpke11 | 16 | 10.6 | 2 | 10.3 | 68.9 | 28 |
| 1069 | RF00691 mir-146 | 17 | 10.6 | 2 | 10.4 | 63.1 | 33 |
| 1070 | RF00034 RprA | 18 | 10.6 | 2 | 10.4 | 66.8 | 13 |
| 1071 | RF00446 mir-133 | 20 | 10.5 | 2 | 10.7 | 67.6 | 46 |

Table S2: **Rfam RNA families with sufficient power but no covariations.** List of 40 Rfam (v14.1) RNA families with more than 10% power but no covariations, ranked by decreasing power. The expected number of basepair to covary is calculated using R-scape with E-value $< 0.05$.
