## Supplementary material for "Estimating the power of sequence covariation for detecting conserved RNA structure": Table S3

| Structural Alignment | # of seqs | avg % identity | avg seq length | proposed pairs | expected cov pairs | observed cov pairs |
|---|---|---|---|---|---|---|
| **HOTAIR RNA** [9] | | | | | | |
| HOTAIR_Pyle15 D1 | 37 | 74 | 450 | 149 | **58.4** | 0 |
| HOTAIR_Pyle15 D2 | 31 | 74 | 505 | 134 | **23.0** | 0 |
| HOTAIR_Pyle15 D3 | 34 | 68 | 407 | 125 | **41.4** | 0 |
| HOTAIR_Pyle15 D4 | 31 | 69 | 601 | 165 | **38.6** | 0 |
| **Steroid Receptor Activator RNA** [10] | | | | | | |
| SRA_Sanbonmatsu12 | 76 | 78 | 764 | 234 | **49.7** | 0 |
| **Xist RNA - Repeat A** | | | | | | |
| Xist_Branlant12 XIST_A.S0 [11] | 10 | 81 | 438 | 53 | 0.0 | 0 |
| Xist_Branlant12 XIST_A.S1 [11] | 10 | 81 | 439 | 90 | 0.5 | 0 |
| Xist_Branlant12 XIST_A.S2 [11] | 10 | 81 | 439 | 72 | 0.4 | 0 |
| Xist_Branlant12 XIST_A.S3 [11] | 10 | 81 | 439 | 83 | 0.4 | 0 |
| Xist_Simon15 Fig.5 [12] | 13 | 75 | 423 | 99 | 2.5 | 0 |
| **Xist RepA lncRNA - Repeat A + Repeat F** | | | | | | |
| Xist_Pyle17 [18] | 57 | 68 | 1,320 | 334 | **108.6** | 1 |
| Xist_Rivas19 h1 | 65 | 68 | 1,076 | 254 | **100.7** | 0 |
| **Xist RNA - other conserved regions** | | | | | | |
| Xist_Rivas19 h2 (exon1, repC frag) | 52 | 68 | 197 | 41 | **25.1** | 0 |
| Xist_Rivas19 h3 (exon1) | 55 | 68 | 257 | 53 | **16.2** | 0 |
| Xist_Rivas19 h4 (exon1) | 61 | 60 | 254 | 64 | **28.6** | 0 |
| Xist_Rivas19 h5 (exon3) | 55 | 70 | 275 | 63 | **18.4** | 0 |
| Xist_Rivas19 h6 (exon4) | 56 | 79 | 360 | 112 | **20.7** | 0 |
| Xist_Rivas19 h7 (exon5) | 73 | 61 | 354 | 123 | **50.4** | 0 |
| Xist_Rivas19 h8 (exon6, repE frag) | 60 | 66 | 323 | 89 | **34.3** | 0 |
| Xist_Rivas19 h9 (exon6) | 47 | 70 | 129 | 32 | **7.2** | 0 |
| Xist_Rivas19 h10 (exon6) | 59 | 73 | 262 | 61 | **15.9** | 0 |
| Xist_Rivas19 h1-h10 (concatenated) | 32 | 72 | 3528 | 980 | **141.3** | 0 |
| **COOLAIR antisense RNA** | | | | | | |
| COOLAIR_Sanbonmatsu16 [21] | 6 | 78 | 394 | 114 | 0.9 | 6 |
| COOLAIR_Rivas19 | 6 | 80 | 547 | 166 | 0.8 | 1 |

Table S3: **Power of covariation versus observed covariations for several lncRNAs.** No observed covariations for an adequate expected number of them constitutes negative evidence for a conserved RNA structure. Expected covariations larger than 10% the number of proposed basepairs representing alignments with sufficient power are displayed in bold. The proposed pairs are the annotated basepairs in the source alignments. The label "Rivas19" indicates alignments produced in this work using nhmmer[19] (for Xist) or Infernal[22] (for COOLAIR). The structures for "Rivas19" alignments are produced with R-scape. The expected and observed covarying pairs were obtained with R-scape v1.2.3 using default parameters (E-value $< 0.05$). The expected number of covarying pairs is the sum of the power of the basepairs in the proposed structures (Methods, Eq. 2). The average pairwise percentage identity is calculated after removing positions with more than 50% gaps. All alignments are included in the supplemental materials.
